# Cancer Prevalence Across Vertebrates

**DOI:** 10.1101/2023.02.15.527881

**Authors:** Zachary T. Compton, Walker Mellon, Valerie Harris, Shawn Rupp, Diego Mallo, Stefania E. Kapsetaki, Mallory Wilmot, Ryan Kennington, Kathleen Noble, Cristina Baciu, Lucia Ramirez, Ashley Peraza, Brian Martins, Sushil Sudhakar, Selin Aksoy, Gabriela Furukawa, Orsolya Vincze, Mathieu Giraudeau, Elizabeth G. Duke, Simon Spiro, Edmund Flach, Hannah Davidson, Christopher Li, Ashley Zehnder, Trevor A. Graham, Brigid Troan, Tara M. Harrison, Marc Tollis, Joshua D. Schiffman, Athena Aktipis, Lisa M. Abegglen, Carlo C. Maley, Amy M. Boddy

## Abstract

Cancer is pervasive across multicellular species, but what explains differences in cancer prevalence across species? Using 16,049 necropsy records for 292 species spanning three clades (amphibians, sauropsids and mammals) we found that neoplasia and malignancy prevalence increases with adult weight (contrary to Peto’s Paradox) and somatic mutation rate, but decreases with gestation time. Evolution of cancer susceptibility appears to have undergone sudden shifts followed by stabilizing selection. Outliers for neoplasia prevalence include the common porpoise (<1.3%), the Rodrigues fruit bat (<1.6%) the black-footed penguin (<0.4%), ferrets (63%) and opossums (35%). Discovering why some species have particularly high or low levels of cancer may lead to a better understanding of cancer syndromes and novel strategies for the management and prevention of cancer.

**Statement of Significance:** Evolution has discovered mechanisms for suppressing cancer in a wide variety of species. By analyzing veterinary necropsy records we can identify species with exceptionally high or low cancer prevalence. Discovering the mechanisms of cancer susceptibility and resistance may help improve cancer prevention and explain cancer syndromes.

## Introduction

Cancer is a ubiquitous problem for multicellular species ^1^ and a leading cause of death in humans^2^. Every multicellular body is a cooperative cellular system, with cells suppressing replication^3^, dividing labor ^4^, sharing resources ^5^, regulating cell death ^6^ and taking care of the extracellular environment ^1^. However, cooperative systems are susceptible to cheaters, which emerge as cancers in multicellular organisms ^7^. Because cancer cells can outcompete normal cells with respect to replication, survival, resource use and other cellular behaviors, natural selection within the body can favor cancer cells via somatic evolution.

Cancer has been a strong selective pressure on multicellular organisms and mechanisms for cancer suppression likely co-evolved along with the evolution of multicellularity ^8,9^. Despite this persistent selective pressure of cancer, species vary in their investment in cancer defenses across the tree of life. Sir Richard Peto predicted in 1977 that the risk of cancer should scale with the number of cells in an organism and the length of its lifespan ^10^. This prediction is based on the fact that tumors evolve from single cells, partially due to the accumulation of somatic mutations over time ^10^. His observation that cancer risk does not appear to increase with increases in body mass and longevity across species ^10^, a phenomenon known as ‘Peto’s paradox’, launched the field of comparative oncology ^11^.

Early work in comparative oncology found that birds, and to a lesser extent reptiles, develop fewer neoplasms than mammals ^12–14^. While single case studies have been reported ^15^, it has been difficult to estimate true neoplasia prevalence in these taxa. In 2015, we published neoplasia prevalence estimates in 37 mammal species and reported support for Peto’s Paradox, that is, bigger, longer-lived species do not get more cancer ^16^. Follow up studies have supported Peto’s Paradox and demonstrated the ubiquity of cancer across mammals ^17,18^. A recent study of 131 species of mammals with at least 5 necropsies per species, found no relationship between body mass and neoplasia prevalence, though they did observe an increase in both neoplasia and malignancy prevalence in 25 species of amphibians and 71 species of squamates^19^. Our study of 110,148 animals across 191 species of mammals found no relationship between body mass and death due to cancer^18^.

The extensive variation in cancer risk across vertebrates provides a unique opportunity to identify species with exceptional cancer resistance that can lead to new discoveries of cancer resistance mechanisms outside the traditional human and murine studies. Additionally, the discovery of cancer vulnerable species could lead to new insights into cancer syndromes as well as provide spontaneous ‘natural’ animal models of disease that can help us gain a better understanding of various types of cancer and their treatments. Here we present a large, curated database of tetrapod veterinary necropsy records, including 16,049 individual animals across 292 species of animals, encompassing reptiles, birds, amphibians, and mammals. Because necropsies typically are diagnosed with “neoplasia” which includes both benign and malignant tumors, we developed a terminology dictionary to distinguish benign from malignant neoplasms in the necropsy reports. We calculate and analyze both neoplasia prevalence as well as malignancy (cancer) prevalence. Only a subset of benign neoplasms evolve into cancers over a lifetime, so neoplasia prevalence is always greater than or equal to malignancy prevalence. We also tested for age bias in the animals that died with neoplasias or cancers. To test for general mechanisms of cancer suppression, we tested for an association with DNA damage response or somatic mutation rate and neoplasia.

## Results

### Variation in Cancer Prevalence Across Clades

We found evidence of neoplastic disease in necropsies across all analyzed taxonomic clades (Fig.1). For all vertebrates, the median prevalence of neoplasia at death was 4.89% (range = 0% – 62.86%) and median malignancy prevalence was 3.20% (range=0% – 40.95%). Mammals had the most neoplasia at death with a median of 12% (range = 0% - 63%) and median malignancy prevalence of 7% (range=0% – 41%). Sauropsids followed with a median neoplasia prevalence of 4% (range = 0% – 39%) and median malignancy prevalence of 1.6% (range=0% – 35%). Lastly, amphibians had a median neoplasia prevalence of 1.2% (range = 0% – 46%) and median malignancy prevalence of 0% (range=0% – 33%). The ranking of prevalence by clade is consistent with previous studies^12,14^. In Fig. 2 we have shown Aves and Squamata separately, but because reptiles are not a monophyletic clade, we have grouped reptiles with birds in the Sauropsida clade for the purposes of further analyses. Despite a lower mean prevalence for both benign and malignant tumors, sauropsids and amphibians show a wide range of neoplastic disease burden across species. There is a small but highly statistically significant correlation between the prevalence of benign neoplasms and the prevalence of malignant neoplasms across species (*r*=0.34, *p*<0.0001, Fig. S57). Supplementary Tables ST1 and ST2 list the species with the highest and lowest neoplasia and malignancy prevalences, as well as the proportion of neoplasms that are malignant. Among the vertebrates with the highest prevalence of neoplasia, 63% of ferrets (*Mustela putorius*) died with a neoplasm (45% of which was lymphoma), 56% of opossums (*Didelphis marsupialis*) died with a neoplasm (46% of which was in the lung), and 45% of four-toed hedgehogs (*Atelerix albiventris*) died with a neoplasm (42% of which was in the alimentary tract) (ST8).

**Fig. 1.**
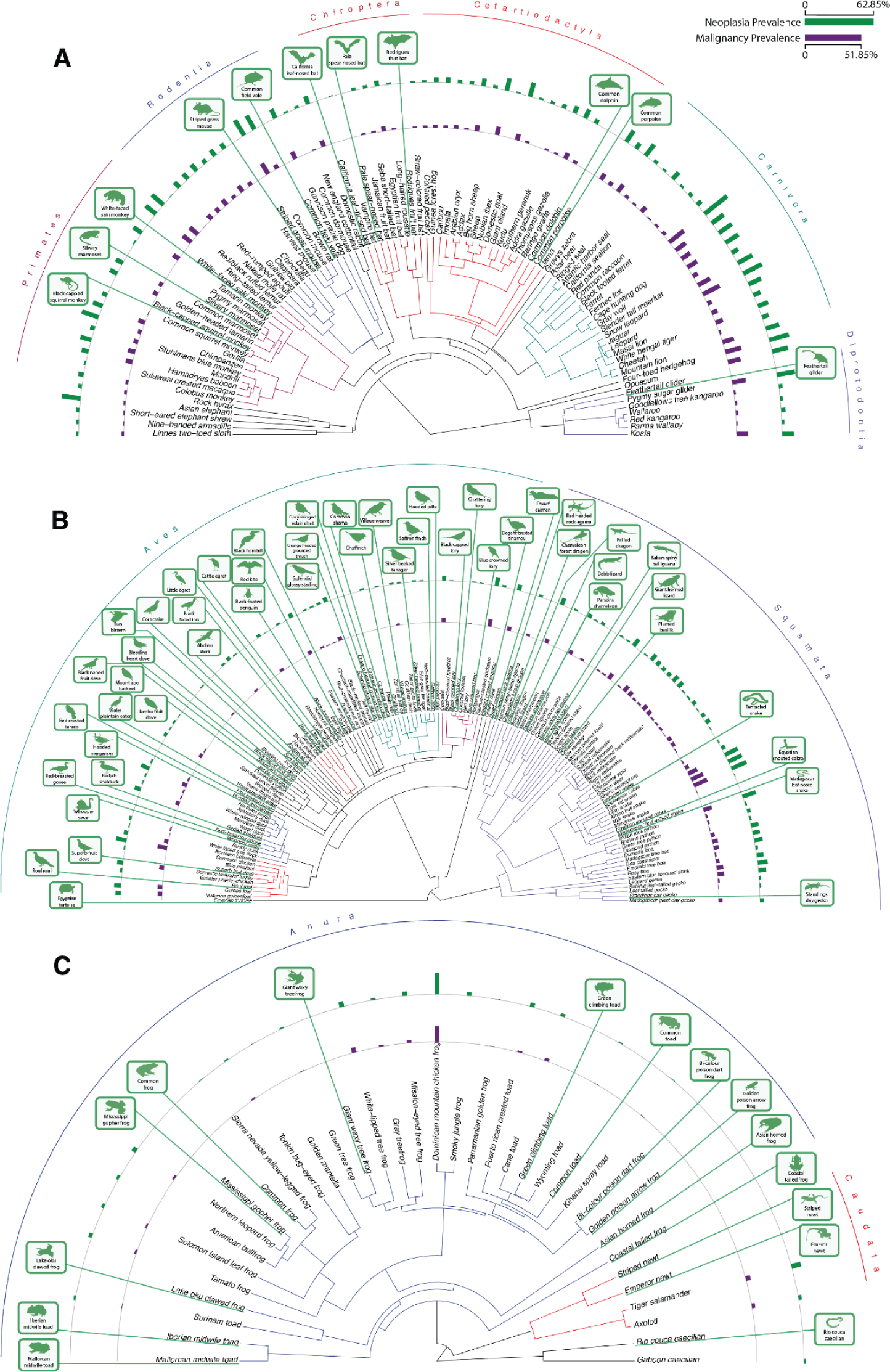
Neoplasia and malignancy prevalence across (**A**) mammals, (**B**) sauropsids (Aves, Squamata, Testudines and Crocodylia) and (**C**) amphibians. Silhouetted species indicated that zero neoplasms were reported in our data.

**Fig. 2.**
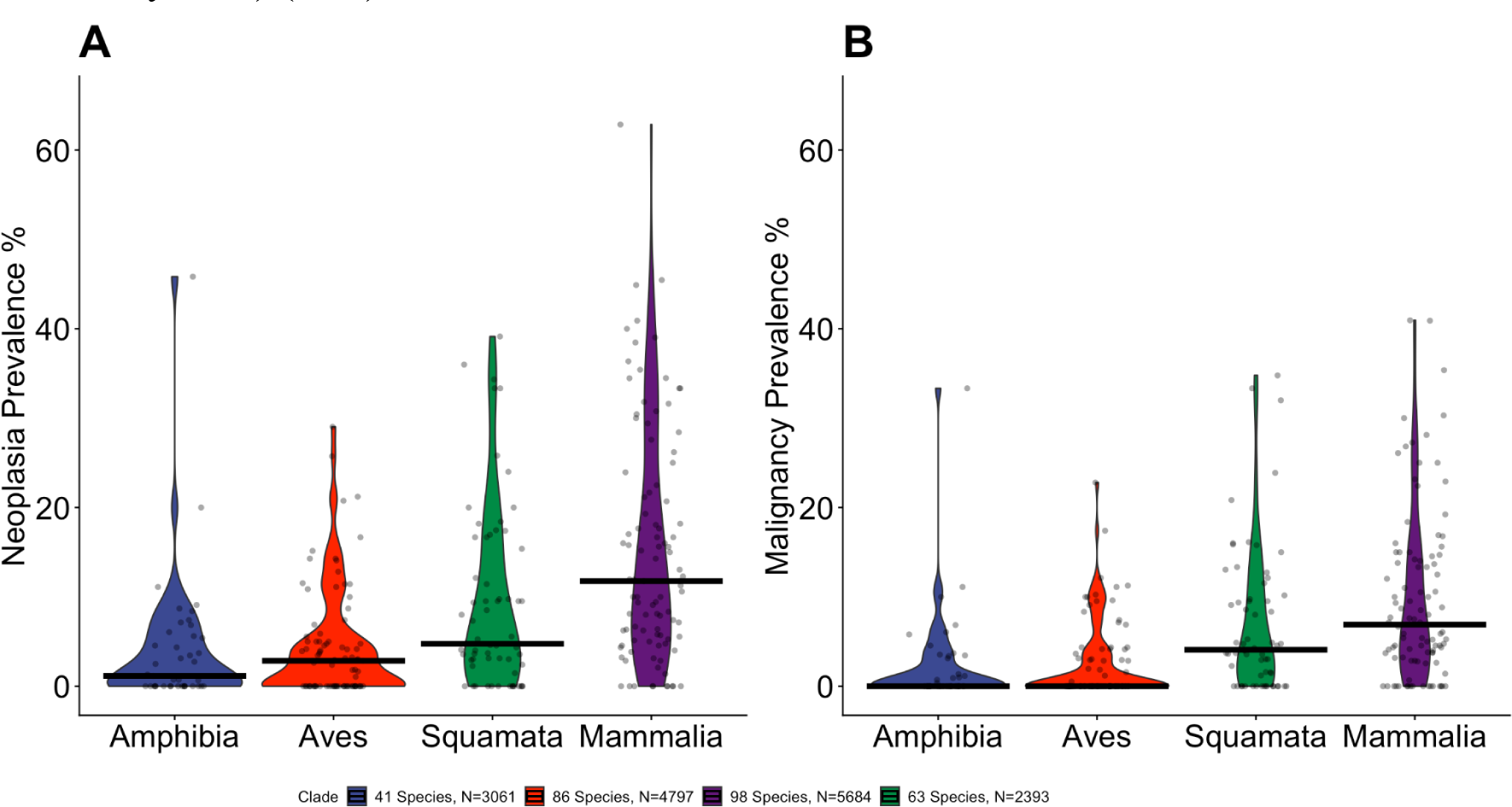
Distributions of **A.** neoplasia (Kruskal-Wallace test: p = 2.906 x 10^-12^) and **B.** malignancy (Kruskal-Wallace test: p = 6.519 x 10^-11^) prevalences are different across four clades, Amphibia, Mammalia, Aves and Squamata. Dots show the estimated species neoplasia prevalence and bars show the median for the clade. Neoplasia and malignancy prevalence for species were calculated by the proportion of the reported lesions among the total number of necropsies for that species.

### Life History Analyses of Neoplasia Prevalence

Evolutionary life history theory provides a framework for understanding the tradeoffs governing species’ survival and reproduction ^20,21^. Life history theory can be used to explain how species level traits shape organismal cancer risk based on trade-offs between investment in somatic maintenance (e.g., cancer suppression) and reproduction or growth. Several smaller studies have shown that specific life history traits can serve as prognostic indicators of neoplasia prevalence in animals managed under human care ^17,22^. We used a phylogenetic regression to test for relationships between life history factors and neoplasia or malignancy prevalence, controlling for phylogenetic relatedness, and weighting species data points by the number of necropsies in our dataset. In contrast to two of three previous studies^16–19^, we found an increase in neoplasia prevalence with increases in body mass (2.1% neoplasia per Log_10_(g), *p* = 0.007; 0.65% malignancy per Log_10_g, *p* = 0.287) and maximum longevity (0.01% neoplasia per Log_10_ months, *p* = 0.02; 0.005% malignancy per Log_10_ months, *p* = 0.276), not supporting Peto’s Paradox (Fig. 3). Animals with longer gestation times also get fewer malignancies (N=151 species, −5.3% neoplasia per Log_10_ months, *p*=0.1; −5.65% malignancies per Log_10_ months, *p* = 0.02; Fig. 3C).

**Fig. 3.**
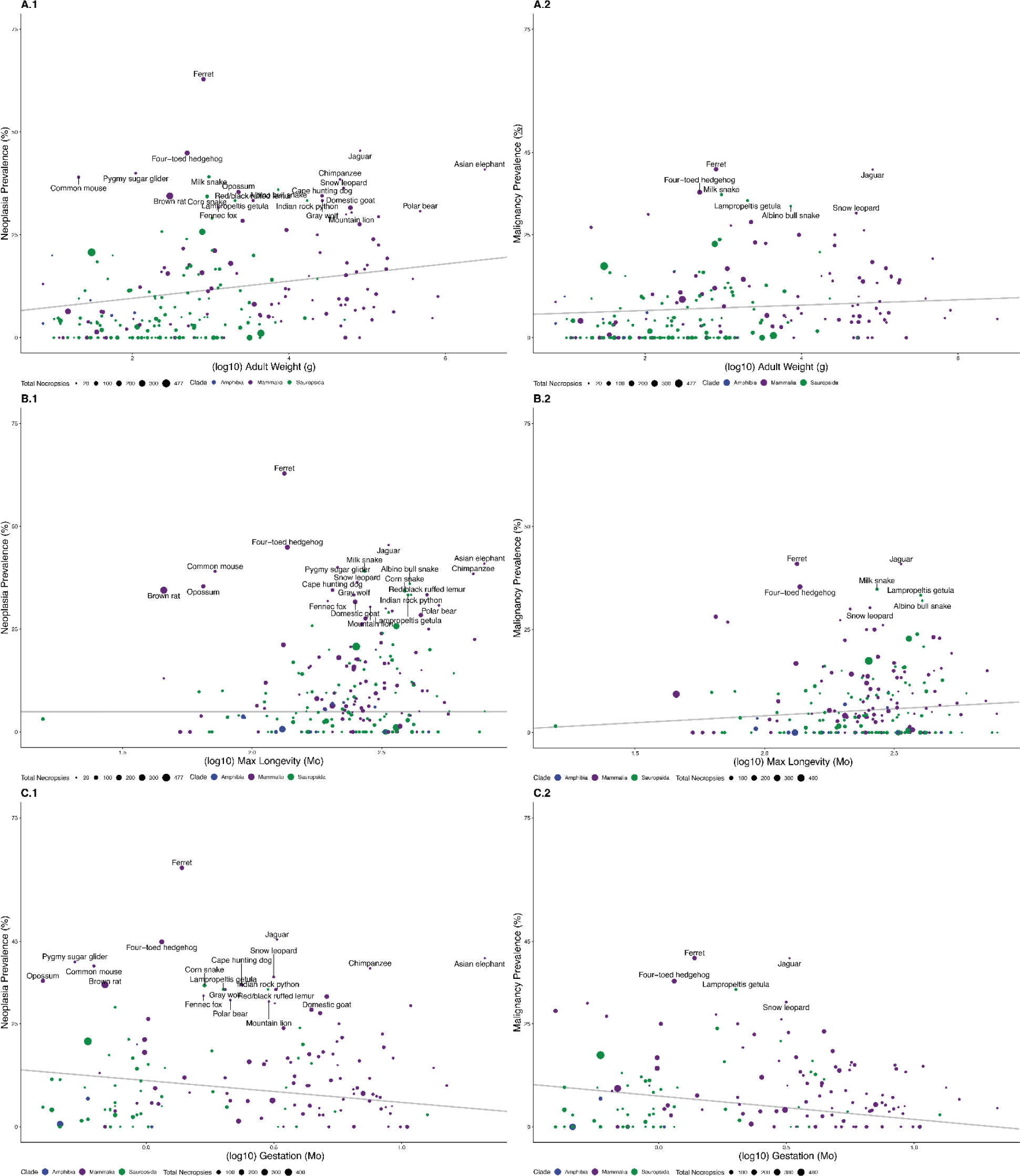
Significant life history factors associated with neoplasia and malignancy prevalence. **A.1** Larger organisms have a higher neoplasia prevalence than smaller organisms (2.1% neoplasia per Log_10_g adult body mass, *p* = 0.007, R^2^ = 0.26, λ = 0.46). **A.2** Larger organisms have a higher malignancy prevalence than smaller organisms (0.65% malignancies per Log_10_g adult body mass, *p* = 0.29, R^2^ = 0.28, λ = 0.61). **B.1** Longer-lived organisms have more neoplasia (0.01% neoplasia per Log_10_ month lifespan, p = 0.02, R^2^ = 0.19, λ = 0.34). **B.2** Longer-lived organisms also have more malignancies (3.3% malignancies per Log_10_ month lifespan, p = 0.17, R^2^ = 0.21, λ = 0.47). **C.1** Organisms with longer gestation times have a lower neoplasia prevalence (−5.30% neoplasia per Log_10_ months, *p* = 0.10, R^2^ = 0.11, λ = 0.34). **C.2** Organisms with longer gestation times have a lower malignancy prevalence (−5.65% malignancies per Log_10_ months, *p* = 0.02, R^2^= 0.16, λ = 0.41). When controlling for adult body mass, organisms with longer gestation times also have fewer neoplasias at death (−15.8% neoplasia per Log_10_ months gestation, *p* = 0.0002).

A multivariate model containing all significant predictors of neoplasia or malignancy (adult weight, maximum longevity, and gestation time) shows that both adult body weight (2.9% neoplasia per Log_10_g, *p* = 0.01) and gestation time (−18.6% neoplasia per Log_10_month, *p* = 0.0001) provide independent information for estimating neoplasia prevalence, but not longevity (*p=*0.12). Because gestation time and adult weight are correlated (r = 0.50, *p* < 10^-10^), but have the opposite relationship to neoplasia and malignancy prevalence, we tested the two-variable model and found that when controlling for adult weight (3.8% neoplasia per Log_10_(g), *p* = 0.0006), gestation time is also a significant predictor of neoplasia prevalence (−15.8% neoplasia per Log_10_ months, *p* = 0.0002; R^2^ = 0.20), and vice versa. When controlling for gestation time, adult weight predicts malignancy prevalence, and when controlling for adult weight, gestation time also predicts malignancy prevalence (1.9% malignancies per Log_10_(g), *p* = 0.02; −12% malignancies per Log_10_ months of gestation, *p* = 0.0002). Adult weight and gestation time were still statistically significant (adjusted *p*<0.05) predictors of neoplasia and malignancy prevalence after a 10% false discovery rate correction for multiple testing.

We validated our results by implementing phylogenetic binomial regressions^19^ on records that include the age of the animal. All the above statistically significant relationships are still significant at p < 0.05 level using binomial regressions. In addition, binomial regressions reveal a significant positive relationship in mammals between body mass and neoplasia (OR=9.5% neoplasia per ln(g), *p*=0.012 but *p*=0.06 with PGLS) as well as malignancy prevalence (OR=1.5% malignancy per ln(g), *p*=0.015 but *p*=0.458 with PGLS; ST4). Within mammals, analyses of adult body mass and gestation time together showed the same relationships as all vertebrates (OR=-18% neoplasia and −15% malignancy per Log_10_ months of gestation, *p*=0.036 and p<10^-10^; 3.4% neoplasia and 2.6% malignancy per Log_10_(g) of adult body mass, *p*=0.017 and 0.0066 respectively with PGLS). These relationships were even stronger with binomial regressions (OR=-44% neoplasia and −48% malignancy per Log_10_ months of gestation, p=0.0009 and 0.001; OR=14% neoplasias and 14% malignancy per Log_10_(g) adult body mass, *p*=0.002 and p=0.0035).

We found no evidence of a relationship between litter or clutch size and neoplasia prevalence (Figs. S1, S2). However, when we restrict the analysis to mammals, litter size is positively associated with both neoplasia and malignancy prevalence (neoplasia: *p* = 0.02, R^2^=0.07; malignancy: *p* = 0.03, R^2^ = 0.11; Suppl. Figs. 21 & 22), supporting our earlier analysis of 37 mammals from the San Diego Zoo^17^. We also found that time to sexual maturity, growth rate and basal metabolic rates (which were only available for mammals) were not significant predictors of neoplasia or malignancy prevalence (Figs. S3, S4, S9, S10, S11, S12, S15, and S16). In addition to calculating the prevalence of neoplasms and malignancies, we also calculated the proportion of neoplasms that were malignant, which is a measure of the likelihood that a benign neoplasm transforms into a malignant one. We found no statistically significant relationships between any of those life history factors and the proportion of neoplasms that were malignant (Figs. S62-S74).

**Fig. 4.**
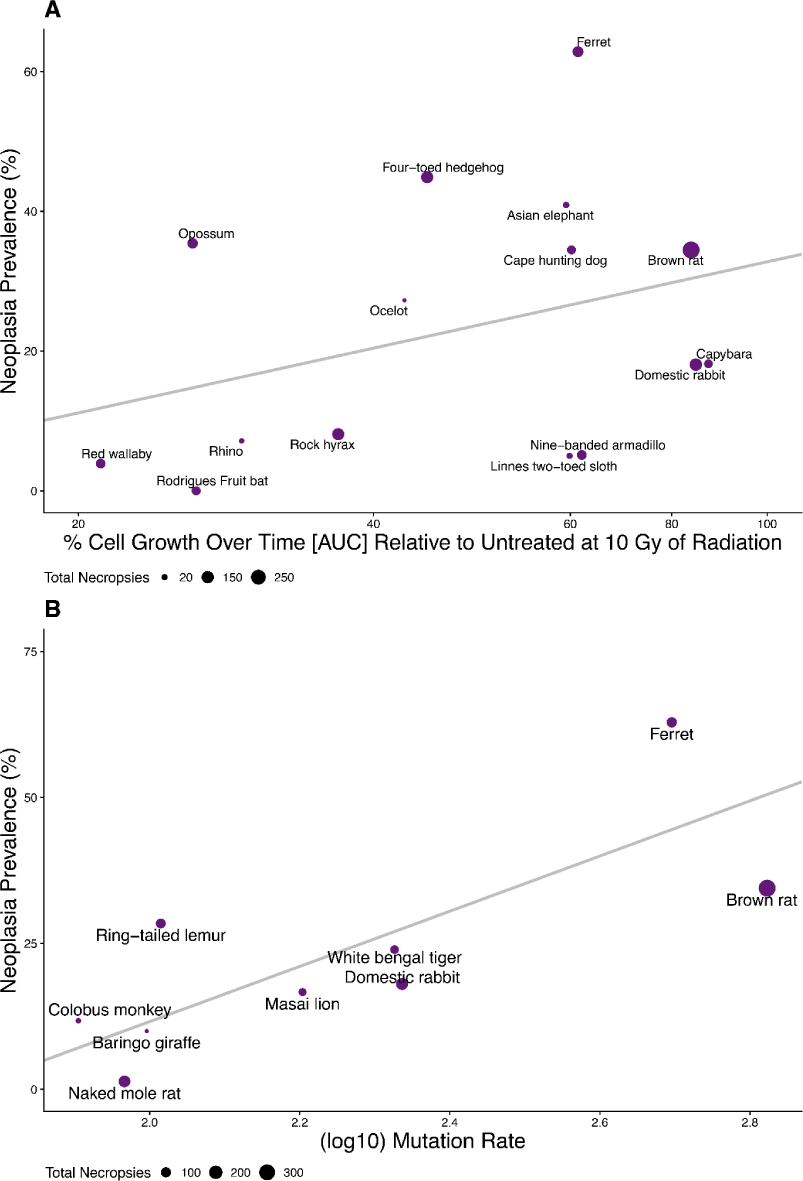
A. Cell cycle arrest as measured by % Cell Growth Over Time [AUC] Relative to Untreated at 10 Gy of Radiation (plotted and analyzed on a Log_10_ scale) as a predictor of neoplasia prevalence in species’ fibroblast cell lines. (30.89% neoplasia per Log_10_ Cell Count Area Change, *p* = 0.22, R^2^ = 0.12, λ = 6.6 x 10^-5^) **B.** Log_10_ Mean Mutation Rate as a predictor of neoplasia prevalence (47.26% per single base substitution per genome per year, *p* = 0.0059, R^2^ = 0.63, λ = 1.00).

### DNA damage response and somatic mutation rates

DNA damage response is an important anti-cancer mechanism. Accurate repair of damaged DNA or clearance of damaged cells through apoptosis ensures that mutations capable of driving tumorigenesis do persist or accumulate. For example, we previously reported that increased apoptosis in response to DNA damage in elephant cells was associated with low cancer mortality^16^. To determine if DNA damage response is a general mechanism of cancer defense across species, we measured the ability of primary fibroblasts from 15 species to respond to DNA damage (Fig. 4A; Suppl. Figs. 41-56). DNA damage response was assessed by measuring cell cycle arrest, which halts cell division until damage is repaired, and apoptosis, which kills cells that cannot be repaired. ^16,23^. We hypothesized that animals with less neoplasia and malignancy would respond more robustly to DNA damage. Cell cycle arrest in response to increasing doses of ionizing radiation and apoptosis in response to increasing doses of a chemotherapeutic drug (doxorubicin) was measured over time. We observed variability in DNA damage response across the species that we tested, and this variability was not associated with neoplasia or malignancy (Figure 4A and Figures S41 - 56), although cell cycle arrest in response to 10Gy radiation trended toward an association with neoplasia prevalence (Figure 4A).

Our mechanism based cellular assays tested the initial response to DNA damage; cell cycle arrest and apoptosis. Perhaps the most important role of DNA damage response is the suppression of somatic mutation accumulation, because mutations drive tumor formation and cancer progression^24–26^. Therefore, we hypothesized that lower somatic mutation rates across species would correlate with lower cancer prevalence. We obtained previously reported somatic mutation rates from 9 species in our dataset and tested for a correlation with neoplasia prevalence ^27^. We discovered a positive association between somatic mutation rates and neoplasia prevalence, and that species with fewer somatic mutations also developed less neoplasia (Figure 4B, pgls *p* = 0.0059). We also observed a similar trend for somatic mutation rates and malignancy prevalence but it was not statistically significant (S50, pgls *p* = 0.087).

### Age at Death with Cancer in Animals

Age is the single biggest risk factor for the development of cancer in humans ^28^. Most mechanisms of somatic maintenance, including immune cell surveillance, DNA damage response, and telomere shortening, decrease in efficacy as we age ^29–33^. To test if observed neoplasms in animals under human care may be due to the animals living beyond their natural lifespans, we plotted the age of the animals with neoplasia at death, compared to the animals that died without neoplasias, scaled by their average lifespan (Fig. 5). The vast majority of animal deaths with neoplasia diagnoses occur before the average lifespan in most animals. Only amphibians seem to be developing more neoplasms as they live past their normal lifespan under human care (Fig. 5C). The distribution of tumor diagnoses across lifespan in these three clades also demonstrates that cancer is not limited to a disease solely of extended lifespan, and in sauropsids, neoplasia is not particularly a disease of old age (Fig. 5B).

**Fig. 5.**
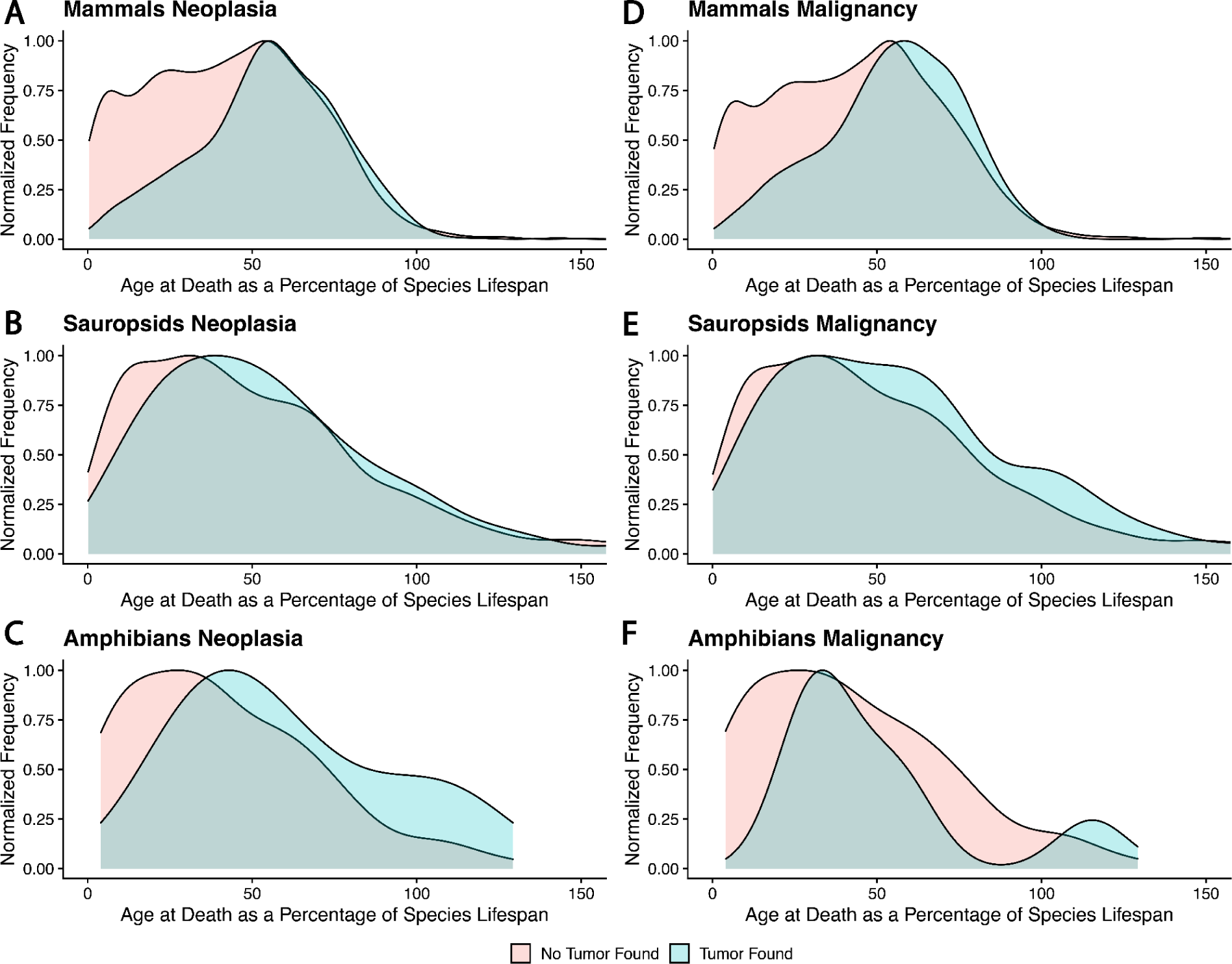
The density distribution of ages at death in animals with neoplasia versus non-neoplasia, adjusted for each species’ lifespan as specified in PanTHERIA. While the distributions of ages at death are different between necropsies showing neoplasia versus those that don’t (Two Sample Kolmogorov-Smirnov Test: Mammals: D=0.11, p =1.81 x 10^-6^; Sauropsids: D= 0.18, p = 4.48 x 10^-8^; Amphibians: D=0.5, p = 0.011), we found few neoplasias that could be explained by an organism living an extraordinarily long time in captivity, except in amphibians.

## Phenotypic Models of Cancer Risk

### Evolution of cancer suppression and susceptibility

Comparative phylogenetics provides a wealth of computational tools to model species’ trait evolution across a phylogeny ^34^. To explore how cancer susceptibility evolved across the tree of life (Fig. 6), we fit three of the most common phenotype evolution models (Ornstein-Uhlenbeck, Brownian Motion, and Early Burst) to neoplasia prevalence as a continuous trait. We found that a model of stabilizing selection on neoplasia prevalence (Ornstein-Uhlenbeck) fits the distribution of neoplasia prevalence the best (ST7). Malignancy prevalence evolution is also best explained by the Ornstein-Uhlenbeck model of sudden shifts followed by stasis in the phenotype.

**Fig. 6.**
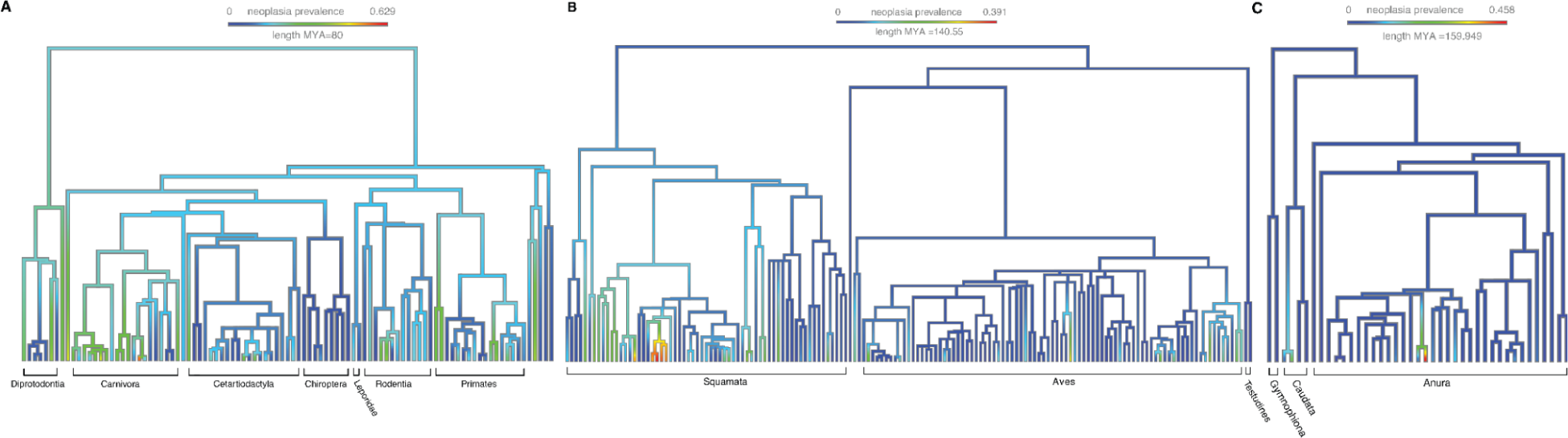
Cladogram depiction of cancer incidence within **A.** Mammals, **B.** Sauropsids, and **C.** Amphibians. Cladograms with the species labels at each tip can be found in Suppl. Fig. 63. Heat map coloration indicates relative prevalence of cancer within each branch, illustrating the diversity of neoplastic disease amongst closely related species. The scale is the same for each panel so that the differences between the clades are apparent.

## Discussion

We estimated cancer prevalence across a wide range of tetrapod species that includes mammals, amphibians, reptiles and birds. Importantly, and contrary to previous studies, our analyses highlight limitations to Peto’s paradox, by showing that large animals do tend to get somewhat more neoplasms, and malignancies, compared with smaller animals. This is particularly apparent when we control for the fact that animals with longer gestation times tend to get both fewer neoplasms and fewer cancers. However, large animals only get slightly more cancer than small animals. Whether or not they get as much cancer as one would expect from their body size and longevity depends on the model one uses to predict cancer prevalence as a function of body size ^35–37^. We hypothesize that animals with a longer gestation time than predicted based on their body size, may be investing more resources towards cell cycle regulation and differentiation, and thereby reducing their vulnerability to cancer, compared to animals with shorter than expected gestation times. They may also prevent “jackpot” somatic mutations in gestation, which expand to large clones through the process of development and can significantly contribute to the risk of progressing to cancer later in life^38,39^. Lastly, it may be the case that hormonal exposures during pregnancy help protect against cancer in the mothers, as we see for some forms of cancer in humans^40,41^. These hypotheses could be tested by examining the relationship between gestation time and cancer in females versus males, across species.

Cancer prevalence across species varies greatly. Here we have used a large collection of species, and expanded our analyses beyond mammals^16–18^, to test for patterns in cancer prevalence. We only include species with at least 20 necropsies (median 35), compared with 10 individuals per species in our original study^16^, and weighted species more in our regression analyses if their cancer prevalence estimate is more accurate because it is based on more necropsies. When Bulls et al. restricted their data to species with at least 5 necropsies, they also found a positive relationship between body size with both neoplasia and malignancy prevalence in amphibians and squamates. When Bulls et al. reanalyzed *cancer mortality* data from Vincze et al^18^ across 189 species, they found that larger animals were more likely to die due to cancer than smaller mammals (3% cancer mortality per ln(g), *p*<0.001). Using a binomial regression, both our data and Bulls et al. found a negative relationship between neoplasia and longevity, but this relationship is non-significant in our dataset.

The fact that neoplasia prevalence seems to evolve by sudden shifts followed by stabilizing selection (the Ornstein-Uhlenbeck model of phenotypic evolution) is consistent with life history theory predictions that investment in somatic maintenance should be under selection in specific ecological conditions^20^, rather than drifting neutrally consistent with random Brownian motion. Some of the variation in cancer prevalence may be noise, due to estimating cancer prevalence from tens of individuals. However, much of that variation comes from the vast diversity of species across amphibians, reptiles, birds and mammals. We have explained only a small portion (∼20%) of the variation in species vulnerability and suppression of cancer. There is clearly more to be discovered.

Peto’s Paradox is based on the expectation that large, long-lived animals should get more cancer because they have more cells that divide for a longer amount of time, increasing the likelihood that cancer will arise^10,11^. Although adult body weight is positively correlated with both neoplasia and malignancy prevalence, partially resolving Peto’s paradox, the effect size is larger for neoplasia (2.1% neoplasia per Log_10_g) than for malignancies (1.9% malignancies per Log_10_g), when controlling for gestation time. There may be several explanations for this. The simplest is that malignancies are less common than neoplasias, which include both benign and malignant neoplasms. This reduces the statistical power and the expected size of the effects. However, the blunted relationship between body size and malignancy prevalence may also be due to natural selection acting to evolve mechanisms to suppress malignant transformation. Cancer suppression mechanisms are likely under stronger selection among these larger and longer-lived organisms because it was critical to suppress cancer for longer in order for these organisms to successfully reproduce. Thus, we might expect a relatively constant cancer rate across species with more cancer suppression mechanisms in large, long-lived organisms ^7,16,42–46^, and fewer in small, short-lived organisms.

Further, there are at least four transitions in neoplastic progression that natural selection might alter to increase the survivability of cancer in a species: 1) initiation of a neoplasm, 2) transformation of that neoplasm into malignancy (*i.e.*, invasion through the basement membrane), 3) metastasis, and 4) death caused by the cancer. Our data bear on the first two transitions. Specifically, we quantify the prevalence of neoplasms in a species, the prevalence of malignant neoplasms, and the proportion of neoplasms detected that are malignant. However, the selective pressure of cancer is ultimately due to its effects on mortality, and so quantifying the prevalence of cancer as a cause of death is also important for evolutionary studies of comparative oncology ^18^. The discrepancy between our earlier analysis of a different data set of 191 mammal species that found no relationship between body mass and cancer mortality and our results here may be due to at least three differences: 1) Here we analyze cancer prevalence, not cancer mortality, 2) our current results are based on all vertebrates, not just mammals, and 3) in our previous analyses, we separately analyzed the chance that a species had non-zero cancer mortality and then tested for a relationship between log body mass and cancer mortality in species with non-zero cancer mortality, so that the species with very low cancer mortality (zero in that sample) were excluded from the second analysis.

We included cross-species functional assays highlighted in Fig. 4A to test for general mechanisms of cancer defense across species. Our results demonstrate variation in the fibroblast response to radiation and chemotherapy induced DNA damage. Here we provide the first test of association between DNA damage response and cancer prevalence across species. While response to DNA damage was not a significant predictor of neoplasia or malignancy prevalence after 4 or 10 grays radiation, the trend follows our hypothesis that sensitivity to DNA damage may be one mechanism of cancer suppression ^16^. As we continue to accumulate DNA damage response data across more species and more individuals from each species, future studies may reveal a clear relationship. On the other hand, the variation in DNA damage response may suggest that different species evolved unique mechanisms of cancer suppression and in some cases, they do not require enhanced DNA damage response. For example, some species may rely more on enhanced mechanisms of immune surveillance, thereby eliminating a relationship between DNA damage response measurements and neoplasia or malignancy prevalence.

However, related to DNA damage response, we did find a connection between somatic mutation rates^27^ and neoplasia prevalence (Figure 4B). This result was predicted due to the role of somatic mutations in tumor initiation^47–49^ and progression^25,26^. Even with only 9 species in our analysis, a strong relationship between somatic mutation rate and neoplasia prevalence was apparent. This relationship should be validated with the addition of both more somatic mutation rate data and more cancer prevalence data. Furthermore, future research should determine what mechanisms evolved in some species to suppress the accumulation of somatic mutations. Potential mechanisms include prevention of DNA damage, accurate repair of DNA damage, death of damaged cells, or use of high fidelity DNA polymerases^16,50–52^.

### Insights from comparative oncology for human cancers

Species with a high prevalence of particular cancers may help to generate targeted studies to elucidate the biological basis of those cancers, help draw informative parallels to particular cancer syndromes in humans, and serve as more realistic models for studying those cancers^53^. For instance 46% of the malignant tumors diagnosed in the opossum (*Didelphis marsupialis*) were lung adenocarcinomas (ST8), which is a leading cause of human cancer mortality in the United States ^54^. 45% of the cancers in four-toed hedgehogs (*Atelerix albiventris*) were gastrointestinal cancers. They may hold insights for colorectal cancer, the third leading cause of cancer mortality in the US^55^. 42% of cancers in ferrets (*Mustela putorius*) were lymphomas, which may make them a good animal model animal for that disease. These spontaneous cancers may be more similar to human cancers compared to those that develop in genetically engineered mouse models, though that remains to be tested.

There is an exciting opportunity to discover cancer suppression mechanisms in species with few to no observed neoplasms, and in species that seem to prevent benign neoplasms from progressing to malignancy (Fig. 1). For example, the paucity of neoplasms in dolphins and porpoises may result from having large, long-lived cetacean ancestors that were under strong selection to suppress cancer ^42^. Our earlier analysis of cancer gene evolution in cetaceans found evidence of positive selection in a large number of tumor suppressor genes and proto-oncogenes^42^.

We previously found that animals that live longer than would be expected for their body size, like bats, tend to have more copies of tumor suppressor genes ^56^. In support of these observations, we find 9 bats, with an average lifespan of 16 years, have low neoplasia prevalence. We had hoped to discover species that are able to prevent malignant transformation by finding species that get a fair number of benign neoplasms, but few to no malignant neoplasms. The common paradigm in understanding the evolution of cancer suppression emphasizes the importance of protecting against tumor initiation. However, mechanisms that suppress malignant transformation could be similarly important in maintaining an organism’s fitness. Unfortunately, only a few species in our dataset fit that description. The species with the lowest proportion of malignant to benign neoplasms was the common squirrel, with only 12% of their tumors being malignant.

### Challenges for Comparative Oncology

There are a number of potential sources of bias in comparative oncology data. The protection against predation that zoological institutions offer fast life history animals may be extending their lifespan, and thereby unmasking a vulnerability to cancer at an age that they would never attain in the wild. However, Fig. 5 shows that the neoplasms were diagnosed prior to average lifespan in most cases, suggesting the extended lifespan due to managed care is not a large factor in these data. In fact, one surprise was that neoplasms appear in birds and reptiles at relatively young ages, although some birds like chickens are known to be prone to virally induced cancers ^57^.

Our data results from the combination of the intrinsic cancer susceptibility of a species with the effect of the artificial conditions of managed populations, which is sometimes called an evolutionary mismatch ^58^. These animals were generally protected from predators, provided with veterinary care, had different diets and exercise from their wild counterparts, many lived in an urban environment, and interacted with different species and microbes than a free-ranging animal. It is striking to us that four of the species with the lowest prevalence of neoplasias in our dataset, the gray squirrel, the common dormouse, the striped grass mouse, and the common field vole are all from wild, urban populations. These necropsies come from the London Zoo which has a policy of performing a necropsy on any animal they recover that dies on its grounds, not just the animals under its care. This is a hint that cancer may well be less common in the wild, although this observation may be dependent on the species and their wildlife habitat^59^.

If the “gross” cause of death was obvious for an animal, an institution may not have submitted the animal’s samples for histopathology, and would not be included in our data collection. Similarly, if a particular type of neoplasia is difficult to detect in a necropsy (including some leukemias and intracerebral tumors), or was only present at a microscopic level, it may have been undercounted.

The functional assays also had some limitations. Currently, the most available cell type from animals is fibroblasts, and as more cell lines become available in the future, it will be important to test DNA damage response in other cell types. Additionally, with more sample availability in the future, important biological factors can be controlled for, like age and sex.The data analyzed here represent the first attempts to determine if mechanisms of cancer suppression are shared across species, and our data suggests that suppression of somatic mutations may be a common defense mechanism.

### The Future of Comparative Oncology

In the future, it will be important to collect additional data to validate our discoveries of species with particularly low and high cancer prevalence, such as those highlighted in Fig. 1. Several life history traits, such as basal metabolic rate, may explain cancer risk but with BMR measured in only a few species, we lacked statistical power to detect a relationship with neoplasia or malignancy prevalence. Here we have dramatically expanded the amount of data on cancer prevalence in non-human animals, but the accumulation of data must continue if we are to match the robustness seen in human cancer epidemiology. In particular, much could be learned from analyzing the age-incidence curves of cancer ^18,60^, but that would require significantly more individuals for each species.

There is a large gap in comparative oncology data on wild animals. Gathering data from free-ranging populations is challenging, as it is difficult to detect cancer due to the animals decomposing or being eaten before they can be necropsied. Additionally, accurate age estimates are much more difficult to obtain in wild populations compared to those managed by humans. However, wild animal populations would greatly enhance the field of comparative oncology by validating species that have low cancer prevalence and testing for evolutionary mismatches for animals in captivity.

## Methods

### Analysis of Veterinary Necropsy Records

We collected necropsy records with permission from 99 zoological institutions, aquariums, and other facilities that house animals under managed care. Gross necropsies had all been conducted by veterinary pathologists who specialize in nondomestic species and neoplasia was identified histologically by board-certified veterinary pathologists. Cases where suspect neoplasms were not examined histologically were excluded from the dataset. We used a terminology dictionary (ST6) to distinguish benign from malignant neoplasms based on the diagnoses in the necropsy reports. We excluded neonatal records to reduce bias from high levels of neonate and infant mortality that is common in many species. Because only common names were recorded for most records, we developed a tool, kestrel, to translate common names into scientific names which is available at https://pkg.go.dev/github.com/icwells/kestrel.

All of the institutions that provided prior approval for the use of their data in these analyses are Association of Zoos and Aquariums (AZA) accredited. AZA accreditation encourages the institution to perform a necropsy on all animals that die under their care to determine cause of death and to monitor morbidity and mortality of each species. Furthermore, each institution had IACUC approval with the Exotic Species Cancer Research Alliance (ESCRA) and the Arizona Cancer and Evolution Center (ACE) for the use of their deceased animal’s records of animals with neoplasia for use in this study. Previous analyses included both necropsies for animals diagnosed with neoplasia and animals that were still alive ^18^. In this study, we restricted our analyses to only necropsies - for both cancer and non-cancer diagnosed animals, because alive animals may harbor undetected cancer or might be eventually diagnosed with cancer, thus skewing estimates in cancer prevalence.

The data for our analyses are available in supplemental file 1.

### Comparative Phylogenetic Methods in Comparative Oncology

Interspecies comparisons must account for the shared ancestry and the constraint of natural selection on species’ traits before a determinant of any correlations can be made. For the life history models of neoplasia and malignancy prevalence, the R programming ^61^ packages “phytools” ^62^, “ape” ^63^, and “caper” ^64^ were all used for phylogenetic comparisons and the handling of phylogenetic data. To accomplish this we wrote the function *pglsSEyPagel* which is built upon phytool’s *pglsSEy* (phylogenetic generalized least squares for uncertainty in Y). pglsSEyPagel expands upon the pglsSEy function by adding the estimate of Pagel’s lambda ^65^ to the regression, rather than assuming it is fixed at 1 (i.e., Brownian motion). Compar.gee is another method from the “ape” package that utilizes phylogenetic input, but uses a Generalized Estimating Equation to carry out binomial regressions^19^ which take into account the sample size for each species. The p-values for malignancy and neoplasia prevalence varied minimally when univariate tests for body mass, longevity, and gestation length were conducted. We did see larger coefficients with the results from this model (see ST4).

In order to validate our *pglsSEyPagel* findings, we carried out a bootstrap analysis. For each species, we randomly selected half of its individual records to be aggregated at the species level. Each re-sample would then be regressed using *pglsSEyPagel*, testing for the relationship between malignancy prevalence or neoplasia prevalence and the chosen life history variable. This iterative process was repeated 100 times each for body mass, longevity, and gestation length. Additionally, we controlled our *pglsSEyPagel* results by individual age for those which age records available, instead of depending upon longevity records from life history databases. Values did change, but conclusions remained the same (see ST3).

### Testing for relationships with life history factors

We extracted data for maximum lifespan, adult body weight, basal metabolic rate, gestation length, litter size, time to sexual maturity, and growth rate from PanTHERIA, AnAge and Amniote ^66–68^. We used a weighted phylogenetic regression to control for non-independence of phenotypes (e.g. neoplasia prevalence) in closely related species. We report the phylogenetic signal, lambda, for each analysis, along with the p-value and R^2^. Due to the nature of the PGLS model, R2 is not a standard output. To report the fit of the model, we employed a pseudo R2 approach using the ‘rr2’ R package. The function ‘R2’ from the package can utilize phylogenetic relationships within the R^2^ calculation. A single phylogenetic tree encompassing the three clades was collected from timetree.org. We pruned the tree to the 292 species in our data set using the setdiff and keep.tip/drop.tip functions in the APE R package. Estimates for neoplasia and malignancy prevalence are more accurate in species with more necropsies. To address the differences in number of necropsies, and to limit the noise from prevalence estimates based on few individuals, we weighted the species data points by the square-root of the number of necropsies records we have. Our R code for all analyses and figures included in this manuscript is freely available at https://github.com/zacharycompton/cancerAcrossVertebrates.git. In addition, we only analyzed species for which we had at least 20 necropsy results (previous studies had used 10 ^16^ or 20 ^17,18^ for the lower bound number of individuals). The main *pglsSEyPagel* analyses were done with all species together, including mammals, sauropsids and amphibians. In the analyses of litter size and gestation time, we also tested for a relationship with neoplasia prevalence in mammals alone. We carried out a total of 28 *pglsSEyPagel* analyses. To control for multiple testing, we used a false discovery rate (FDR) of 10%.

### DNA Damage Response Assays

Established, primary cells from mammals were obtained from San Diego Zoo Wildlife Alliance (Capybara, Linne’s Two Toed Sloth, Red Necked Wallaby, Rock Hyrax, Rodrigues Fruit Bat, Six Banded Armadillo, Southern White Rhino, and Virginia Opossum) or generated at Huntsman Cancer Institute from tissues collected from African Pygmy Hedgehog, Domestic Rabbit, Leopard, Asian Elephant, and Cape hunting dog. Brown rat (Cell Applications) and Normal Human Dermal Fibroblasts (Lonza) were commercially available. Detailed information on culture conditions, primary donor demographics, and passage numbers can be found in the supplementary information. Cells were seeded in 96-well plates at 2,000 cells per well in cell growth media and allowed to adhere overnight. The following day, doxorubicin was added at one of four concentrations (0μM [DMSO vehicle control], 0.11μM, 0.33μM, and 1μM). Each condition was tested in triplicate in three separate experiments. Cell proliferation and apoptosis were measured by real-time fluorescence microscopy (IncuCyte, Sartorius) at two-hour intervals for three days. Apoptosis was measured using a fluorescent cell death marker, Annexin V Dye (Sartorius). Images were processed and analyzed using IncuCyte software. The number of dead or dying cells were identified by counting Annexin V positive cells. In addition, cell count overtime was calculated using IncuCyte cell-by-cell software. To measure response to radiation-induced DNA damage, cells were irradiated with one of four doses: 0Gy, 0.4Gy, 2Gy, and 10 Gy. Radiation dose was delivered using an RS-2000 X-Ray Irradiator (Radsource). Cells were then seeded in 96-well plates in cell growth media containing Annexin V Dye (Sartorius). Cells were imaged by real-time fluorescence microscopy (IncuCyte, Sartorius) at two-hour intervals for five days. We estimated cell cycle arrest by normalizing the cell count of irradiated cells to untreated cells by dividing the area under the curve (AUC) of cell count over-time for treated cells by the AUC of cell count over time for the untreated (UT) cells. We converted that number into a percentage that represents the percent of cell proliferation relative to untreated cells. We then tested if this normalized amount of cell growth was predictive of neoplasia prevalence using the phylogenetically controlled *pglsSEyPagel* regression (Fig. 4A). Cagan et al. published somatic mutation rates (single base substitution per genome per year) for 16 species based on sequencing 208 individual colonic crypts from 56 animals from the London Zoo^27^. They divided the number of somatic mutations, detected by whole genome sequencing, by the age (in years) of the individual at the time that the tissue was taken, to estimate the mutation rate per year. Nine of those species are in our dataset, allowing us to use a pgls regression to test for an association between mean somatic mutation rates and neoplasia prevalence.

## Conclusion

Cancer is a problem of multicellularity ^1^. While we found a relationship between both body mass and gestation time with cancer prevalence, we are just beginning to discover patterns of cancer susceptibility and cancer defenses across species. It is likely species evolved a variety of mechanisms to suppress cancer. The discovery of particular species with extremely low neoplasia prevalence provides opportunities to elucidate cancer suppression mechanisms that are compatible with life and reproductive success. Investigation of species with extreme vulnerability to a particular cancer may also help us understand those cancers as well as human syndromes that predispose to those cancers. We hope to learn from nature how to better prevent cancer in both humans and non-human animals.

## Supporting information

Supplemental figures, tables, and data dictionary

## Acknowledgements

We would like to thank our dear colleague Drury Reavill who passed away prior to the preparation of this manuscript. We are forever grateful to her contribution to the field of veterinary science and her dedication to the health and survival of all animals. Many thanks to our continued collaboration with Michael Garner and Northwest Zoo Pathology who provided the annotated veterinarian pathology database. Liam Revell, the developer of the R software package “phytools”, was extremely helpful in providing input for the application of his methods in novel ways. We would also like to thank the zoos and aquaria who cared for these animals and gave us permission to analyze their data. These discoveries could not have been made without them. Among these contributing institutions we would like to specifically thank these zoos: Akron Zoo, Atlanta Zoo, Audubon Nature Institute, Bergen County Zoo, Birmingham Zoo, Buffalo Zoo, Capron Park Zoo, Central Florida Zoo, Dallas Zoo, El Paso Zoo, Elmwood Park Zoo, Fort Worth Zoo, Gladys Porter Zoo, Greensboro Science Center, Henry Doorly Zoo, Utah’s Hogle Zoo, Jacksonville Zoo, John Ball Zoo, Los Angeles Zoo, Louisville Zoo, Mesker Park Zoo, Miami Zoo, Oakland Zoo, Oklahoma City Zoo, Philadelphia Zoo, Phoenix Zoo, Pueblo Zoo, San Antonio Zoo, Santa Ana Zoo, Santa Barbara Zoo, Sedgwick County Zoo, Seneca Park Zoo, The Brevard Zoo, The Detroit Zoo, The Oregon Zoo, and Toledo Zoo.

## Funding

National Institutes of Health grant U54 CA217376 (CCM, ZTC, JDS, LMA, AMB, TMH)

National Institutes of Health grant U2C CA233254 (CCM)

National Institutes of Health grant P01 CA91955 (CCM)

National Institutes of Health grant R01 CA140657 (CCM)

National Institutes of Health grant T32 CA272303 (ZTC)

CDMRP Breast Cancer Research Program Award BC132057 (CCM)

Arizona Biomedical Research Commission grant ADHS18-198847 (CCM)

Hyundai Hope On Wheels (Hope On Wheels) (JDS)

Soccer for Hope Foundation (JDS)

Li-Fraumeni Syndrome Association (JDS)

Kneaders Hope Fights Childhood Cancer (JDS)

Helen Clise Presidential Endowed Chair in Li-Fraumeni Syndrome Research (JDS)

Primary Children’s Hospital Foundation (LMA)

University of Utah Department of Pediatrics Research Enterprise (LMA)

The findings, opinions and recommendations expressed here are those of the authors and not necessarily those of the universities where the research was performed or the National Institutes of Health.

## Author Contributions

Conceptualization: ZTC, CCM, AA, AMB

Data Collection: AMB, HD, TMH, DM, MG, VH, LMA, KN, MW, RK, GF

Data Curation: SR, ZTC, JL, DM, VH, SEK, CB, BT, LMA, KN, MW, RK, GF

Veterinary Pathology Support: MG, BT, SS, ED, AZ, LMA, KN, MW

Methodology: ZTC, LR, MW, DM, OV, RK

Investigation: ZTC, SR, WM, AP, SA, LMA, KN, MW, RK, GF

Visualization: WM, BM, SA, SS, SA

Supervision: CCM, AMB, AA, MT, TAG, MG, LA, JS, TMH

Writing - original draft: ZTC, VH, CCM, AA, AMB, LMA

Writing - review and editing: All Authors

## Data and materials availability

All data and code are available at <u>zacharycompton/cancerAcrossVertebrates</u> (github.com), with the exception of the ages of individual animals and the locations of their tumors, which are restricted due to privacy agreements with the contributing zoos. Access to that data may be granted with permissions of the zoos.

## Supplementary Materials

Figs S1 to S74

Tables ST1 to ST8

