## Supplemental figures, tables, and data dictionary for "Cancer Prevalence Across Vertebrates": S58arcsinMin20.html

arcsineMin20all


### arcsineMin20all

###### Walker M

#### 2024-04-17

```
library(nlme)
library(rms)
```

```
## Loading required package: Hmisc
```

```
## 
## Attaching package: 'Hmisc'
```

```
## The following objects are masked from 'package:base':
## 
##     format.pval, units
```

```
library(phytools)
```

```
## Loading required package: ape
```

```
## 
## Attaching package: 'ape'
```

```
## The following object is masked from 'package:Hmisc':
## 
##     zoom
```

```
## Loading required package: maps
```

```
library(geiger)
library(caper)
```

```
## Loading required package: MASS
```

```
## Loading required package: mvtnorm
```

```
library(tidyverse)
```

```
## ── Attaching core tidyverse packages ──────────────────────── tidyverse 2.0.0 ──
## ✔ dplyr     1.1.4     ✔ readr     2.1.5
## ✔ forcats   1.0.0     ✔ stringr   1.5.1
## ✔ ggplot2   3.5.0     ✔ tibble    3.2.1
## ✔ lubridate 1.9.3     ✔ tidyr     1.3.1
## ✔ purrr     1.0.2
```

```
## ── Conflicts ────────────────────────────────────────── tidyverse_conflicts() ──
## ✖ dplyr::collapse()  masks nlme::collapse()
## ✖ dplyr::filter()    masks stats::filter()
## ✖ dplyr::lag()       masks stats::lag()
## ✖ purrr::map()       masks maps::map()
## ✖ dplyr::select()    masks MASS::select()
## ✖ dplyr::src()       masks Hmisc::src()
## ✖ dplyr::summarize() masks Hmisc::summarize()
## ✖ dplyr::where()     masks ape::where()
## ℹ Use the conflicted package (<http://conflicted.r-lib.org/>) to force all conflicts to become errors
```

```
library(cowplot)
```

```
## 
## Attaching package: 'cowplot'
## 
## The following object is masked from 'package:lubridate':
## 
##     stamp
```

```
library(ggrepel)
library(ggsci)
library(patchwork)
```

```
## 
## Attaching package: 'patchwork'
## 
## The following object is masked from 'package:cowplot':
## 
##     align_plots
## 
## The following object is masked from 'package:MASS':
## 
##     area
```

```
#make sure to run all of this before you get to work.
#pgls sey base (just run all of this)
modPgls.SEy = function (model, data, corClass = corBrownian, tree, se = NULL, 
                        method = c("REML", "ML"), interval = c(0, 1000), corClassValue=1, sig2e=NULL, ...) 
{
  Call <- match.call()
  corfunc <- corClass
  spp <- rownames(data)
  data <- cbind(data, spp)
  if (is.null(se)) 
    se <- setNames(rep(0, Ntip(tree)), tree$tip.label)[spp]
  else se <- se[spp]
  
  lk <- function(sig2e, data, tree, model, ve, corfunc, spp) {
    tree$edge.length <- tree$edge.length * sig2e
    ii <- sapply(1:Ntip(tree), function(x, e) which(e == 
                                                      x), e = tree$edge[, 2])
    tree$edge.length[ii] <- tree$edge.length[ii] + ve[tree$tip.label]
    vf <- diag(vcv(tree))[spp]
    w <- varFixed(~vf)
    COR <- corfunc(corClassValue, tree, form = ~spp, ...)
    fit <- gls(model, data = cbind(data, vf), correlation = COR, 
               method = method, weights = w)
    -logLik(fit)
  }
  
  if (is.null(sig2e)) {
    fit <- optimize(lk, interval = interval, data = data, tree = tree, 
                    model = model, ve = se^2, corfunc = corfunc, spp = spp)
    sig2e=fit$minimum
  }
  
  tree$edge.length <- tree$edge.length * sig2e
  ii <- sapply(1:Ntip(tree), function(x, e) which(e == x), 
               e = tree$edge[, 2])
  tree$edge.length[ii] <- tree$edge.length[ii] + se[tree$tip.label]^2
  vf <- diag(vcv(tree))[spp]
  w <- varFixed(~vf)
  obj <- gls(model, data = cbind(data, vf), correlation = corfunc(corClassValue, 
                                                                  tree, form = ~spp, ...), weights = w, method = method)
  obj$call <- Call
  obj$sig2e <- sig2e
  obj
}

#Internal function
pglsSEyPagelToOptimizeLambda=function(lambda,model,data,tree,...) {
  -logLik(modPgls.SEy(model=model,data=data,tree=tree,corClassValue=lambda,corClass=corPagel,fixed=T,...)) #Returns -logLikelihood of the pgls.SEy model with lambda fixed to the value of the lambda argument. sig2e will be optimized within modPgls.SEy unless given as an argument here
}

#Function intended for users
pglsSEyPagel=function(model, data, tree, lambdaInterval=c(0,1),...){
  optimizedModel=optimize(pglsSEyPagelToOptimizeLambda,lambdaInterval,model=model,data=data,tree=tree,...) #Optimizes lambda in the lambdaInterval using the pglsSEyPagelToOptimizeLambda function
  return(modPgls.SEy(model=model,data=data,tree=tree,corClass=corPagel,fixed=T,corClassValue=optimizedModel$minimum,...)) #Returns the final model fit
}


#read data
Data <- read.csv("min20-2022.05.16.csv")
```

adult weight

```
#adult weight models
#adult weight neo

cutData <- Data[,c(5,9,10,11,13,38,42),drop=FALSE] 
cutData[cutData$adult_weight == -1, ] <-NA
cutData <- na.omit(cutData)
tree <- read.tree("min20Fixed516.nwk")

cutData$Species <- gsub(" ", "_", cutData$Species) 
cutData$common_name<-gsub("_", "", cutData$common_name)
includedSpecies<-cutData$Species
pruned.tree<-drop.tip(
  tree, setdiff(
    tree$tip.label, includedSpecies))
pruned.tree <- keep.tip(pruned.tree,pruned.tree$tip.label)
cutData$Keep <- cutData$Species %in% pruned.tree$tip.label
cutData <- cutData[!(cutData$Keep==FALSE),]
rownames(cutData)<-cutData$Species
SE<-setNames(cutData$SE_simple,cutData$Species)[rownames(cutData)]
view(cutData)

cutData$NeoplasiaPrevalenceArc <- asin(sqrt(cutData$NeoplasiaPrevalence))

#pgls model
adult.weight.neo<-pglsSEyPagel(NeoplasiaPrevalenceArc~log10(adult_weight.g.),data=cutData,tree=pruned.tree,se=SE,method = "ML")

summary(adult.weight.neo)
```

```
## Generalized least squares fit by maximum likelihood
##   Model: model 
##   Data: data 
##         AIC       BIC  logLik
##   -134.1556 -123.8544 70.0778
## 
## Correlation Structure: corPagel
##  Formula: ~spp 
##  Parameter estimate(s):
##    lambda 
## 0.3899542 
## Variance function:
##  Structure: fixed weights
##  Formula: ~vf 
## 
## Coefficients:
##                             Value  Std.Error  t-value p-value
## (Intercept)            0.17530272 0.07012631 2.499814  0.0131
## log10(adult_weight.g.) 0.04144421 0.01299026 3.190408  0.0016
## 
##  Correlation: 
##                        (Intr)
## log10(adult_weight.g.) -0.457
## 
## Standardized residuals:
##        Min         Q1        Med         Q3        Max 
## -1.8199548 -0.7718386 -0.2102776  0.5080215  2.9792343 
## 
## Residual standard error: 0.0003501939 
## Degrees of freedom: 229 total; 227 residual
```

```
#grab r squared, lambda, and p values from summary 

r.v.adult.weight.neo <- summary(adult.weight.neo)$corBeta
r.v.adult.weight.neo <- format(r.v.adult.weight.neo[2,1])
r.v.adult.weight.neo <-signif(as.numeric(r.v.adult.weight.neo)^2, digits= 2)
ld.v.adult.weight.neo<- summary(adult.weight.neo)$modelStruct$corStruct
ld.v.adult.weight.neo <- signif(ld.v.adult.weight.neo[1], digits = 2)
p.v.adult.weight.neo<-summary(adult.weight.neo)$tTable
p.v.adult.weight.neo<-signif(p.v.adult.weight.neo[2,4], digits = 2)
```

```
#gestation mal
cutData <- Data[,c(5,9,10,11,17,30,42),drop=FALSE] 
cutData[cutData$Gestation.months. < 0, ] <-NA
cutData <- na.omit(cutData)
tree <- read.tree("min20Fixed516.nwk")

cutData$Species <- gsub(" ", "_", cutData$Species) 
cutData$common_name<-gsub("_", "", cutData$common_name)
includedSpecies <- cutData$Species
pruned.tree<-drop.tip(
  tree, setdiff(
    tree$tip.label, includedSpecies))
pruned.tree <- keep.tip(pruned.tree,pruned.tree$tip.label)
cutData$Keep <- cutData$Species %in% pruned.tree$tip.label
cutData <- cutData[!(cutData$Keep==FALSE),]
rownames(cutData)<-cutData$Species
SE<-setNames(cutData$SE_simple,cutData$Species)[rownames(cutData)]
view(cutData)

cutData$MalignancyPrevalenceArc <- asin(sqrt(cutData$MalignancyPrevalence))

#pgls model
gestation.mal<-pglsSEyPagel(MalignancyPrevalenceArc~log10(Gestation.months.),data=cutData,
                            tree=pruned.tree,method="ML",se=SE)
summary(gestation.mal)
```

```
## Generalized least squares fit by maximum likelihood
##   Model: model 
##   Data: data 
##         AIC       BIC   logLik
##   -103.7917 -94.73984 54.89584
## 
## Correlation Structure: corPagel
##  Formula: ~spp 
##  Parameter estimate(s):
##    lambda 
## 0.4809645 
## Variance function:
##  Structure: fixed weights
##  Formula: ~vf 
## 
## Coefficients:
##                                Value  Std.Error   t-value p-value
## (Intercept)               0.20130880 0.06759191  2.978297  0.0034
## log10(Gestation.months.) -0.07212253 0.04863145 -1.483043  0.1402
## 
##  Correlation: 
##                          (Intr)
## log10(Gestation.months.) -0.095
## 
## Standardized residuals:
##        Min         Q1        Med         Q3        Max 
## -1.1209844 -0.3595640  0.4201036  0.9938786  2.5763669 
## 
## Residual standard error: 0.000346148 
## Degrees of freedom: 151 total; 149 residual
```

```
#grab r squared, lambda, and p values from summary 

r.v.gestmal <- summary(gestation.mal)$corBeta
r.v.gestmal <- format(r.v.gestmal[2,1])
r.v.gestmal<-signif(as.numeric(r.v.gestmal)^2, digits= 2)
ld.v.gestmal<- summary(gestation.mal)$modelStruct$corStruct
ld.v.gestmal<- signif(ld.v.gestmal[1], digits = 2)
p.v.gestmal<-summary(gestation.mal)$tTable
p.v.gestmal<-signif(p.v.gestmal[2,4], digits = 3)
```

```
### Longevity model
#longevity neo
cutData <- Data[,c(5,9,10,11,13,40,42),drop=FALSE] 
cutData[cutData$max_longevity.months. < 0,] <-NA
cutData <- na.omit(cutData)
tree <- read.tree("min20Fixed516.nwk")

view(cutData)

cutData$Species <- gsub(" ", "_", cutData$Species) 
cutData$common_name<-gsub("_", "", cutData$common_name)
includedSpecies<-cutData$Species
pruned.tree<-drop.tip(
  tree, setdiff(
    tree$tip.label, includedSpecies))
pruned.tree <- keep.tip(pruned.tree,pruned.tree$tip.label)
cutData$Keep <- cutData$Species %in% pruned.tree$tip.label
cutData <- cutData[!(cutData$Keep==FALSE),]
rownames(cutData)<-cutData$Species
SE<-setNames(cutData$SE_simple,cutData$Species)[rownames(cutData)]
view(cutData)

cutData$NeoplasiaPrevalenceArc <- asin(sqrt(cutData$NeoplasiaPrevalence))

#pgls model
longevity.neo<-pglsSEyPagel(NeoplasiaPrevalenceArc~max_longevity.months.,data=cutData,
                            tree=pruned.tree,method="ML",se=SE)
summary(longevity.neo)
```

```
## Generalized least squares fit by maximum likelihood
##   Model: model 
##   Data: data 
##         AIC       BIC   logLik
##   -104.9895 -94.94814 55.49473
## 
## Correlation Structure: corPagel
##  Formula: ~spp 
##  Parameter estimate(s):
##    lambda 
## 0.4136901 
## Variance function:
##  Structure: fixed weights
##  Formula: ~vf 
## 
## Coefficients:
##                            Value  Std.Error  t-value p-value
## (Intercept)           0.16333159 0.06636922 2.460954  0.0147
## max_longevity.months. 0.00021279 0.00009663 2.202093  0.0288
## 
##  Correlation: 
##                       (Intr)
## max_longevity.months. -0.364
## 
## Standardized residuals:
##        Min         Q1        Med         Q3        Max 
## -1.4851709 -0.2439729  0.2663664  0.9175001  3.3207412 
## 
## Residual standard error: 0.0003669875 
## Degrees of freedom: 210 total; 208 residual
```

```
#grab r squared, lambda, and p values from summary 

r.v.longneo <- summary(longevity.neo)$corBeta
r.v.longneo <- format(r.v.longneo[2,1])
r.v.longneo <-signif(as.numeric(r.v.longneo)^2, digits= 2)
ld.v.longneo<- summary(longevity.neo)$modelStruct$corStruct
ld.v.longneo <- signif(ld.v.longneo[1], digits = 2)
p.v.longneo<-summary(longevity.neo)$tTable
p.v.longneo<-signif(p.v.longneo[2,4], digits = 3)
```
