## Supplemental figures, tables, and data dictionary for "Cancer Prevalence Across Vertebrates": ST1 Mammal Leaderboard.pptx

### Slide 1
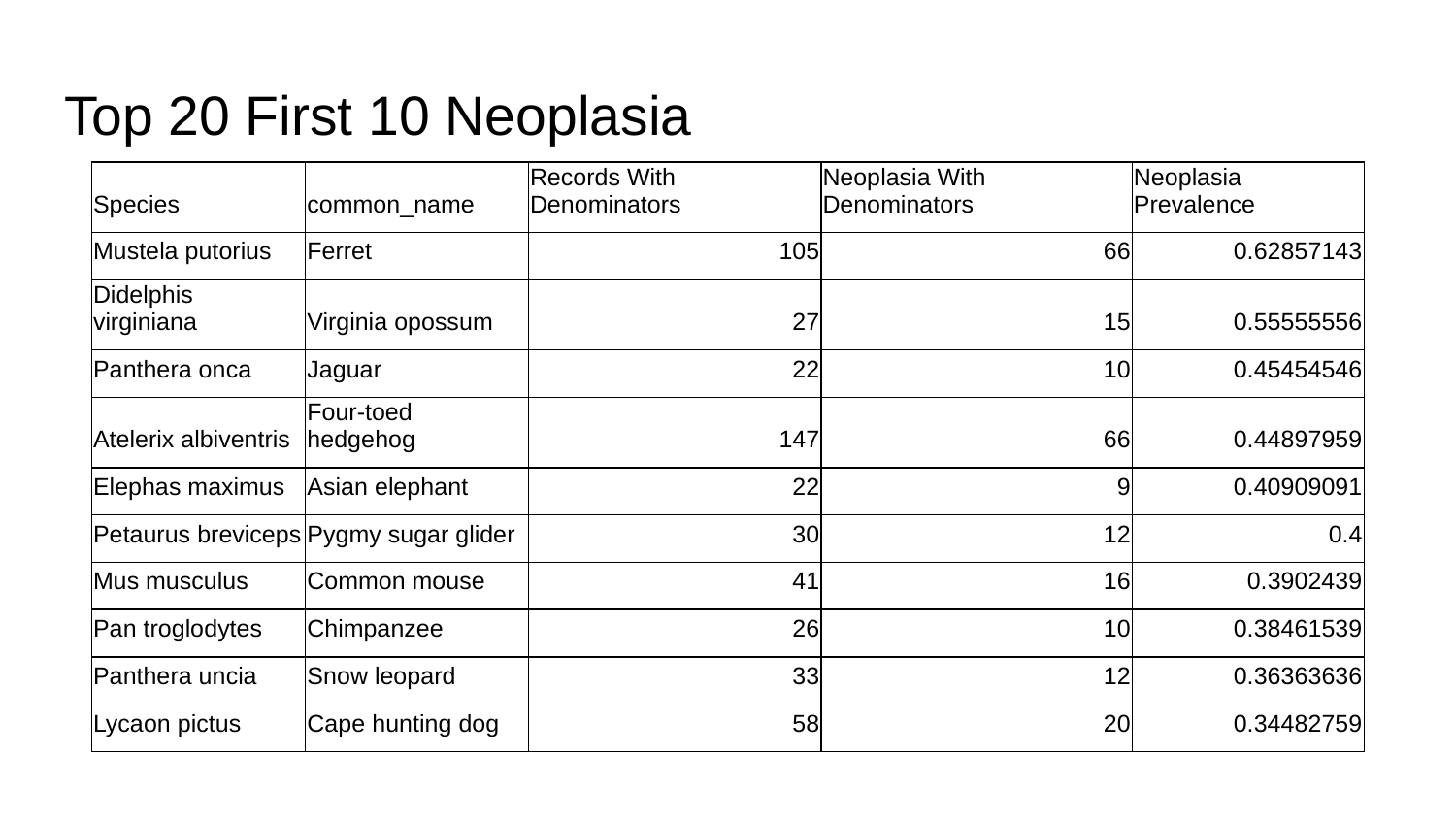

#### Top 20 First 10 Neoplasia
| Species | common\_name | Records With Denominators | Neoplasia With Denominators | Neoplasia Prevalence |
| --- | --- | --- | --- | --- |
| Mustela putorius | Ferret | 105 | 66 | 0.62857143 |
| Didelphis virginiana | Virginia opossum | 27 | 15 | 0.55555556 |
| Panthera onca | Jaguar | 22 | 10 | 0.45454546 |
| Atelerix albiventris | Four-toed hedgehog | 147 | 66 | 0.44897959 |
| Elephas maximus | Asian elephant | 22 | 9 | 0.40909091 |
| Petaurus breviceps | Pygmy sugar glider | 30 | 12 | 0.4 |
| Mus musculus | Common mouse | 41 | 16 | 0.3902439 |
| Pan troglodytes | Chimpanzee | 26 | 10 | 0.38461539 |
| Panthera uncia | Snow leopard | 33 | 12 | 0.36363636 |
| Lycaon pictus | Cape hunting dog | 58 | 20 | 0.34482759 |

### Slide 2
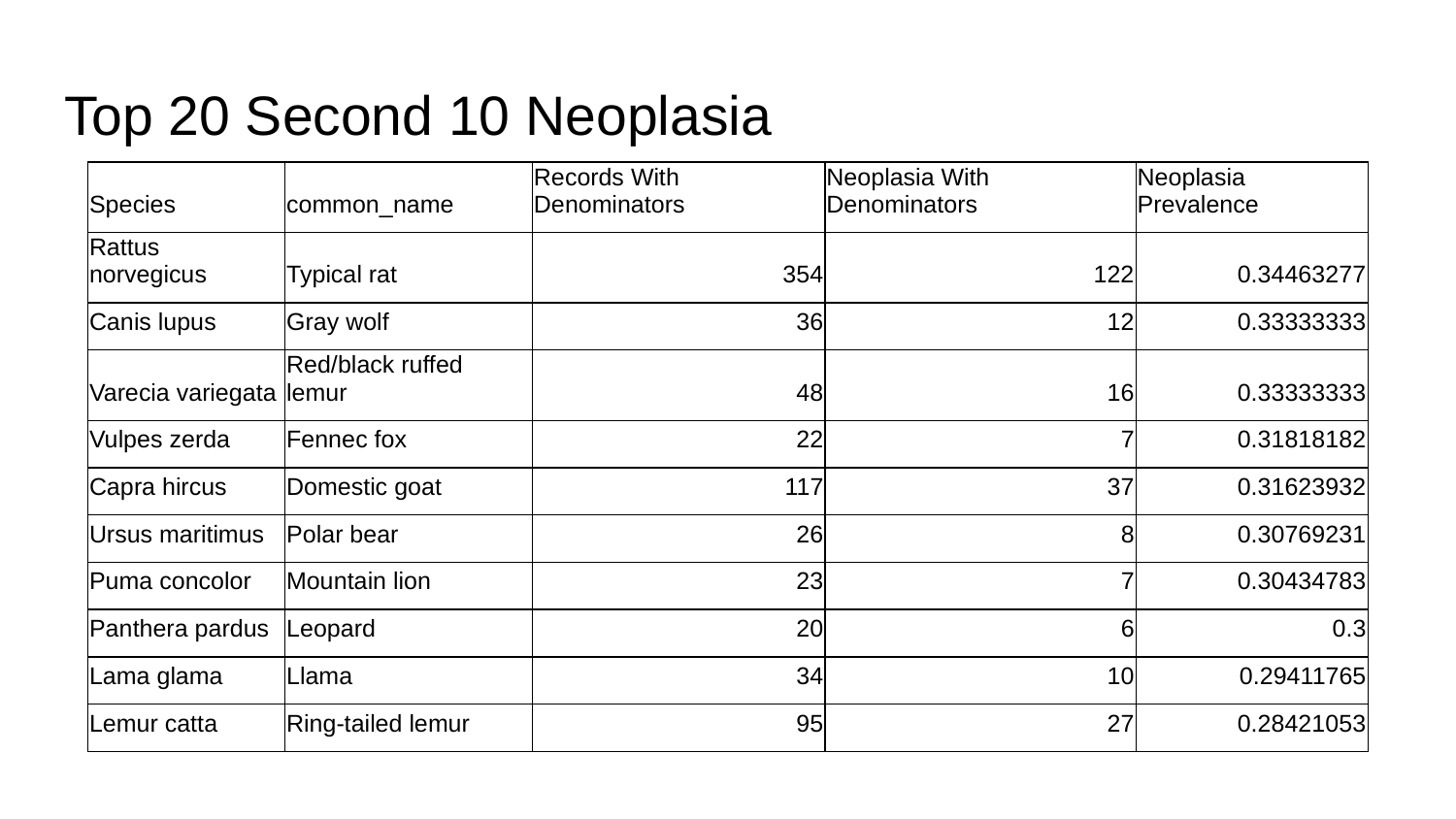

#### Top 20 Second 10 Neoplasia
| Species | common\_name | Records With Denominators | Neoplasia With Denominators | Neoplasia Prevalence |
| --- | --- | --- | --- | --- |
| Rattus norvegicus | Typical rat | 354 | 122 | 0.34463277 |
| Canis lupus | Gray wolf | 36 | 12 | 0.33333333 |
| Varecia variegata | Red/black ruffed lemur | 48 | 16 | 0.33333333 |
| Vulpes zerda | Fennec fox | 22 | 7 | 0.31818182 |
| Capra hircus | Domestic goat | 117 | 37 | 0.31623932 |
| Ursus maritimus | Polar bear | 26 | 8 | 0.30769231 |
| Puma concolor | Mountain lion | 23 | 7 | 0.30434783 |
| Panthera pardus | Leopard | 20 | 6 | 0.3 |
| Lama glama | Llama | 34 | 10 | 0.29411765 |
| Lemur catta | Ring-tailed lemur | 95 | 27 | 0.28421053 |

### Slide 3
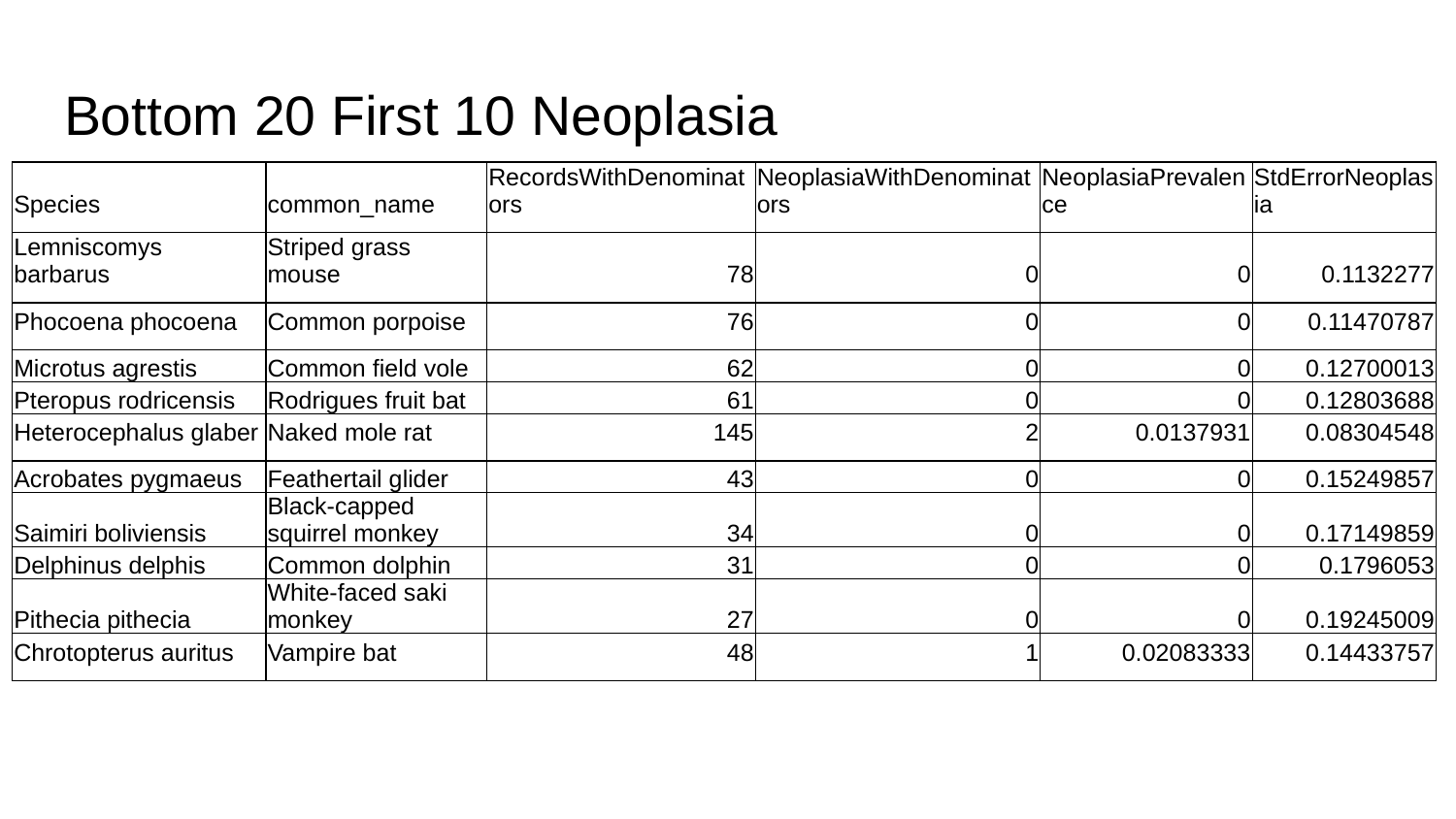

#### Bottom 20 First 10 Neoplasia
| Species | common\_name | RecordsWithDenominators | NeoplasiaWithDenominators | NeoplasiaPrevalence | StdErrorNeoplasia |
| --- | --- | --- | --- | --- | --- |
| Lemniscomys barbarus | Striped grass mouse | 78 | 0 | 0 | 0.1132277 |
| Phocoena phocoena | Common porpoise | 76 | 0 | 0 | 0.11470787 |
| Microtus agrestis | Common field vole | 62 | 0 | 0 | 0.12700013 |
| Pteropus rodricensis | Rodrigues fruit bat | 61 | 0 | 0 | 0.12803688 |
| Heterocephalus glaber | Naked mole rat | 145 | 2 | 0.0137931 | 0.08304548 |
| Acrobates pygmaeus | Feathertail glider | 43 | 0 | 0 | 0.15249857 |
| Saimiri boliviensis | Black-capped squirrel monkey | 34 | 0 | 0 | 0.17149859 |
| Delphinus delphis | Common dolphin | 31 | 0 | 0 | 0.1796053 |
| Pithecia pithecia | White-faced saki monkey | 27 | 0 | 0 | 0.19245009 |
| Chrotopterus auritus | Vampire bat | 48 | 1 | 0.02083333 | 0.14433757 |

### Slide 4
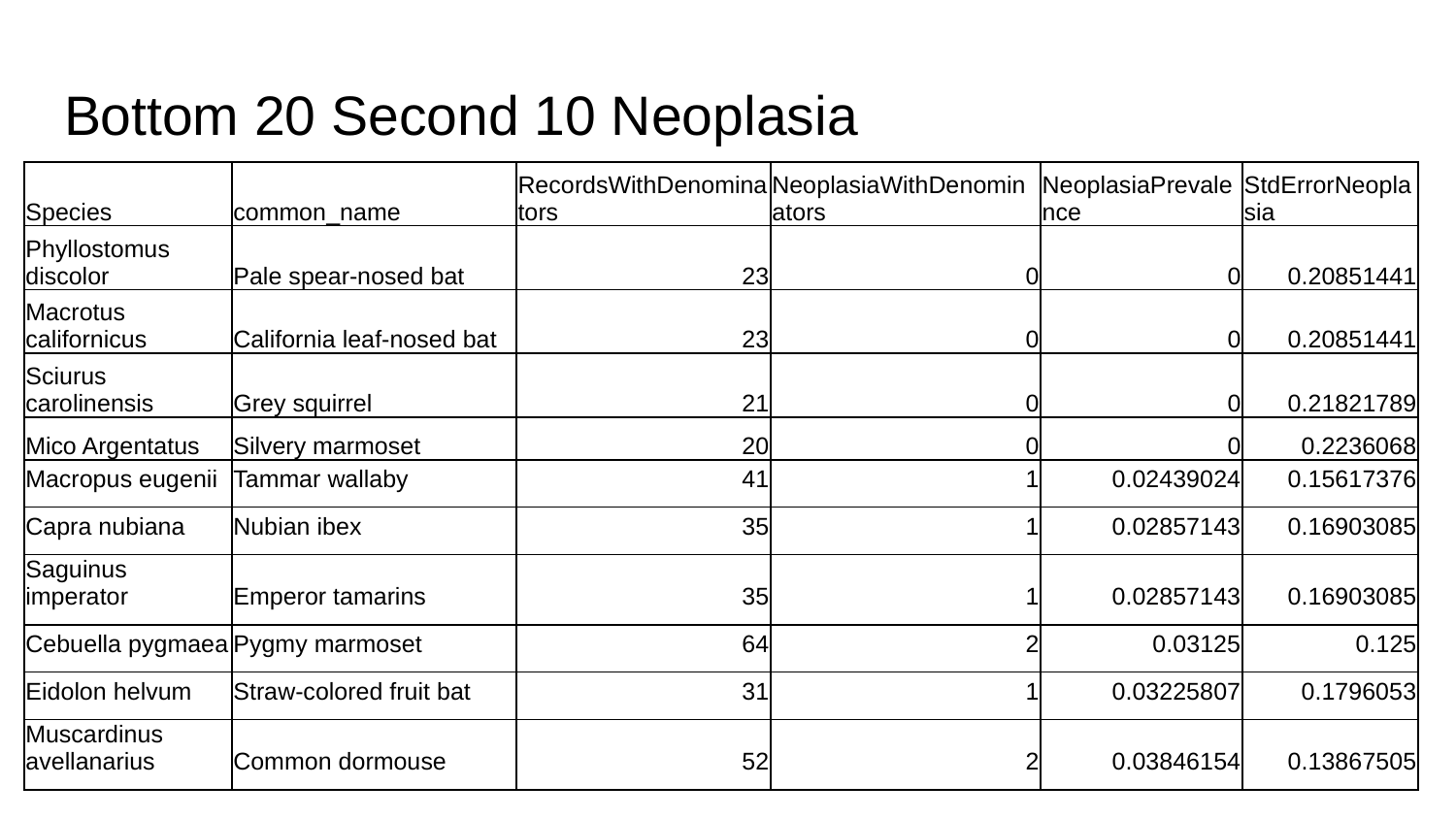

#### Bottom 20 Second 10 Neoplasia
| Species | common\_name | RecordsWithDenominators | NeoplasiaWithDenominators | NeoplasiaPrevalence | StdErrorNeoplasia |
| --- | --- | --- | --- | --- | --- |
| Phyllostomus discolor | Pale spear-nosed bat | 23 | 0 | 0 | 0.20851441 |
| Macrotus californicus | California leaf-nosed bat | 23 | 0 | 0 | 0.20851441 |
| Sciurus carolinensis | Grey squirrel | 21 | 0 | 0 | 0.21821789 |
| Mico Argentatus | Silvery marmoset | 20 | 0 | 0 | 0.2236068 |
| Macropus eugenii | Tammar wallaby | 41 | 1 | 0.02439024 | 0.15617376 |
| Capra nubiana | Nubian ibex | 35 | 1 | 0.02857143 | 0.16903085 |
| Saguinus imperator | Emperor tamarins | 35 | 1 | 0.02857143 | 0.16903085 |
| Cebuella pygmaea | Pygmy marmoset | 64 | 2 | 0.03125 | 0.125 |
| Eidolon helvum | Straw-colored fruit bat | 31 | 1 | 0.03225807 | 0.1796053 |
| Muscardinus avellanarius | Common dormouse | 52 | 2 | 0.03846154 | 0.13867505 |

### Slide 5
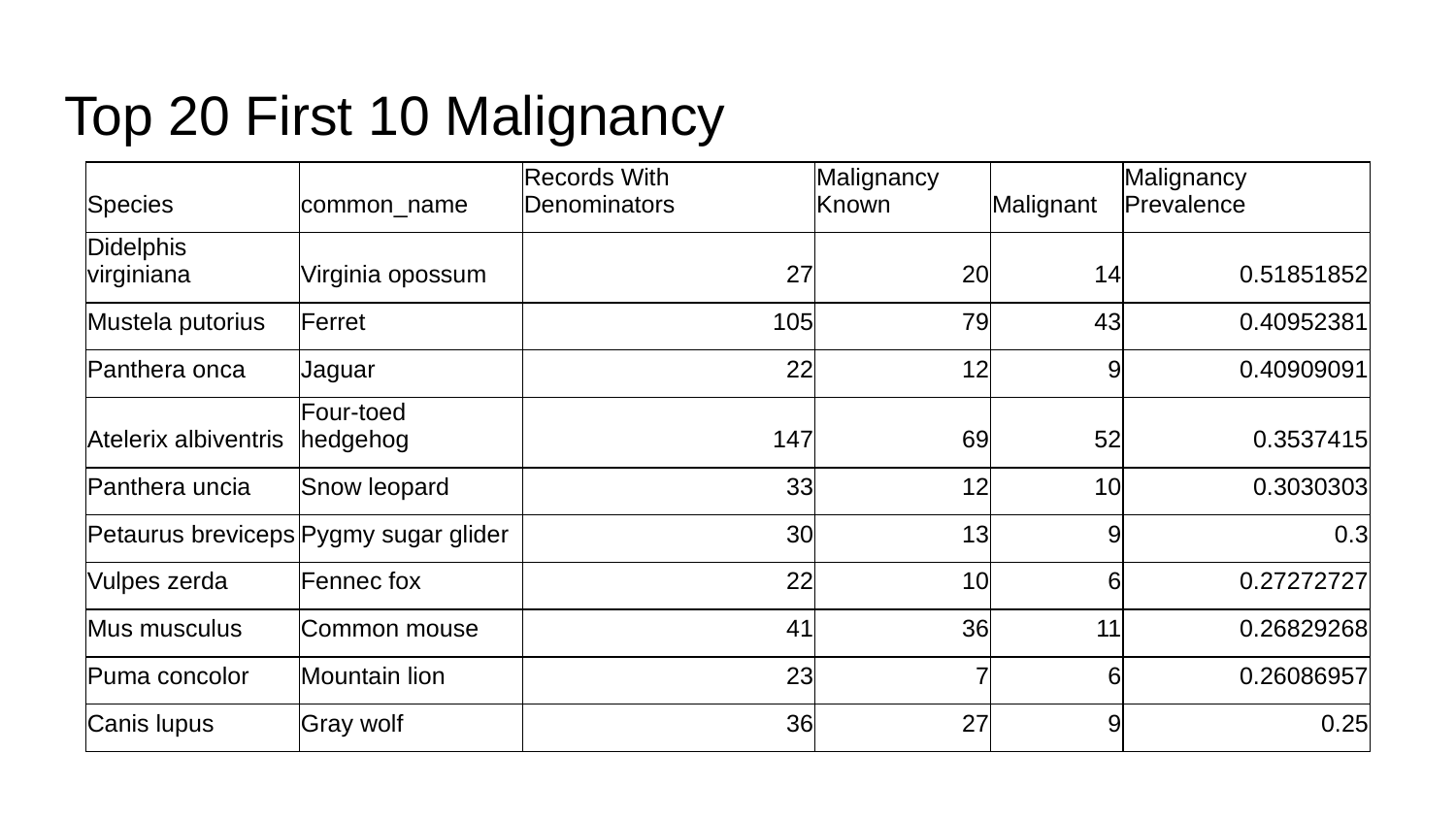

#### Top 20 First 10 Malignancy
| Species | common\_name | Records With Denominators | Malignancy Known | Malignant | Malignancy Prevalence |
| --- | --- | --- | --- | --- | --- |
| Didelphis virginiana | Virginia opossum | 27 | 20 | 14 | 0.51851852 |
| Mustela putorius | Ferret | 105 | 79 | 43 | 0.40952381 |
| Panthera onca | Jaguar | 22 | 12 | 9 | 0.40909091 |
| Atelerix albiventris | Four-toed hedgehog | 147 | 69 | 52 | 0.3537415 |
| Panthera uncia | Snow leopard | 33 | 12 | 10 | 0.3030303 |
| Petaurus breviceps | Pygmy sugar glider | 30 | 13 | 9 | 0.3 |
| Vulpes zerda | Fennec fox | 22 | 10 | 6 | 0.27272727 |
| Mus musculus | Common mouse | 41 | 36 | 11 | 0.26829268 |
| Puma concolor | Mountain lion | 23 | 7 | 6 | 0.26086957 |
| Canis lupus | Gray wolf | 36 | 27 | 9 | 0.25 |

### Slide 6
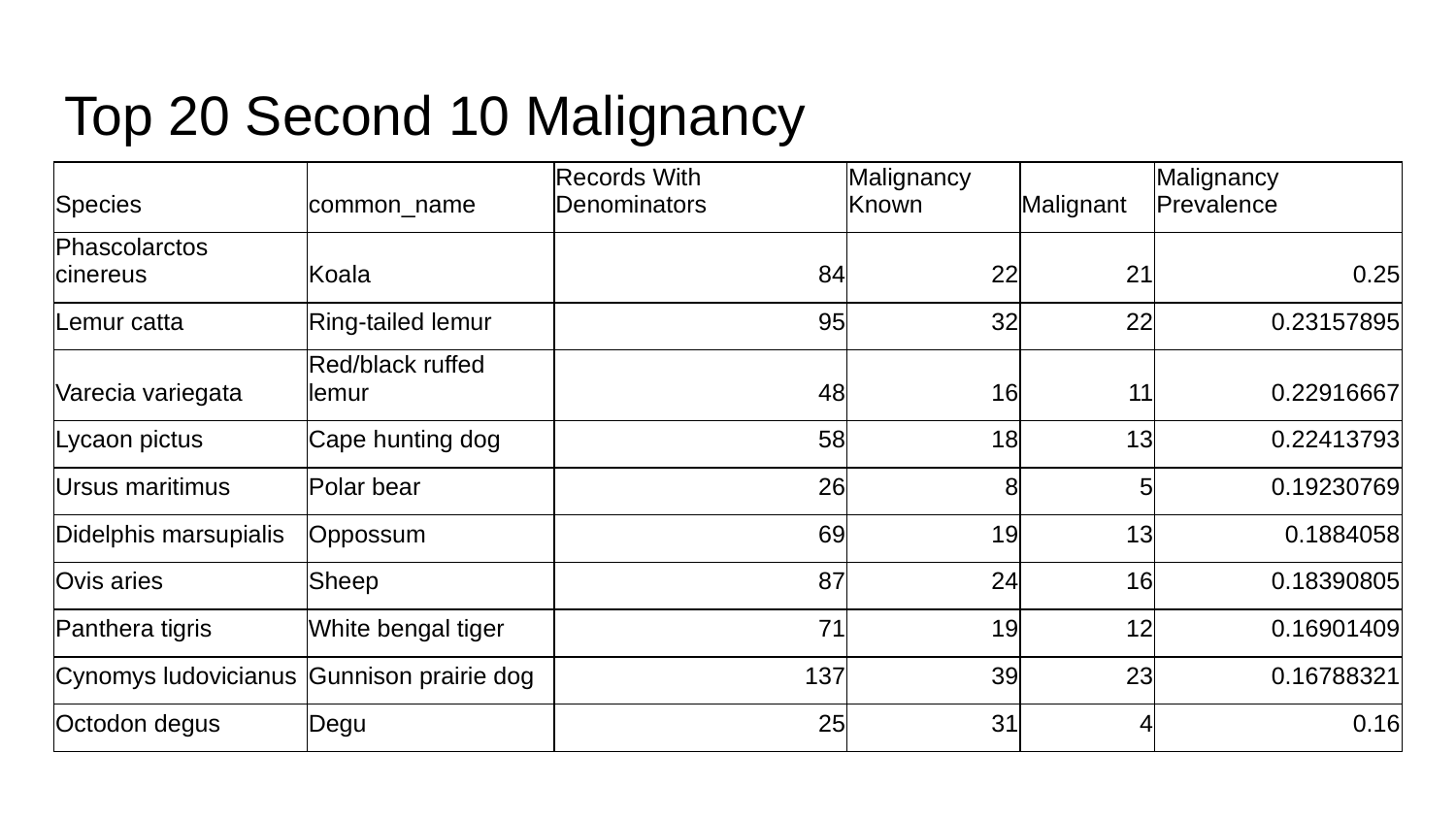

#### Top 20 Second 10 Malignancy
| Species | common\_name | Records With Denominators | Malignancy Known | Malignant | Malignancy Prevalence |
| --- | --- | --- | --- | --- | --- |
| Phascolarctos cinereus | Koala | 84 | 22 | 21 | 0.25 |
| Lemur catta | Ring-tailed lemur | 95 | 32 | 22 | 0.23157895 |
| Varecia variegata | Red/black ruffed lemur | 48 | 16 | 11 | 0.22916667 |
| Lycaon pictus | Cape hunting dog | 58 | 18 | 13 | 0.22413793 |
| Ursus maritimus | Polar bear | 26 | 8 | 5 | 0.19230769 |
| Didelphis marsupialis | Oppossum | 69 | 19 | 13 | 0.1884058 |
| Ovis aries | Sheep | 87 | 24 | 16 | 0.18390805 |
| Panthera tigris | White bengal tiger | 71 | 19 | 12 | 0.16901409 |
| Cynomys ludovicianus | Gunnison prairie dog | 137 | 39 | 23 | 0.16788321 |
| Octodon degus | Degu | 25 | 31 | 4 | 0.16 |

### Slide 7
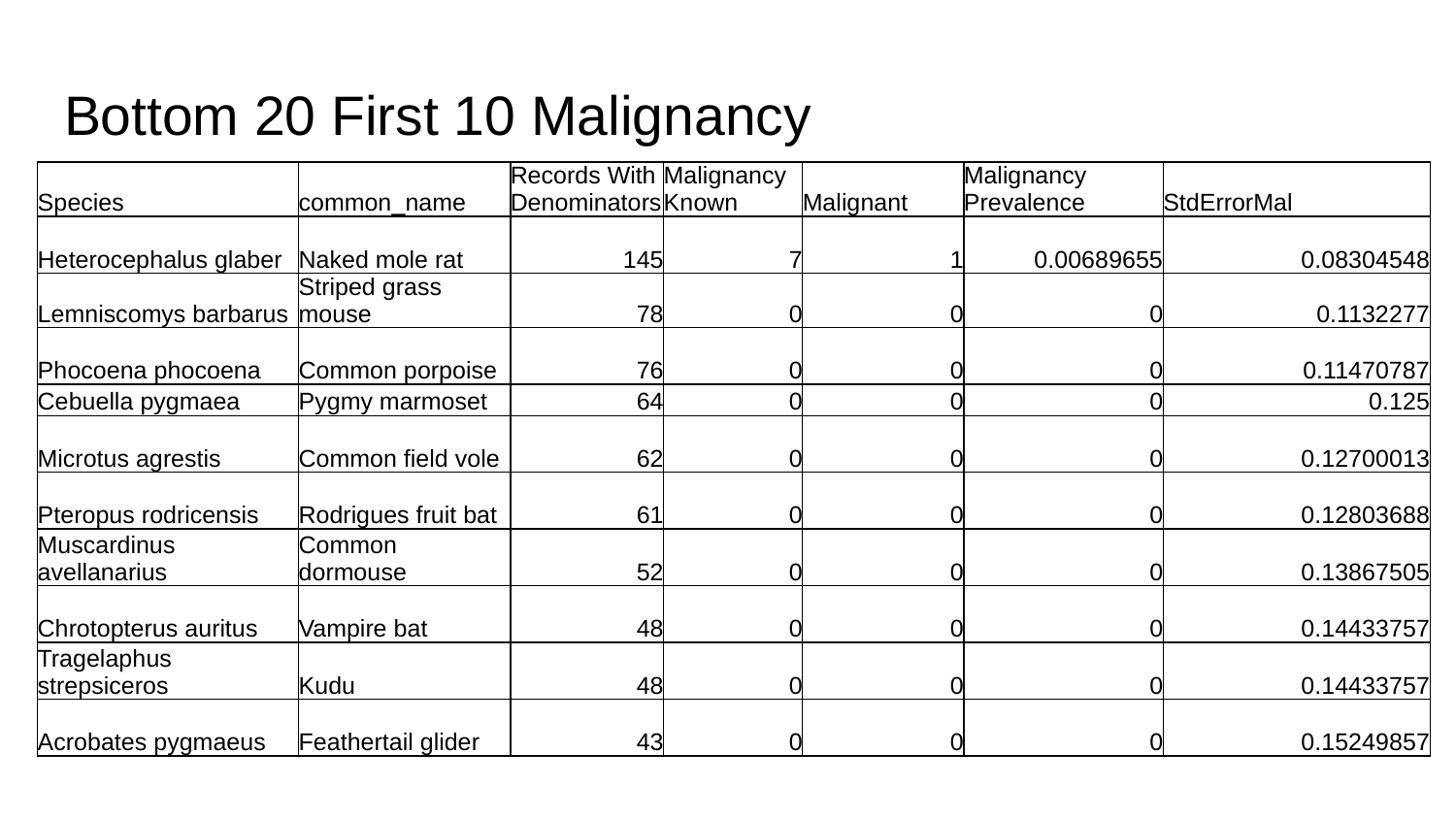

#### Bottom 20 First 10 Malignancy
| Species | common\_name | Records With Denominators | Malignancy Known | Malignant | Malignancy Prevalence | StdErrorMal |
| --- | --- | --- | --- | --- | --- | --- |
| Heterocephalus glaber | Naked mole rat | 145 | 7 | 1 | 0.00689655 | 0.08304548 |
| Lemniscomys barbarus | Striped grass mouse | 78 | 0 | 0 | 0 | 0.1132277 |
| Phocoena phocoena | Common porpoise | 76 | 0 | 0 | 0 | 0.11470787 |
| Cebuella pygmaea | Pygmy marmoset | 64 | 0 | 0 | 0 | 0.125 |
| Microtus agrestis | Common field vole | 62 | 0 | 0 | 0 | 0.12700013 |
| Pteropus rodricensis | Rodrigues fruit bat | 61 | 0 | 0 | 0 | 0.12803688 |
| Muscardinus avellanarius | Common dormouse | 52 | 0 | 0 | 0 | 0.13867505 |
| Chrotopterus auritus | Vampire bat | 48 | 0 | 0 | 0 | 0.14433757 |
| Tragelaphus strepsiceros | Kudu | 48 | 0 | 0 | 0 | 0.14433757 |
| Acrobates pygmaeus | Feathertail glider | 43 | 0 | 0 | 0 | 0.15249857 |

### Slide 8
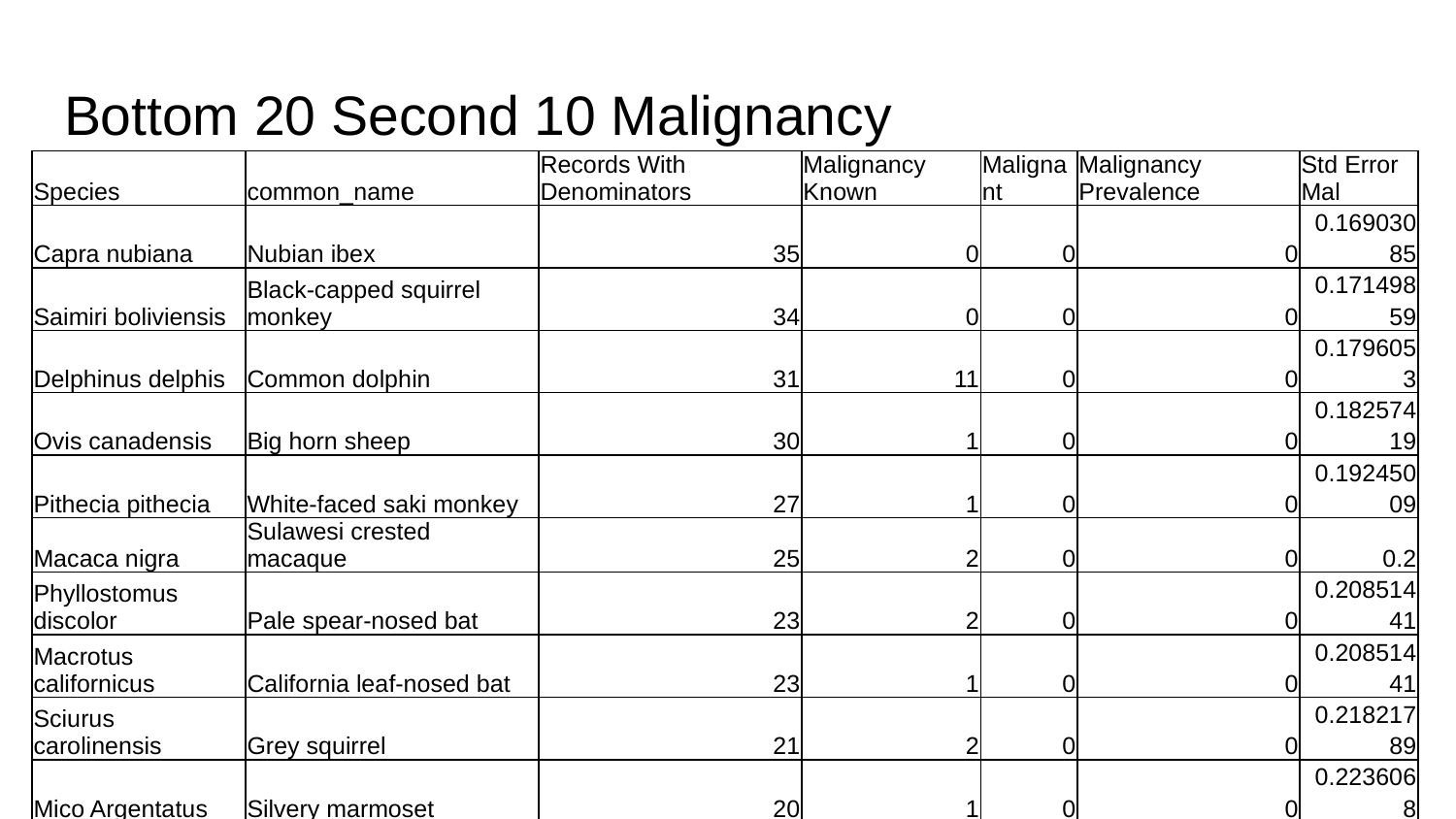

#### Bottom 20 Second 10 Malignancy
| Species | common\_name | Records With Denominators | Malignancy Known | Malignant | Malignancy Prevalence | Std Error Mal |
| --- | --- | --- | --- | --- | --- | --- |
| Capra nubiana | Nubian ibex | 35 | 0 | 0 | 0 | 0.16903085 |
| Saimiri boliviensis | Black-capped squirrel monkey | 34 | 0 | 0 | 0 | 0.17149859 |
| Delphinus delphis | Common dolphin | 31 | 11 | 0 | 0 | 0.1796053 |
| Ovis canadensis | Big horn sheep | 30 | 1 | 0 | 0 | 0.18257419 |
| Pithecia pithecia | White-faced saki monkey | 27 | 1 | 0 | 0 | 0.19245009 |
| Macaca nigra | Sulawesi crested macaque | 25 | 2 | 0 | 0 | 0.2 |
| Phyllostomus discolor | Pale spear-nosed bat | 23 | 2 | 0 | 0 | 0.20851441 |
| Macrotus californicus | California leaf-nosed bat | 23 | 1 | 0 | 0 | 0.20851441 |
| Sciurus carolinensis | Grey squirrel | 21 | 2 | 0 | 0 | 0.21821789 |
| Mico Argentatus | Silvery marmoset | 20 | 1 | 0 | 0 | 0.2236068 |
