## Supplemental figures, tables, and data dictionary for "Cancer Prevalence Across Vertebrates": ST2 min20 LeaderBoard.pptx

### Slide 1
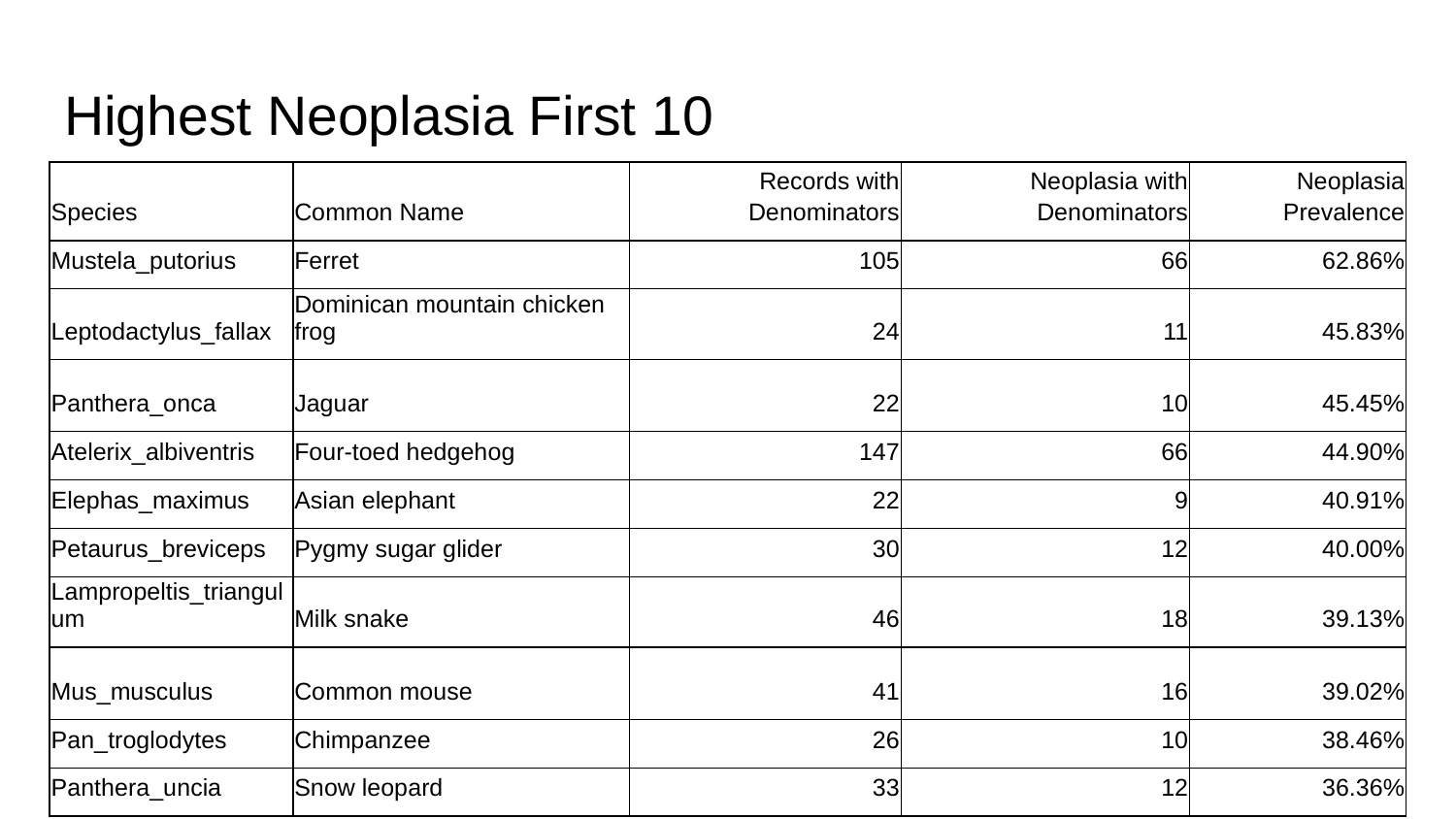

#### Highest Neoplasia First 10
| Species | Common Name | Records with Denominators | Neoplasia with Denominators | Neoplasia Prevalence |
| --- | --- | --- | --- | --- |
| Mustela\_putorius | Ferret | 105 | 66 | 62.86% |
| Leptodactylus\_fallax | Dominican mountain chicken frog | 24 | 11 | 45.83% |
| Panthera\_onca | Jaguar | 22 | 10 | 45.45% |
| Atelerix\_albiventris | Four-toed hedgehog | 147 | 66 | 44.90% |
| Elephas\_maximus | Asian elephant | 22 | 9 | 40.91% |
| Petaurus\_breviceps | Pygmy sugar glider | 30 | 12 | 40.00% |
| Lampropeltis\_triangulum | Milk snake | 46 | 18 | 39.13% |
| Mus\_musculus | Common mouse | 41 | 16 | 39.02% |
| Pan\_troglodytes | Chimpanzee | 26 | 10 | 38.46% |
| Panthera\_uncia | Snow leopard | 33 | 12 | 36.36% |

### Slide 2
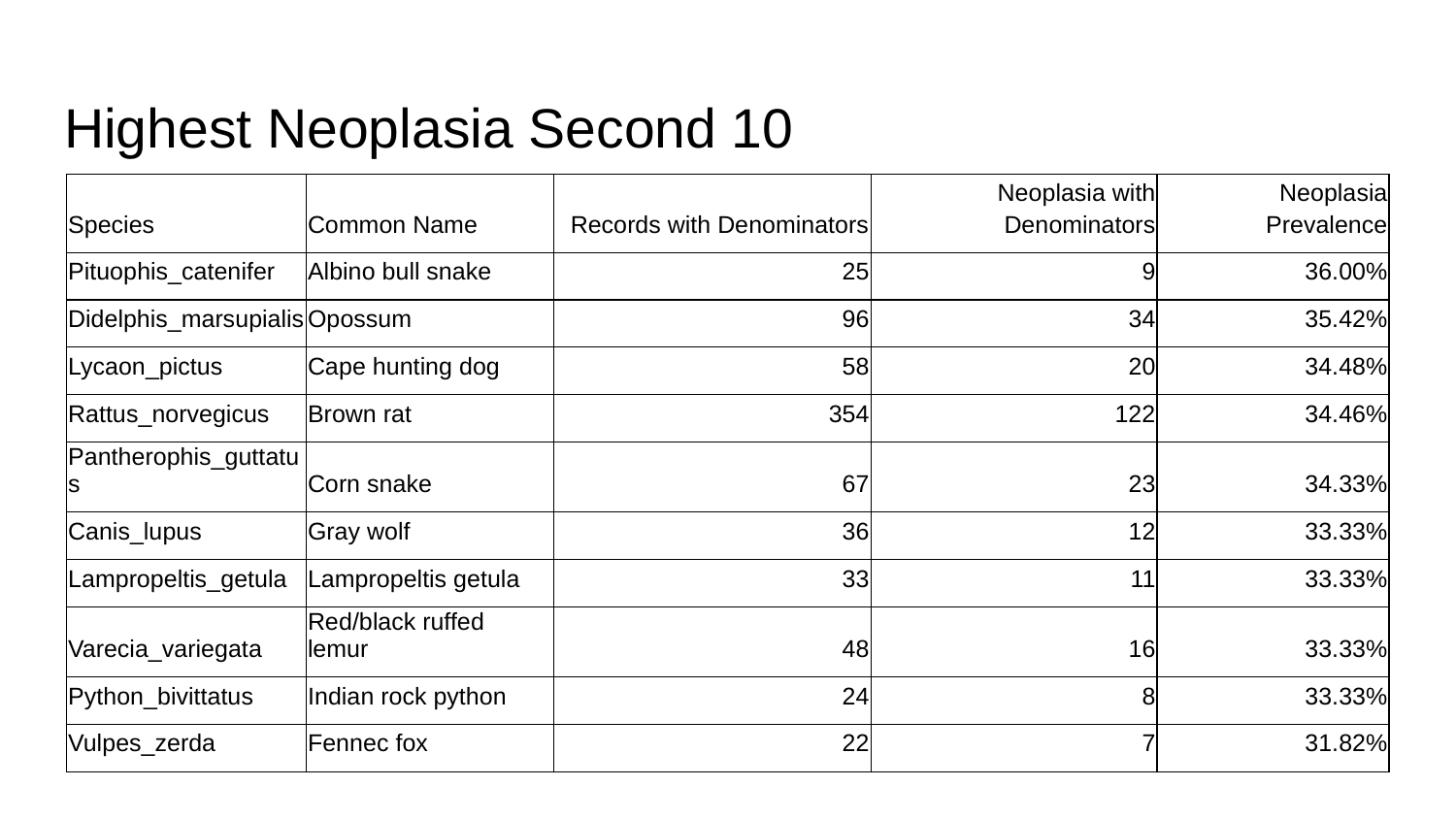

#### Highest Neoplasia Second 10
| Species | Common Name | Records with Denominators | Neoplasia with Denominators | Neoplasia Prevalence |
| --- | --- | --- | --- | --- |
| Pituophis\_catenifer | Albino bull snake | 25 | 9 | 36.00% |
| Didelphis\_marsupialis | Opossum | 96 | 34 | 35.42% |
| Lycaon\_pictus | Cape hunting dog | 58 | 20 | 34.48% |
| Rattus\_norvegicus | Brown rat | 354 | 122 | 34.46% |
| Pantherophis\_guttatus | Corn snake | 67 | 23 | 34.33% |
| Canis\_lupus | Gray wolf | 36 | 12 | 33.33% |
| Lampropeltis\_getula | Lampropeltis getula | 33 | 11 | 33.33% |
| Varecia\_variegata | Red/black ruffed lemur | 48 | 16 | 33.33% |
| Python\_bivittatus | Indian rock python | 24 | 8 | 33.33% |
| Vulpes\_zerda | Fennec fox | 22 | 7 | 31.82% |

### Slide 3
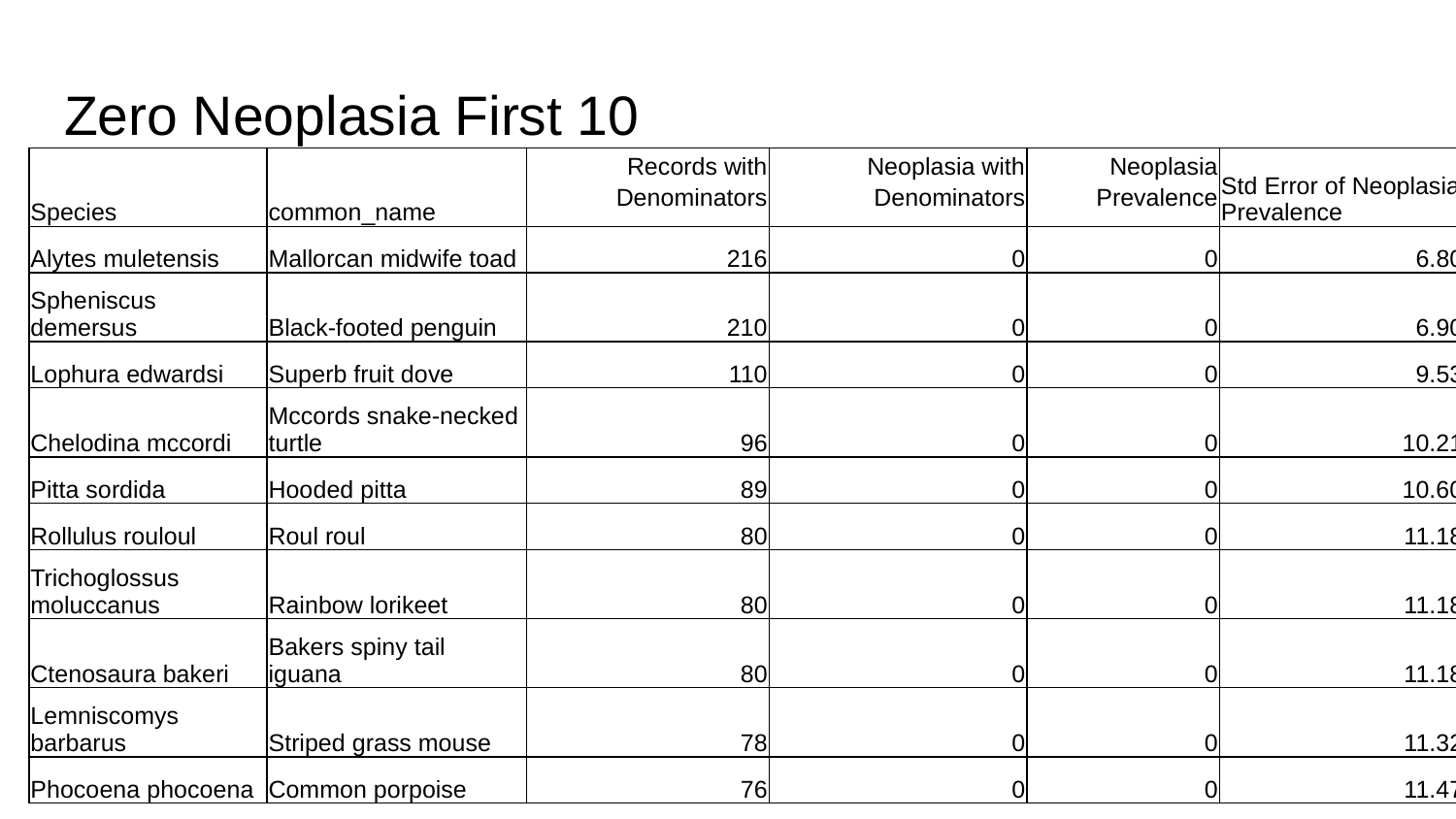

#### Zero Neoplasia First 10
| Species | common\_name | Records with Denominators | Neoplasia with Denominators | Neoplasia Prevalence | Std Error of Neoplasia Prevalence |
| --- | --- | --- | --- | --- | --- |
| Alytes muletensis | Mallorcan midwife toad | 216 | 0 | 0 | 6.80% |
| Spheniscus demersus | Black-footed penguin | 210 | 0 | 0 | 6.90% |
| Lophura edwardsi | Superb fruit dove | 110 | 0 | 0 | 9.53% |
| Chelodina mccordi | Mccords snake-necked turtle | 96 | 0 | 0 | 10.21% |
| Pitta sordida | Hooded pitta | 89 | 0 | 0 | 10.60% |
| Rollulus rouloul | Roul roul | 80 | 0 | 0 | 11.18% |
| Trichoglossus moluccanus | Rainbow lorikeet | 80 | 0 | 0 | 11.18% |
| Ctenosaura bakeri | Bakers spiny tail iguana | 80 | 0 | 0 | 11.18% |
| Lemniscomys barbarus | Striped grass mouse | 78 | 0 | 0 | 11.32% |
| Phocoena phocoena | Common porpoise | 76 | 0 | 0 | 11.47% |

### Slide 4
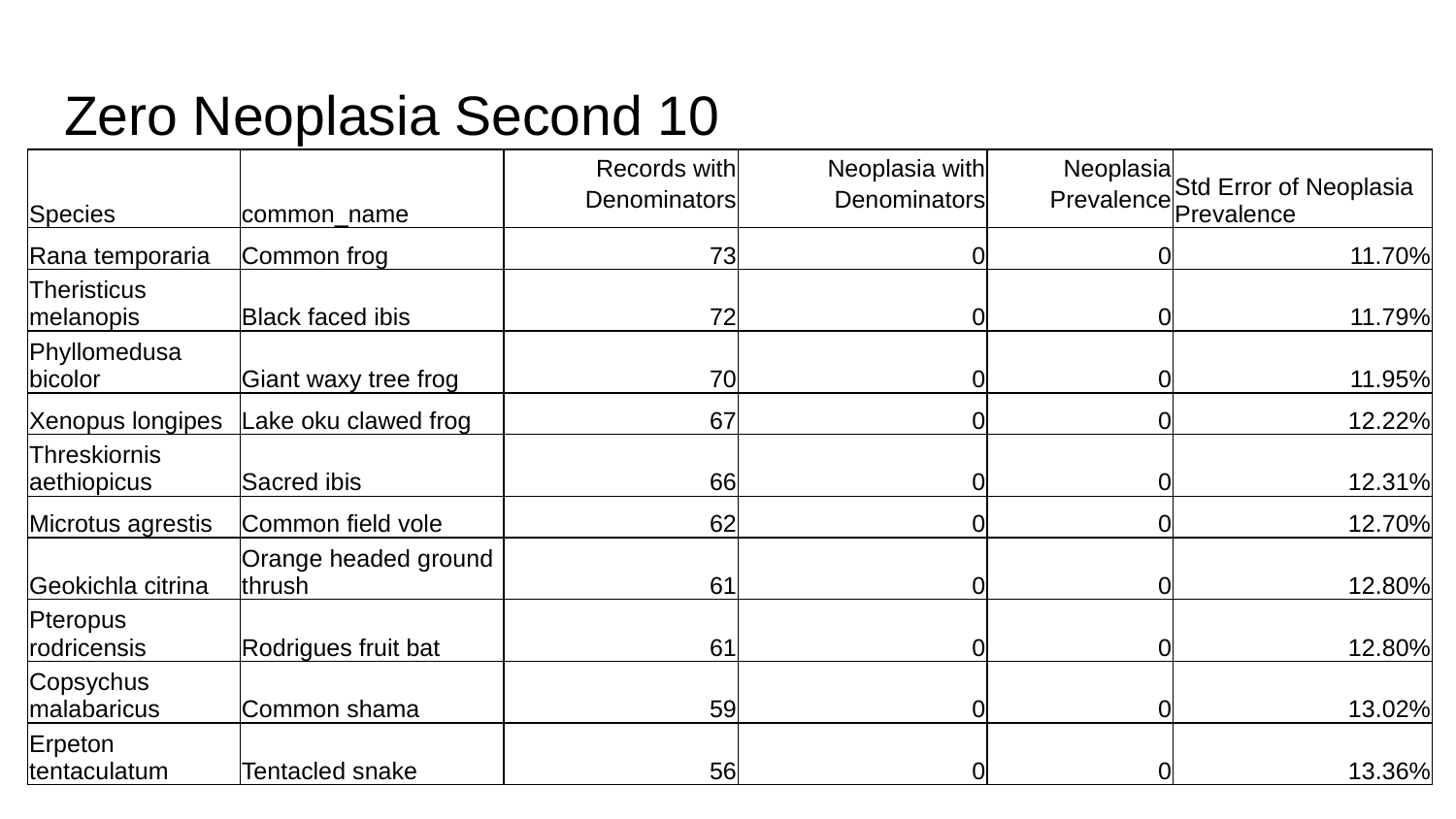

#### Zero Neoplasia Second 10
| Species | common\_name | Records with Denominators | Neoplasia with Denominators | Neoplasia Prevalence | Std Error of Neoplasia Prevalence |
| --- | --- | --- | --- | --- | --- |
| Rana temporaria | Common frog | 73 | 0 | 0 | 11.70% |
| Theristicus melanopis | Black faced ibis | 72 | 0 | 0 | 11.79% |
| Phyllomedusa bicolor | Giant waxy tree frog | 70 | 0 | 0 | 11.95% |
| Xenopus longipes | Lake oku clawed frog | 67 | 0 | 0 | 12.22% |
| Threskiornis aethiopicus | Sacred ibis | 66 | 0 | 0 | 12.31% |
| Microtus agrestis | Common field vole | 62 | 0 | 0 | 12.70% |
| Geokichla citrina | Orange headed ground thrush | 61 | 0 | 0 | 12.80% |
| Pteropus rodricensis | Rodrigues fruit bat | 61 | 0 | 0 | 12.80% |
| Copsychus malabaricus | Common shama | 59 | 0 | 0 | 13.02% |
| Erpeton tentaculatum | Tentacled snake | 56 | 0 | 0 | 13.36% |

### Slide 5
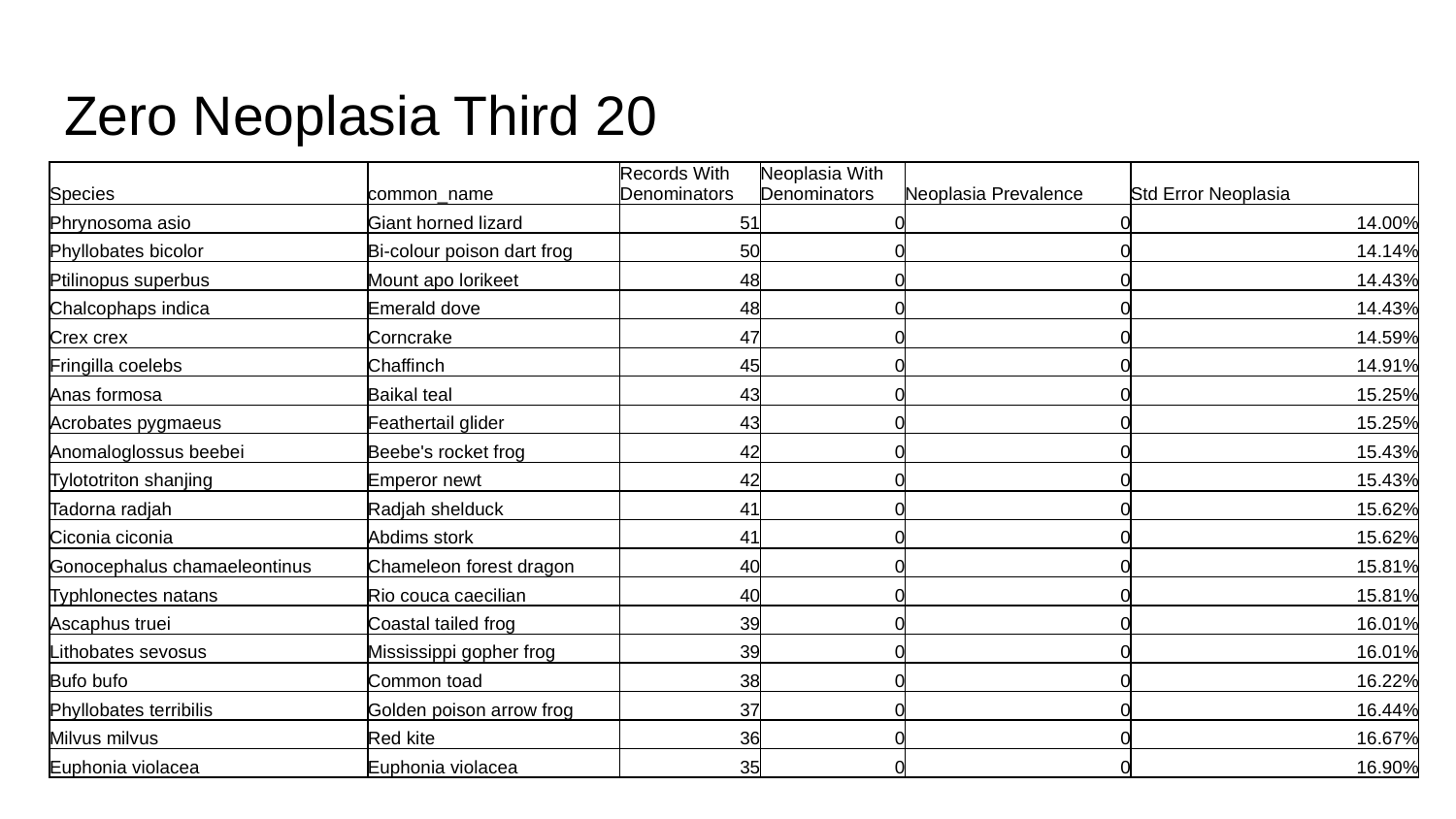

#### Zero Neoplasia Third 20
| Species | common\_name | Records With Denominators | Neoplasia With Denominators | Neoplasia Prevalence | Std Error Neoplasia |
| --- | --- | --- | --- | --- | --- |
| Phrynosoma asio | Giant horned lizard | 51 | 0 | 0 | 14.00% |
| Phyllobates bicolor | Bi-colour poison dart frog | 50 | 0 | 0 | 14.14% |
| Ptilinopus superbus | Mount apo lorikeet | 48 | 0 | 0 | 14.43% |
| Chalcophaps indica | Emerald dove | 48 | 0 | 0 | 14.43% |
| Crex crex | Corncrake | 47 | 0 | 0 | 14.59% |
| Fringilla coelebs | Chaffinch | 45 | 0 | 0 | 14.91% |
| Anas formosa | Baikal teal | 43 | 0 | 0 | 15.25% |
| Acrobates pygmaeus | Feathertail glider | 43 | 0 | 0 | 15.25% |
| Anomaloglossus beebei | Beebe's rocket frog | 42 | 0 | 0 | 15.43% |
| Tylototriton shanjing | Emperor newt | 42 | 0 | 0 | 15.43% |
| Tadorna radjah | Radjah shelduck | 41 | 0 | 0 | 15.62% |
| Ciconia ciconia | Abdims stork | 41 | 0 | 0 | 15.62% |
| Gonocephalus chamaeleontinus | Chameleon forest dragon | 40 | 0 | 0 | 15.81% |
| Typhlonectes natans | Rio couca caecilian | 40 | 0 | 0 | 15.81% |
| Ascaphus truei | Coastal tailed frog | 39 | 0 | 0 | 16.01% |
| Lithobates sevosus | Mississippi gopher frog | 39 | 0 | 0 | 16.01% |
| Bufo bufo | Common toad | 38 | 0 | 0 | 16.22% |
| Phyllobates terribilis | Golden poison arrow frog | 37 | 0 | 0 | 16.44% |
| Milvus milvus | Red kite | 36 | 0 | 0 | 16.67% |
| Euphonia violacea | Euphonia violacea | 35 | 0 | 0 | 16.90% |

### Slide 6
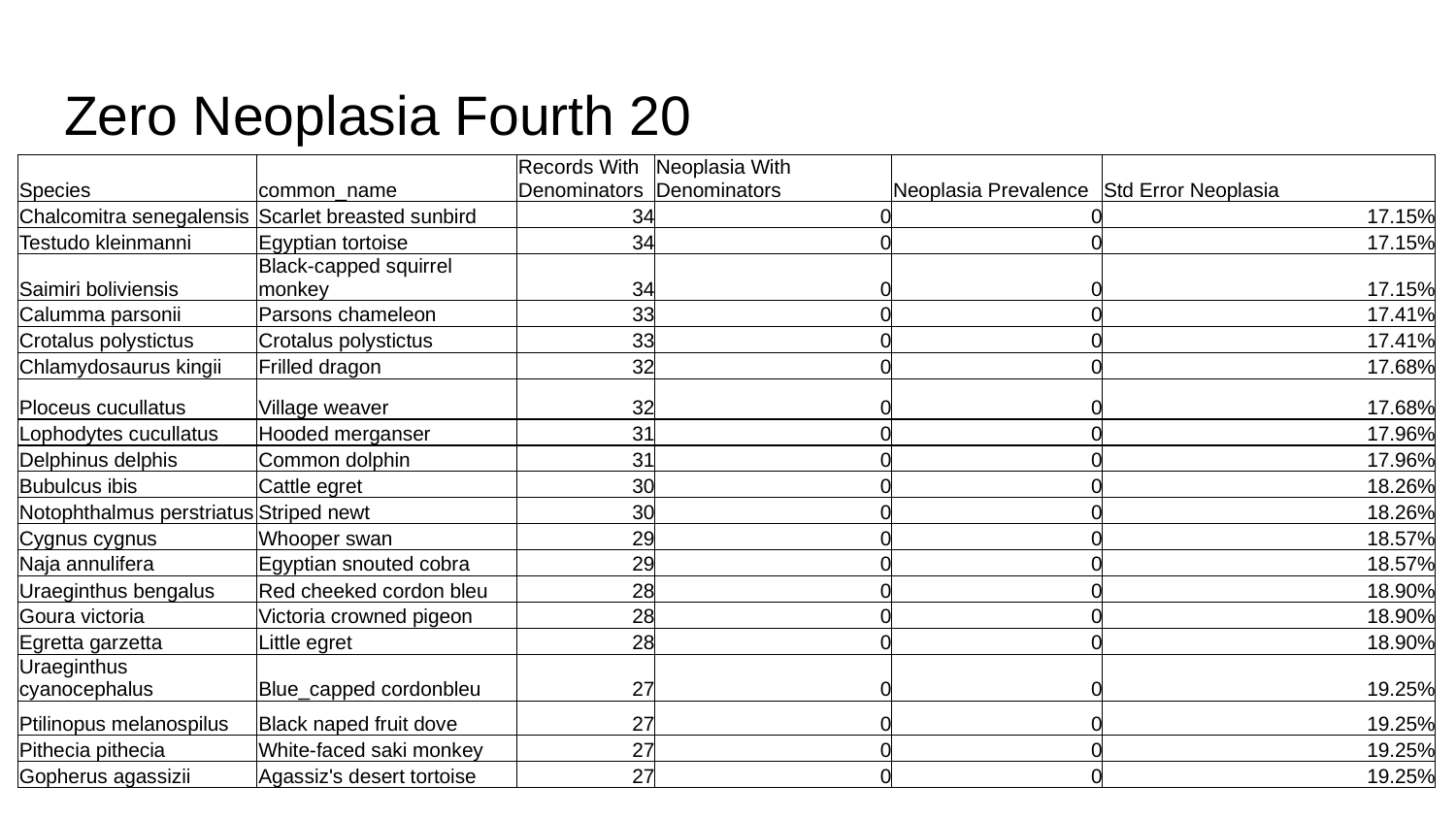

#### Zero Neoplasia Fourth 20
| Species | common\_name | Records With Denominators | Neoplasia With Denominators | Neoplasia Prevalence | Std Error Neoplasia |
| --- | --- | --- | --- | --- | --- |
| Chalcomitra senegalensis | Scarlet breasted sunbird | 34 | 0 | 0 | 17.15% |
| Testudo kleinmanni | Egyptian tortoise | 34 | 0 | 0 | 17.15% |
| Saimiri boliviensis | Black-capped squirrel monkey | 34 | 0 | 0 | 17.15% |
| Calumma parsonii | Parsons chameleon | 33 | 0 | 0 | 17.41% |
| Crotalus polystictus | Crotalus polystictus | 33 | 0 | 0 | 17.41% |
| Chlamydosaurus kingii | Frilled dragon | 32 | 0 | 0 | 17.68% |
| Ploceus cucullatus | Village weaver | 32 | 0 | 0 | 17.68% |
| Lophodytes cucullatus | Hooded merganser | 31 | 0 | 0 | 17.96% |
| Delphinus delphis | Common dolphin | 31 | 0 | 0 | 17.96% |
| Bubulcus ibis | Cattle egret | 30 | 0 | 0 | 18.26% |
| Notophthalmus perstriatus | Striped newt | 30 | 0 | 0 | 18.26% |
| Cygnus cygnus | Whooper swan | 29 | 0 | 0 | 18.57% |
| Naja annulifera | Egyptian snouted cobra | 29 | 0 | 0 | 18.57% |
| Uraeginthus bengalus | Red cheeked cordon bleu | 28 | 0 | 0 | 18.90% |
| Goura victoria | Victoria crowned pigeon | 28 | 0 | 0 | 18.90% |
| Egretta garzetta | Little egret | 28 | 0 | 0 | 18.90% |
| Uraeginthus cyanocephalus | Blue\_capped cordonbleu | 27 | 0 | 0 | 19.25% |
| Ptilinopus melanospilus | Black naped fruit dove | 27 | 0 | 0 | 19.25% |
| Pithecia pithecia | White-faced saki monkey | 27 | 0 | 0 | 19.25% |
| Gopherus agassizii | Agassiz's desert tortoise | 27 | 0 | 0 | 19.25% |

### Slide 7
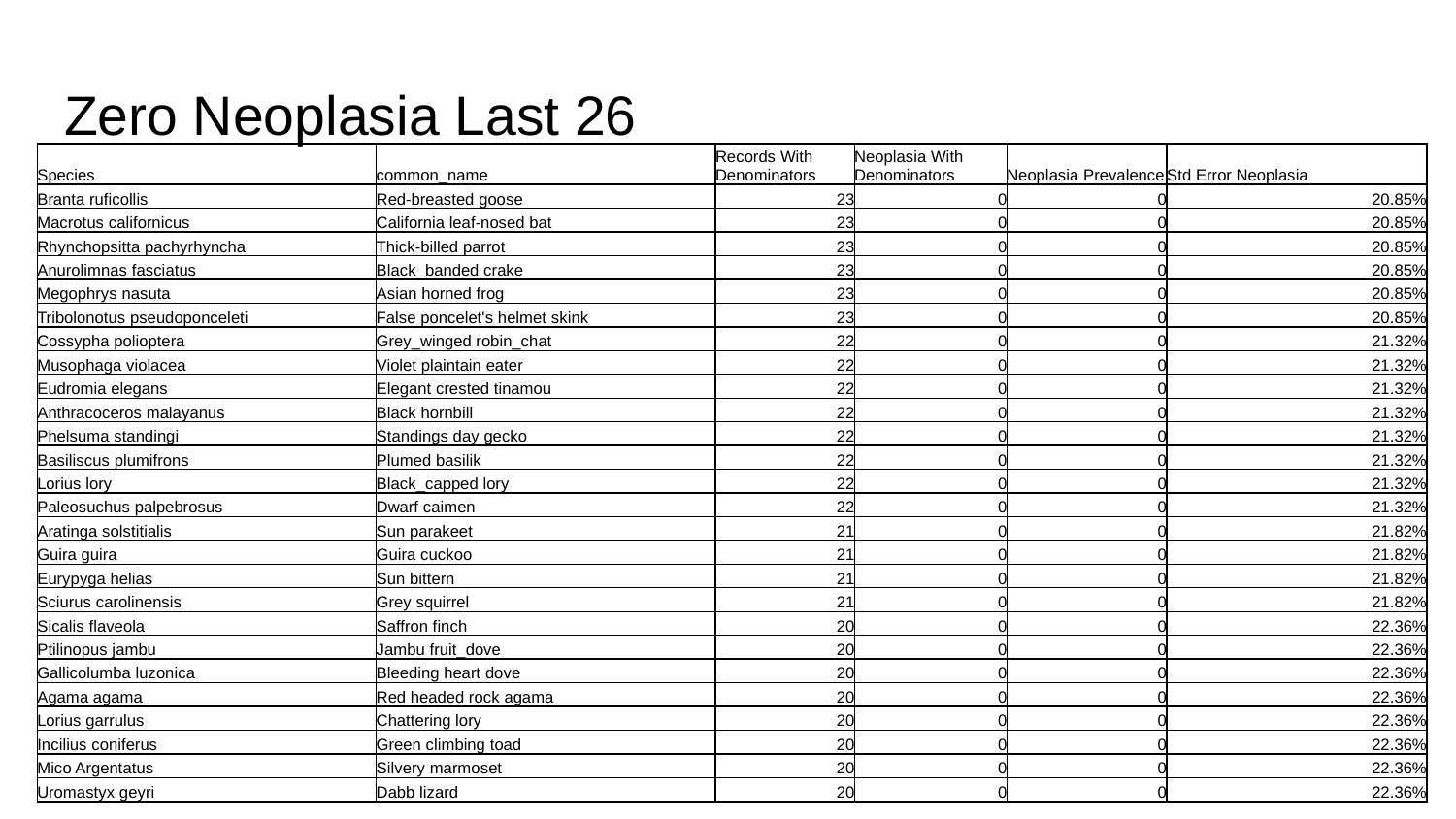

#### Zero Neoplasia Last 26
| Species | common\_name | Records With Denominators | Neoplasia With Denominators | Neoplasia Prevalence | Std Error Neoplasia |
| --- | --- | --- | --- | --- | --- |
| Branta ruficollis | Red-breasted goose | 23 | 0 | 0 | 20.85% |
| Macrotus californicus | California leaf-nosed bat | 23 | 0 | 0 | 20.85% |
| Rhynchopsitta pachyrhyncha | Thick-billed parrot | 23 | 0 | 0 | 20.85% |
| Anurolimnas fasciatus | Black\_banded crake | 23 | 0 | 0 | 20.85% |
| Megophrys nasuta | Asian horned frog | 23 | 0 | 0 | 20.85% |
| Tribolonotus pseudoponceleti | False poncelet's helmet skink | 23 | 0 | 0 | 20.85% |
| Cossypha polioptera | Grey\_winged robin\_chat | 22 | 0 | 0 | 21.32% |
| Musophaga violacea | Violet plaintain eater | 22 | 0 | 0 | 21.32% |
| Eudromia elegans | Elegant crested tinamou | 22 | 0 | 0 | 21.32% |
| Anthracoceros malayanus | Black hornbill | 22 | 0 | 0 | 21.32% |
| Phelsuma standingi | Standings day gecko | 22 | 0 | 0 | 21.32% |
| Basiliscus plumifrons | Plumed basilik | 22 | 0 | 0 | 21.32% |
| Lorius lory | Black\_capped lory | 22 | 0 | 0 | 21.32% |
| Paleosuchus palpebrosus | Dwarf caimen | 22 | 0 | 0 | 21.32% |
| Aratinga solstitialis | Sun parakeet | 21 | 0 | 0 | 21.82% |
| Guira guira | Guira cuckoo | 21 | 0 | 0 | 21.82% |
| Eurypyga helias | Sun bittern | 21 | 0 | 0 | 21.82% |
| Sciurus carolinensis | Grey squirrel | 21 | 0 | 0 | 21.82% |
| Sicalis flaveola | Saffron finch | 20 | 0 | 0 | 22.36% |
| Ptilinopus jambu | Jambu fruit\_dove | 20 | 0 | 0 | 22.36% |
| Gallicolumba luzonica | Bleeding heart dove | 20 | 0 | 0 | 22.36% |
| Agama agama | Red headed rock agama | 20 | 0 | 0 | 22.36% |
| Lorius garrulus | Chattering lory | 20 | 0 | 0 | 22.36% |
| Incilius coniferus | Green climbing toad | 20 | 0 | 0 | 22.36% |
| Mico Argentatus | Silvery marmoset | 20 | 0 | 0 | 22.36% |
| Uromastyx geyri | Dabb lizard | 20 | 0 | 0 | 22.36% |

### Slide 8
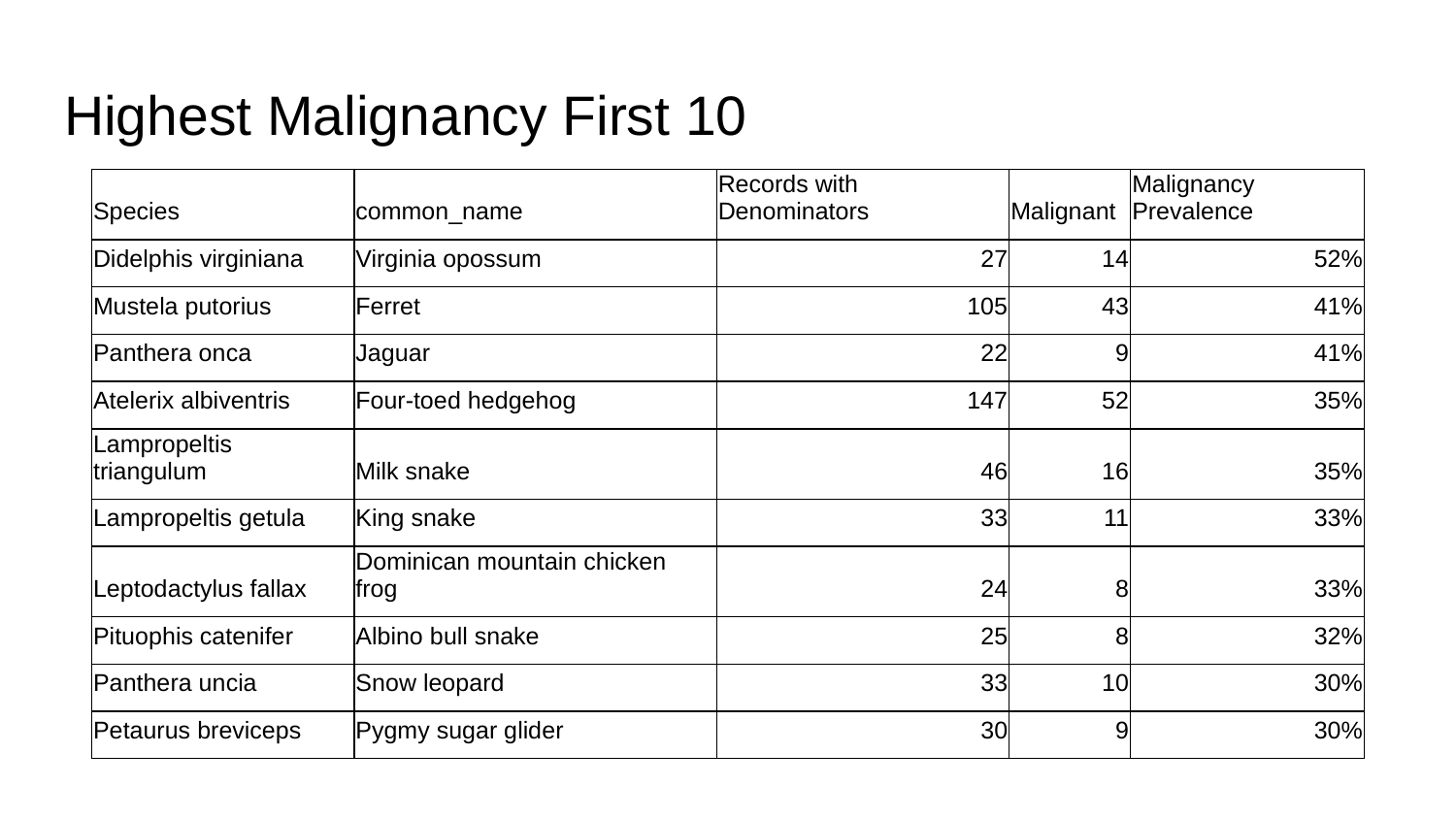

#### Highest Malignancy First 10
| Species | common\_name | Records with Denominators | Malignant | Malignancy Prevalence |
| --- | --- | --- | --- | --- |
| Didelphis virginiana | Virginia opossum | 27 | 14 | 52% |
| Mustela putorius | Ferret | 105 | 43 | 41% |
| Panthera onca | Jaguar | 22 | 9 | 41% |
| Atelerix albiventris | Four-toed hedgehog | 147 | 52 | 35% |
| Lampropeltis triangulum | Milk snake | 46 | 16 | 35% |
| Lampropeltis getula | King snake | 33 | 11 | 33% |
| Leptodactylus fallax | Dominican mountain chicken frog | 24 | 8 | 33% |
| Pituophis catenifer | Albino bull snake | 25 | 8 | 32% |
| Panthera uncia | Snow leopard | 33 | 10 | 30% |
| Petaurus breviceps | Pygmy sugar glider | 30 | 9 | 30% |

### Slide 9
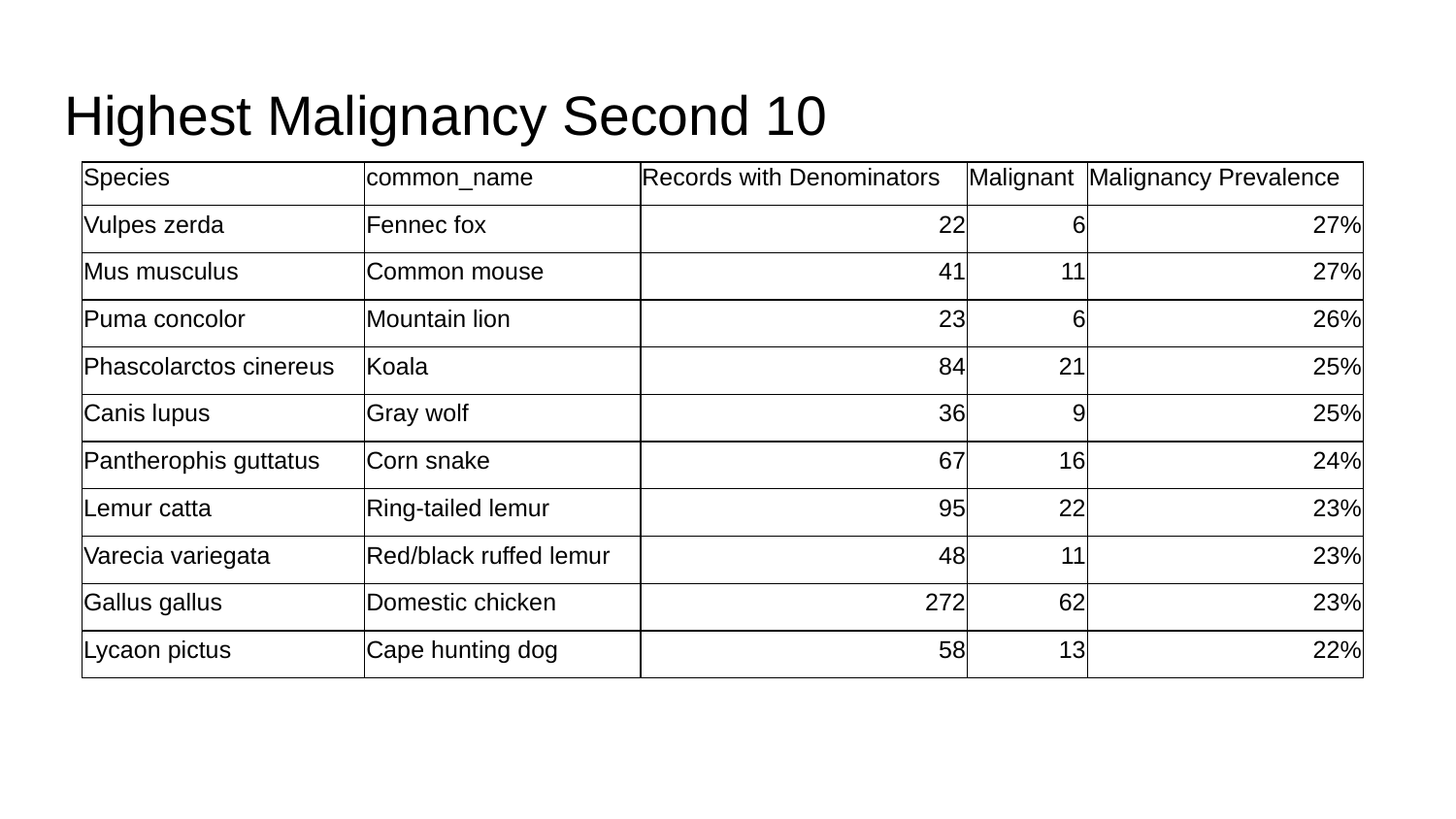

#### Highest Malignancy Second 10
| Species | common\_name | Records with Denominators | Malignant | Malignancy Prevalence |
| --- | --- | --- | --- | --- |
| Vulpes zerda | Fennec fox | 22 | 6 | 27% |
| Mus musculus | Common mouse | 41 | 11 | 27% |
| Puma concolor | Mountain lion | 23 | 6 | 26% |
| Phascolarctos cinereus | Koala | 84 | 21 | 25% |
| Canis lupus | Gray wolf | 36 | 9 | 25% |
| Pantherophis guttatus | Corn snake | 67 | 16 | 24% |
| Lemur catta | Ring-tailed lemur | 95 | 22 | 23% |
| Varecia variegata | Red/black ruffed lemur | 48 | 11 | 23% |
| Gallus gallus | Domestic chicken | 272 | 62 | 23% |
| Lycaon pictus | Cape hunting dog | 58 | 13 | 22% |

### Slide 10
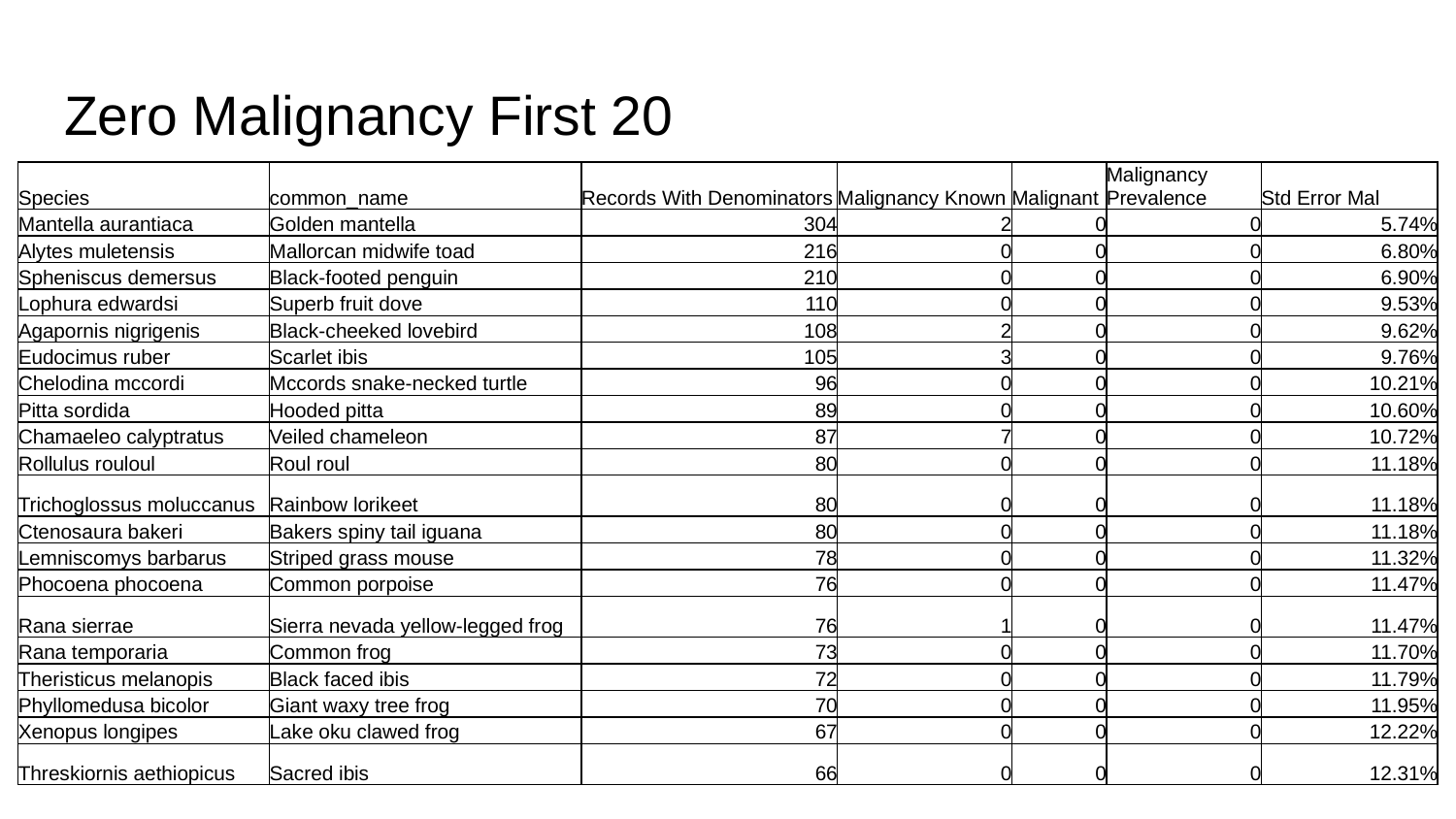

#### Zero Malignancy First 20
| Species | common\_name | Records With Denominators | Malignancy Known | Malignant | Malignancy Prevalence | Std Error Mal |
| --- | --- | --- | --- | --- | --- | --- |
| Mantella aurantiaca | Golden mantella | 304 | 2 | 0 | 0 | 5.74% |
| Alytes muletensis | Mallorcan midwife toad | 216 | 0 | 0 | 0 | 6.80% |
| Spheniscus demersus | Black-footed penguin | 210 | 0 | 0 | 0 | 6.90% |
| Lophura edwardsi | Superb fruit dove | 110 | 0 | 0 | 0 | 9.53% |
| Agapornis nigrigenis | Black-cheeked lovebird | 108 | 2 | 0 | 0 | 9.62% |
| Eudocimus ruber | Scarlet ibis | 105 | 3 | 0 | 0 | 9.76% |
| Chelodina mccordi | Mccords snake-necked turtle | 96 | 0 | 0 | 0 | 10.21% |
| Pitta sordida | Hooded pitta | 89 | 0 | 0 | 0 | 10.60% |
| Chamaeleo calyptratus | Veiled chameleon | 87 | 7 | 0 | 0 | 10.72% |
| Rollulus rouloul | Roul roul | 80 | 0 | 0 | 0 | 11.18% |
| Trichoglossus moluccanus | Rainbow lorikeet | 80 | 0 | 0 | 0 | 11.18% |
| Ctenosaura bakeri | Bakers spiny tail iguana | 80 | 0 | 0 | 0 | 11.18% |
| Lemniscomys barbarus | Striped grass mouse | 78 | 0 | 0 | 0 | 11.32% |
| Phocoena phocoena | Common porpoise | 76 | 0 | 0 | 0 | 11.47% |
| Rana sierrae | Sierra nevada yellow-legged frog | 76 | 1 | 0 | 0 | 11.47% |
| Rana temporaria | Common frog | 73 | 0 | 0 | 0 | 11.70% |
| Theristicus melanopis | Black faced ibis | 72 | 0 | 0 | 0 | 11.79% |
| Phyllomedusa bicolor | Giant waxy tree frog | 70 | 0 | 0 | 0 | 11.95% |
| Xenopus longipes | Lake oku clawed frog | 67 | 0 | 0 | 0 | 12.22% |
| Threskiornis aethiopicus | Sacred ibis | 66 | 0 | 0 | 0 | 12.31% |

### Slide 11
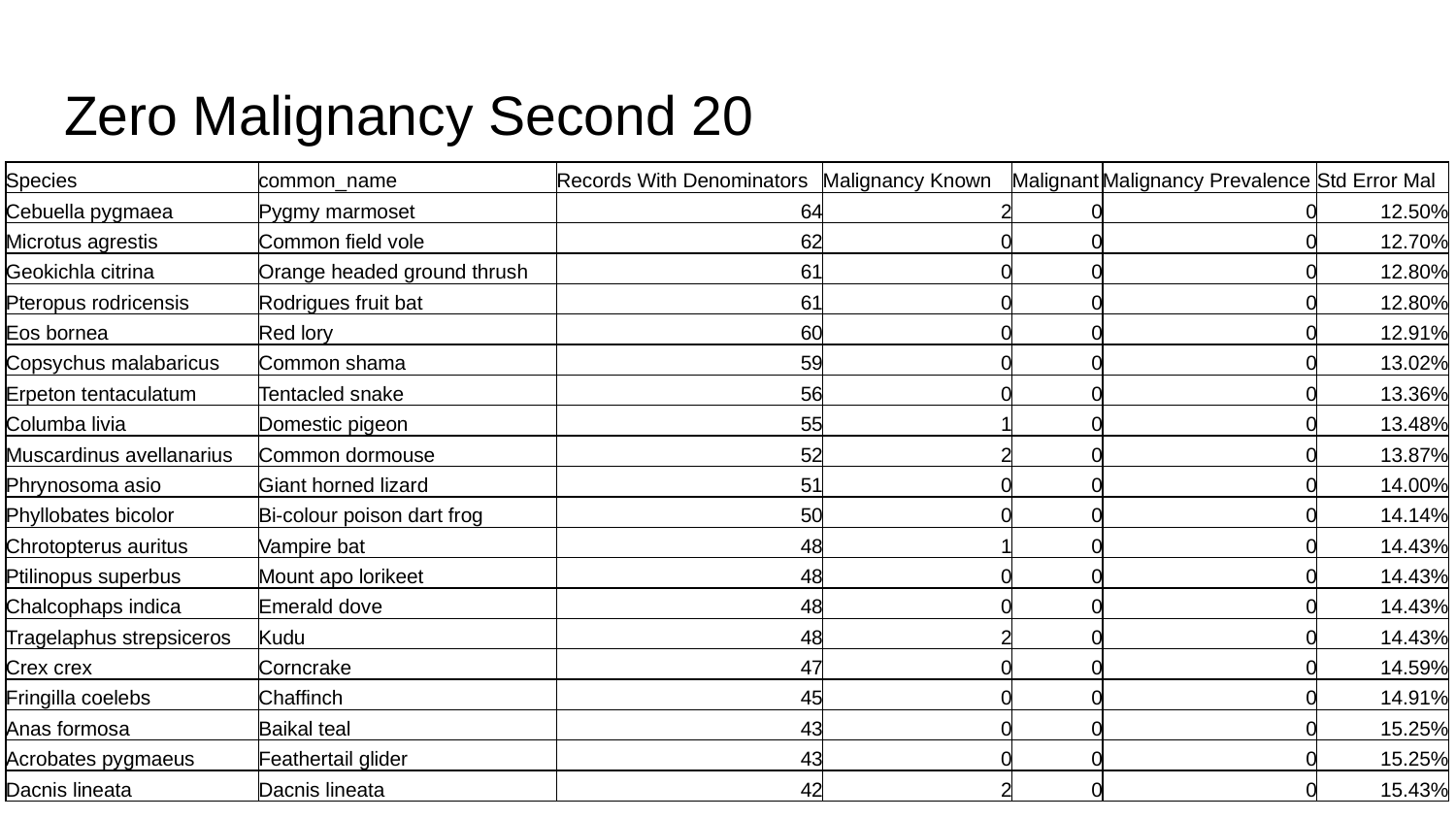

#### Zero Malignancy Second 20
| Species | common\_name | Records With Denominators | Malignancy Known | Malignant | Malignancy Prevalence | Std Error Mal |
| --- | --- | --- | --- | --- | --- | --- |
| Cebuella pygmaea | Pygmy marmoset | 64 | 2 | 0 | 0 | 12.50% |
| Microtus agrestis | Common field vole | 62 | 0 | 0 | 0 | 12.70% |
| Geokichla citrina | Orange headed ground thrush | 61 | 0 | 0 | 0 | 12.80% |
| Pteropus rodricensis | Rodrigues fruit bat | 61 | 0 | 0 | 0 | 12.80% |
| Eos bornea | Red lory | 60 | 0 | 0 | 0 | 12.91% |
| Copsychus malabaricus | Common shama | 59 | 0 | 0 | 0 | 13.02% |
| Erpeton tentaculatum | Tentacled snake | 56 | 0 | 0 | 0 | 13.36% |
| Columba livia | Domestic pigeon | 55 | 1 | 0 | 0 | 13.48% |
| Muscardinus avellanarius | Common dormouse | 52 | 2 | 0 | 0 | 13.87% |
| Phrynosoma asio | Giant horned lizard | 51 | 0 | 0 | 0 | 14.00% |
| Phyllobates bicolor | Bi-colour poison dart frog | 50 | 0 | 0 | 0 | 14.14% |
| Chrotopterus auritus | Vampire bat | 48 | 1 | 0 | 0 | 14.43% |
| Ptilinopus superbus | Mount apo lorikeet | 48 | 0 | 0 | 0 | 14.43% |
| Chalcophaps indica | Emerald dove | 48 | 0 | 0 | 0 | 14.43% |
| Tragelaphus strepsiceros | Kudu | 48 | 2 | 0 | 0 | 14.43% |
| Crex crex | Corncrake | 47 | 0 | 0 | 0 | 14.59% |
| Fringilla coelebs | Chaffinch | 45 | 0 | 0 | 0 | 14.91% |
| Anas formosa | Baikal teal | 43 | 0 | 0 | 0 | 15.25% |
| Acrobates pygmaeus | Feathertail glider | 43 | 0 | 0 | 0 | 15.25% |
| Dacnis lineata | Dacnis lineata | 42 | 2 | 0 | 0 | 15.43% |

### Slide 12
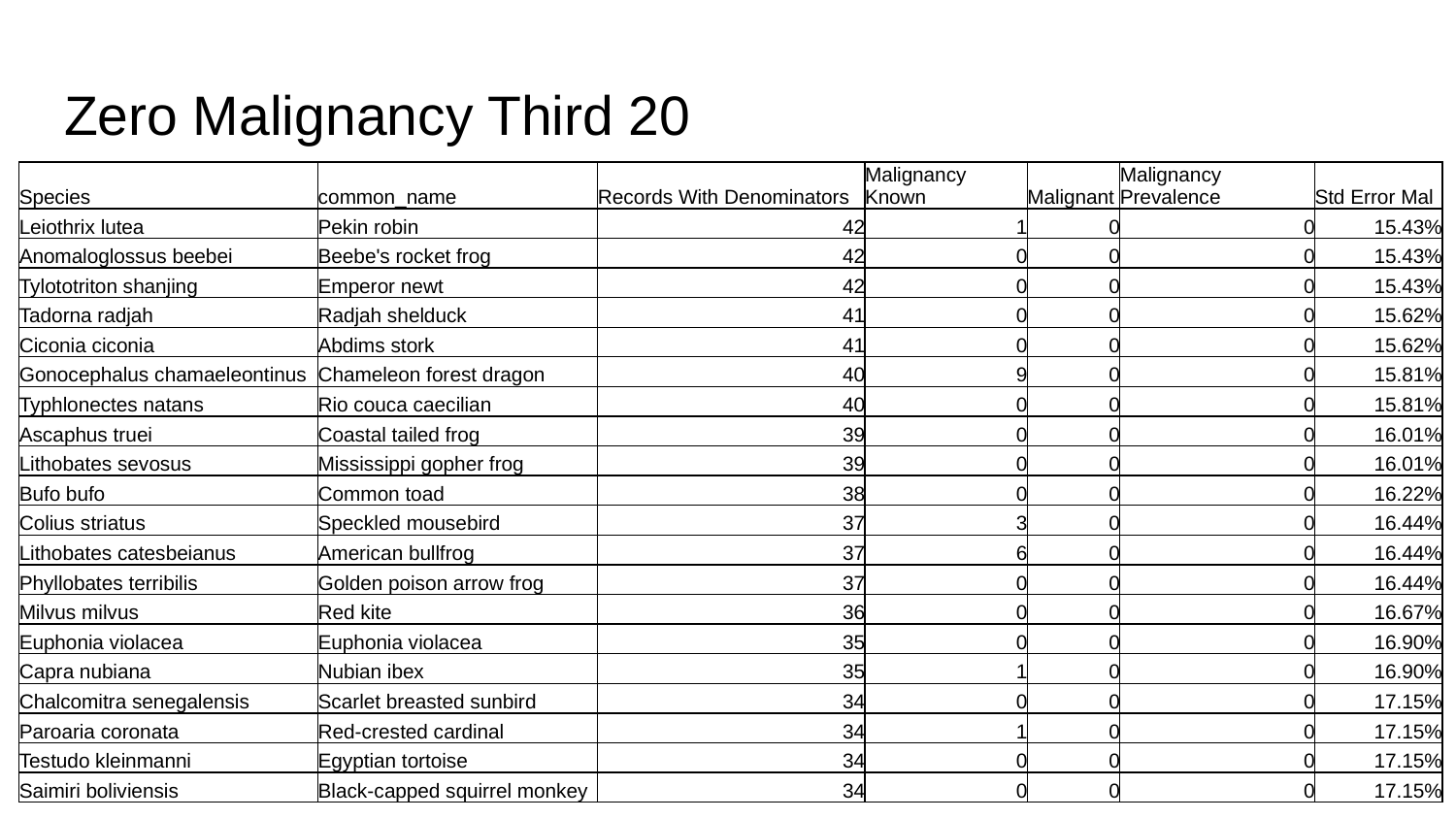

#### Zero Malignancy Third 20
| Species | common\_name | Records With Denominators | Malignancy Known | Malignant | Malignancy Prevalence | Std Error Mal |
| --- | --- | --- | --- | --- | --- | --- |
| Leiothrix lutea | Pekin robin | 42 | 1 | 0 | 0 | 15.43% |
| Anomaloglossus beebei | Beebe's rocket frog | 42 | 0 | 0 | 0 | 15.43% |
| Tylototriton shanjing | Emperor newt | 42 | 0 | 0 | 0 | 15.43% |
| Tadorna radjah | Radjah shelduck | 41 | 0 | 0 | 0 | 15.62% |
| Ciconia ciconia | Abdims stork | 41 | 0 | 0 | 0 | 15.62% |
| Gonocephalus chamaeleontinus | Chameleon forest dragon | 40 | 9 | 0 | 0 | 15.81% |
| Typhlonectes natans | Rio couca caecilian | 40 | 0 | 0 | 0 | 15.81% |
| Ascaphus truei | Coastal tailed frog | 39 | 0 | 0 | 0 | 16.01% |
| Lithobates sevosus | Mississippi gopher frog | 39 | 0 | 0 | 0 | 16.01% |
| Bufo bufo | Common toad | 38 | 0 | 0 | 0 | 16.22% |
| Colius striatus | Speckled mousebird | 37 | 3 | 0 | 0 | 16.44% |
| Lithobates catesbeianus | American bullfrog | 37 | 6 | 0 | 0 | 16.44% |
| Phyllobates terribilis | Golden poison arrow frog | 37 | 0 | 0 | 0 | 16.44% |
| Milvus milvus | Red kite | 36 | 0 | 0 | 0 | 16.67% |
| Euphonia violacea | Euphonia violacea | 35 | 0 | 0 | 0 | 16.90% |
| Capra nubiana | Nubian ibex | 35 | 1 | 0 | 0 | 16.90% |
| Chalcomitra senegalensis | Scarlet breasted sunbird | 34 | 0 | 0 | 0 | 17.15% |
| Paroaria coronata | Red-crested cardinal | 34 | 1 | 0 | 0 | 17.15% |
| Testudo kleinmanni | Egyptian tortoise | 34 | 0 | 0 | 0 | 17.15% |
| Saimiri boliviensis | Black-capped squirrel monkey | 34 | 0 | 0 | 0 | 17.15% |

### Slide 13
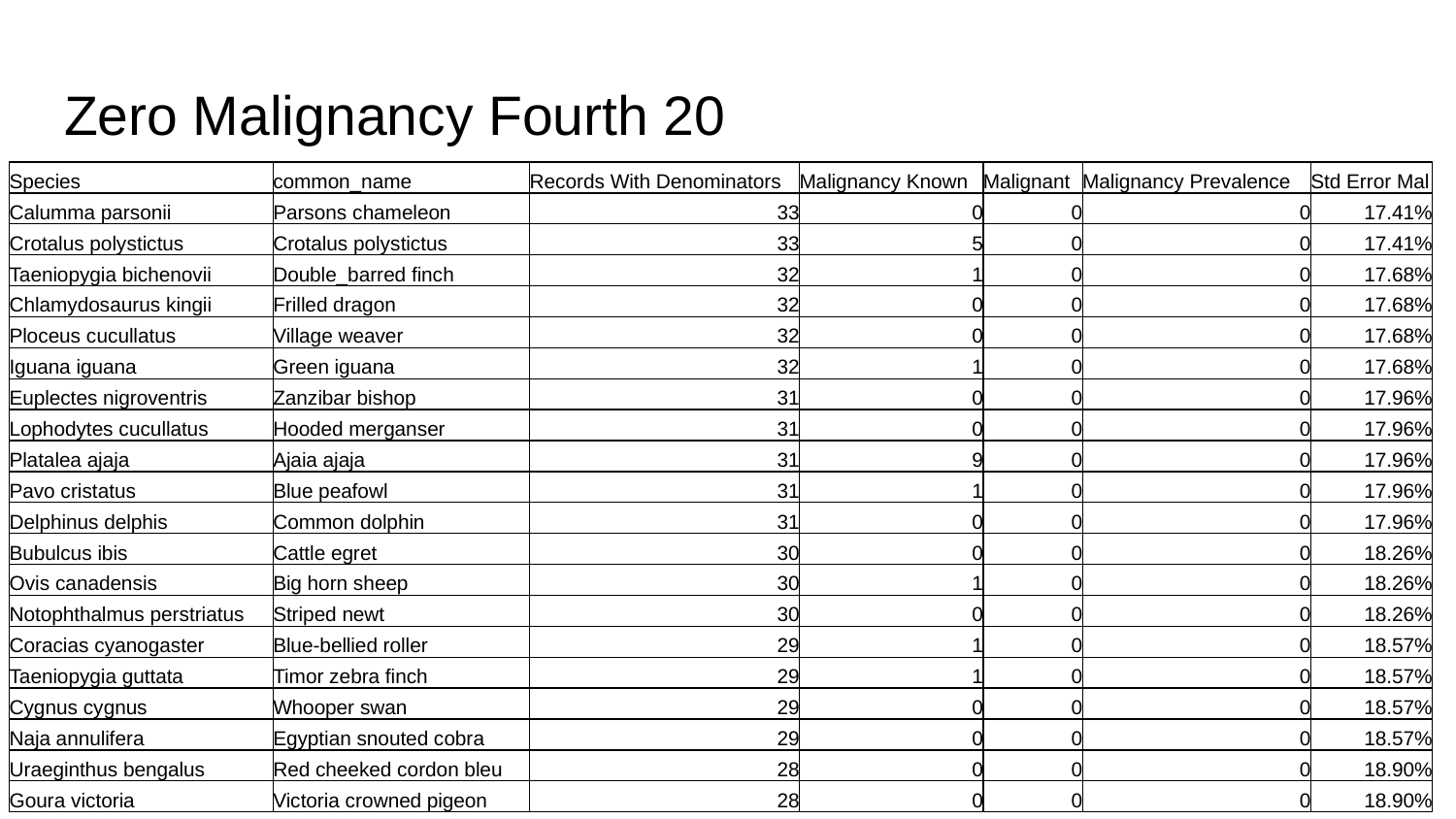

#### Zero Malignancy Fourth 20
| Species | common\_name | Records With Denominators | Malignancy Known | Malignant | Malignancy Prevalence | Std Error Mal |
| --- | --- | --- | --- | --- | --- | --- |
| Calumma parsonii | Parsons chameleon | 33 | 0 | 0 | 0 | 17.41% |
| Crotalus polystictus | Crotalus polystictus | 33 | 5 | 0 | 0 | 17.41% |
| Taeniopygia bichenovii | Double\_barred finch | 32 | 1 | 0 | 0 | 17.68% |
| Chlamydosaurus kingii | Frilled dragon | 32 | 0 | 0 | 0 | 17.68% |
| Ploceus cucullatus | Village weaver | 32 | 0 | 0 | 0 | 17.68% |
| Iguana iguana | Green iguana | 32 | 1 | 0 | 0 | 17.68% |
| Euplectes nigroventris | Zanzibar bishop | 31 | 0 | 0 | 0 | 17.96% |
| Lophodytes cucullatus | Hooded merganser | 31 | 0 | 0 | 0 | 17.96% |
| Platalea ajaja | Ajaia ajaja | 31 | 9 | 0 | 0 | 17.96% |
| Pavo cristatus | Blue peafowl | 31 | 1 | 0 | 0 | 17.96% |
| Delphinus delphis | Common dolphin | 31 | 0 | 0 | 0 | 17.96% |
| Bubulcus ibis | Cattle egret | 30 | 0 | 0 | 0 | 18.26% |
| Ovis canadensis | Big horn sheep | 30 | 1 | 0 | 0 | 18.26% |
| Notophthalmus perstriatus | Striped newt | 30 | 0 | 0 | 0 | 18.26% |
| Coracias cyanogaster | Blue-bellied roller | 29 | 1 | 0 | 0 | 18.57% |
| Taeniopygia guttata | Timor zebra finch | 29 | 1 | 0 | 0 | 18.57% |
| Cygnus cygnus | Whooper swan | 29 | 0 | 0 | 0 | 18.57% |
| Naja annulifera | Egyptian snouted cobra | 29 | 0 | 0 | 0 | 18.57% |
| Uraeginthus bengalus | Red cheeked cordon bleu | 28 | 0 | 0 | 0 | 18.90% |
| Goura victoria | Victoria crowned pigeon | 28 | 0 | 0 | 0 | 18.90% |

### Slide 14
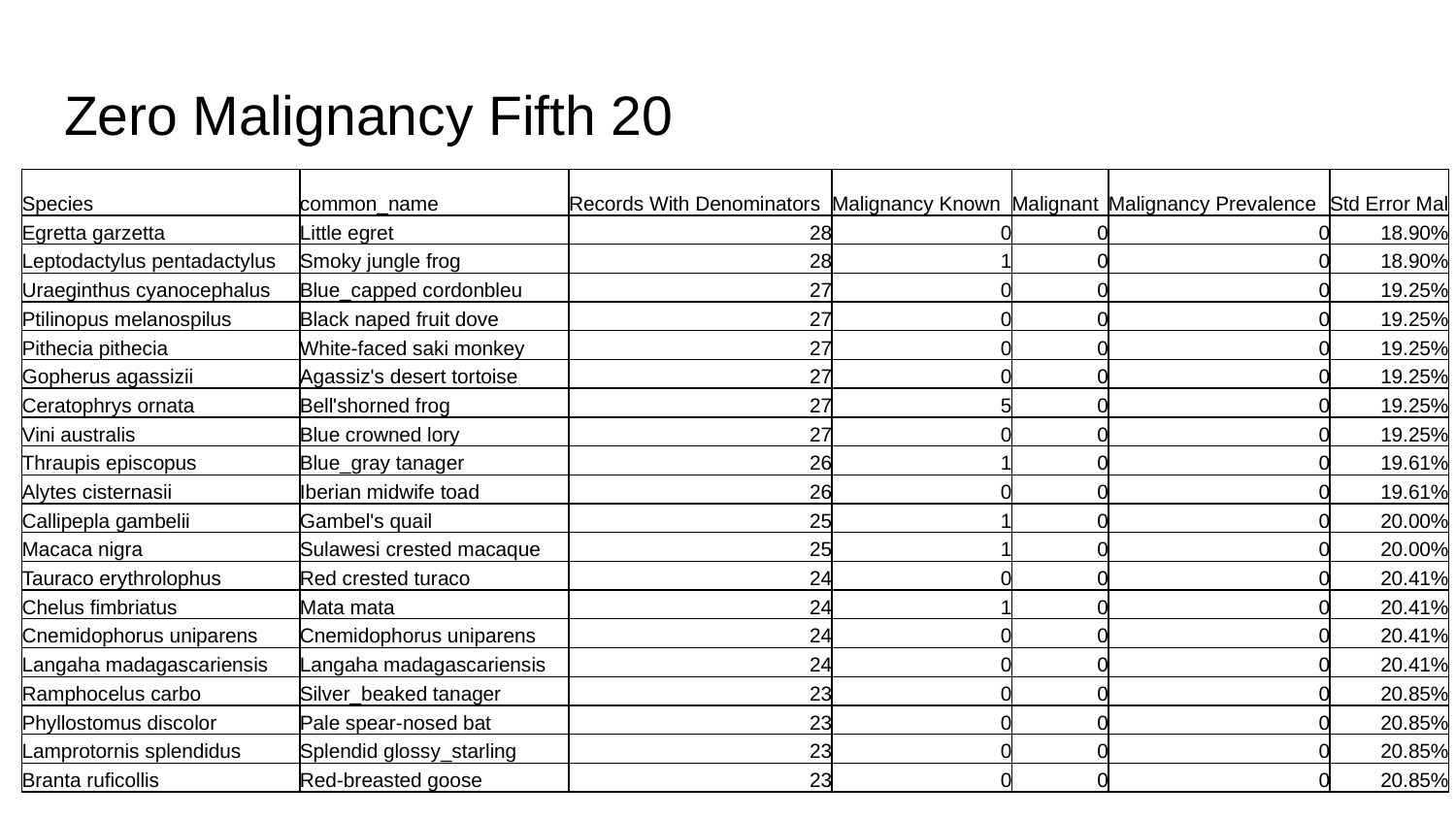

#### Zero Malignancy Fifth 20
| Species | common\_name | Records With Denominators | Malignancy Known | Malignant | Malignancy Prevalence | Std Error Mal |
| --- | --- | --- | --- | --- | --- | --- |
| Egretta garzetta | Little egret | 28 | 0 | 0 | 0 | 18.90% |
| Leptodactylus pentadactylus | Smoky jungle frog | 28 | 1 | 0 | 0 | 18.90% |
| Uraeginthus cyanocephalus | Blue\_capped cordonbleu | 27 | 0 | 0 | 0 | 19.25% |
| Ptilinopus melanospilus | Black naped fruit dove | 27 | 0 | 0 | 0 | 19.25% |
| Pithecia pithecia | White-faced saki monkey | 27 | 0 | 0 | 0 | 19.25% |
| Gopherus agassizii | Agassiz's desert tortoise | 27 | 0 | 0 | 0 | 19.25% |
| Ceratophrys ornata | Bell'shorned frog | 27 | 5 | 0 | 0 | 19.25% |
| Vini australis | Blue crowned lory | 27 | 0 | 0 | 0 | 19.25% |
| Thraupis episcopus | Blue\_gray tanager | 26 | 1 | 0 | 0 | 19.61% |
| Alytes cisternasii | Iberian midwife toad | 26 | 0 | 0 | 0 | 19.61% |
| Callipepla gambelii | Gambel's quail | 25 | 1 | 0 | 0 | 20.00% |
| Macaca nigra | Sulawesi crested macaque | 25 | 1 | 0 | 0 | 20.00% |
| Tauraco erythrolophus | Red crested turaco | 24 | 0 | 0 | 0 | 20.41% |
| Chelus fimbriatus | Mata mata | 24 | 1 | 0 | 0 | 20.41% |
| Cnemidophorus uniparens | Cnemidophorus uniparens | 24 | 0 | 0 | 0 | 20.41% |
| Langaha madagascariensis | Langaha madagascariensis | 24 | 0 | 0 | 0 | 20.41% |
| Ramphocelus carbo | Silver\_beaked tanager | 23 | 0 | 0 | 0 | 20.85% |
| Phyllostomus discolor | Pale spear-nosed bat | 23 | 0 | 0 | 0 | 20.85% |
| Lamprotornis splendidus | Splendid glossy\_starling | 23 | 0 | 0 | 0 | 20.85% |
| Branta ruficollis | Red-breasted goose | 23 | 0 | 0 | 0 | 20.85% |

### Slide 15
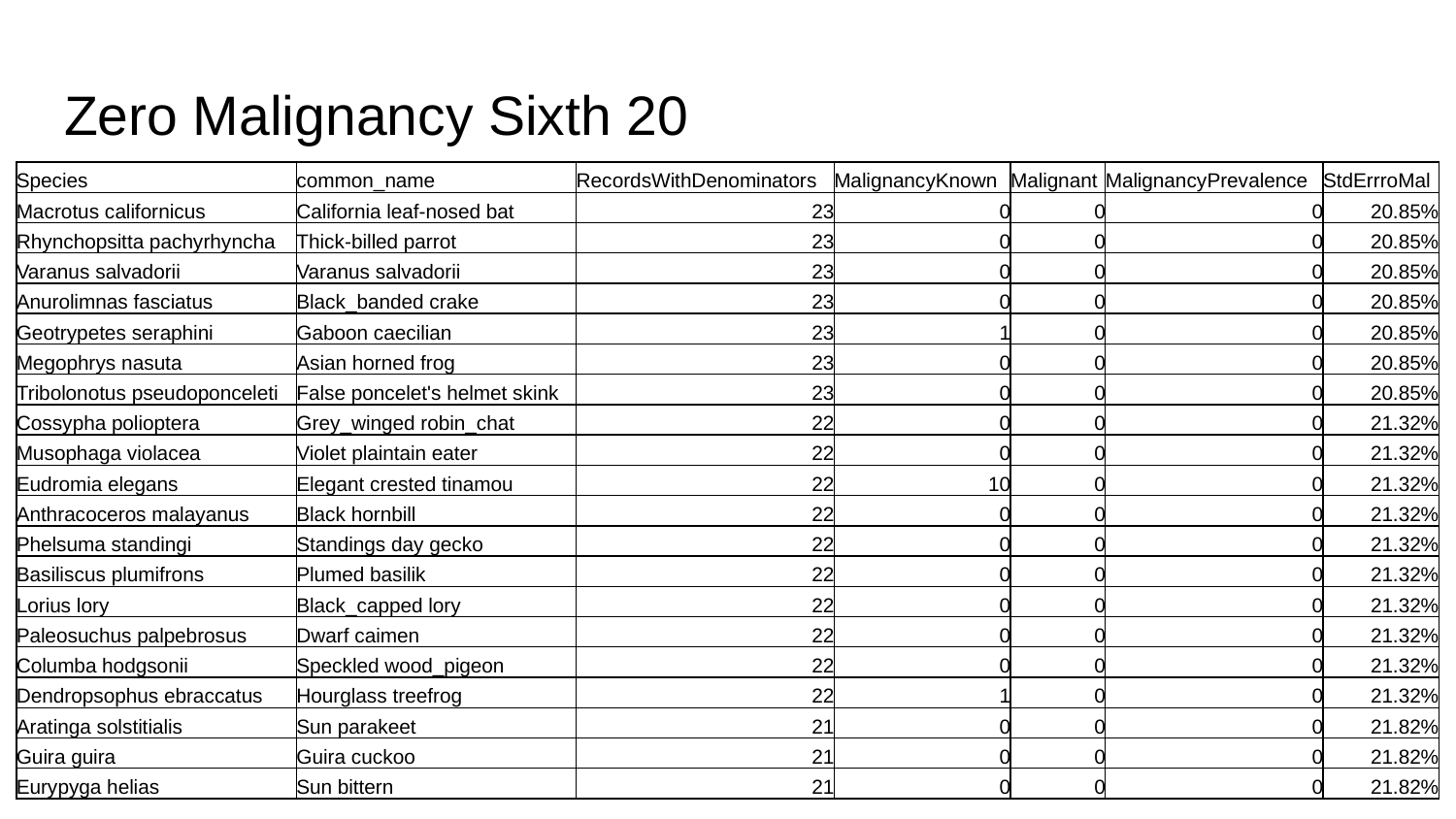

#### Zero Malignancy Sixth 20
| Species | common\_name | RecordsWithDenominators | MalignancyKnown | Malignant | MalignancyPrevalence | StdErrroMal |
| --- | --- | --- | --- | --- | --- | --- |
| Macrotus californicus | California leaf-nosed bat | 23 | 0 | 0 | 0 | 20.85% |
| Rhynchopsitta pachyrhyncha | Thick-billed parrot | 23 | 0 | 0 | 0 | 20.85% |
| Varanus salvadorii | Varanus salvadorii | 23 | 0 | 0 | 0 | 20.85% |
| Anurolimnas fasciatus | Black\_banded crake | 23 | 0 | 0 | 0 | 20.85% |
| Geotrypetes seraphini | Gaboon caecilian | 23 | 1 | 0 | 0 | 20.85% |
| Megophrys nasuta | Asian horned frog | 23 | 0 | 0 | 0 | 20.85% |
| Tribolonotus pseudoponceleti | False poncelet's helmet skink | 23 | 0 | 0 | 0 | 20.85% |
| Cossypha polioptera | Grey\_winged robin\_chat | 22 | 0 | 0 | 0 | 21.32% |
| Musophaga violacea | Violet plaintain eater | 22 | 0 | 0 | 0 | 21.32% |
| Eudromia elegans | Elegant crested tinamou | 22 | 10 | 0 | 0 | 21.32% |
| Anthracoceros malayanus | Black hornbill | 22 | 0 | 0 | 0 | 21.32% |
| Phelsuma standingi | Standings day gecko | 22 | 0 | 0 | 0 | 21.32% |
| Basiliscus plumifrons | Plumed basilik | 22 | 0 | 0 | 0 | 21.32% |
| Lorius lory | Black\_capped lory | 22 | 0 | 0 | 0 | 21.32% |
| Paleosuchus palpebrosus | Dwarf caimen | 22 | 0 | 0 | 0 | 21.32% |
| Columba hodgsonii | Speckled wood\_pigeon | 22 | 0 | 0 | 0 | 21.32% |
| Dendropsophus ebraccatus | Hourglass treefrog | 22 | 1 | 0 | 0 | 21.32% |
| Aratinga solstitialis | Sun parakeet | 21 | 0 | 0 | 0 | 21.82% |
| Guira guira | Guira cuckoo | 21 | 0 | 0 | 0 | 21.82% |
| Eurypyga helias | Sun bittern | 21 | 0 | 0 | 0 | 21.82% |

### Slide 16
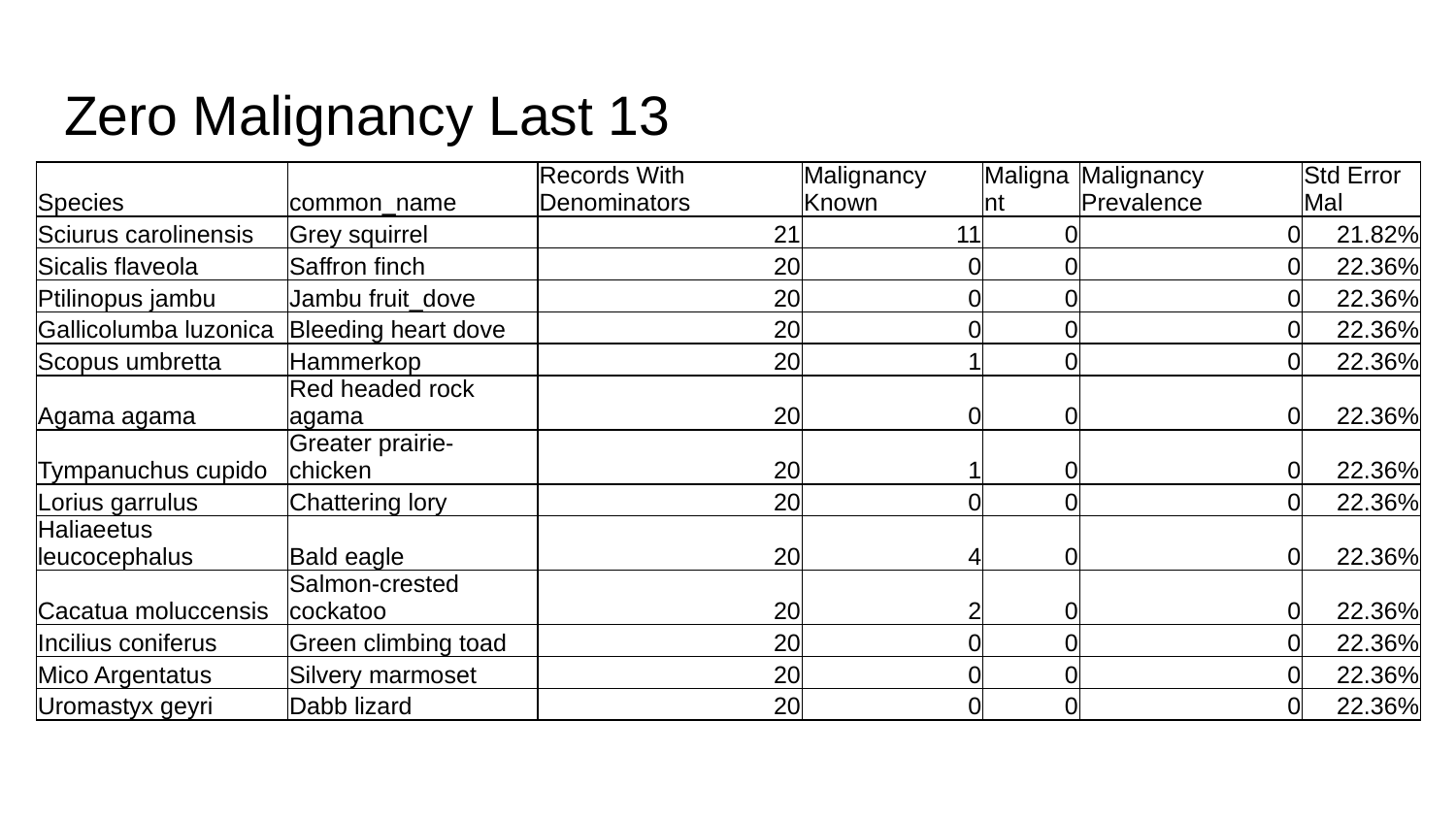

#### Zero Malignancy Last 13
| Species | common\_name | Records With Denominators | Malignancy Known | Malignant | Malignancy Prevalence | Std Error Mal |
| --- | --- | --- | --- | --- | --- | --- |
| Sciurus carolinensis | Grey squirrel | 21 | 11 | 0 | 0 | 21.82% |
| Sicalis flaveola | Saffron finch | 20 | 0 | 0 | 0 | 22.36% |
| Ptilinopus jambu | Jambu fruit\_dove | 20 | 0 | 0 | 0 | 22.36% |
| Gallicolumba luzonica | Bleeding heart dove | 20 | 0 | 0 | 0 | 22.36% |
| Scopus umbretta | Hammerkop | 20 | 1 | 0 | 0 | 22.36% |
| Agama agama | Red headed rock agama | 20 | 0 | 0 | 0 | 22.36% |
| Tympanuchus cupido | Greater prairie-chicken | 20 | 1 | 0 | 0 | 22.36% |
| Lorius garrulus | Chattering lory | 20 | 0 | 0 | 0 | 22.36% |
| Haliaeetus leucocephalus | Bald eagle | 20 | 4 | 0 | 0 | 22.36% |
| Cacatua moluccensis | Salmon-crested cockatoo | 20 | 2 | 0 | 0 | 22.36% |
| Incilius coniferus | Green climbing toad | 20 | 0 | 0 | 0 | 22.36% |
| Mico Argentatus | Silvery marmoset | 20 | 0 | 0 | 0 | 22.36% |
| Uromastyx geyri | Dabb lizard | 20 | 0 | 0 | 0 | 22.36% |
