## Supplementary figures and images for "Cancer Prevalence Across Vertebrates"

### cladesmush2.0.pdf

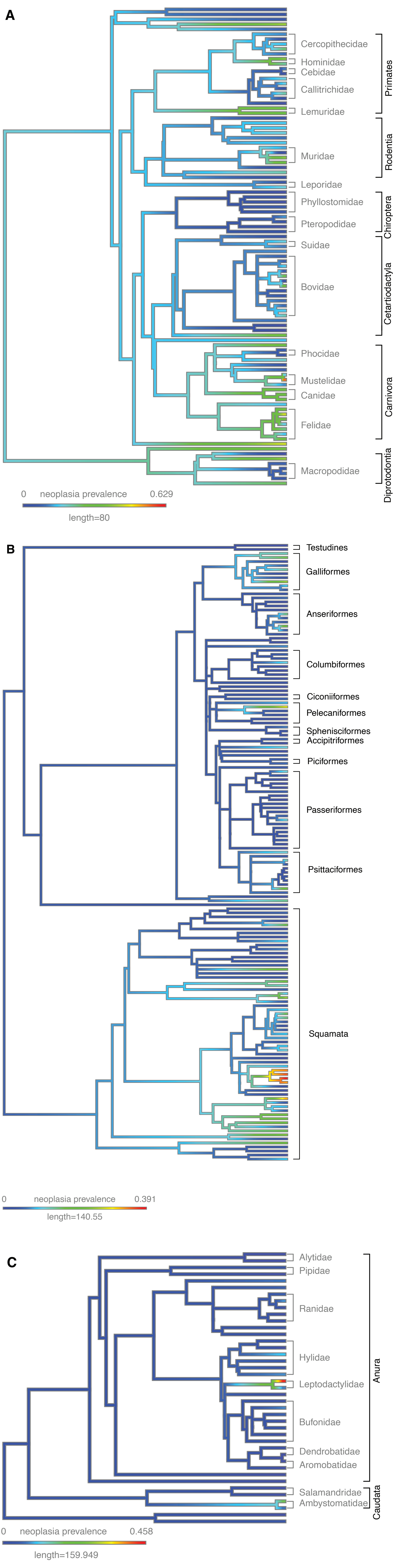

### correctsmushfix.pdf

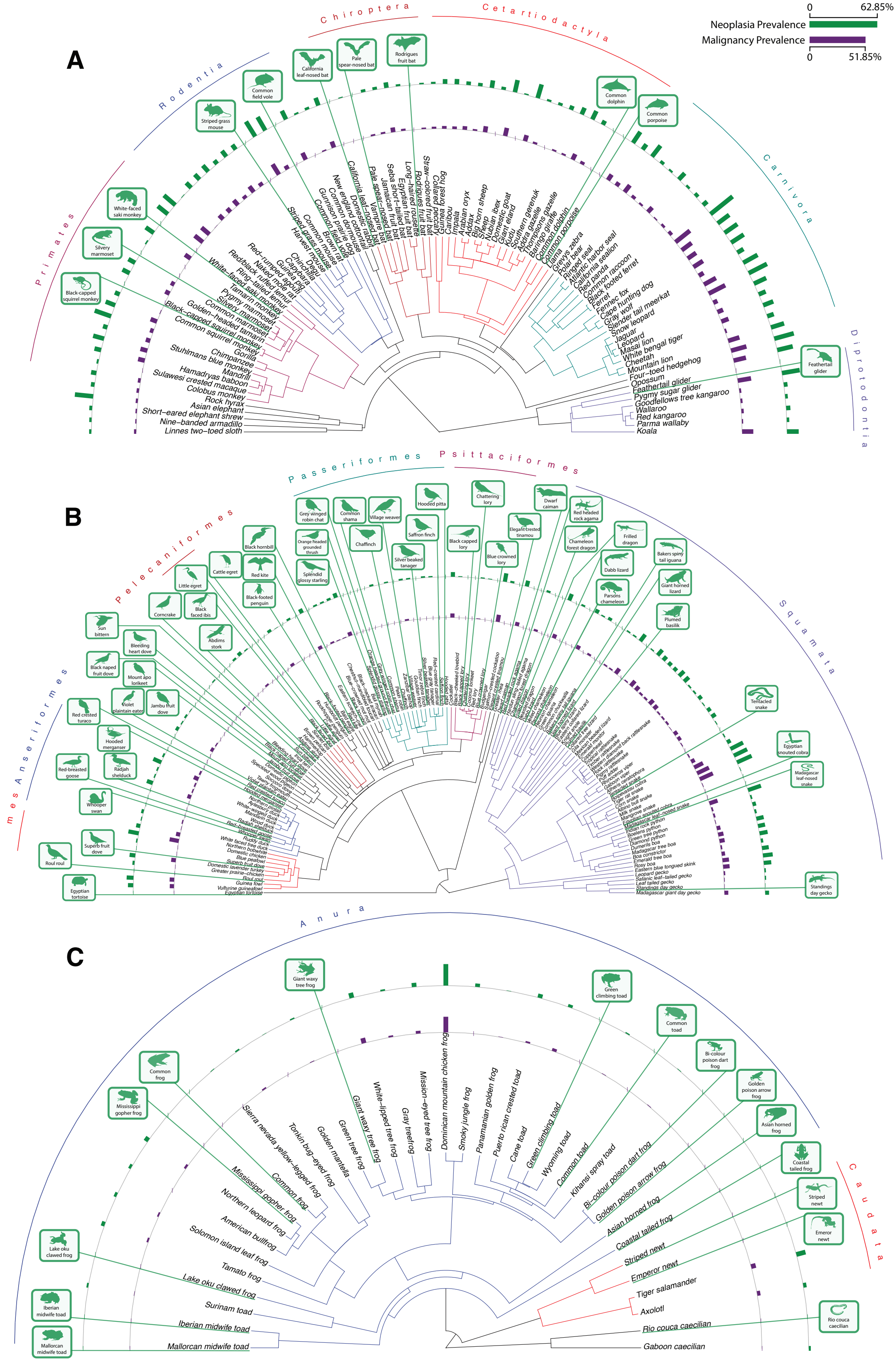

### lifehistorypgls.pdf

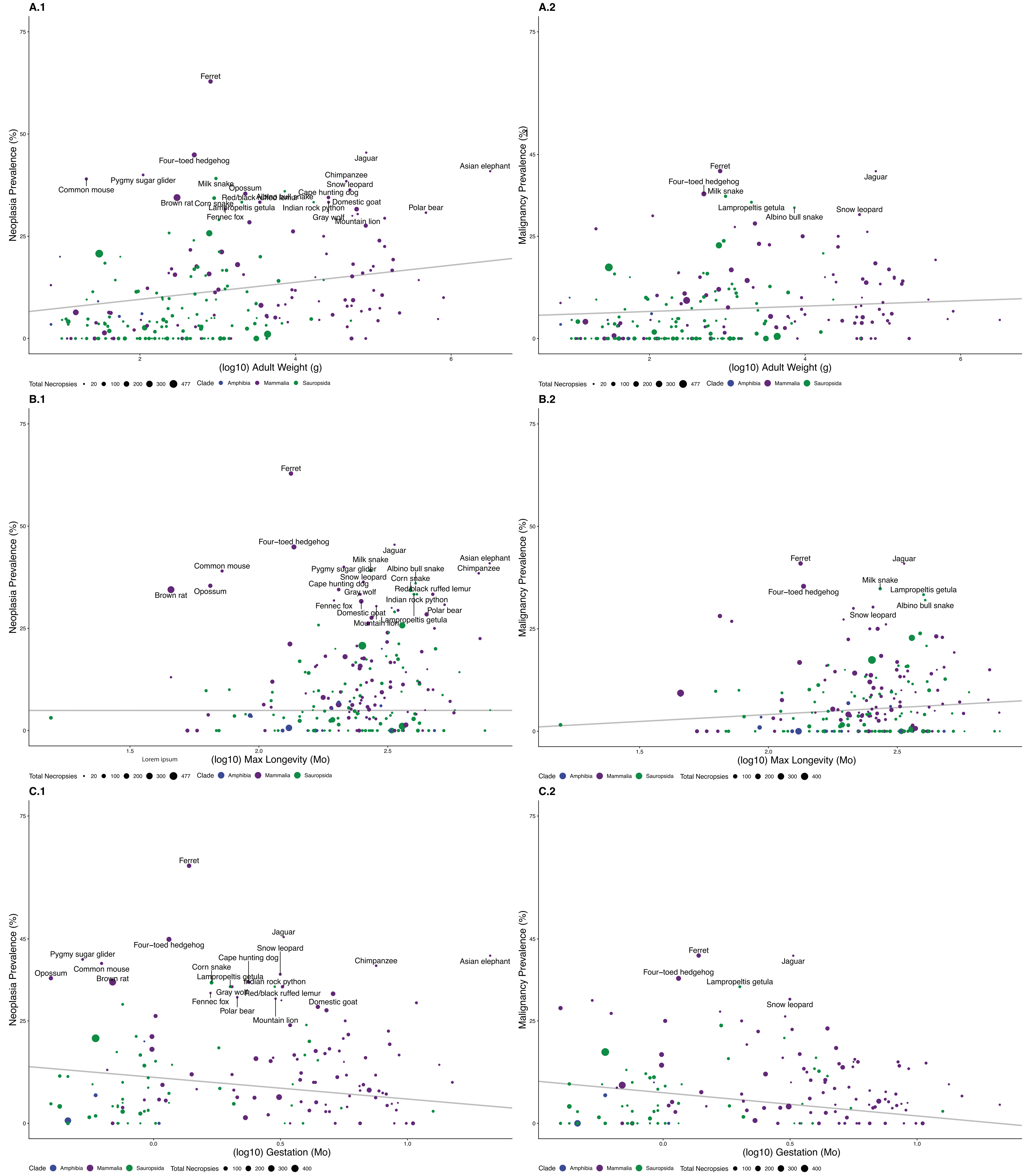

### Maldensity.png

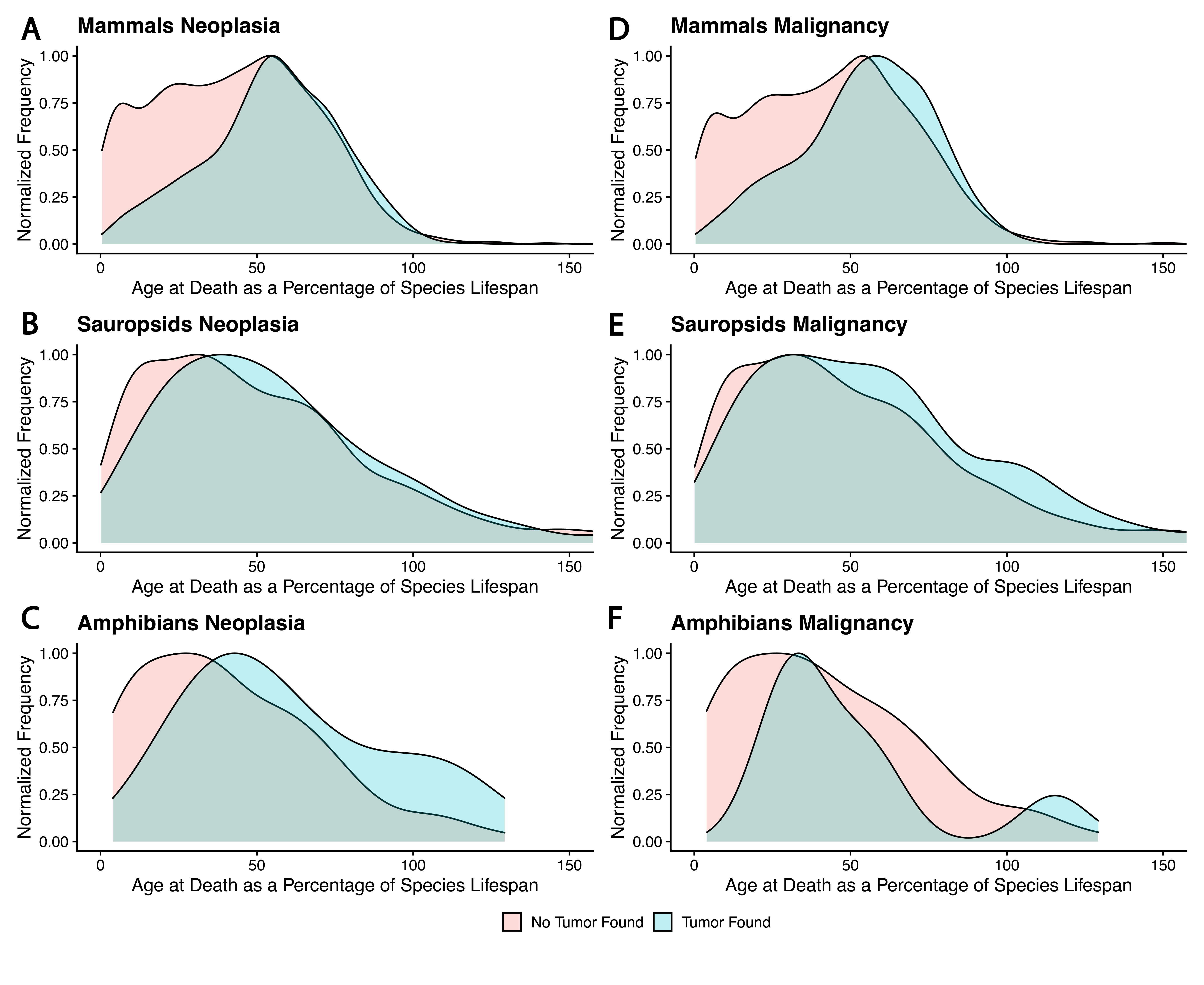

### radymutation.png

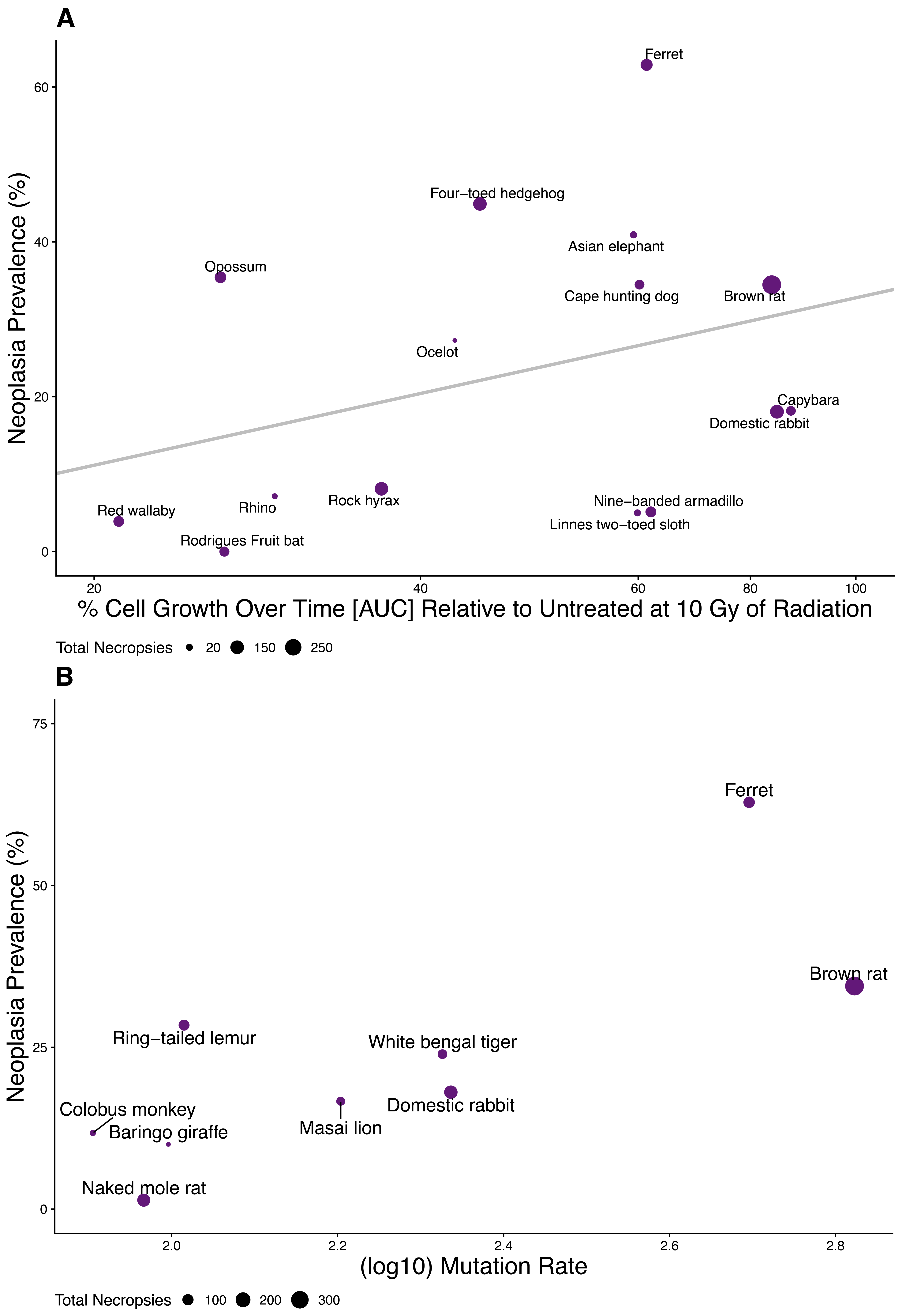

### S1litneo.pdf

1 **Neoplasia Prevalence vs. Litter Size**

p − value : 0.23 $R^2$ : 0.292 $\Lambda$ : 0.4

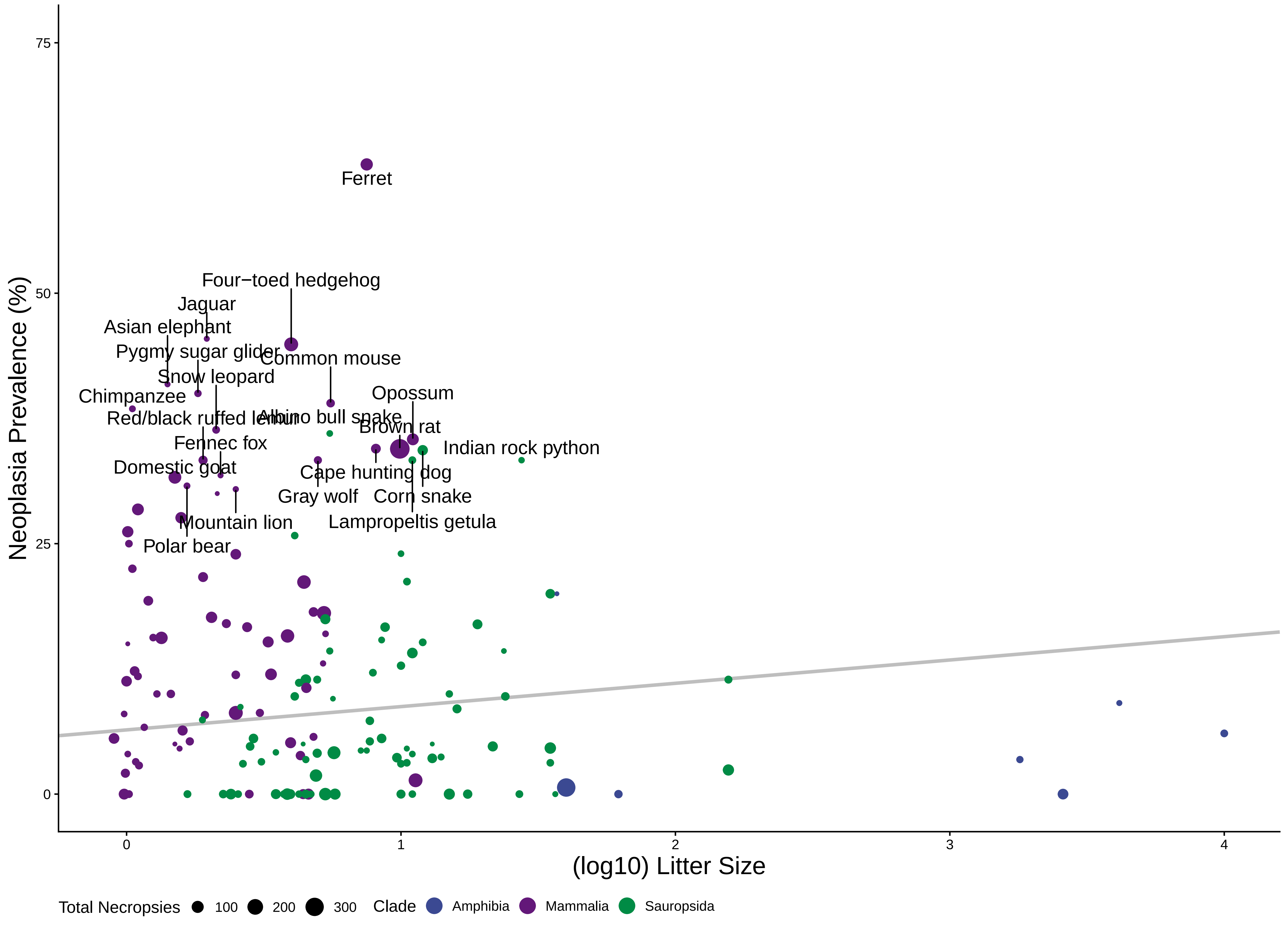

### S2litmal.pdf

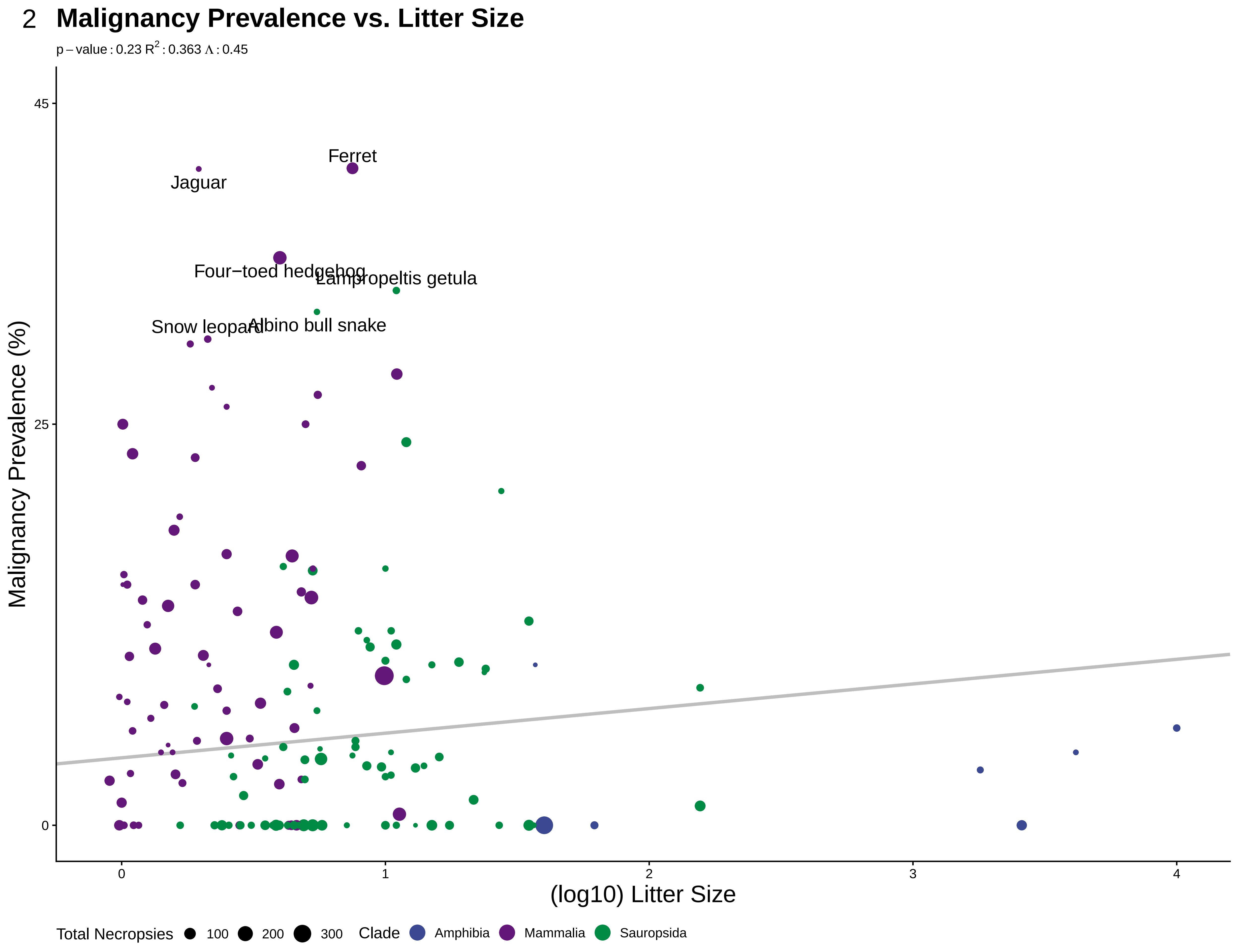

### S3bmrneo.pdf

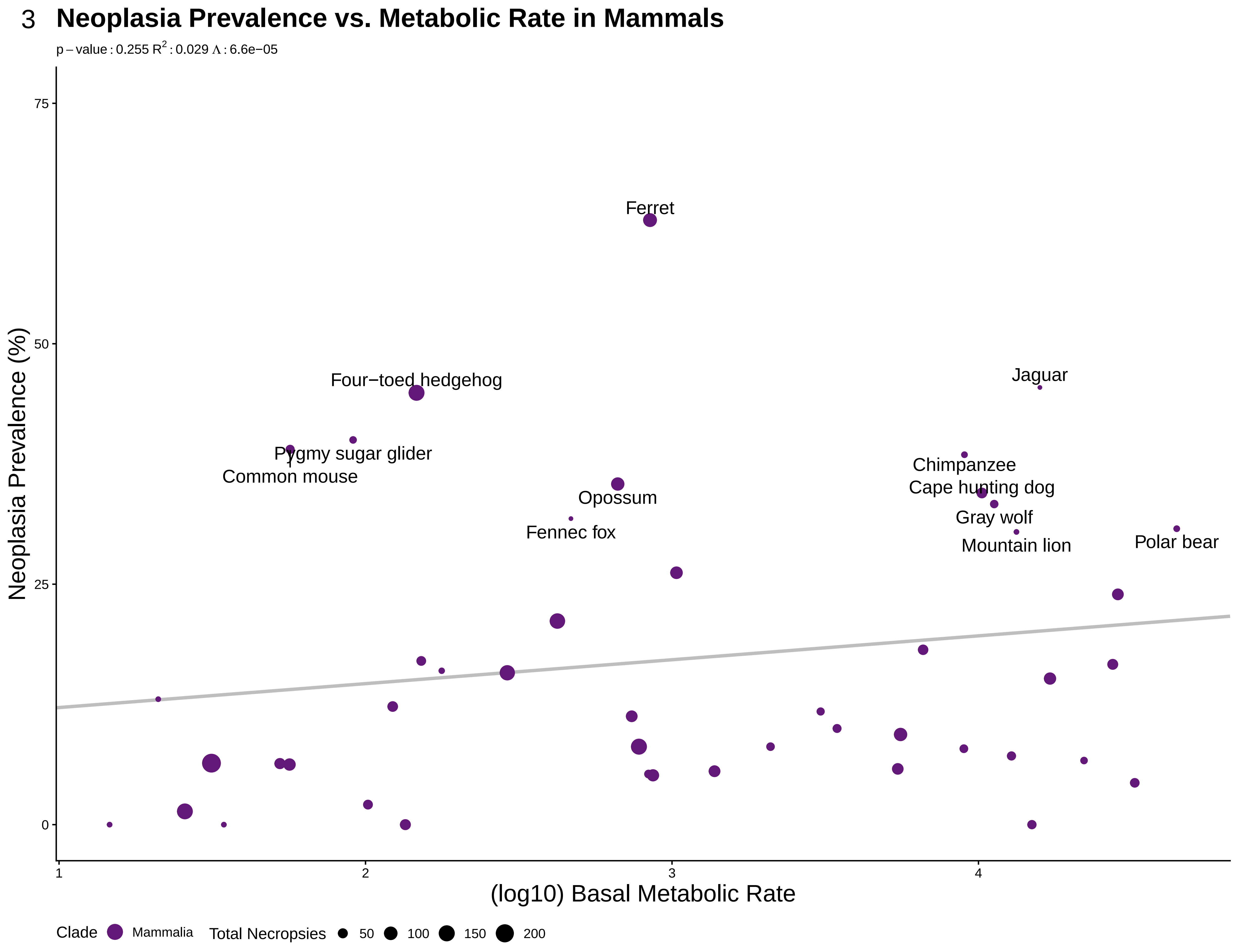

### S4bmrmal.pdf

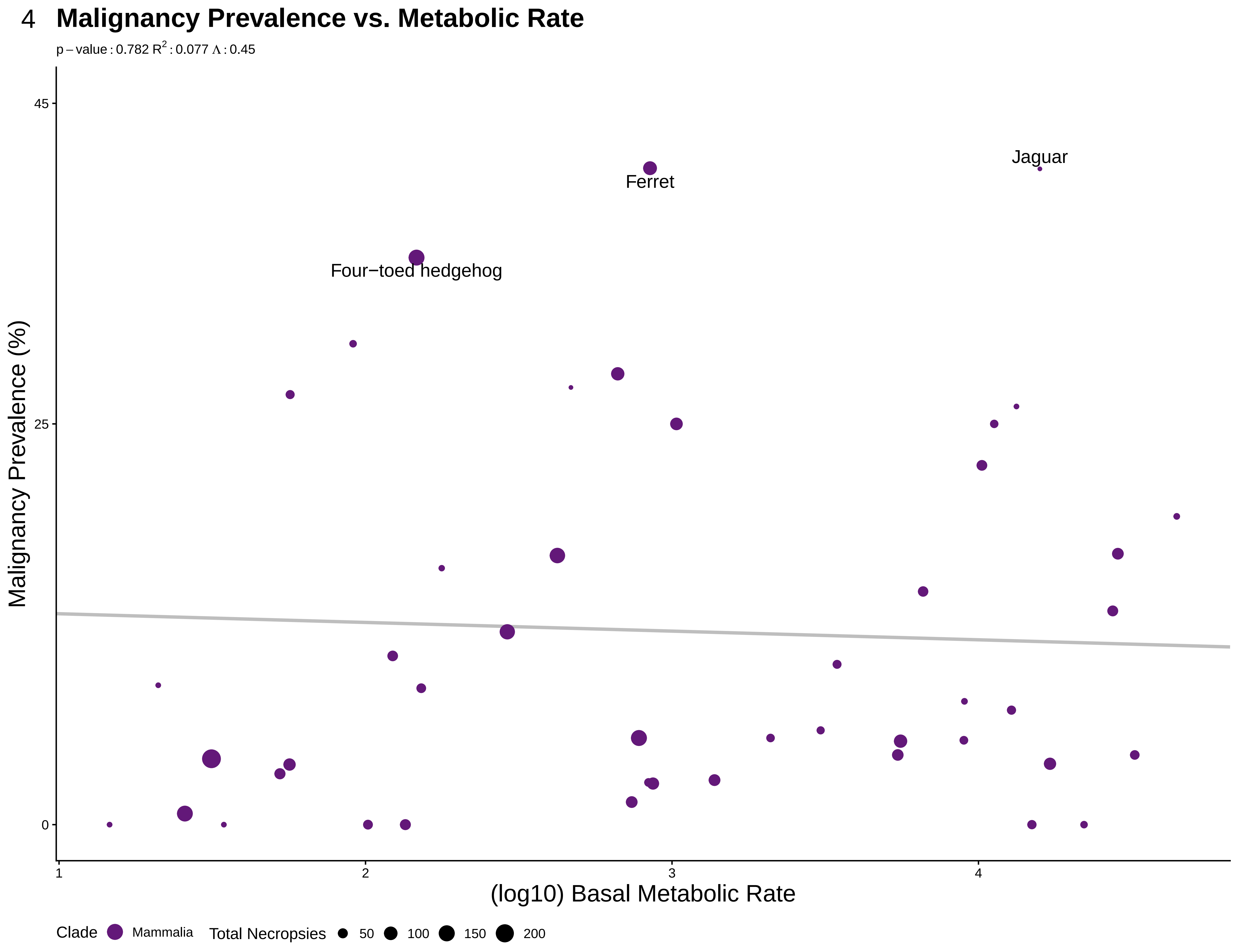

### S5wgtlongneo.pdf

5 **Neoplasia Prevalence vs. Max Longevity*Weight**

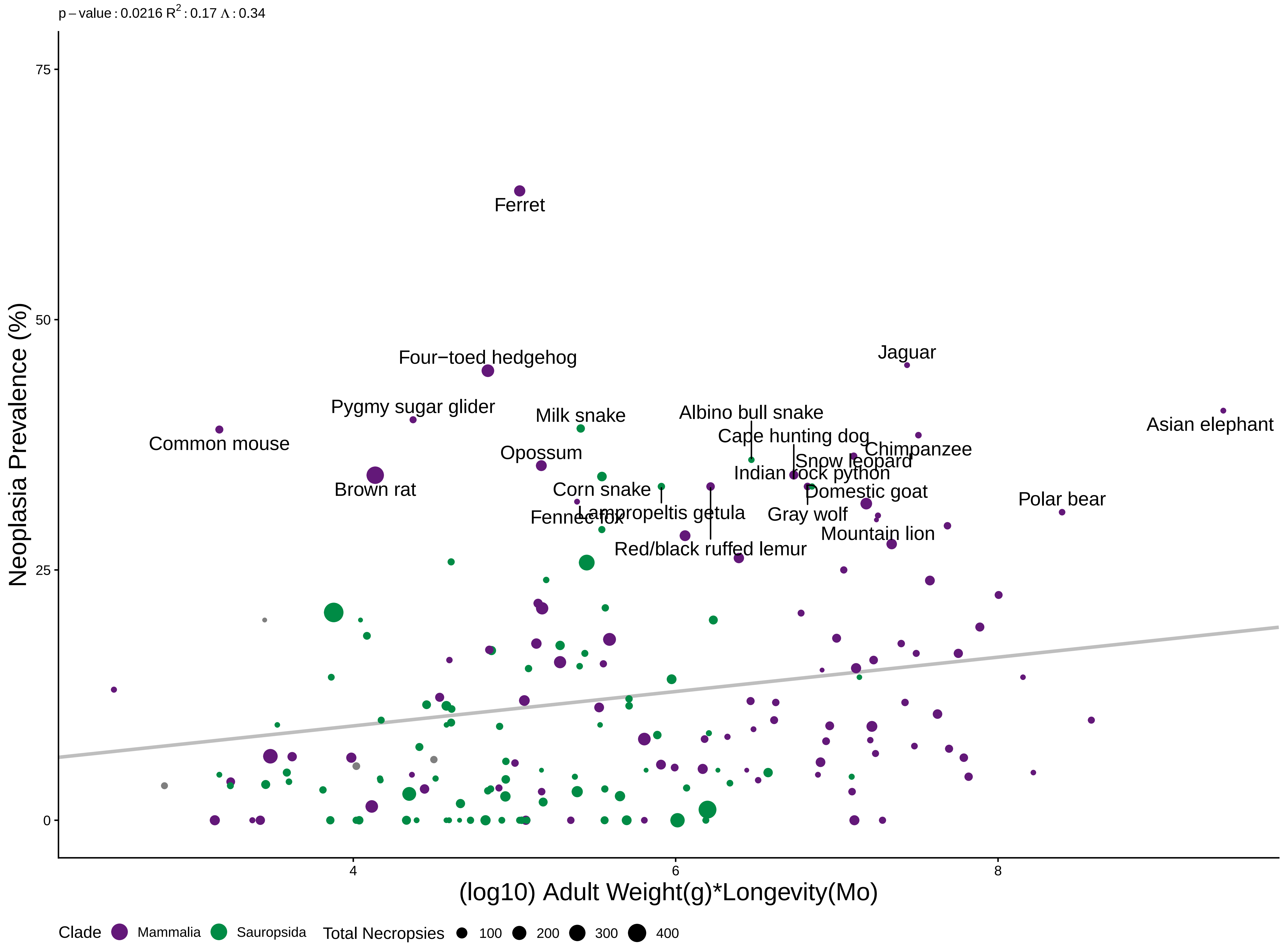

### S6wgtlongmal.pdf

#### 6 Malignancy Prevalence vs. Max Longevity*Weight

p – value : 0.376 $R^2$ : 0.21 $\Lambda$ : 0.55

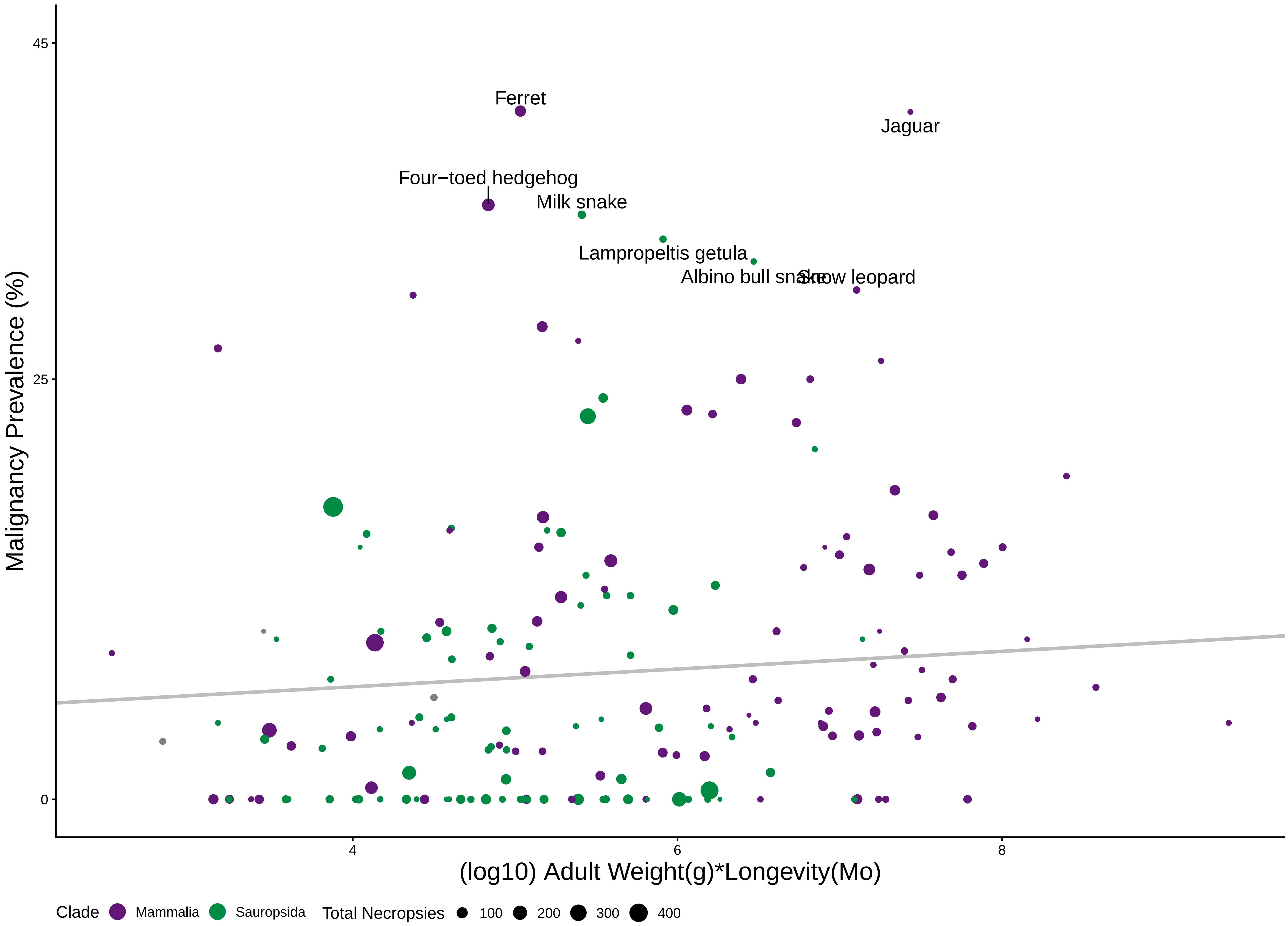

### S7lityearneo.pdf

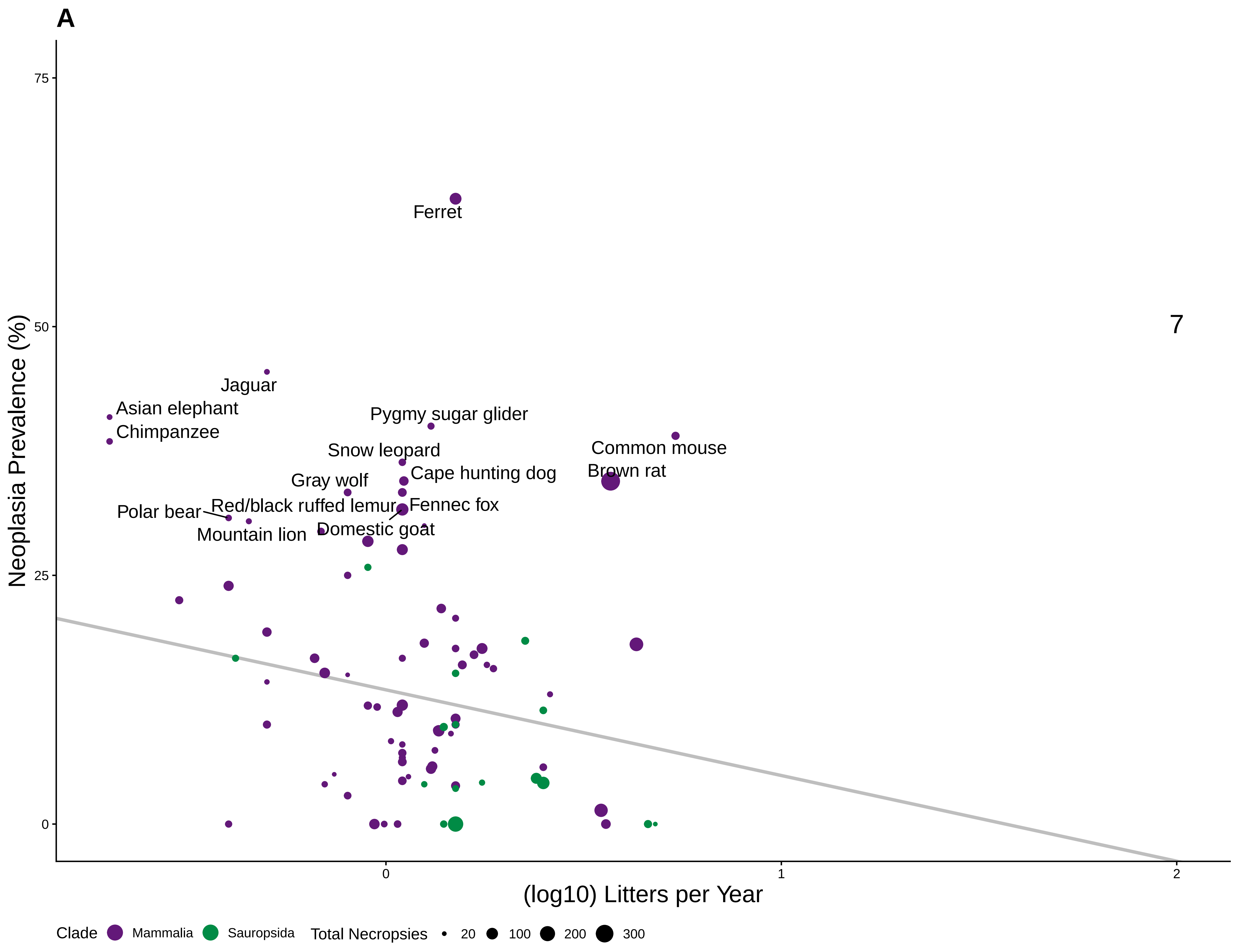

### S8lityearmal.pdf

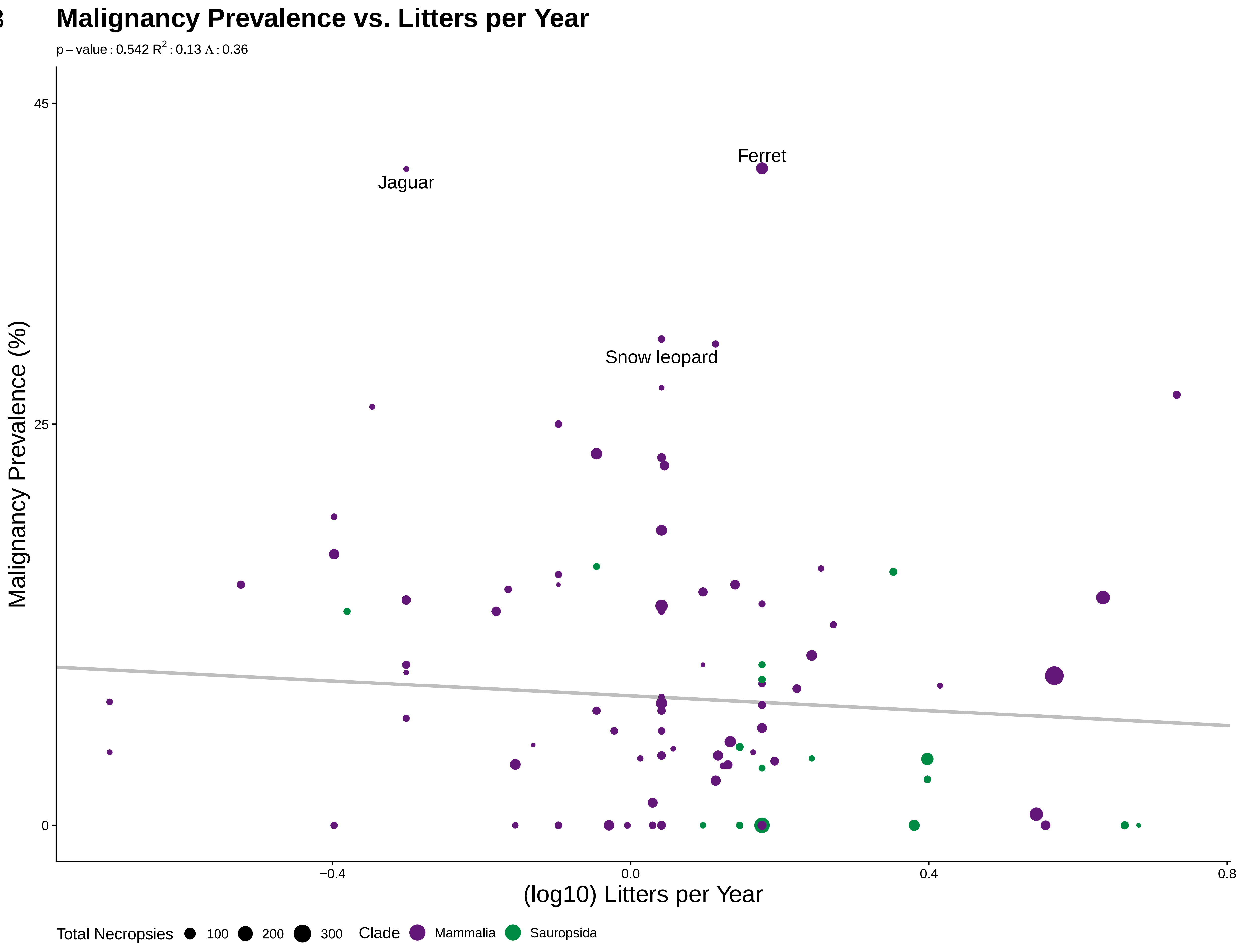

### S9femmatneo.pdf

9 **Neoplasia Prevalence vs. Female Maturity**

p – value : 0.658 $R^2$ : 0.16 $\Lambda$ : 0.43

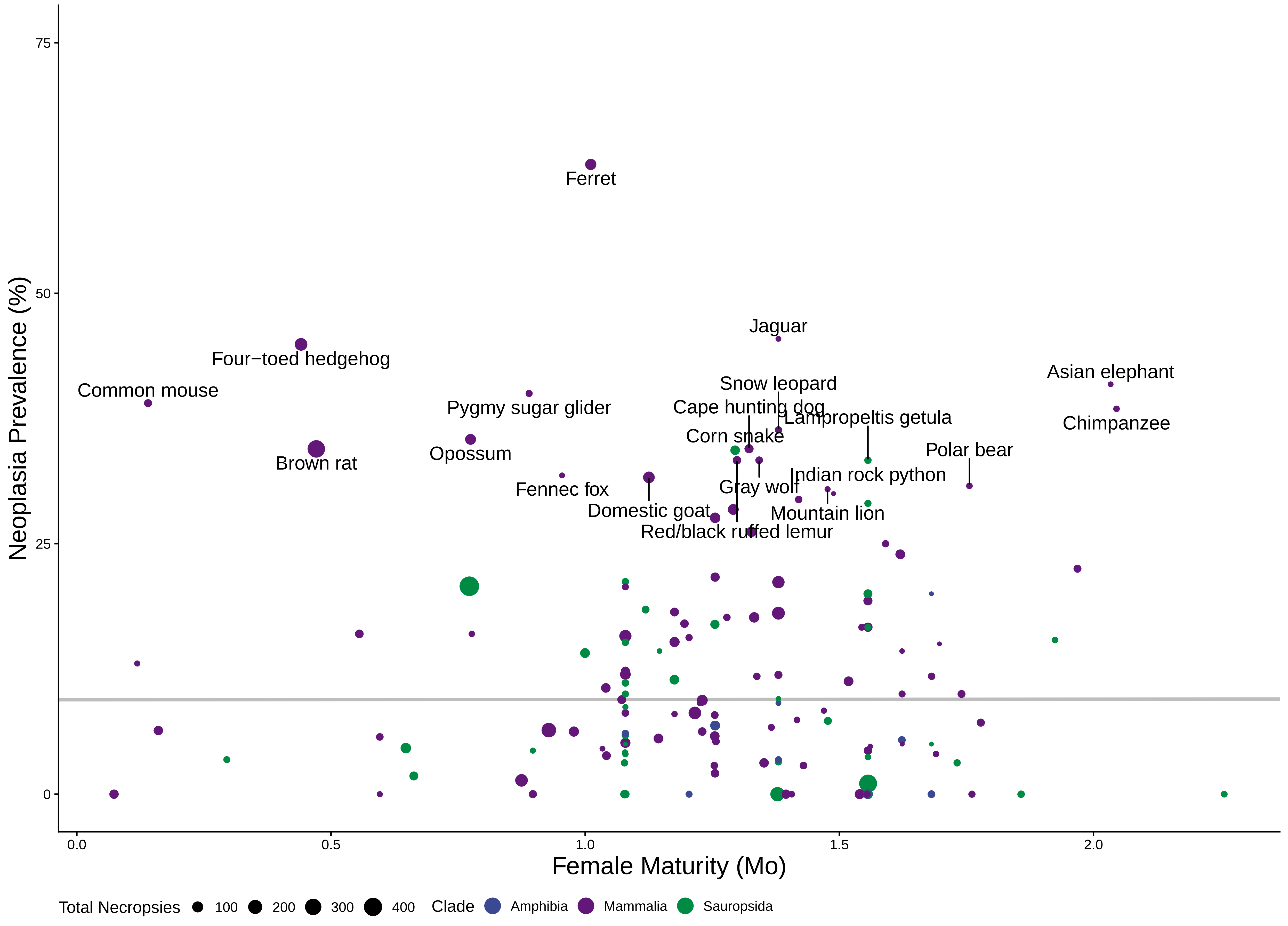

### S10femmatmal.pdf

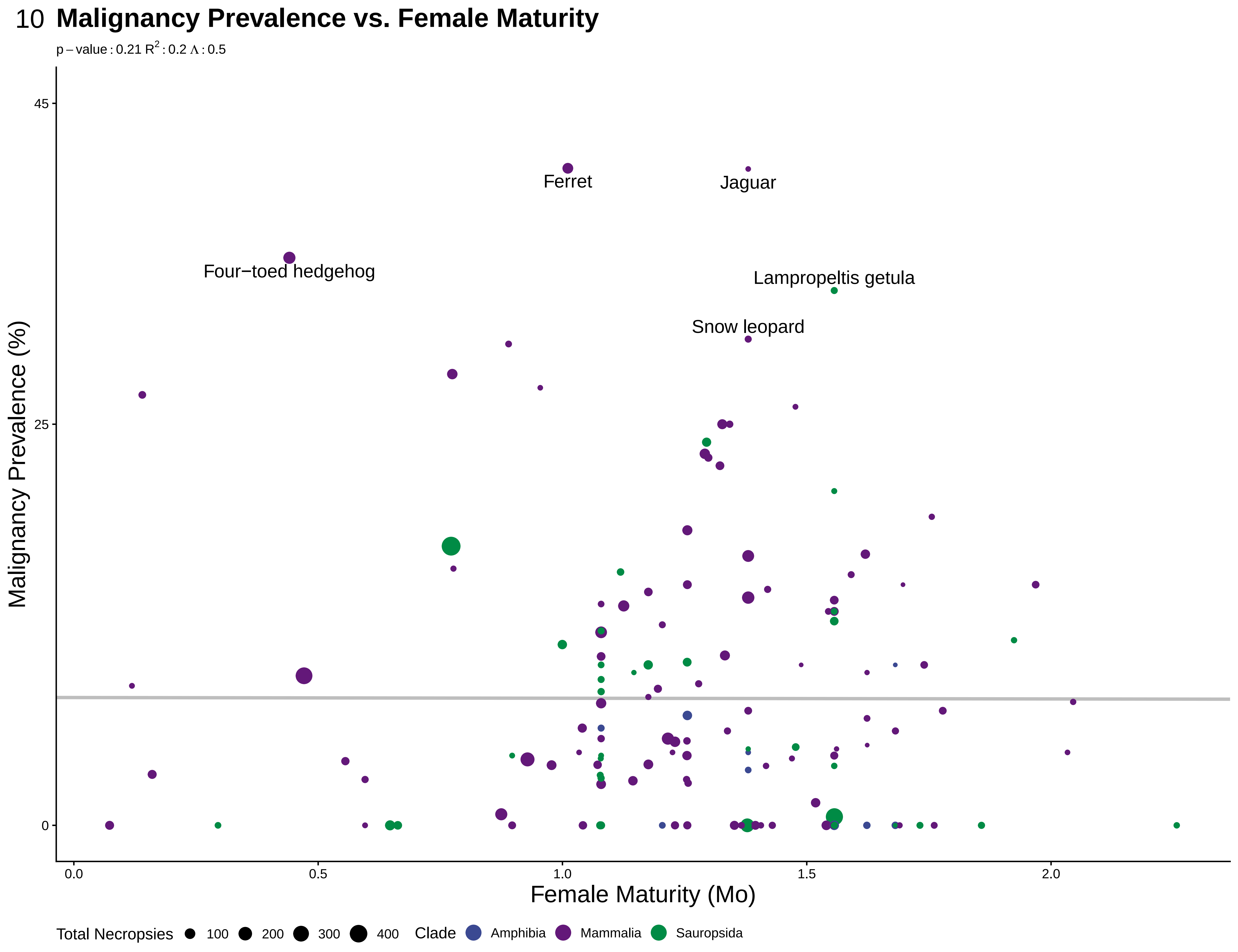

### S11malematneo.pdf

11 **Neoplasia Prevalence vs. Male Maturity**

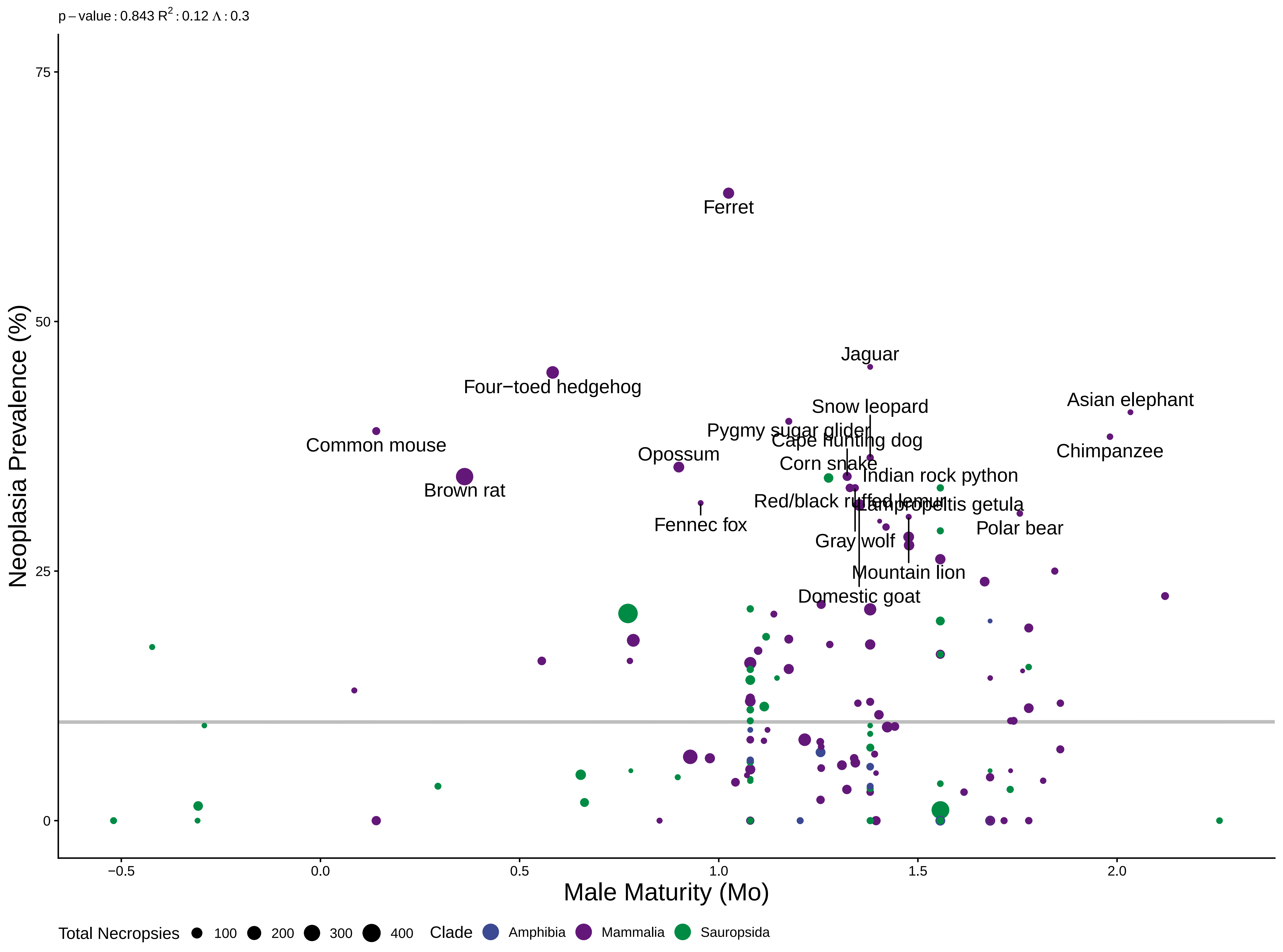

### S12malematmal.pdf

12 **Malignancy Prevalence vs. Male Maturity**

p – value : 0.291 $R^2$ : 0.16 Λ : 0.41

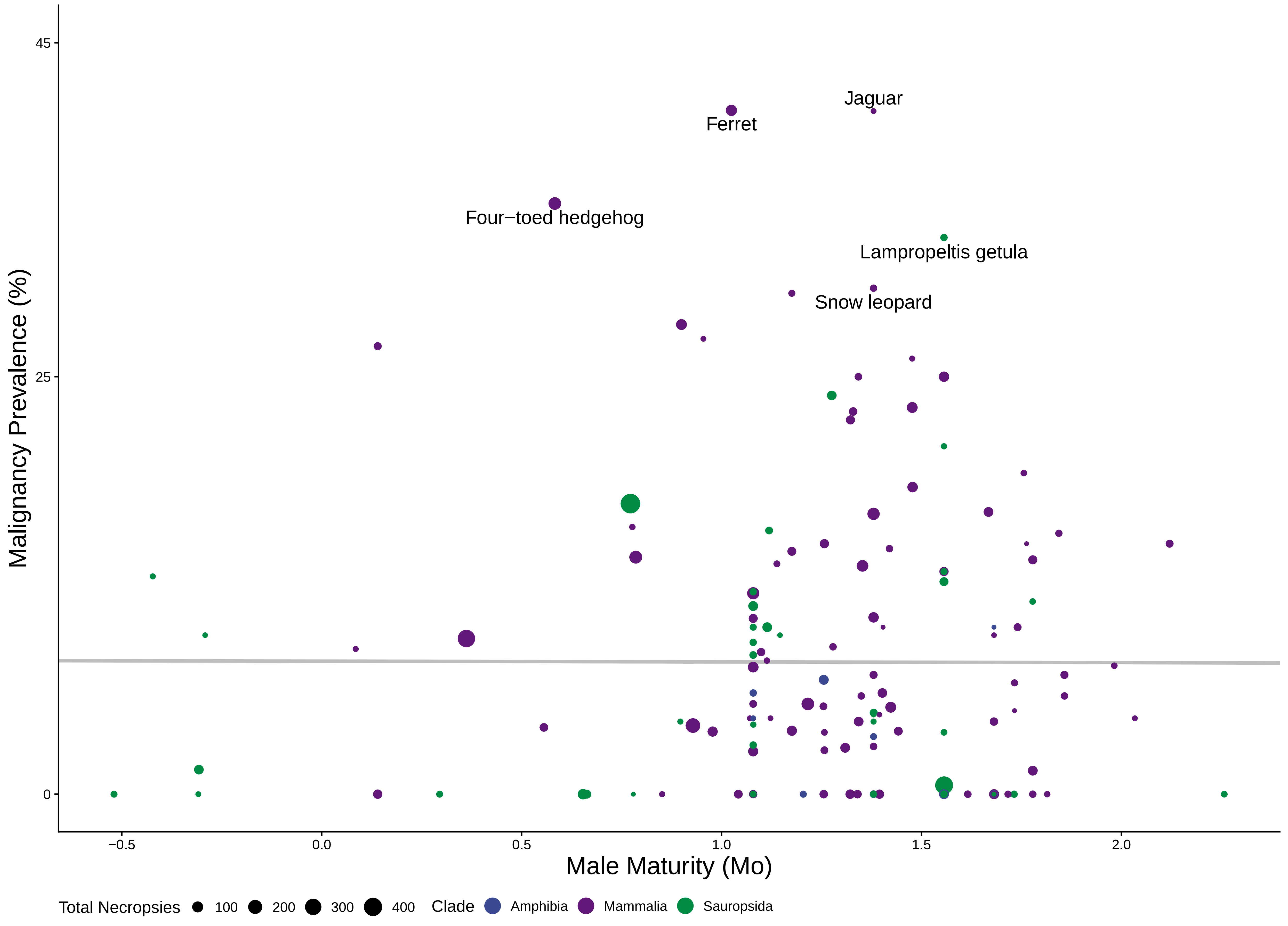

### S13weanneo.pdf

#### Neoplasia Prevalence vs. Weaning Weight

p – value : 0.138 $R^2$ : 0.084 Λ : 0.67

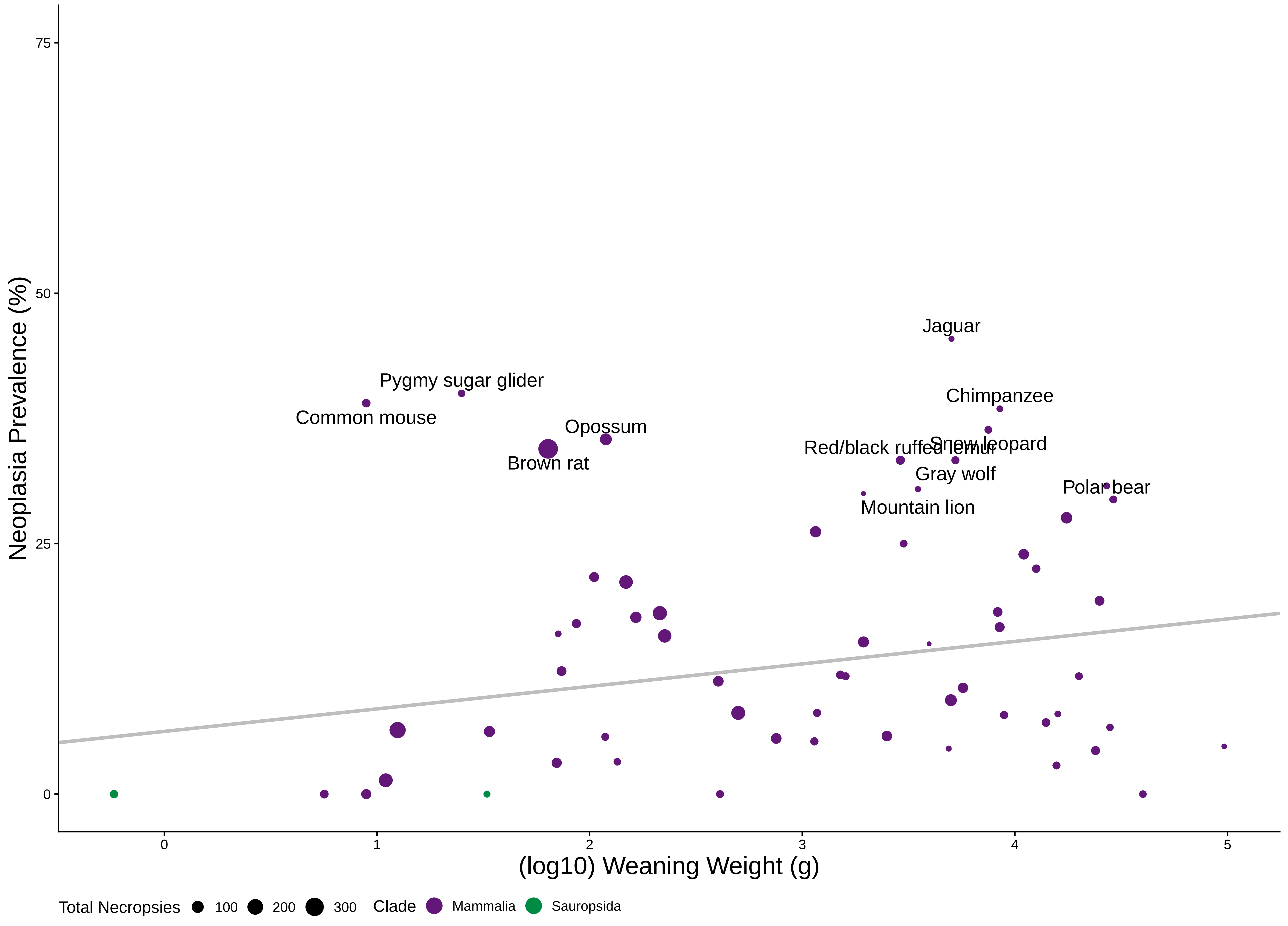

### S14weanmal.pdf

#### 14 Malignancy Prevalence vs. Weaning Weight

p – value : 0.178 $R^2$ : 0.018 $\Lambda$ : 6.6e−05

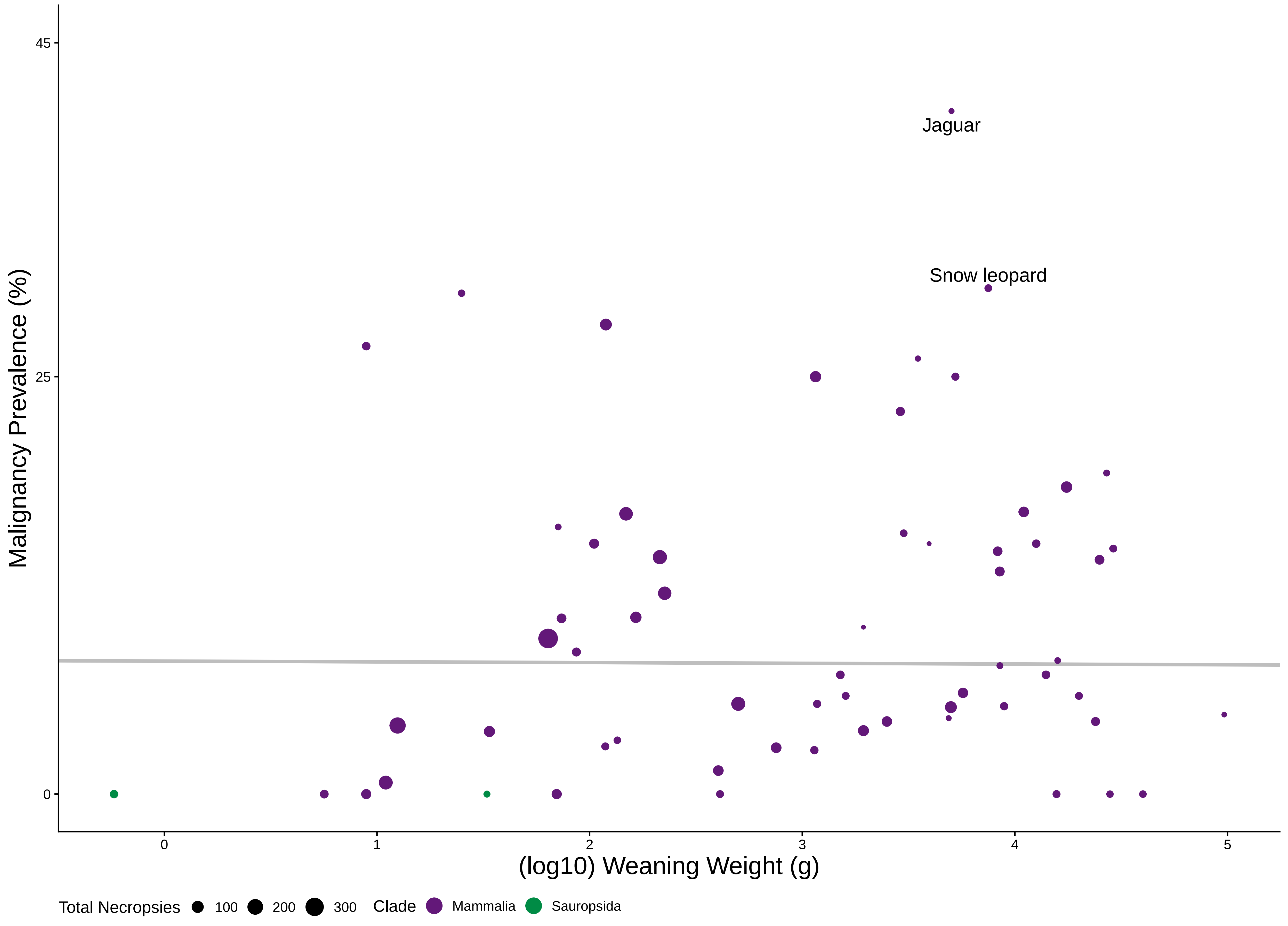

### S15growneo.pdf

#### 15 Neoplasia Prevalence vs. Growth Rate

p − value : 0.258 $R^2$ : 0.026 Λ : 6.6e−05

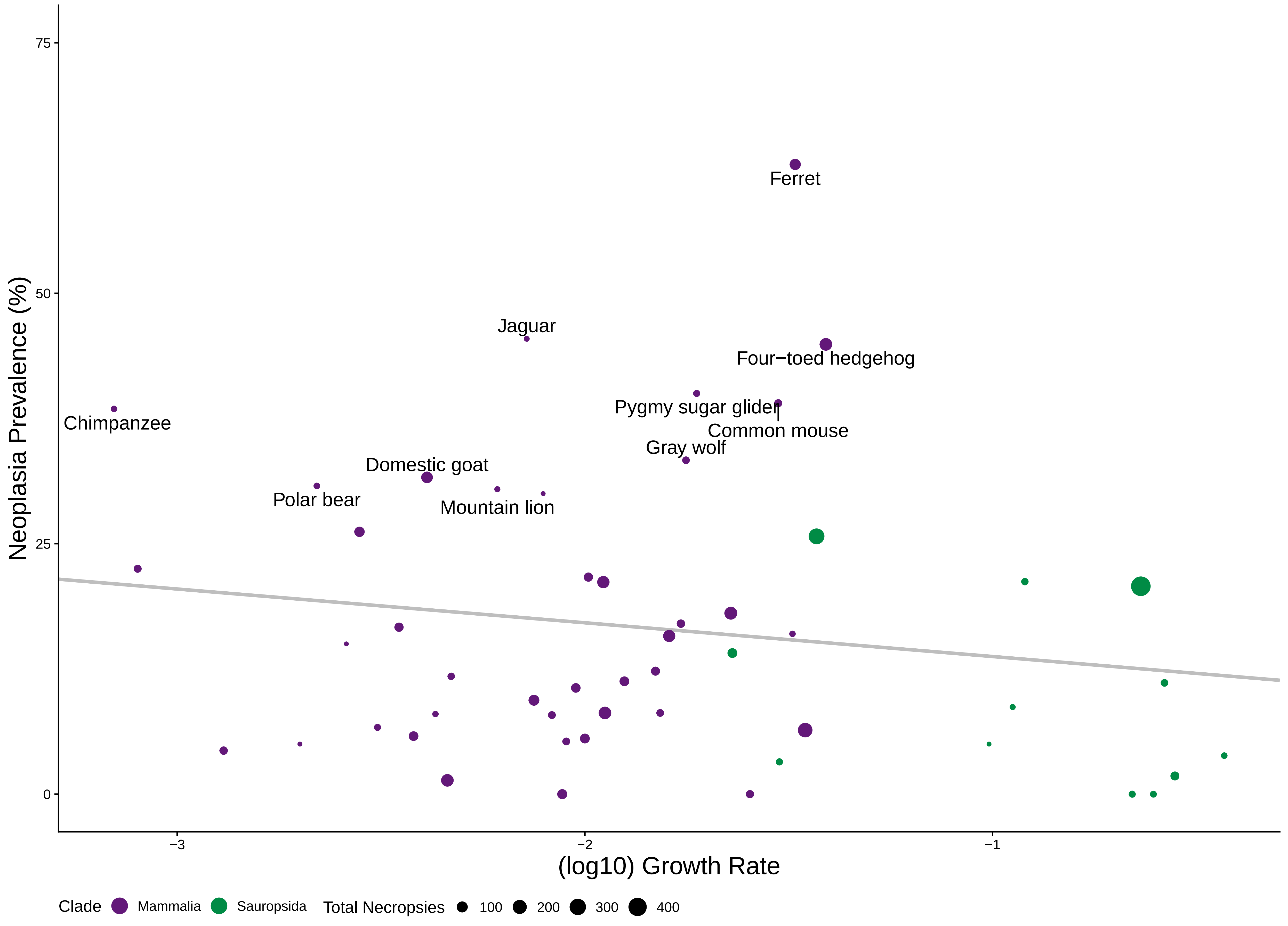

### S16growmal.pdf

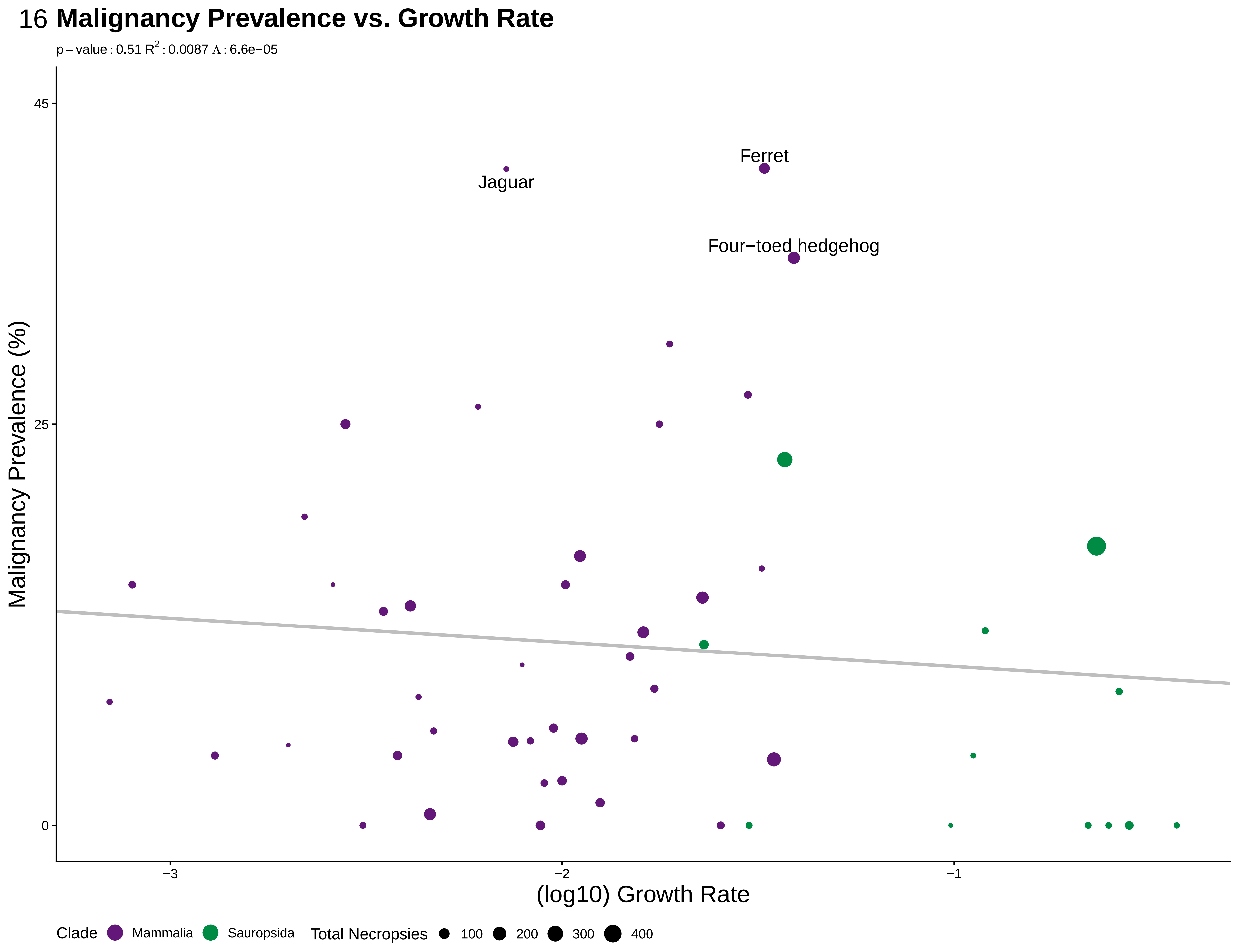

### S17wgtneo.pdf

17 **Neoplasia vs. Adult Weight**

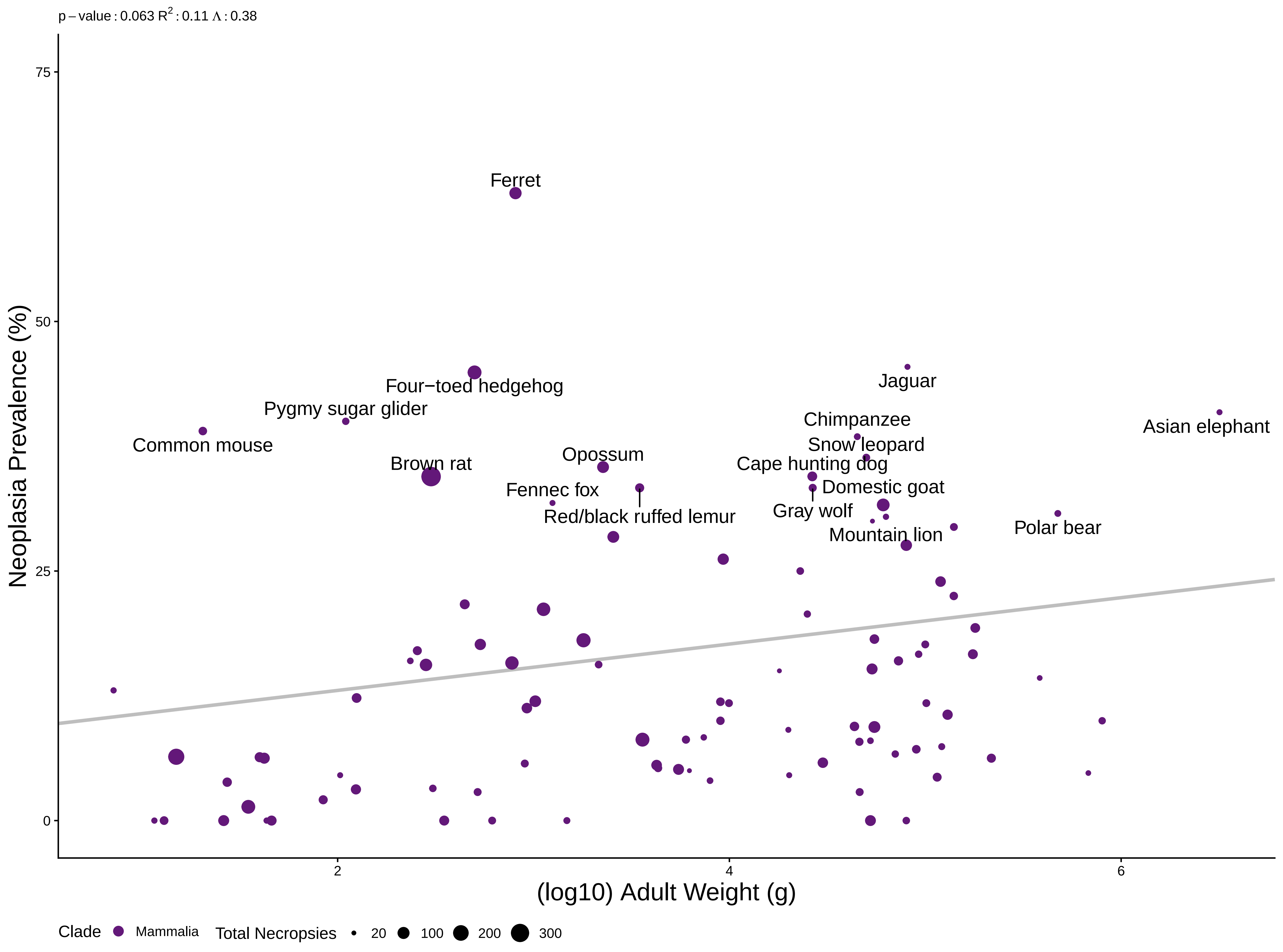

### S18wgtmal.pdf

#### Malignancy Prevalence vs. Adult Weight

p – value : 0.458 $R^2$ : 0.12 Λ : 0.43

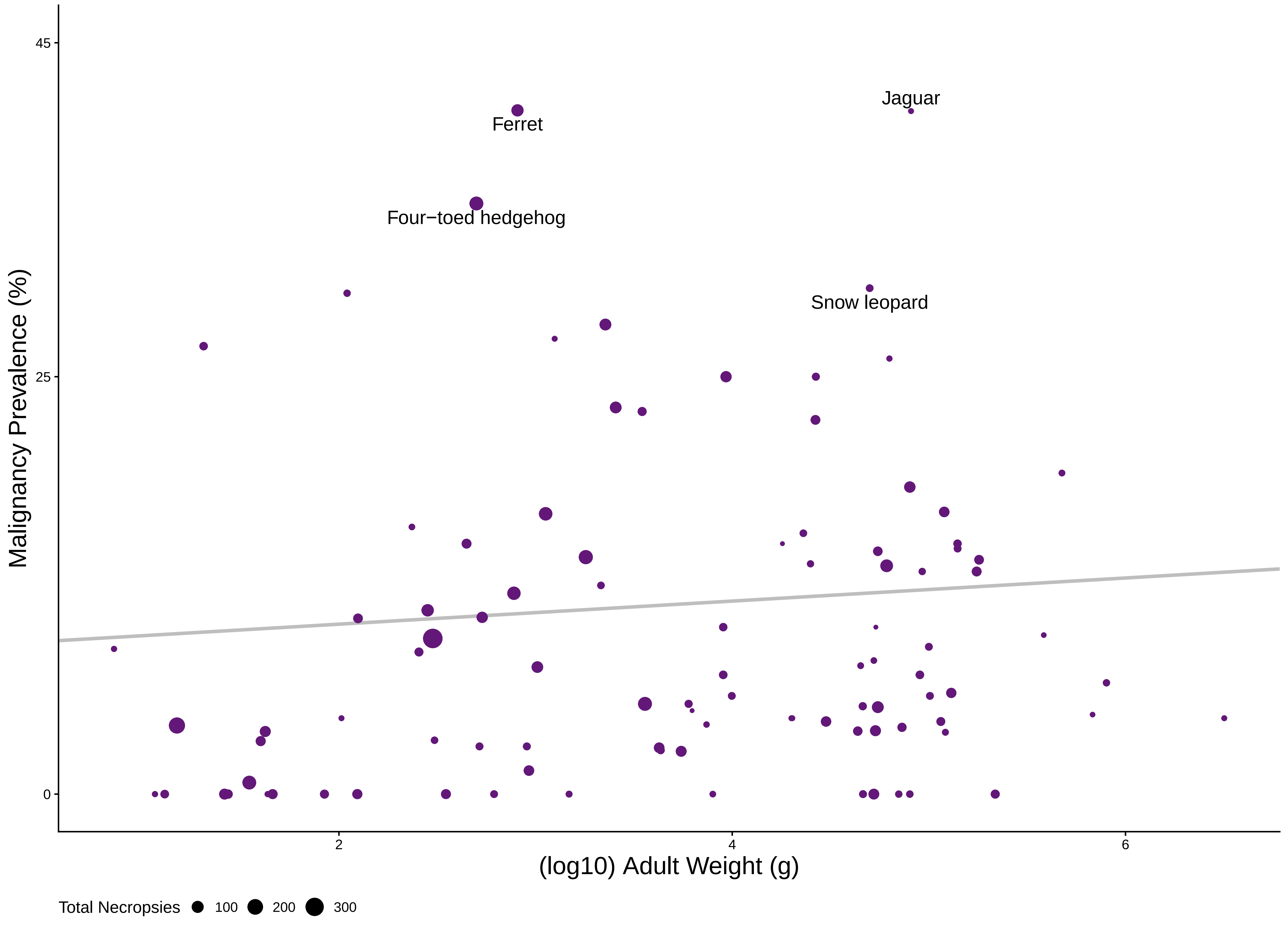

### S19gestneomam.pdf

19 **Neoplasia Prevalence vs. Gestation in Mammals**

p – value : 0.0369 $R^2$ : 0.14 Λ : 0.43

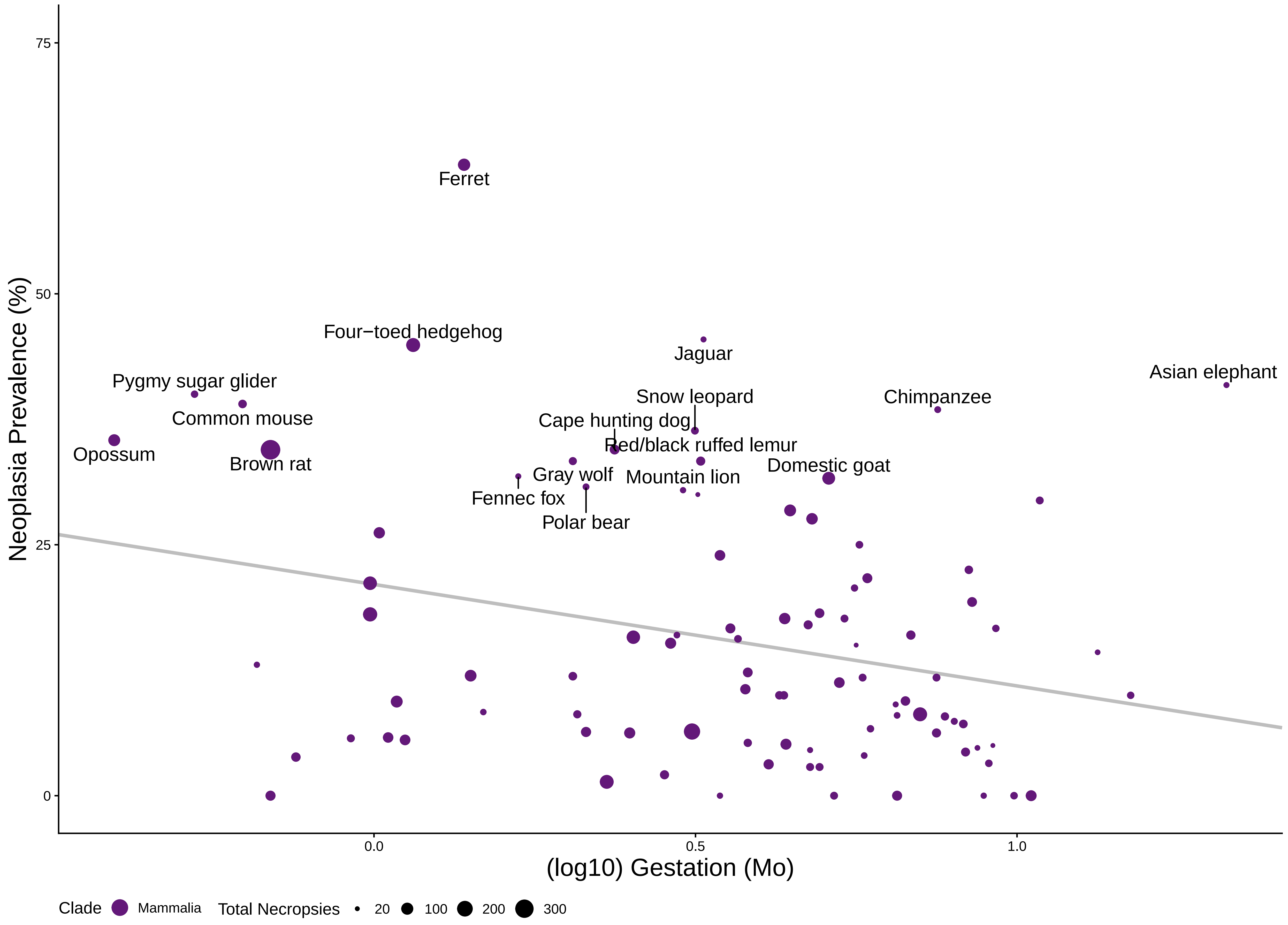

### S20gestmalmam.pdf

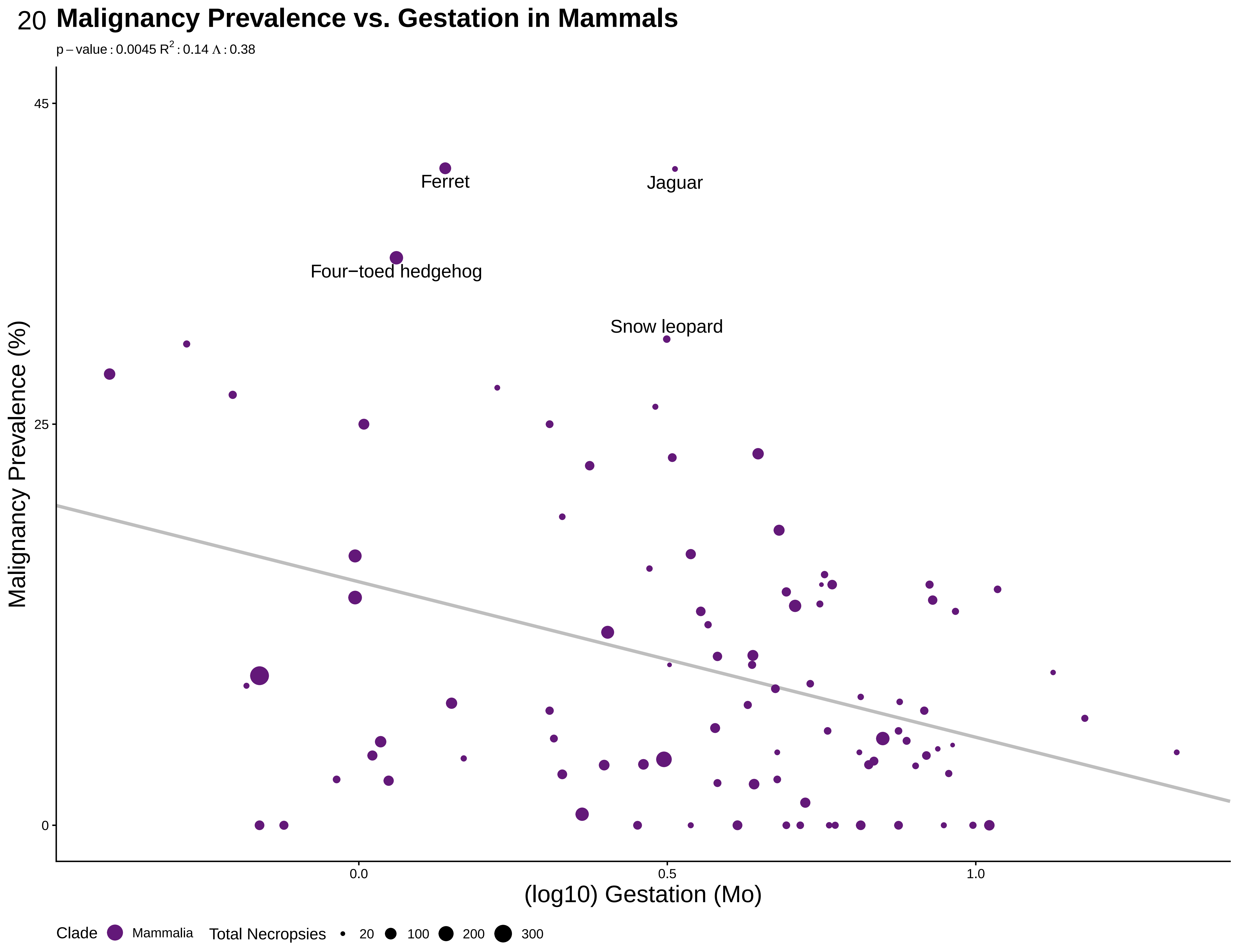

### S21litneomam.pdf

21 **Neoplasia Prevalence vs. Litter Size in Mammals**

p – value : 0.0205 $R^2$ : 0.073 Λ : 8e−05

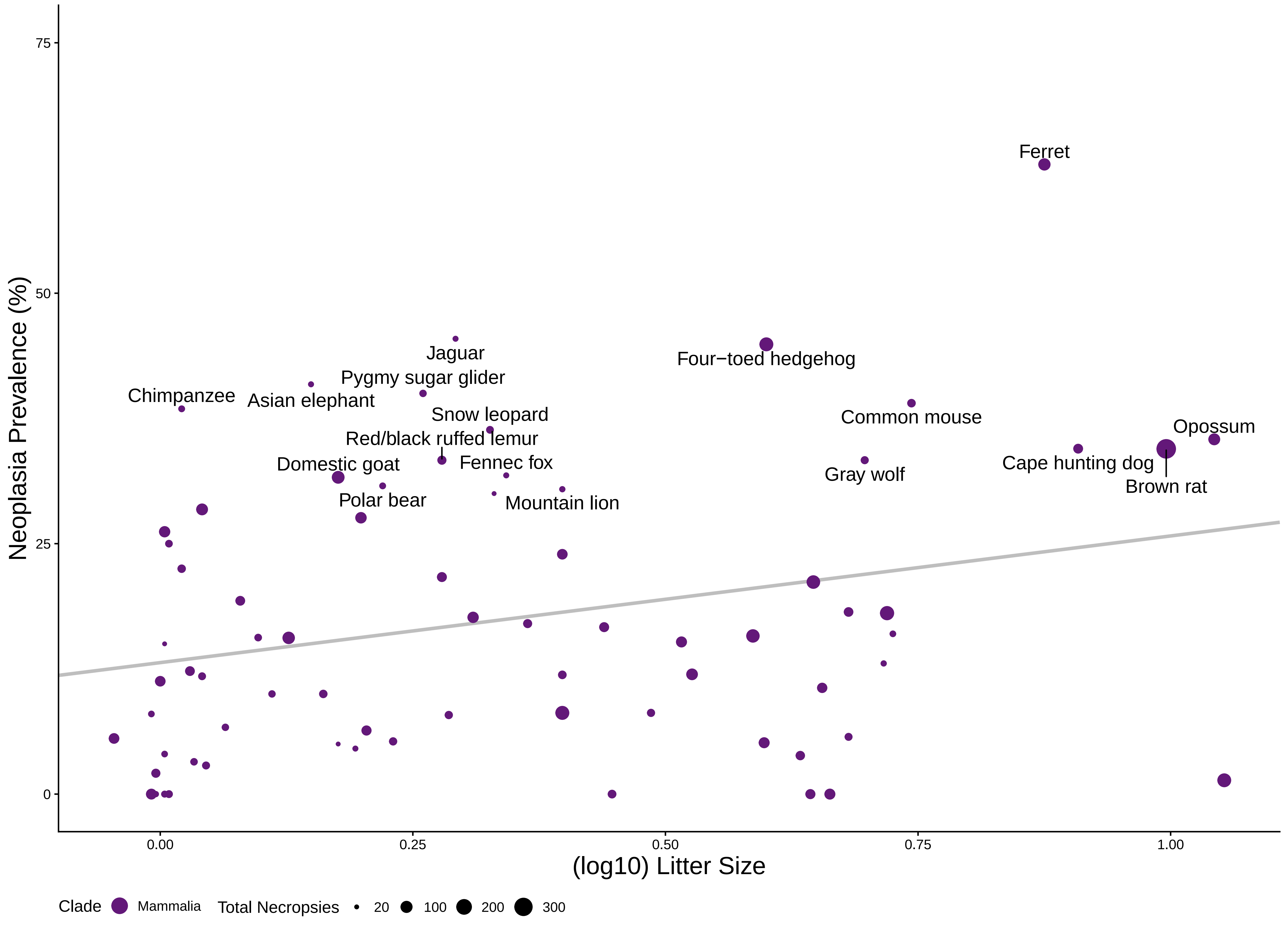

### S22litmalmam.pdf

22 **Malignancy Prevalence vs. Litter Size in Mammals**

p – value : 0.0306 $R^2$ : 0.11 Λ : 0.2

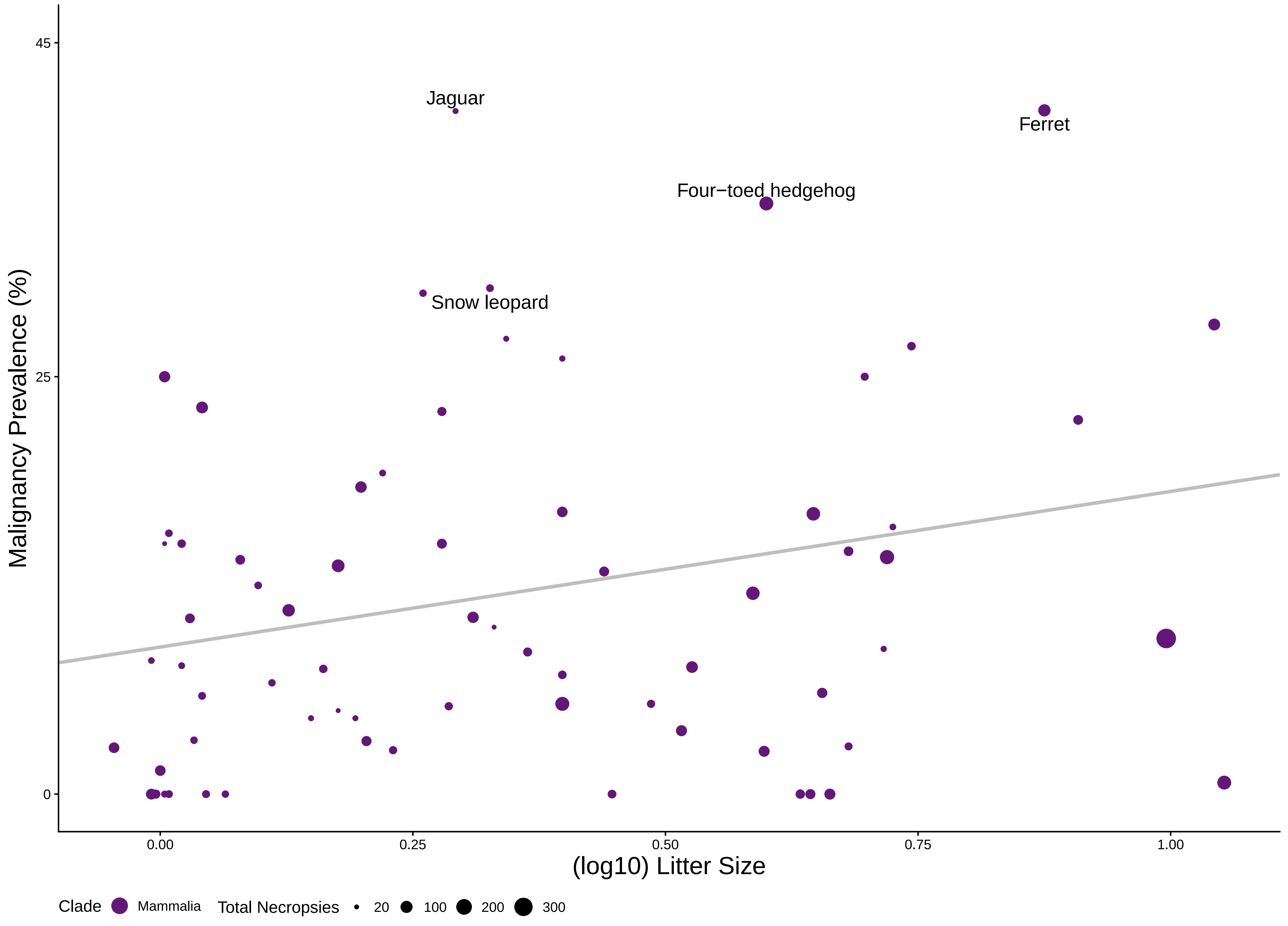

### S23longneomam.pdf

23 **Neoplasia Prevalence vs. Max Longevity in Mammals**

p – value : 0.323 $R^2$ : 0.072 Λ : 0.38

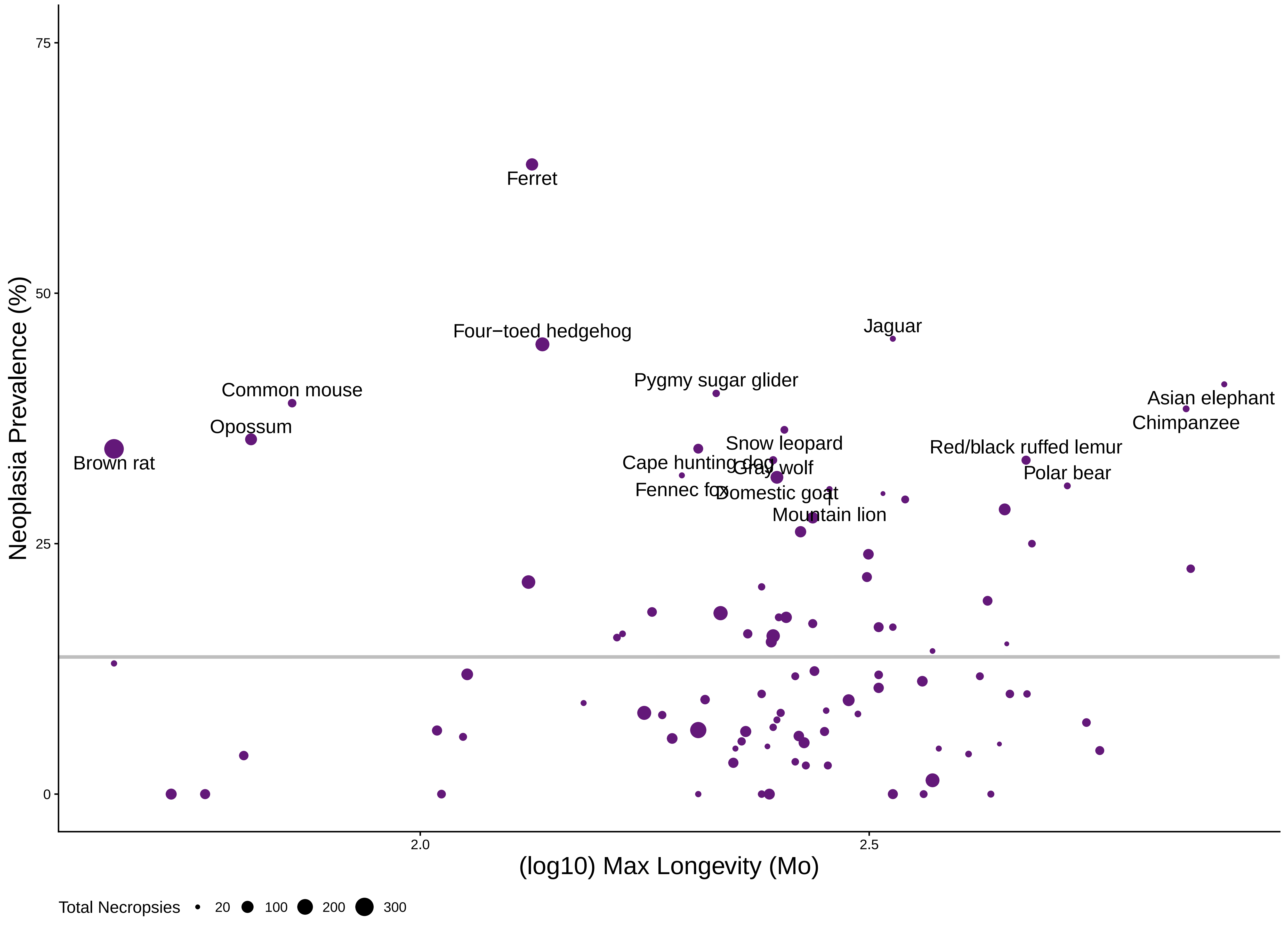

### S24longmalmam.pdf

#### Malignancy Prevalence vs. Max Longevity in Mammals

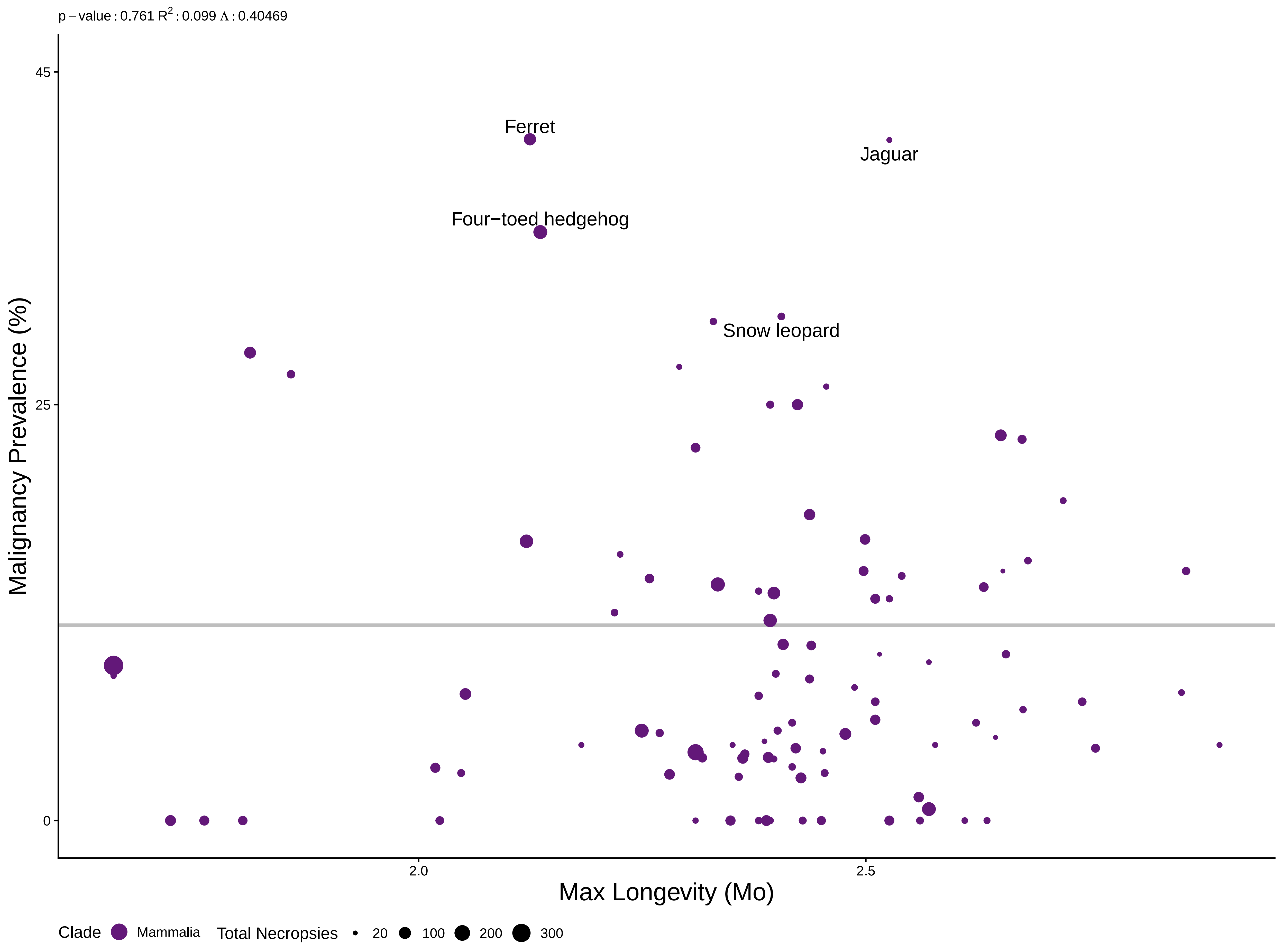

### S25bmrneomam.pdf

25 **Neoplasia Prevalence vs. Metabolic Rate in Mammals**

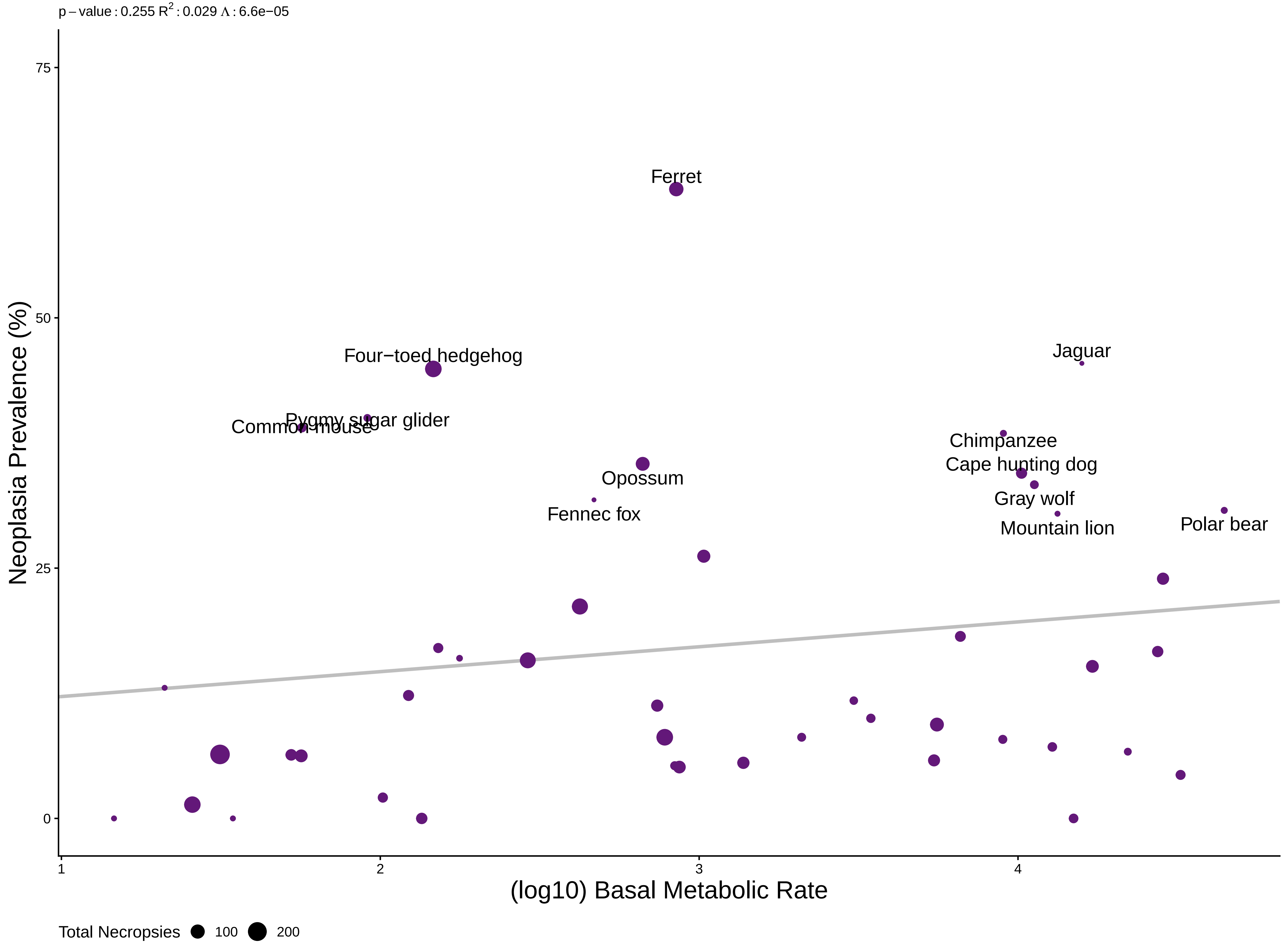

### S26bmrmalmam.pdf

26 **Malignancy Prevalence vs. Metabolic Rate in Mammals**

p – value : 0.782 $R^2$ : 0.077 Λ : 0.45

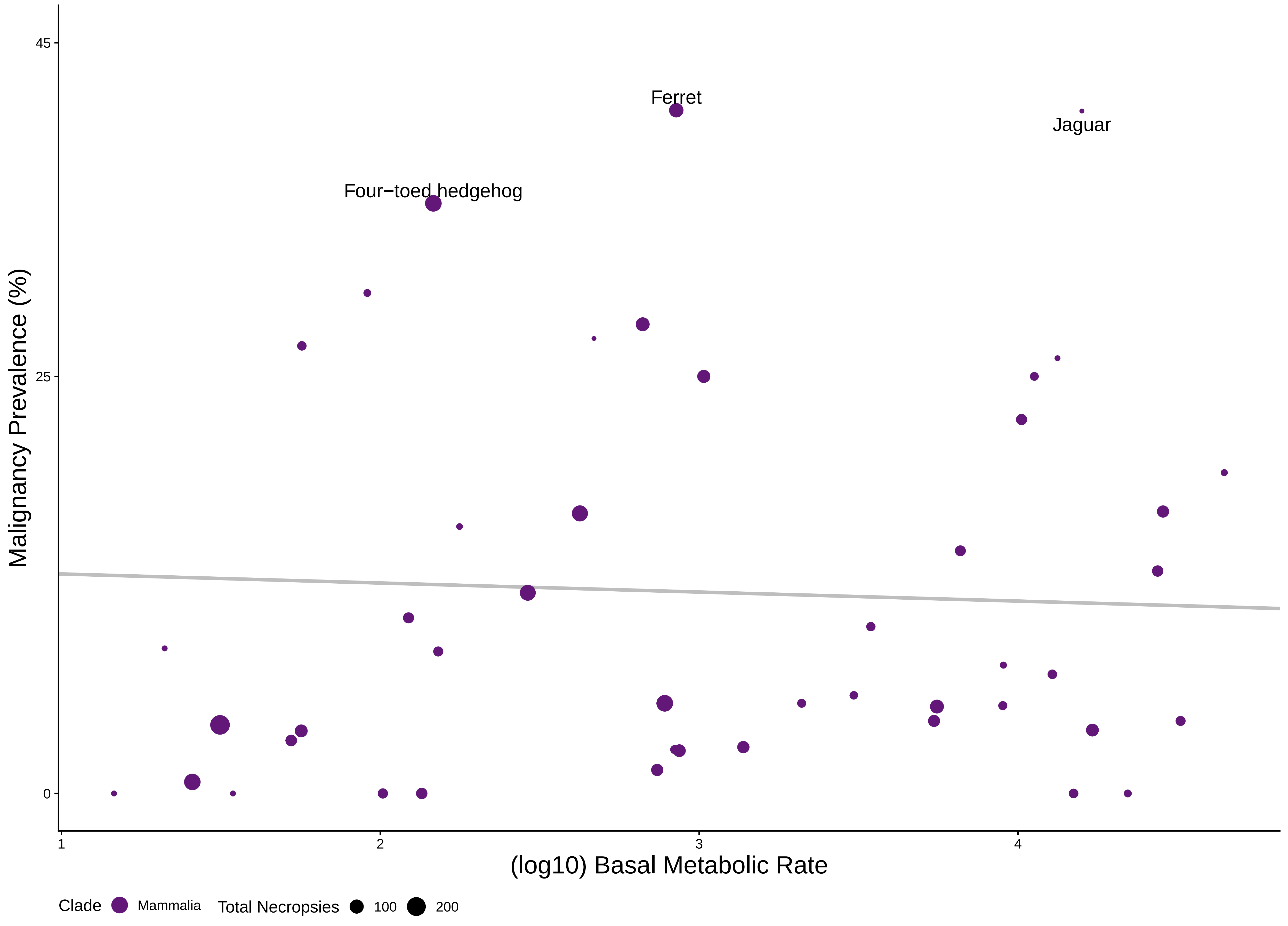

### S27wgtlongneomam.pdf

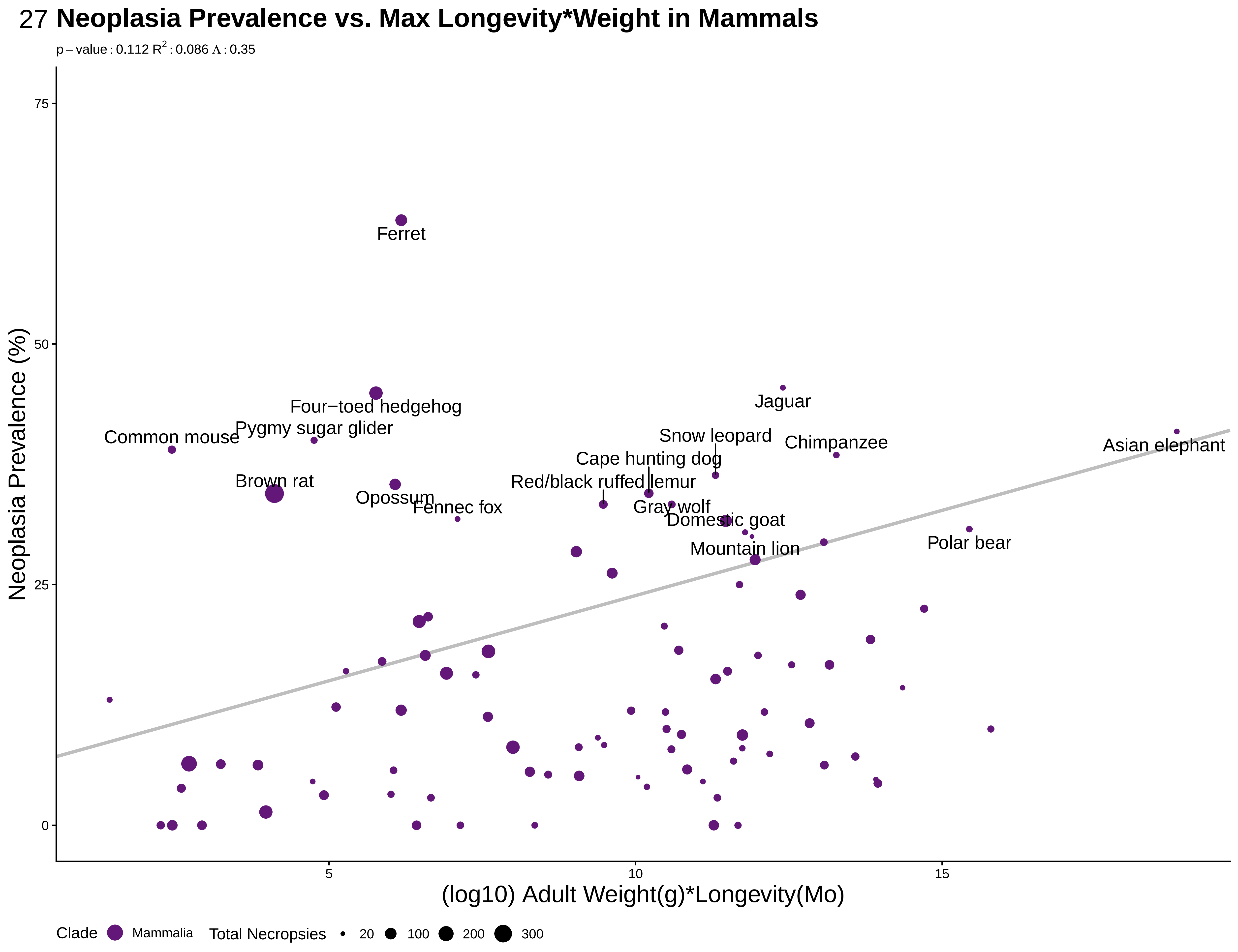

### S28wxlmal.pdf

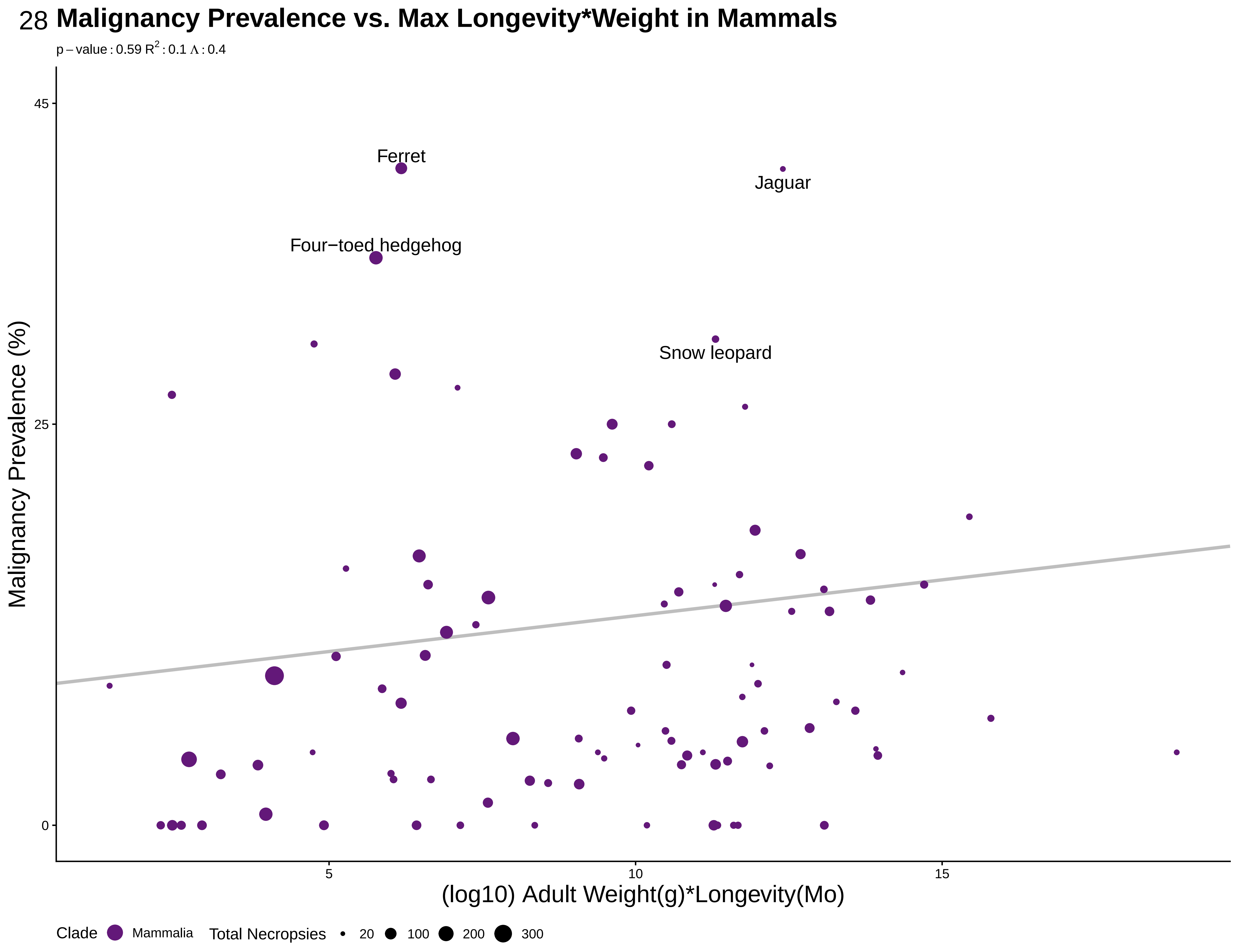

### S29lityearneomam.pdf

29 **Neoplasia Prevalence vs. Litters Per Year in Mammals**

p – value : 0.366 $R^2$ : 0.073 Λ : 0.3

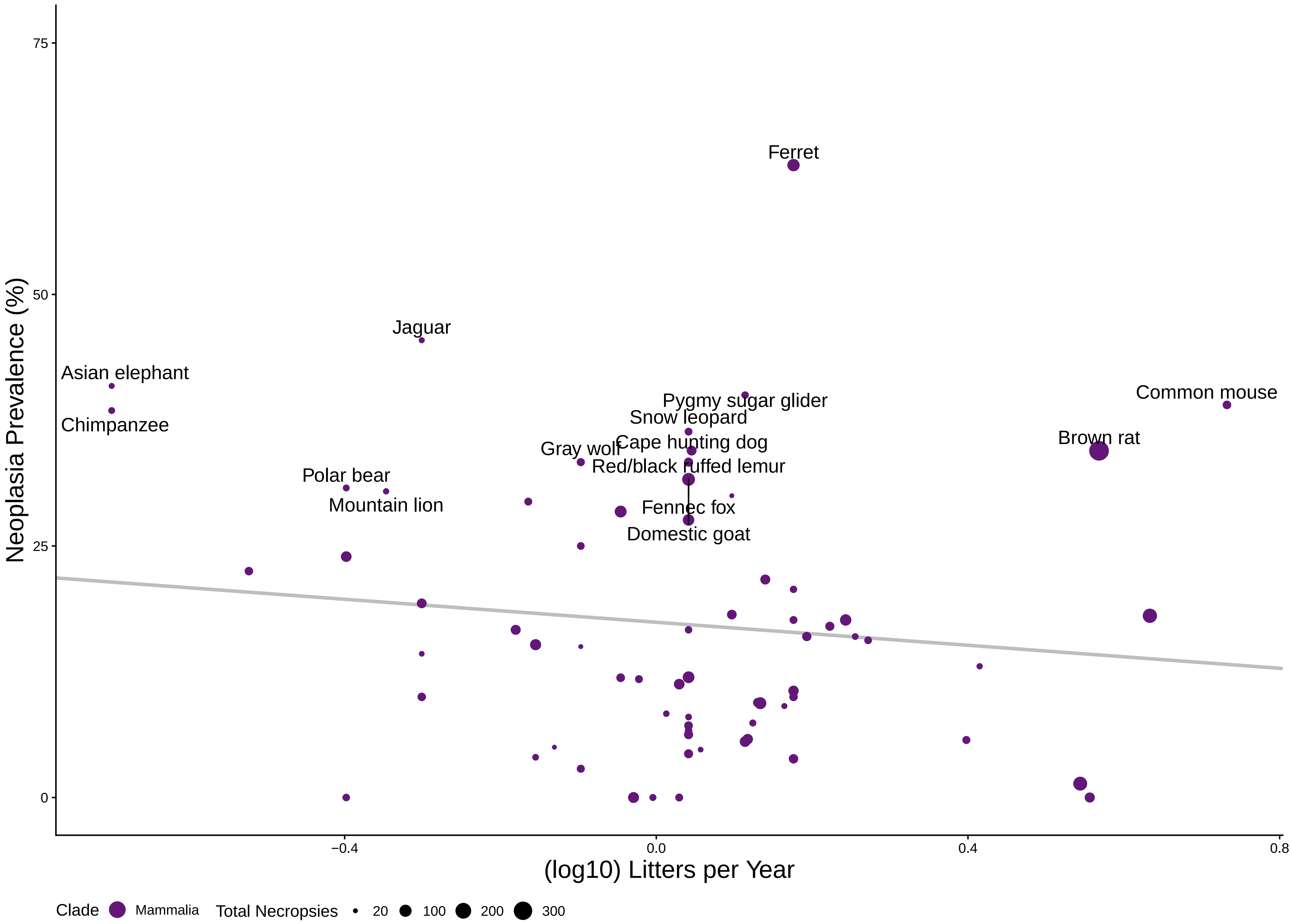

### S30lityearmalmam.pdf

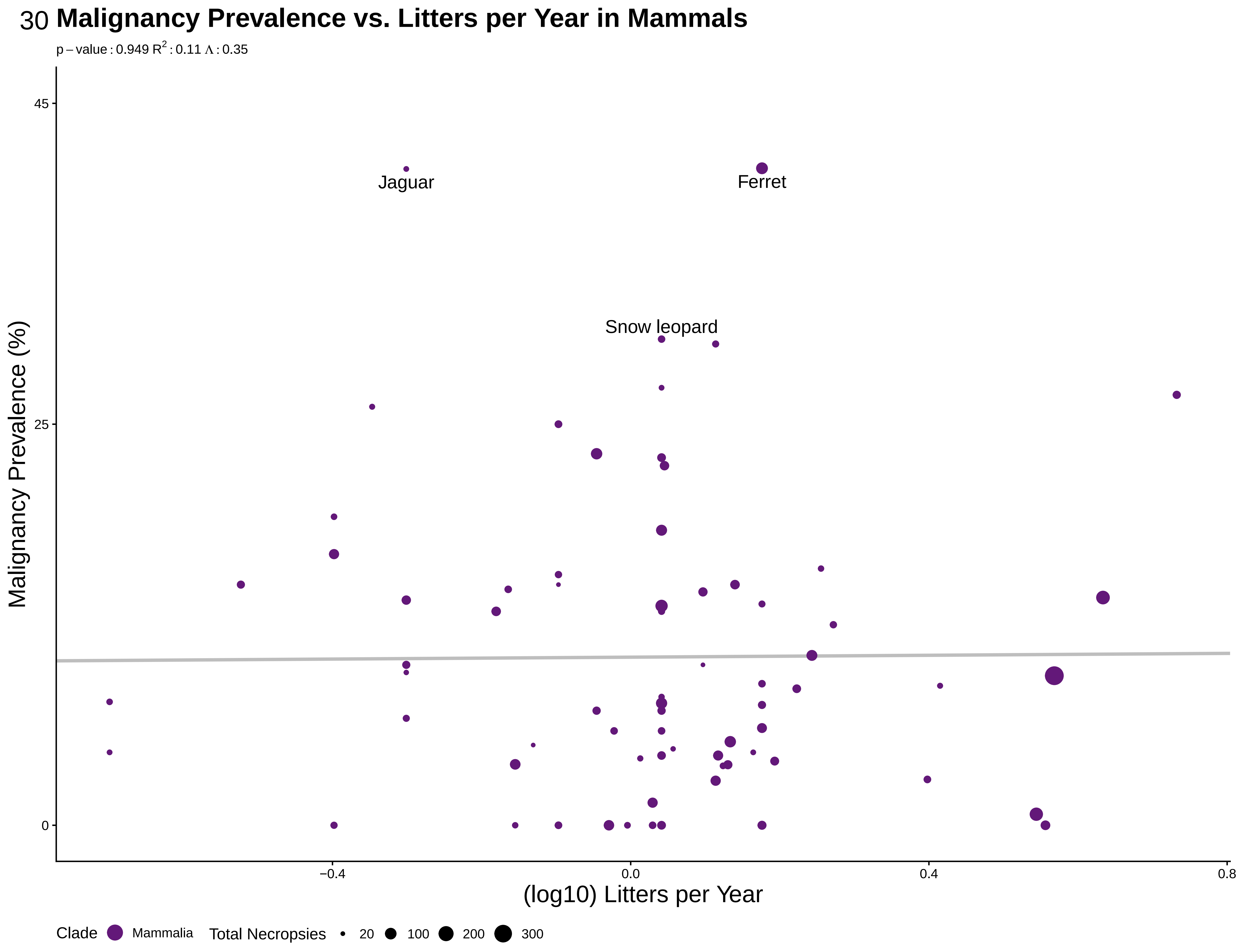

### S31femmatneomam.pdf

31 **Neoplasia Prevalence vs. Female Maturity in Mammals**

p − value : 0.294 $R^2$ : 0.083 Λ : 0.42

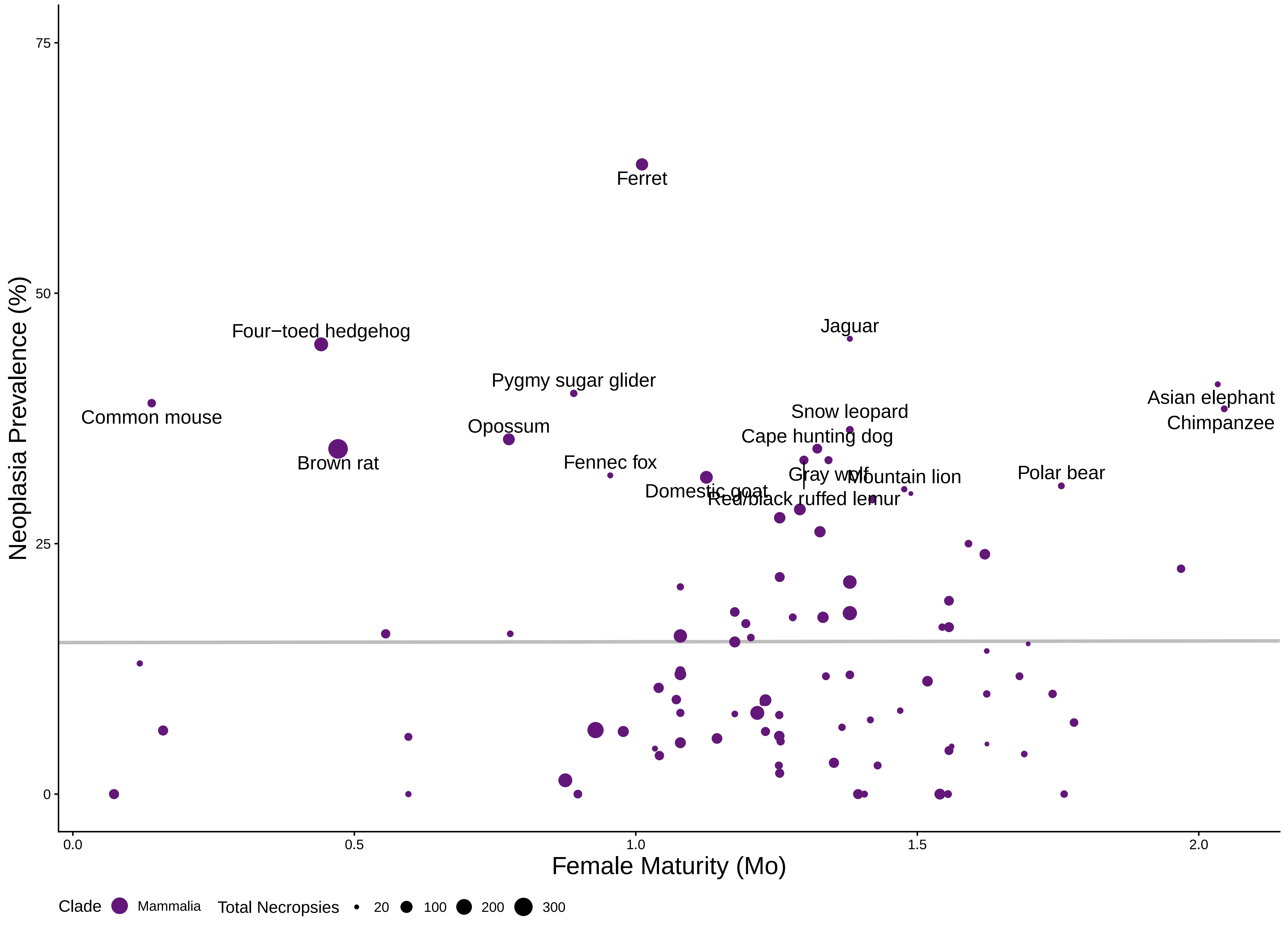

### S32femmatmalmam.pdf

#### Malignancy Prevalence vs. Female Maturity in Mammals

$p-value: 0.475$ $R^2: 0.11$ $\Lambda: 0.43$

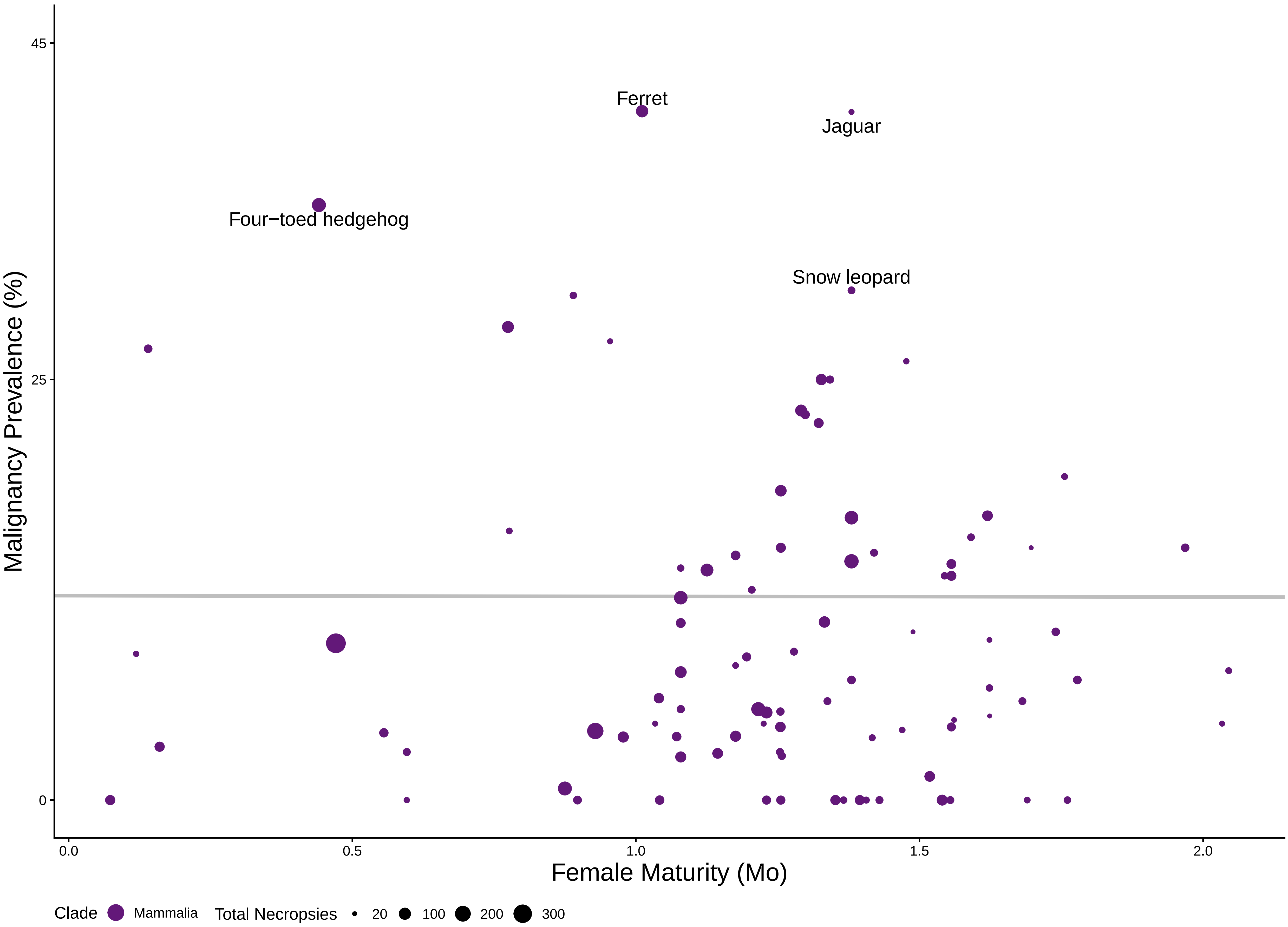

### S33malematneomam.pdf

33 **Neoplasia Prevalence vs. Male Maturity in Mammals**

p – value : 0.804 $R^2$ : 0.073 Λ : 0.42

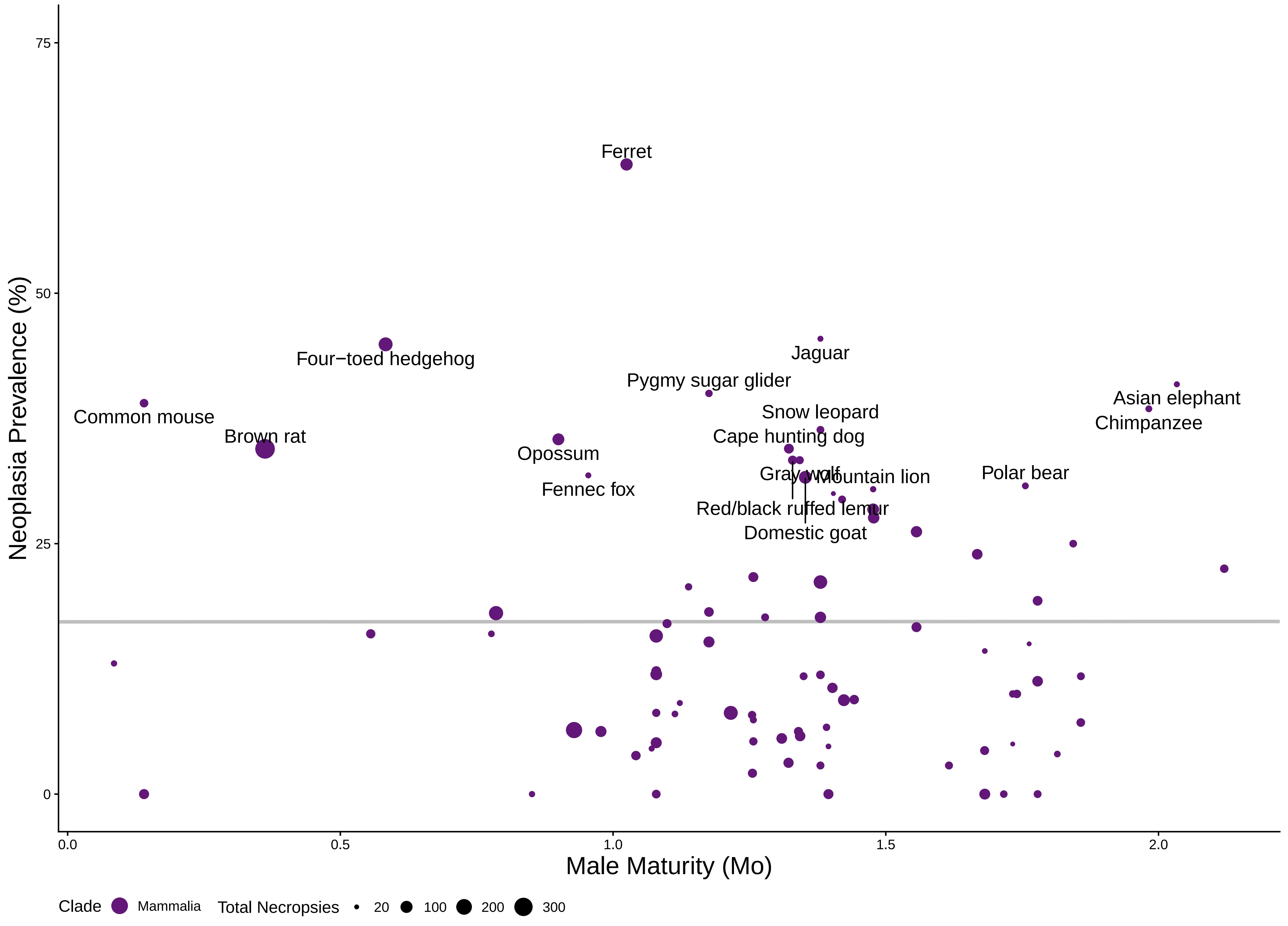

### S34malematmalmam.pdf

#### Malignancy Prevalence vs. Male Maturity in Mammals

p – value : 0.326 $R^2$ : 0.13 Λ : 0.43

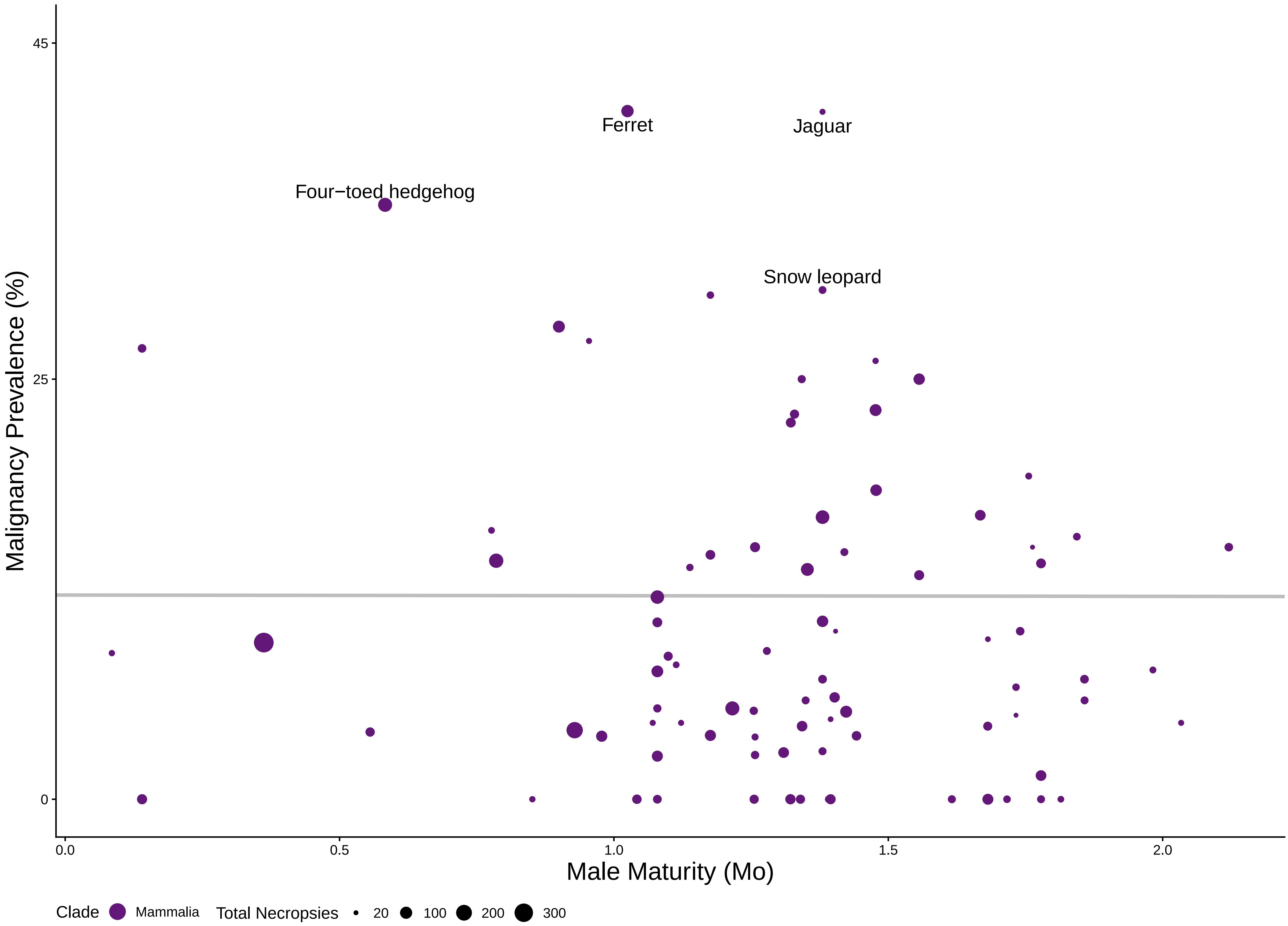

### S35weanneomam.pdf

35 **Neoplasia Prevalence vs. Weaning Weight in Mammals**

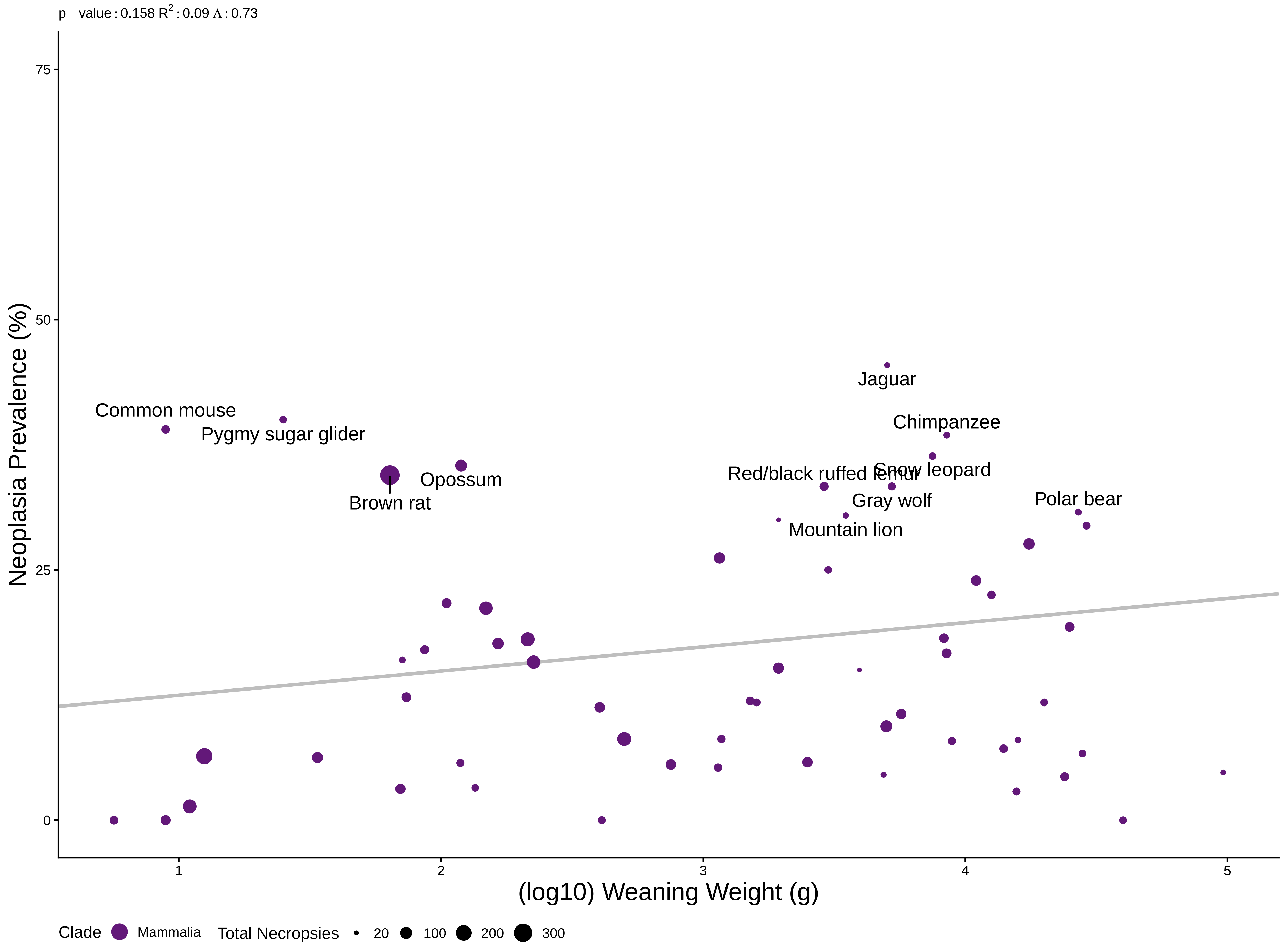

### S36weanmalmam.pdf

36 **Malignancy Prevalence vs. Weaning Weight in Mammals**

p – value : 0.281 $R^2$ : 0.00084 $\Lambda$ : 6.6e−05

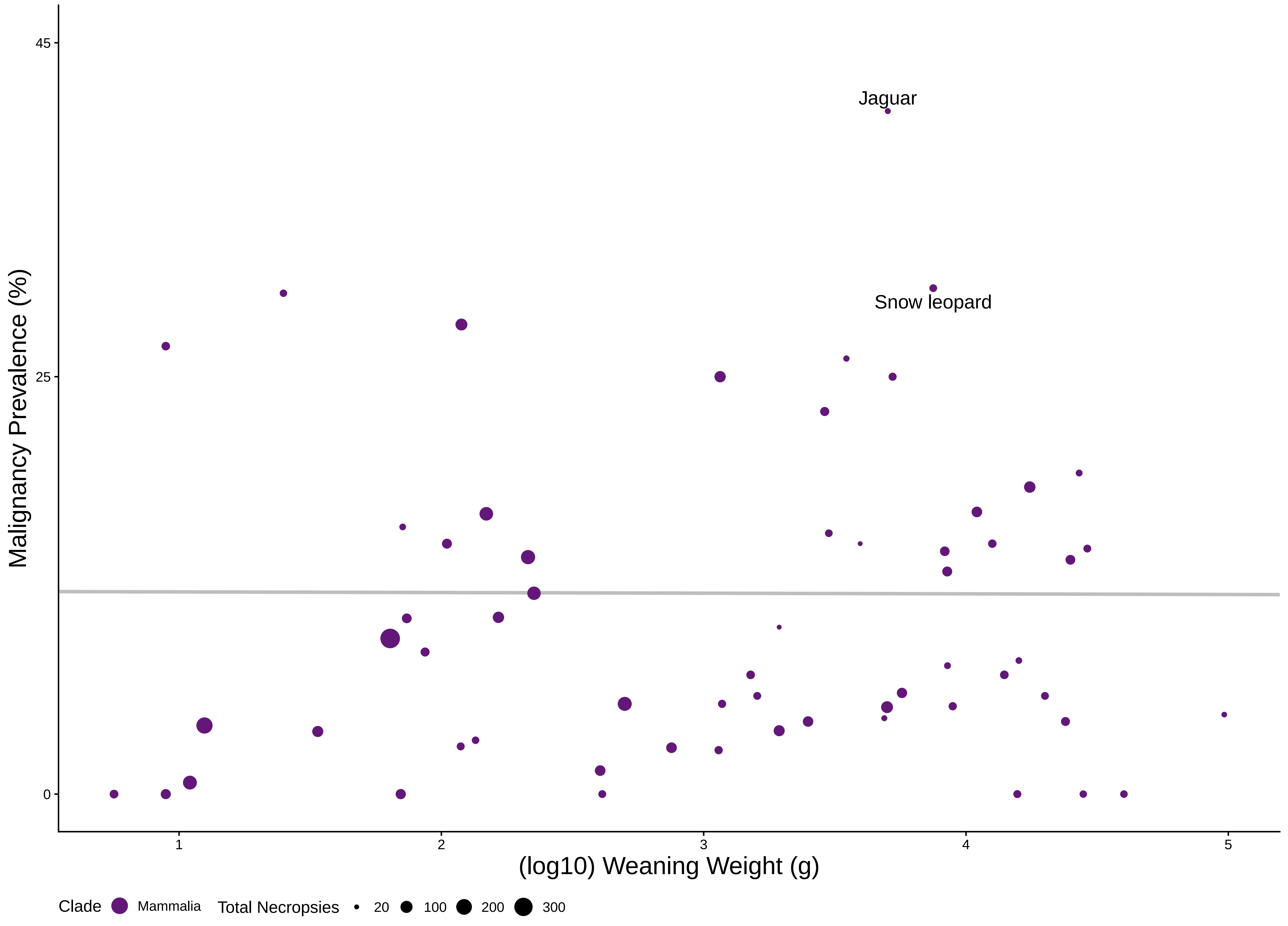

### S37growneomam.pdf

37 **Neoplasia Prevalence vs. Growth Rate in Mammals**

p – value : 0.368 $R^2$ : 0.021 Λ : 6.6e−05

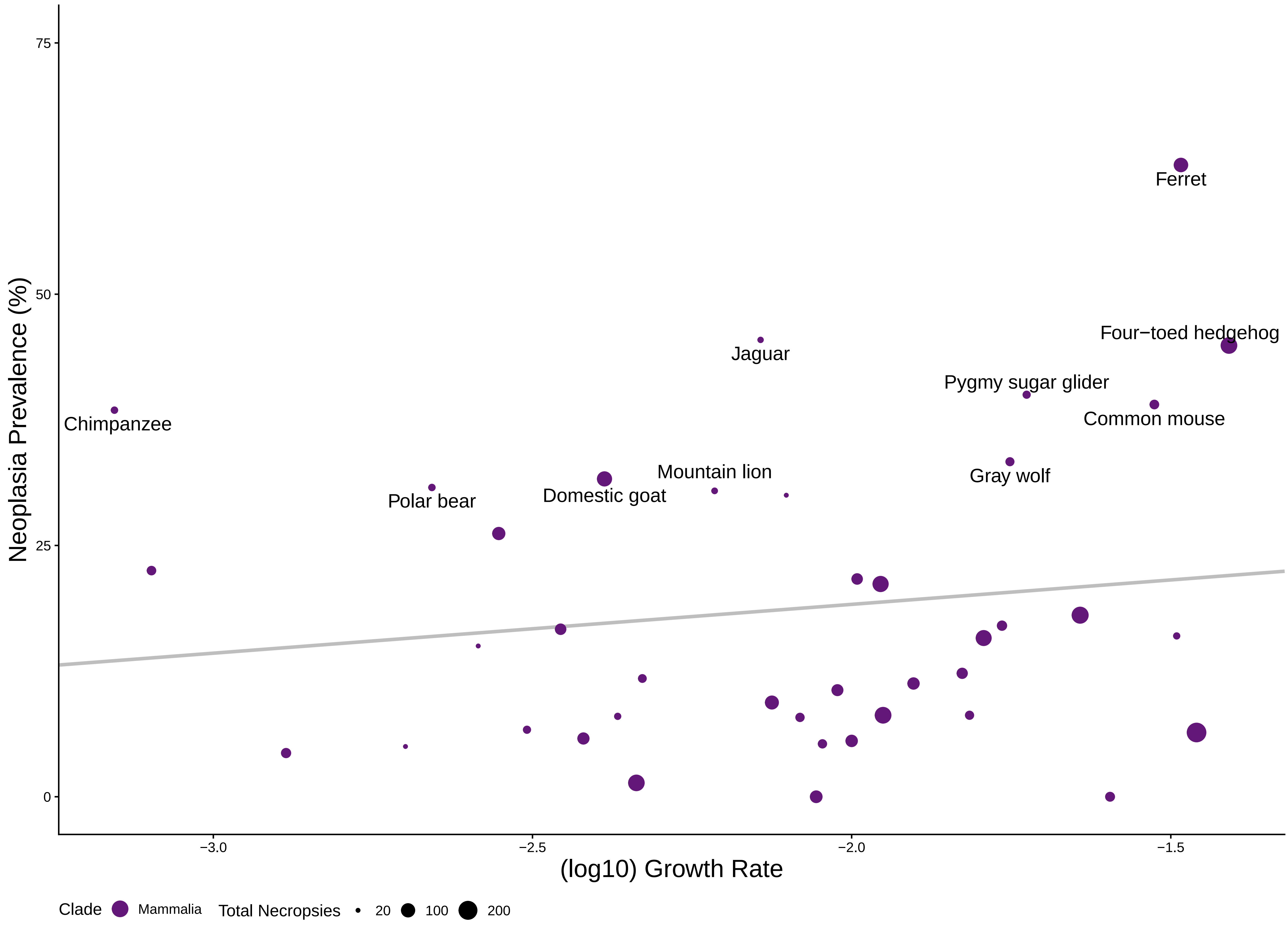

### S38growmalmam.pdf

38 **Malignancy Prevalence vs. Growth Rate in Mammals**

p − value : 0.12 $R^2$ : 0.061 Λ : 6.6e−05

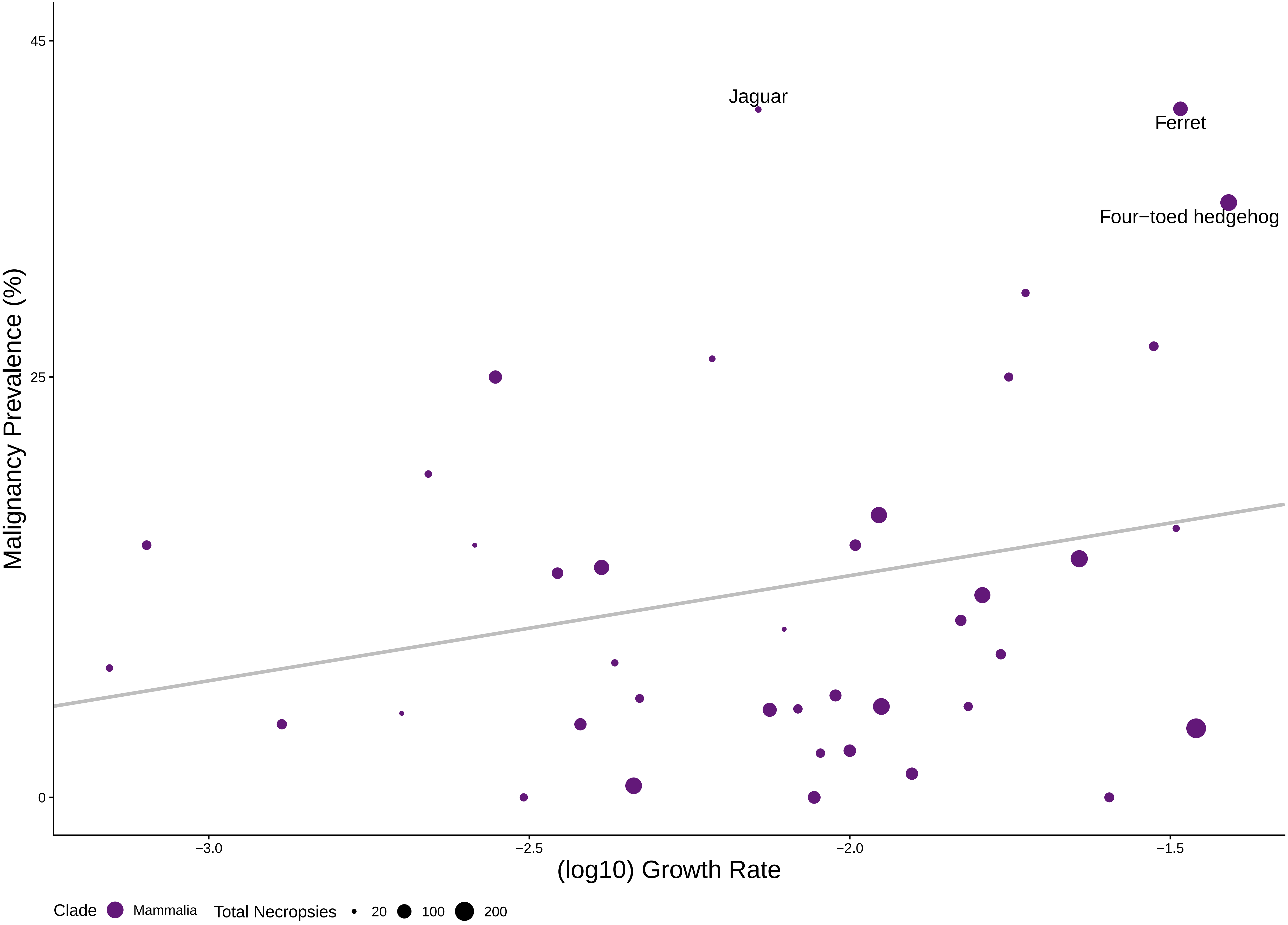

### S39wgtgestneomam.pdf

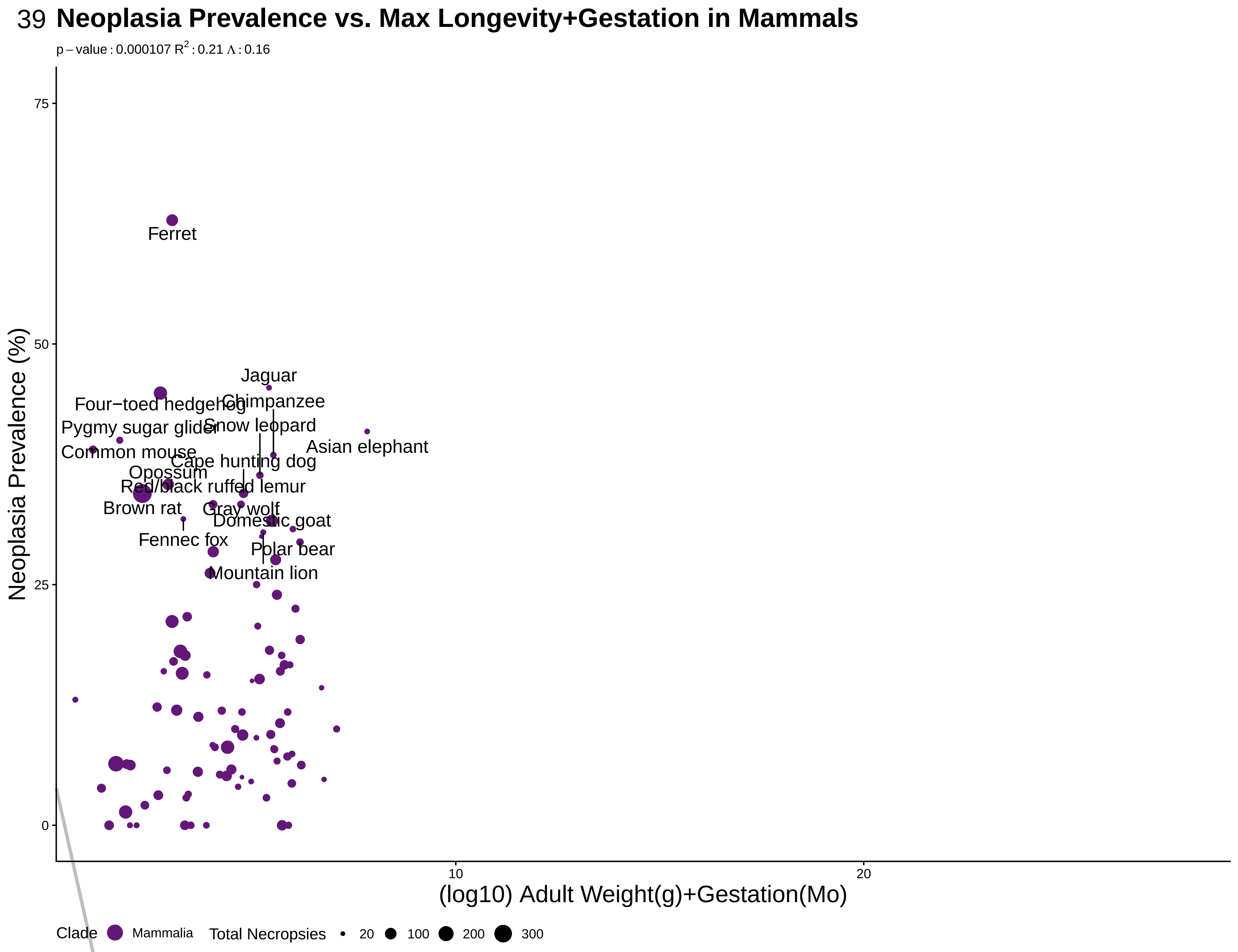

### S40wgtgestmalmam.pdf

40

#### Malignancy Prevalence vs. Max Longevity+Gestation in Mammals

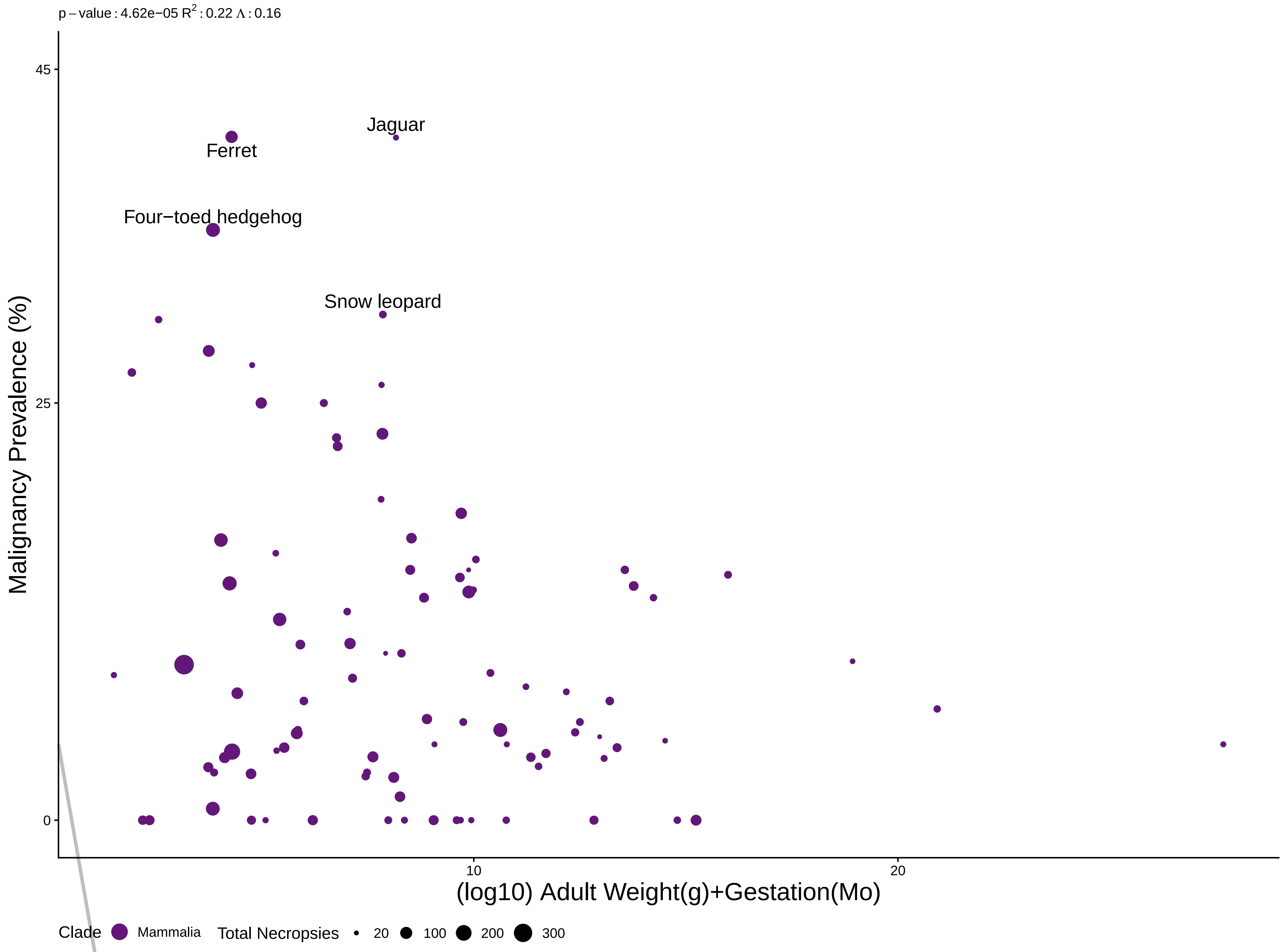

### S41RADAUC10.pdf

#### Neoplasia vs. AUC 10Gy Radiation

p − value : 0.35 $R^2$ : 0.079 Λ : 6.6e−05

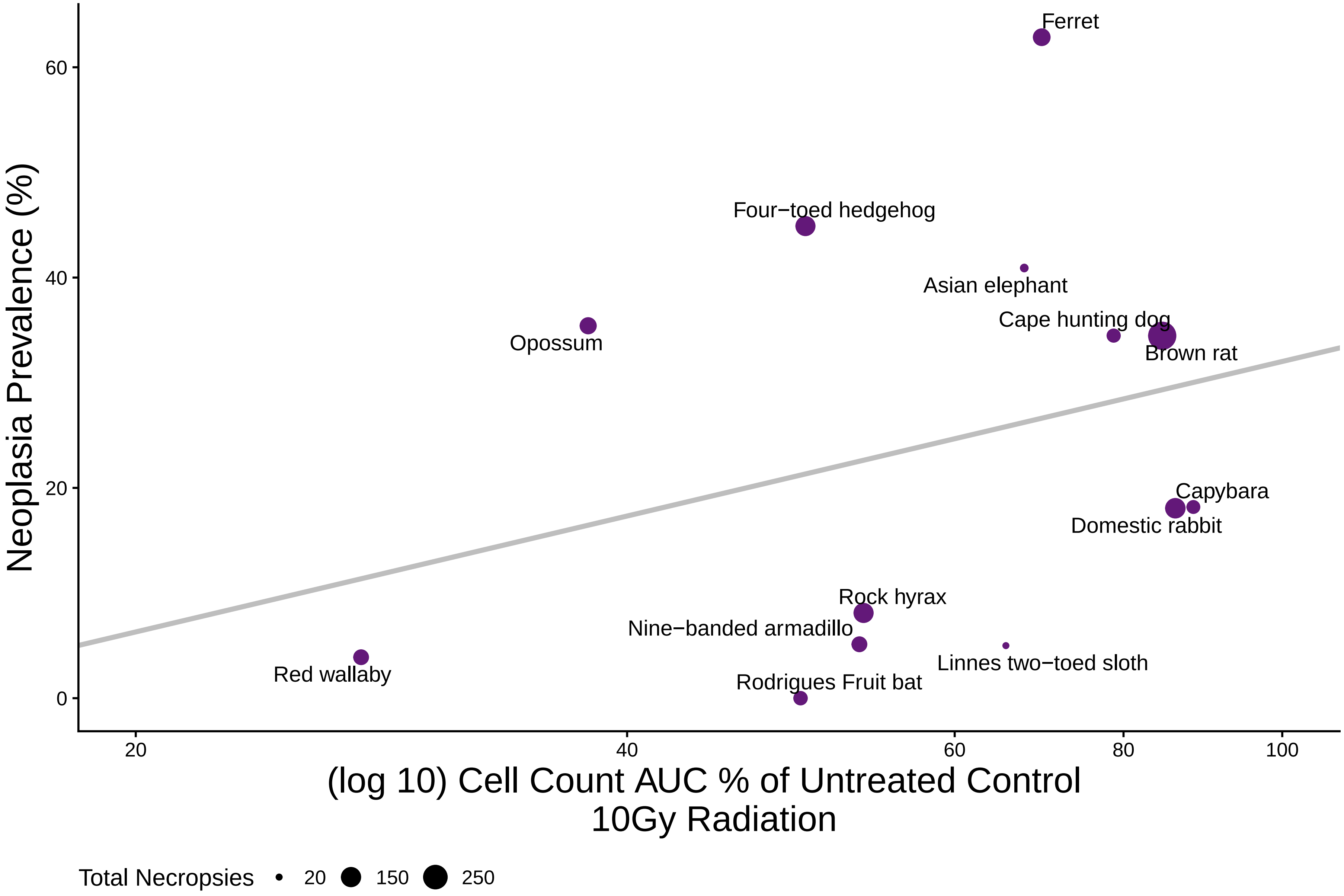

### S42RADAUC2.pdf

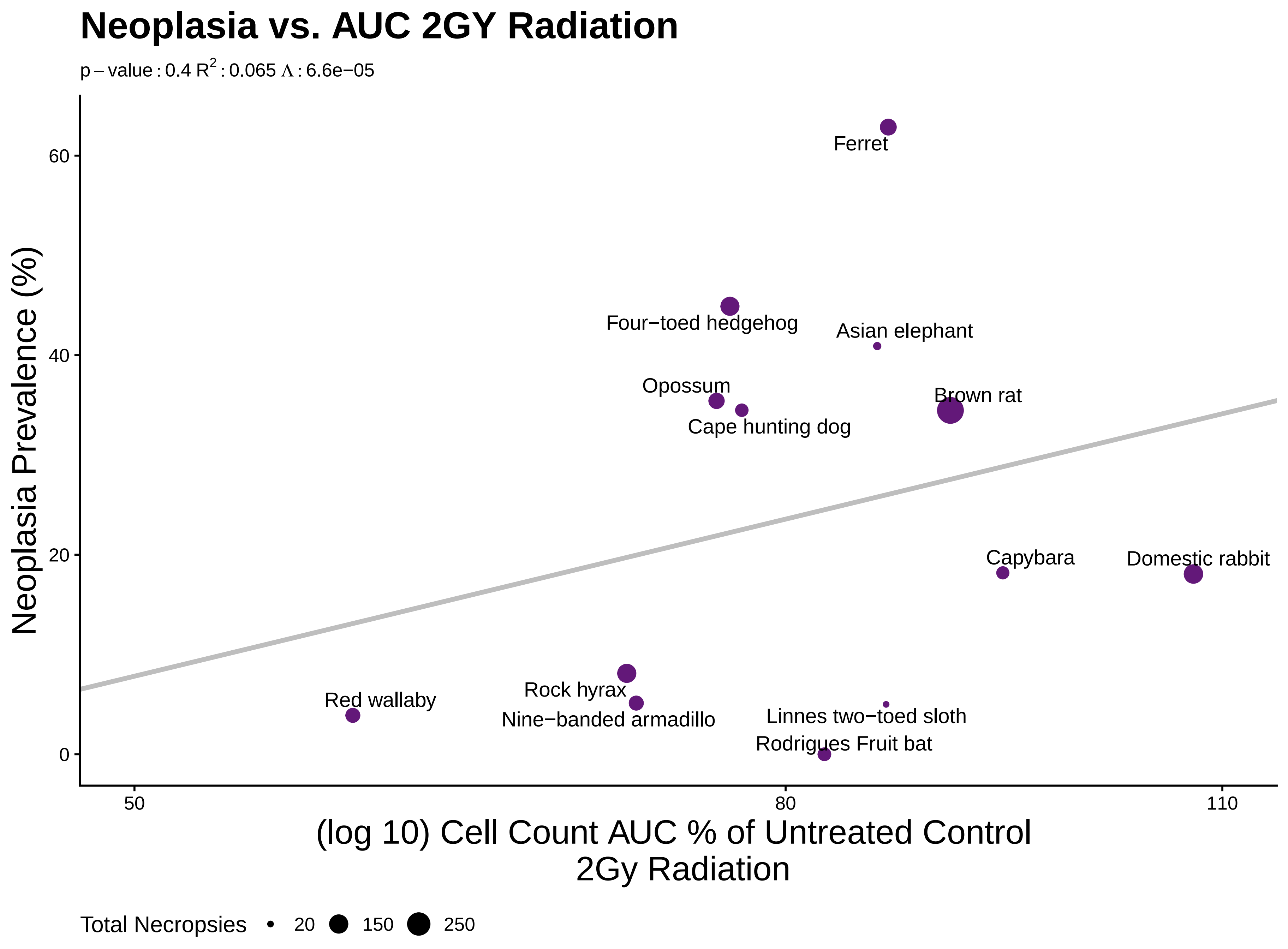

### S43RADAUC04.pdf

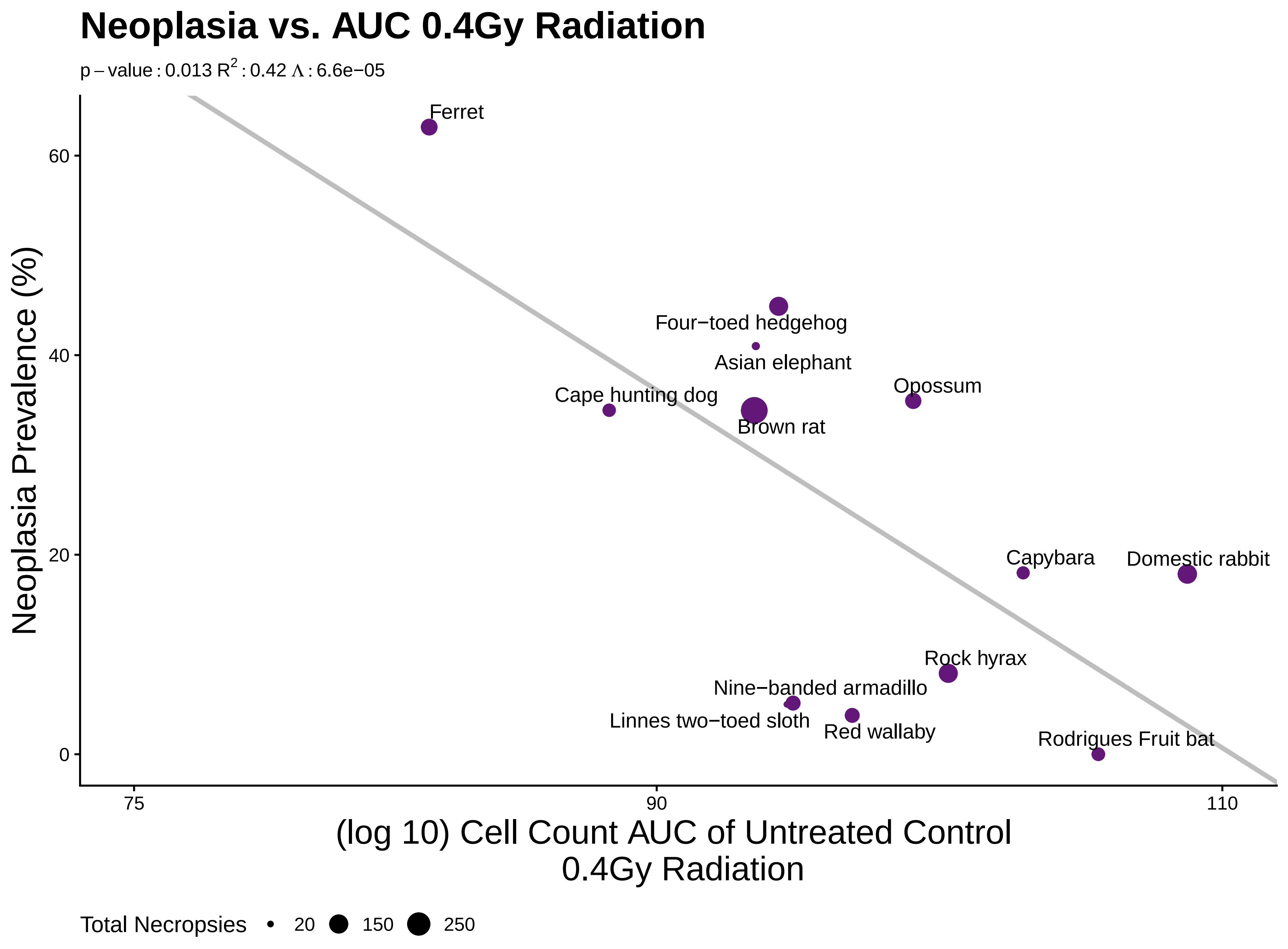

### S44RADChange10.pdf

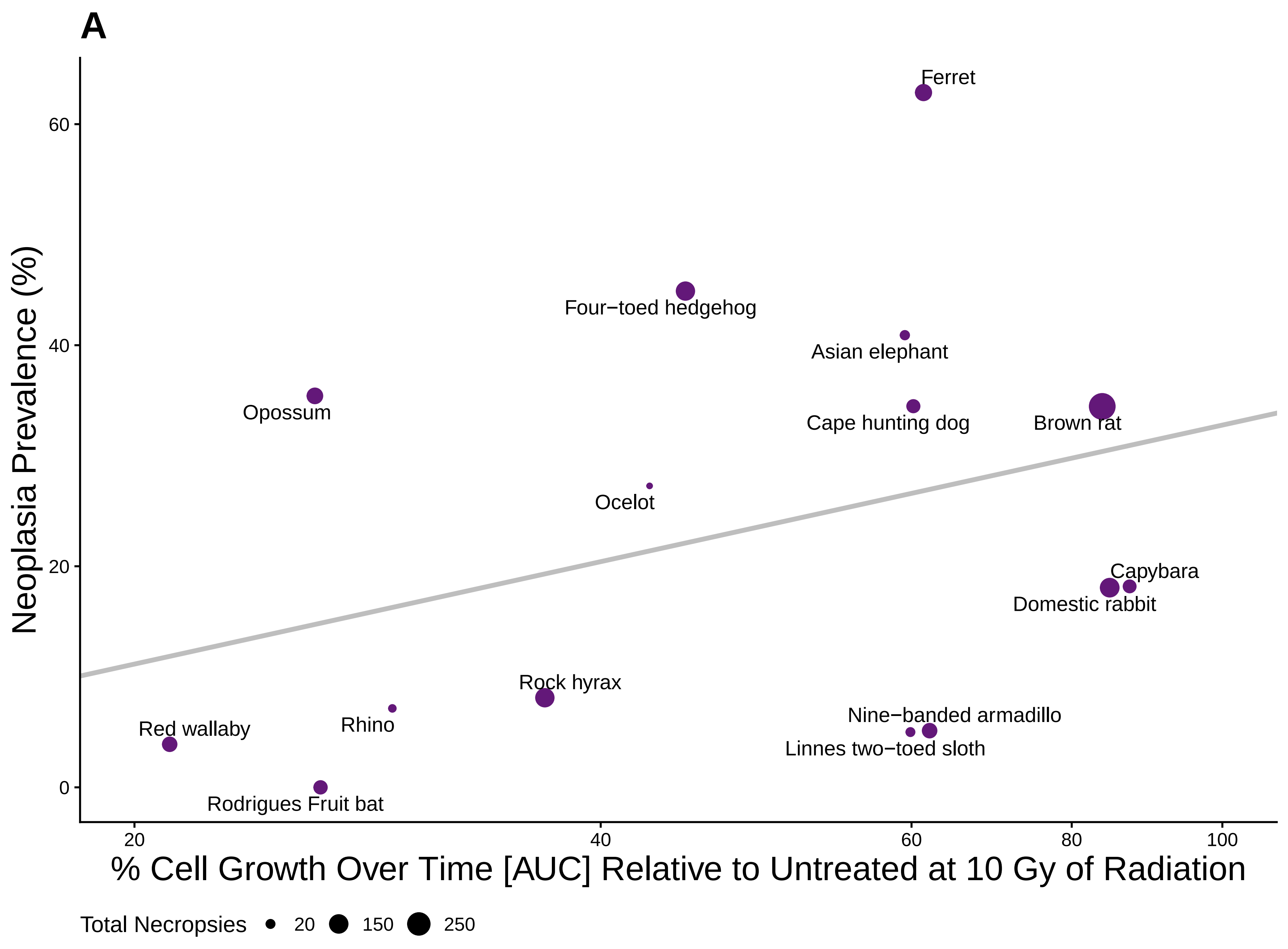

### S45RADChange2.pdf

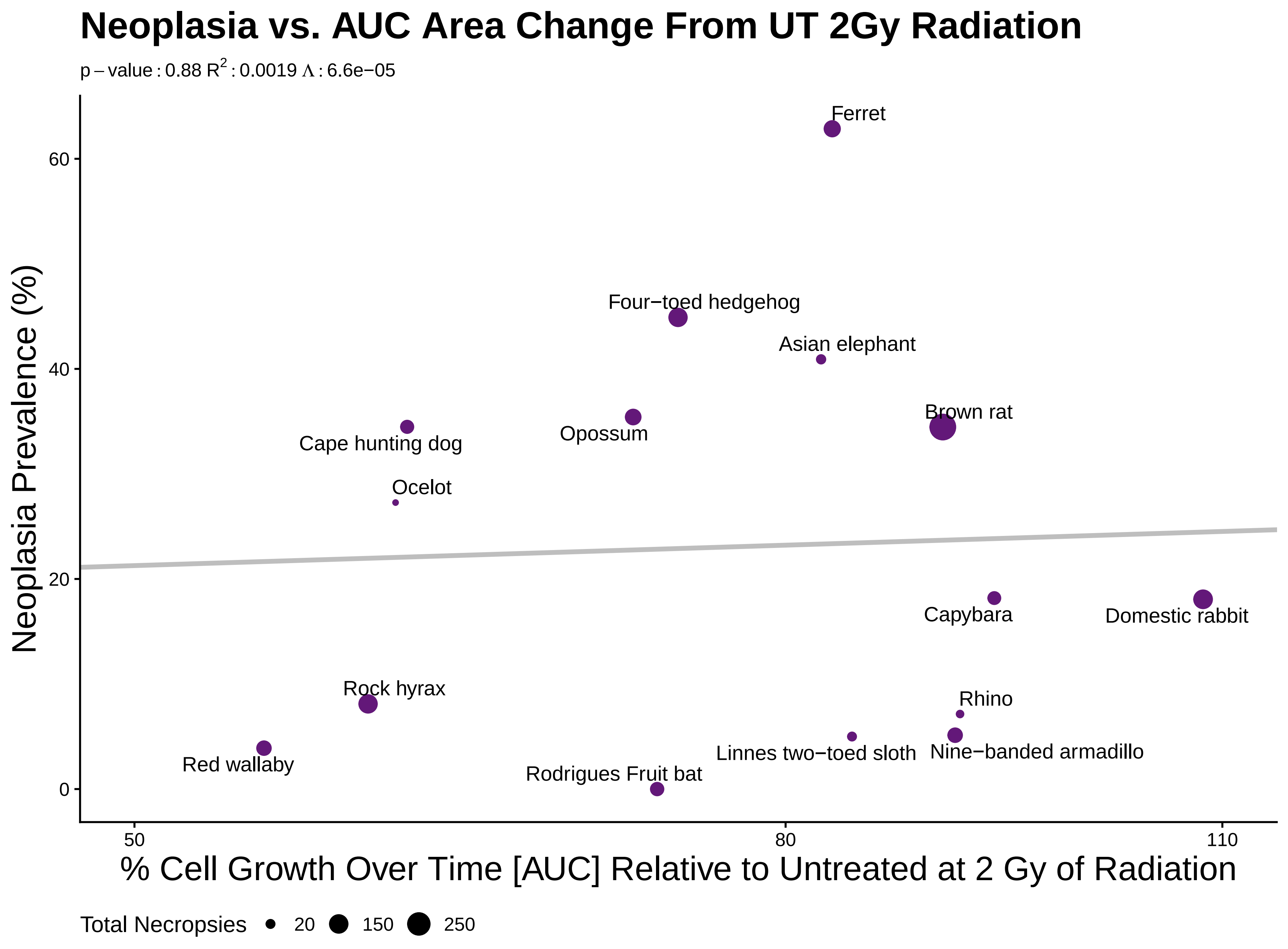

### S46RADChange10mal.pdf

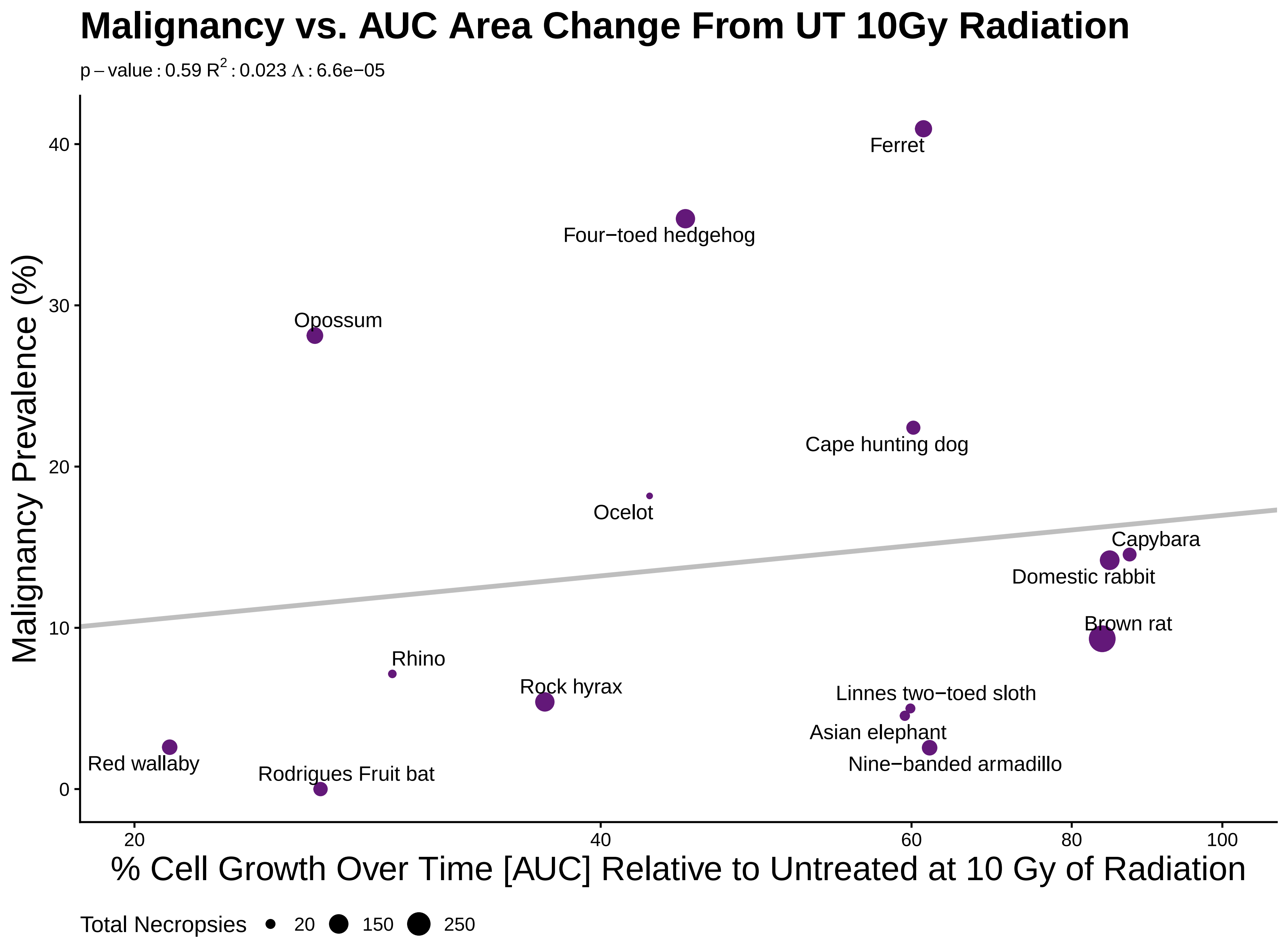

### S47RADChange2mal.pdf

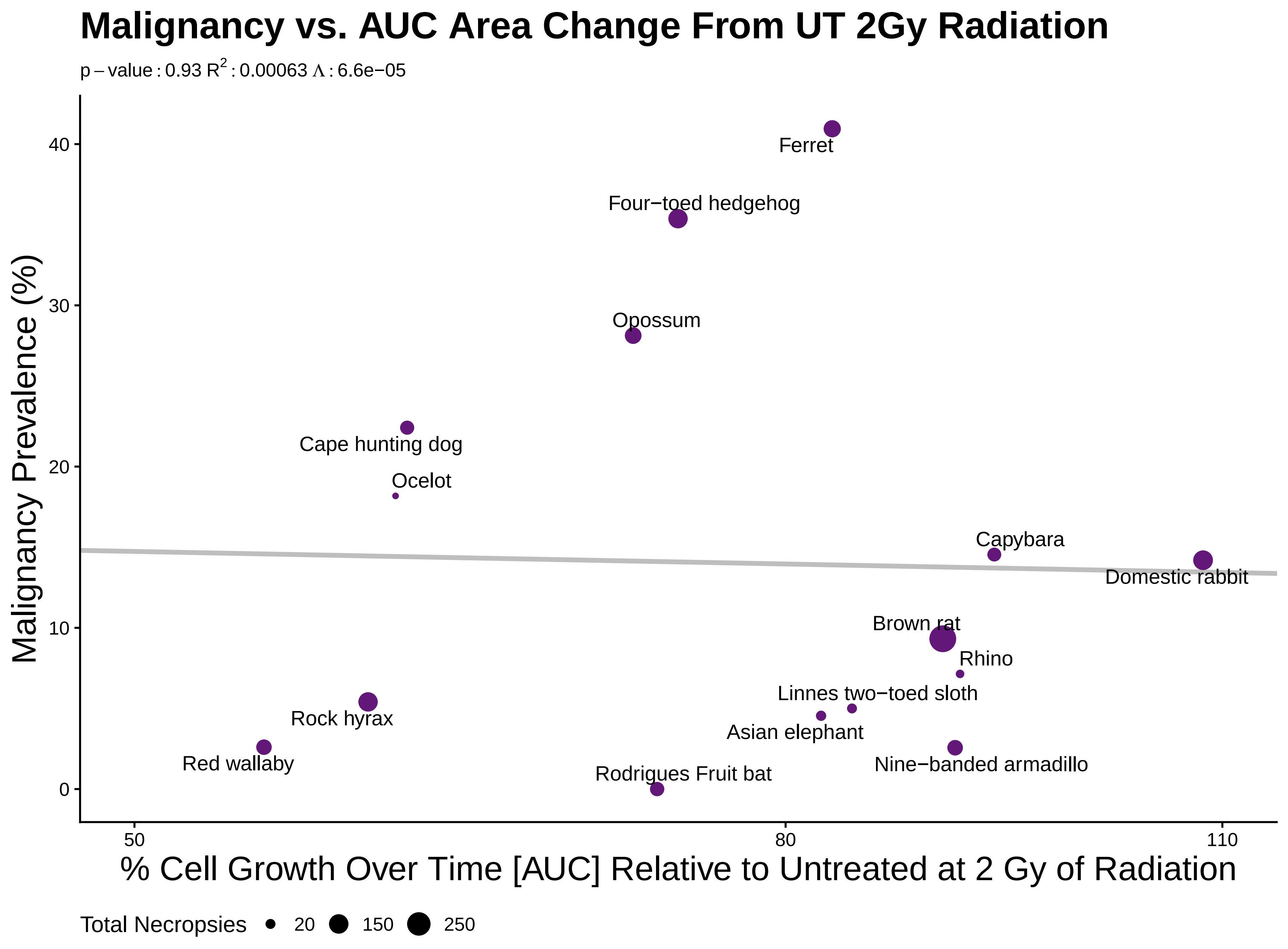

### S48RADChange04mal.pdf

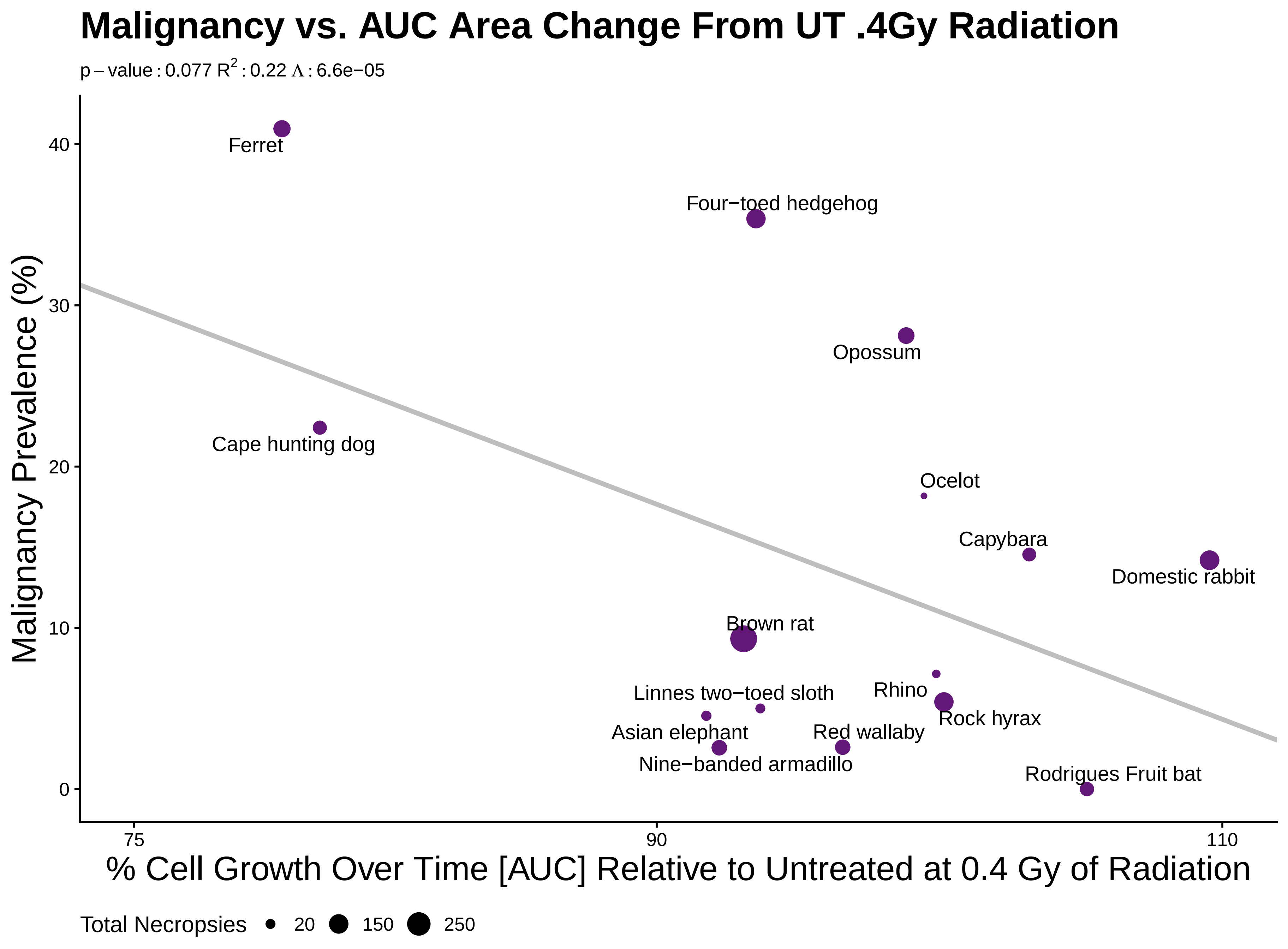

### S49RADChange04.pdf

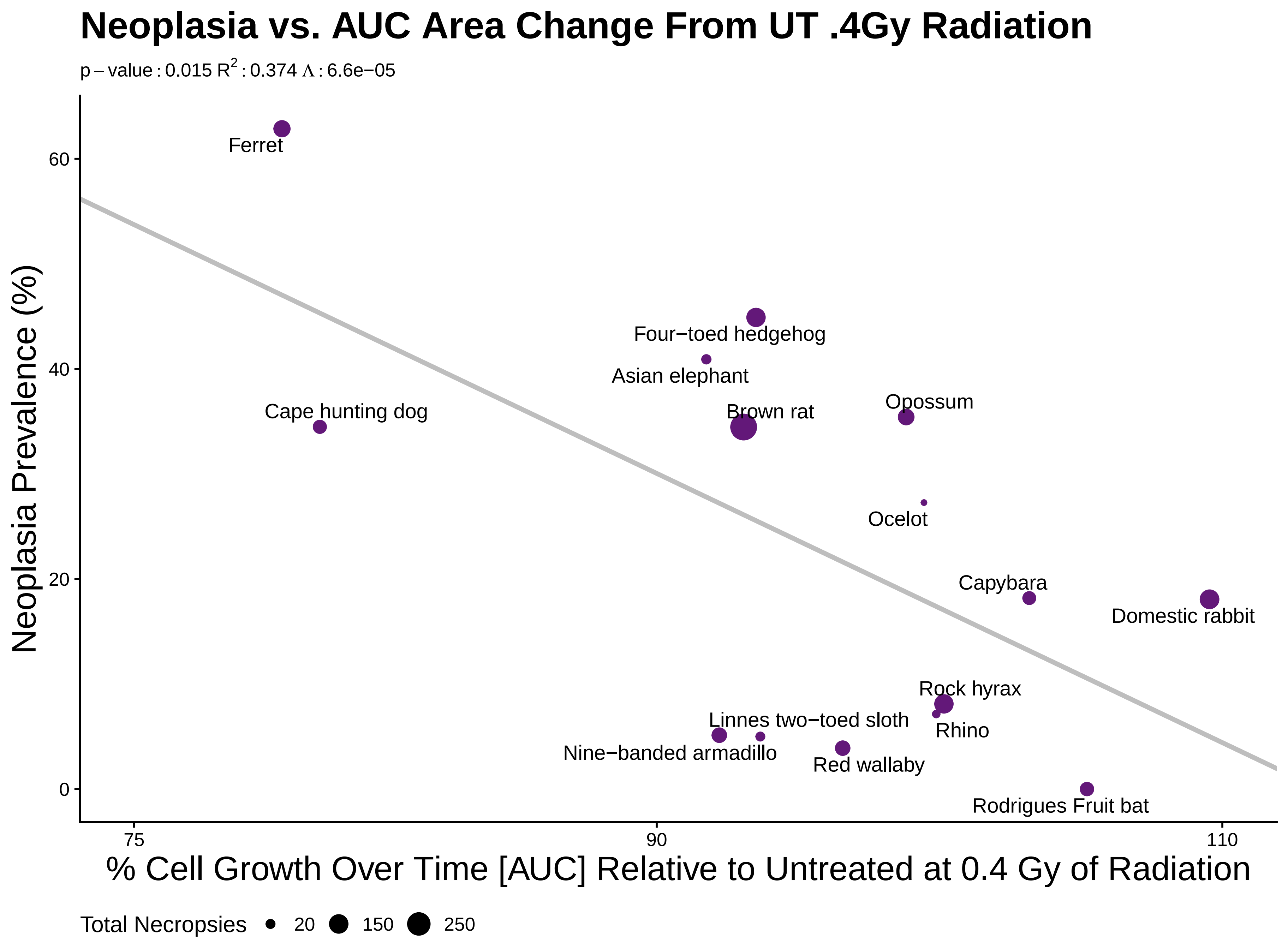

### S50mutation.mal.png

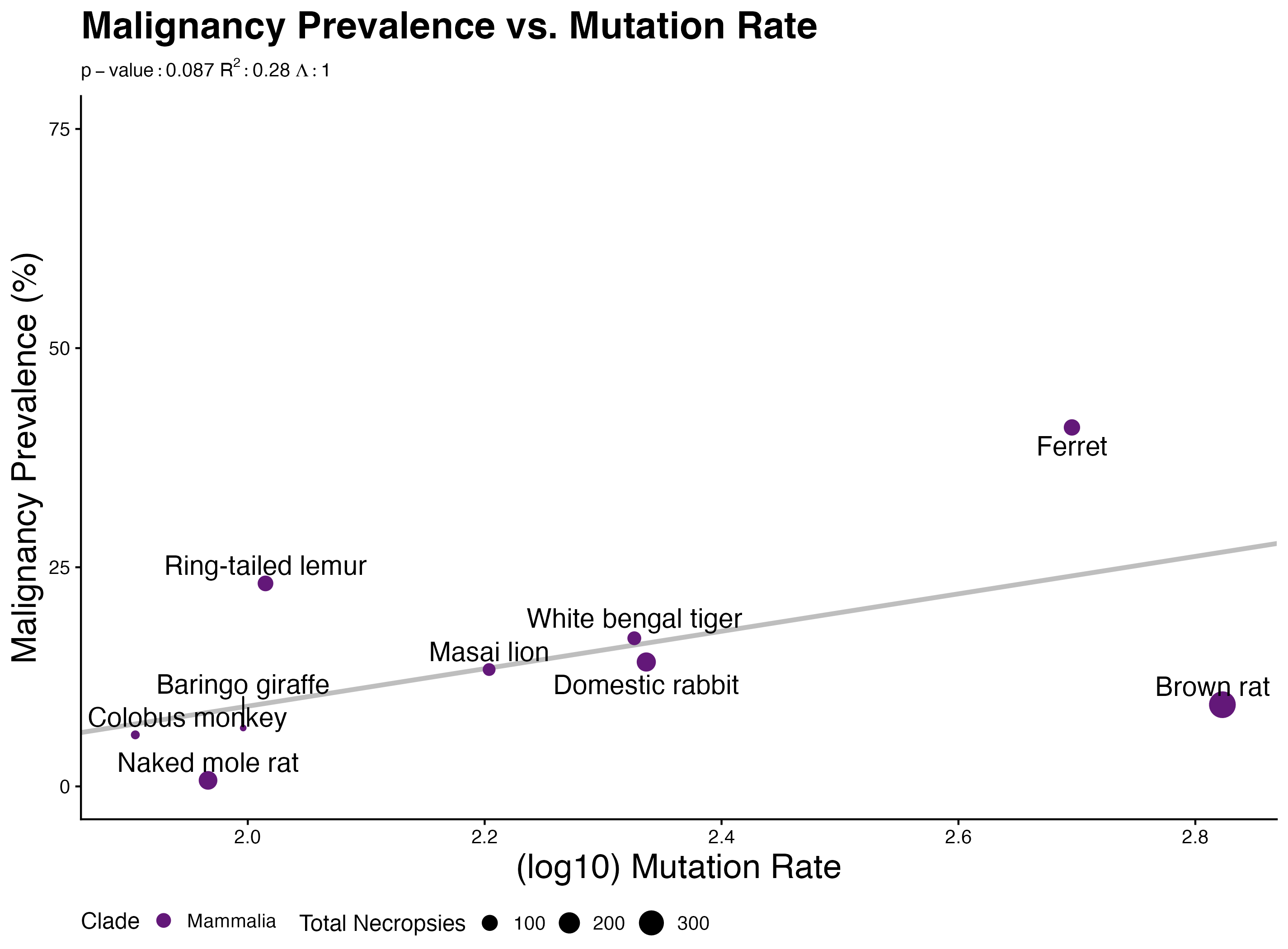

### S51anvfold721.pdf

#### Neoplasia vs. AUC 1uM Doxorubicin

p – value : 0.345 $R^2$ : 0.0814 Λ : 6.6107e−05

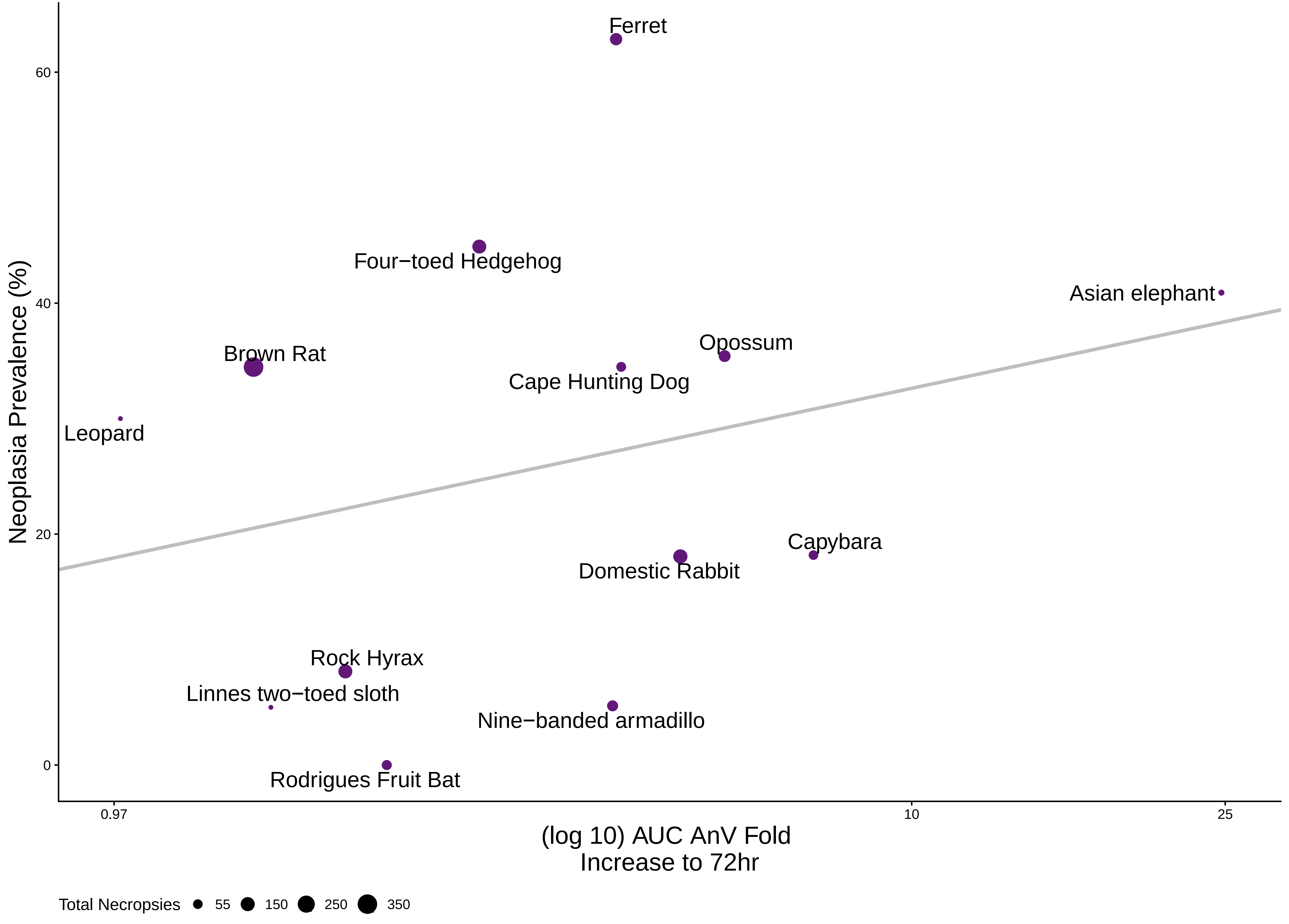

### S52anvfold72.33.pdf

#### Neoplasia vs. AUC 0.33uM Doxorubicin

p – value : 0.189 $R^2$ : 0.151 $\Lambda$ : 6.6107e−05

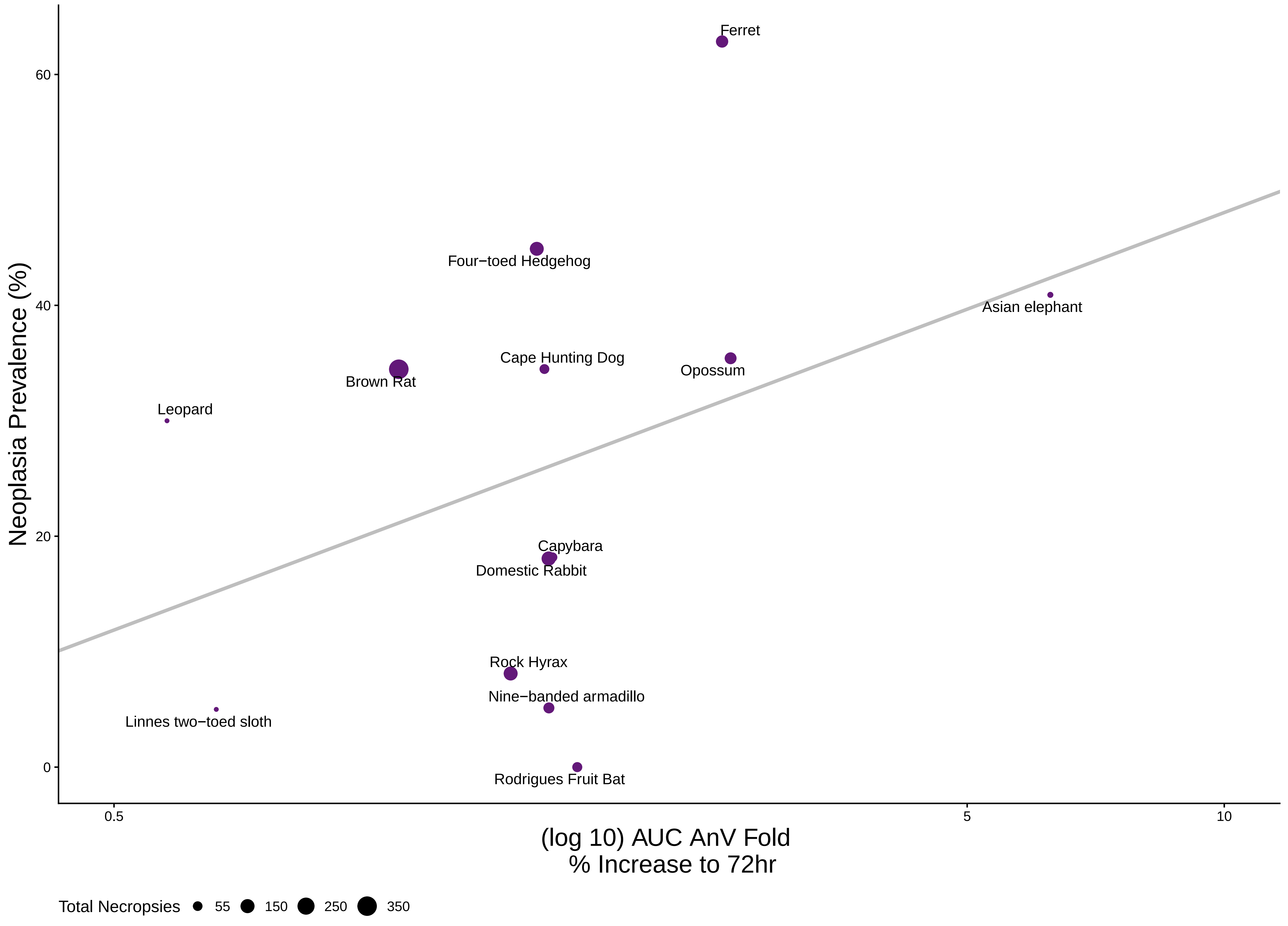

### S53anvfold72.11.pdf

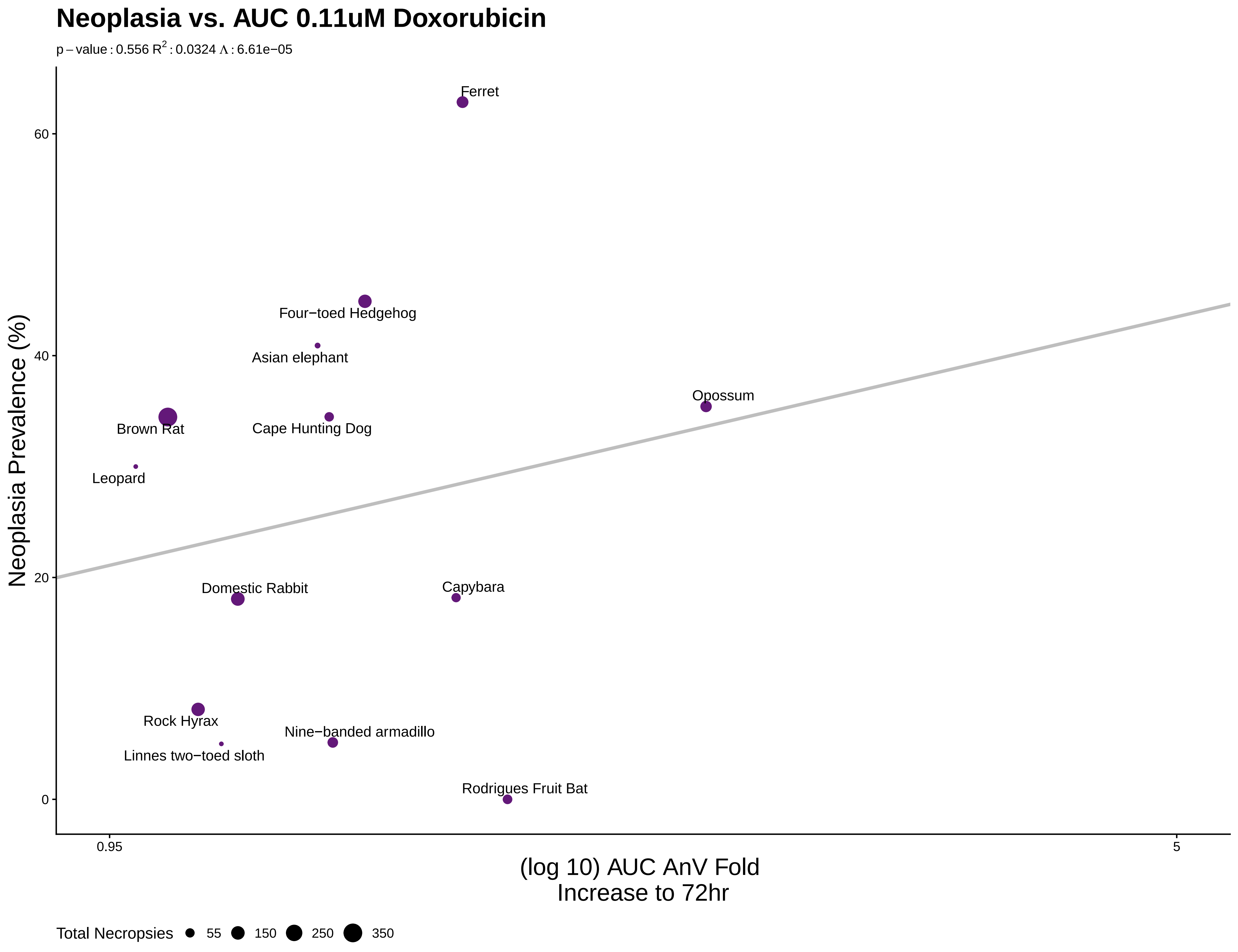

### S54celldeath72.1.pdf

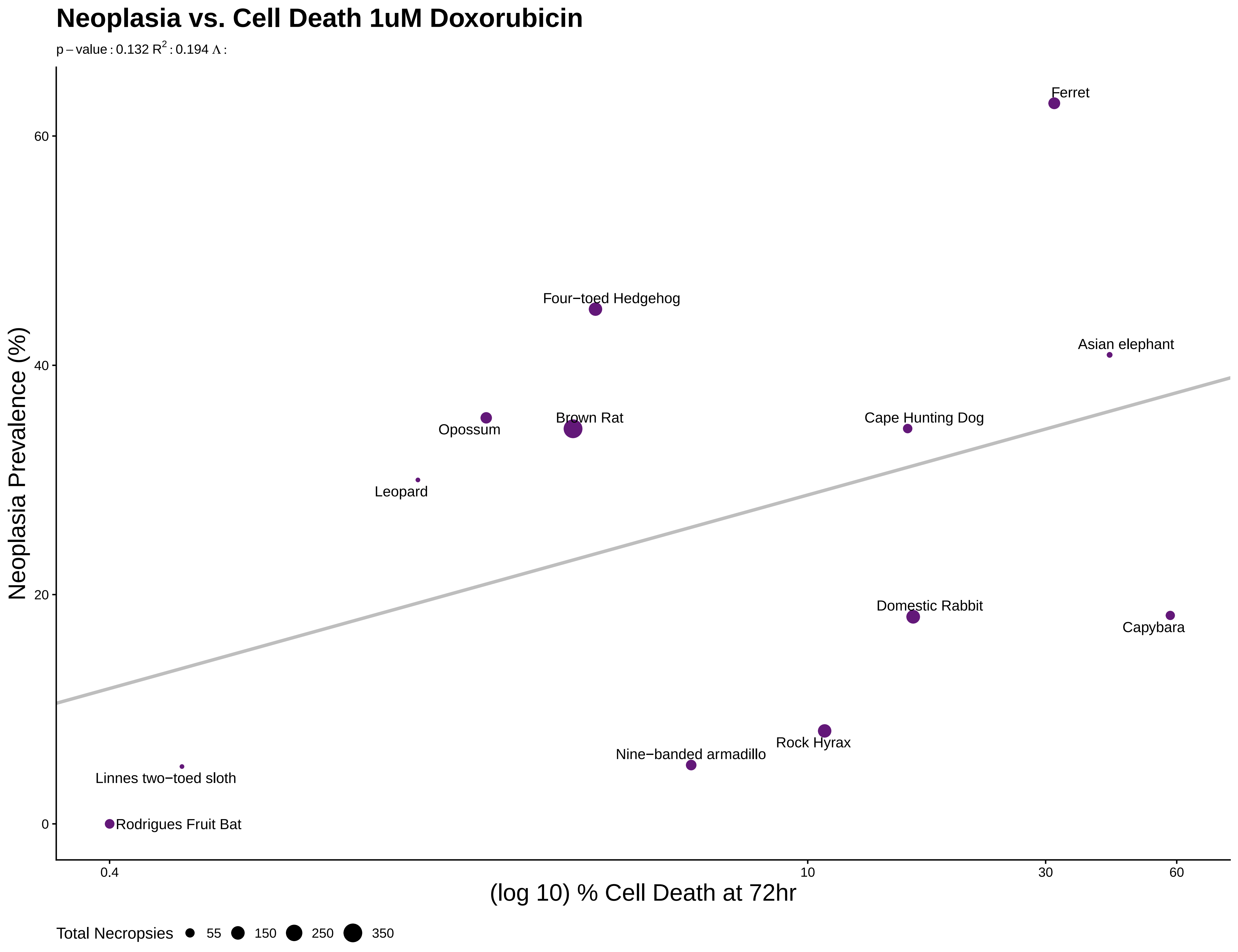

### S55celldeath72.33.pdf

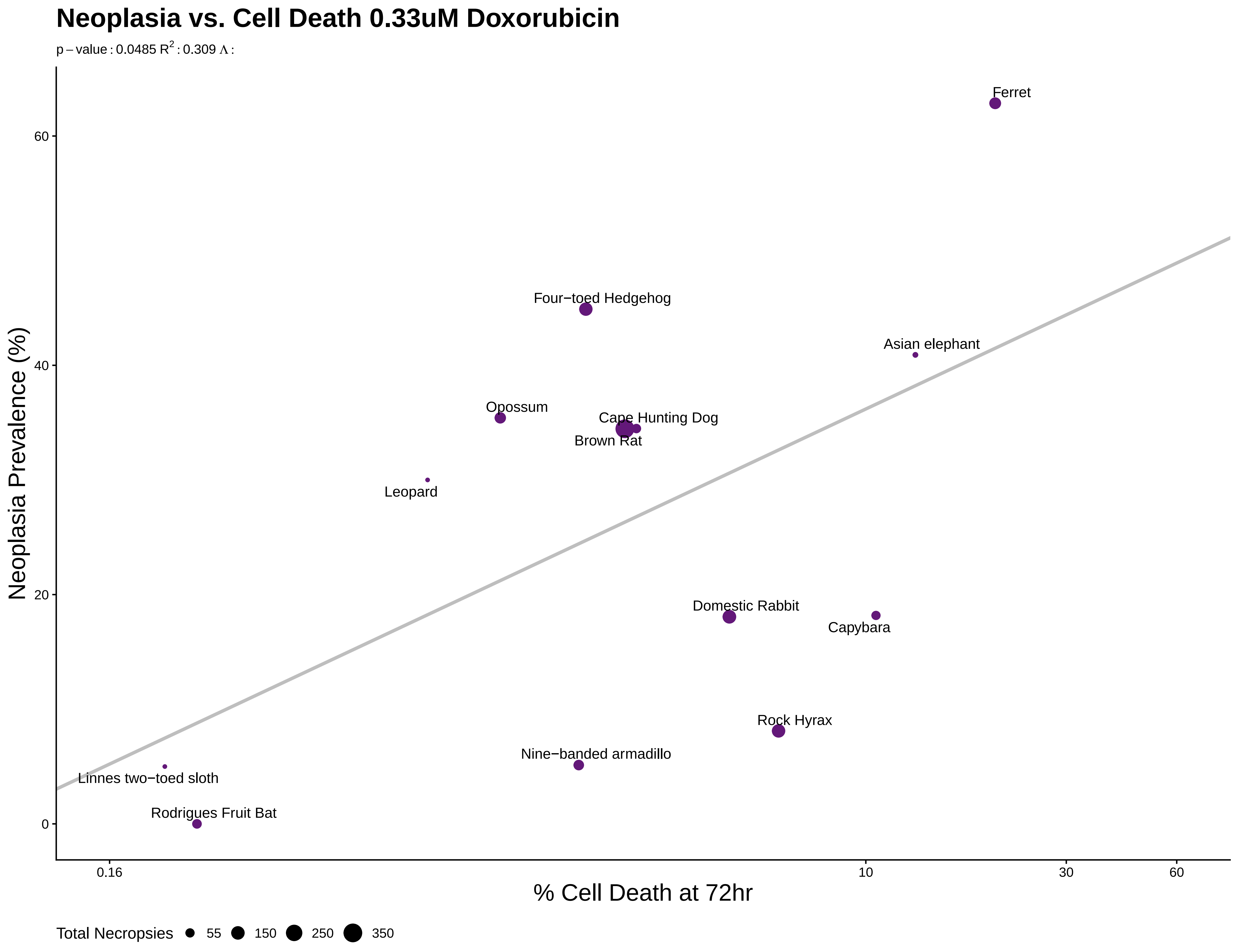

### S56celldeath72.11.pdf

#### Neoplasia vs. Cell Death 0.11uM Doxorubicin

p – value : 0.0503 $R^2$ : 0.305 Λ :

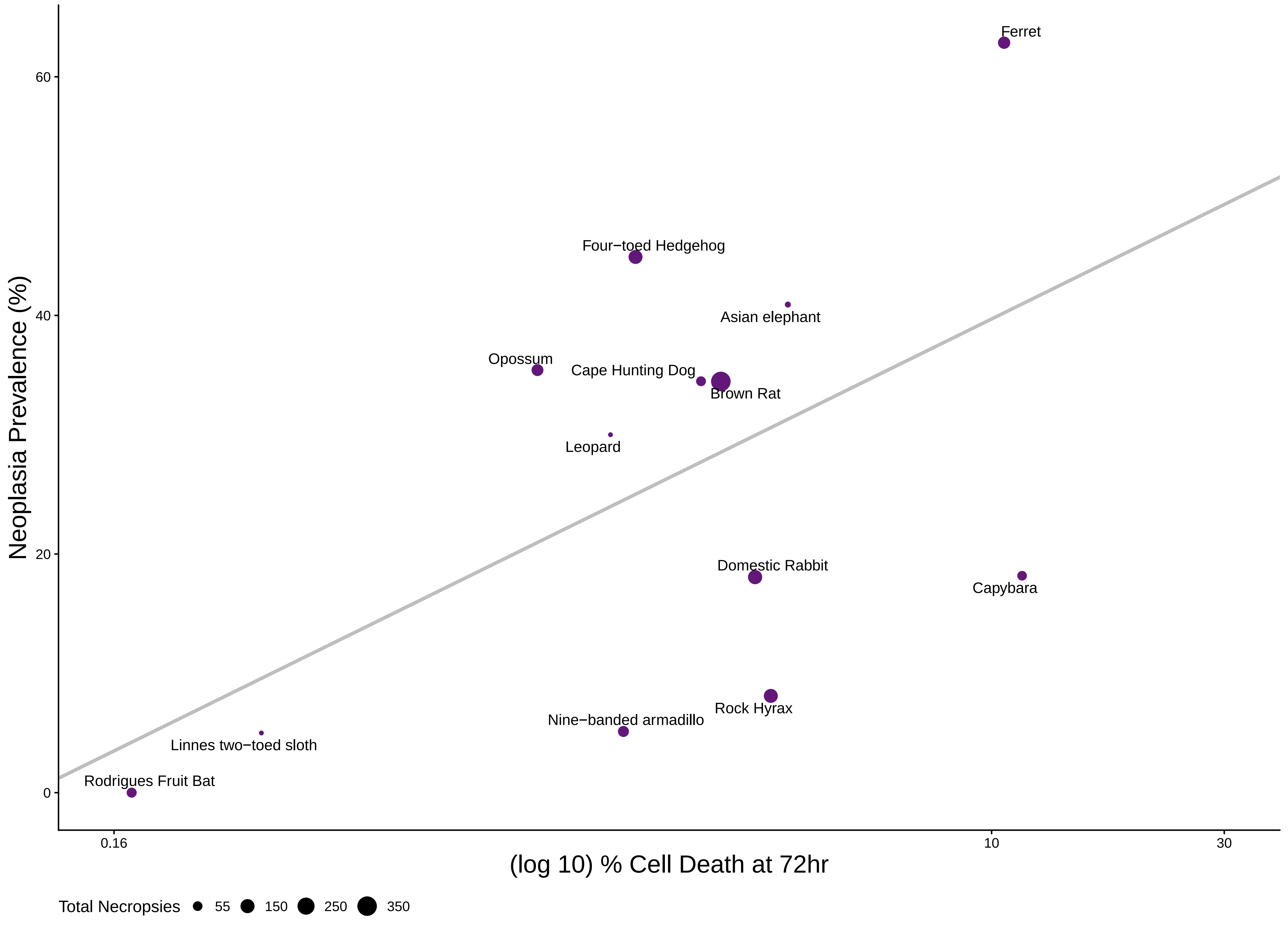

### S57mutation.mal.png

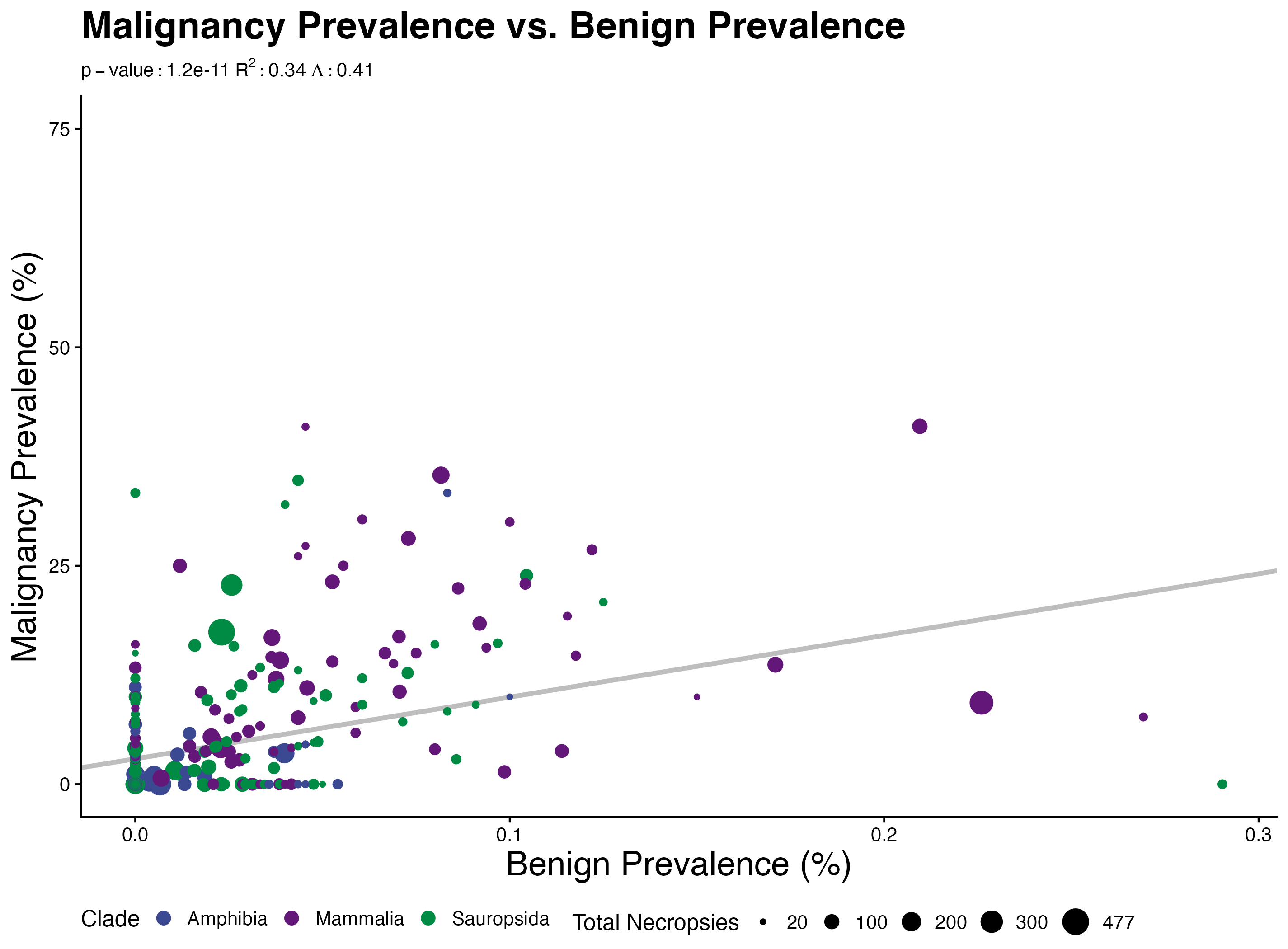

### S58correlationmatrix.jpeg

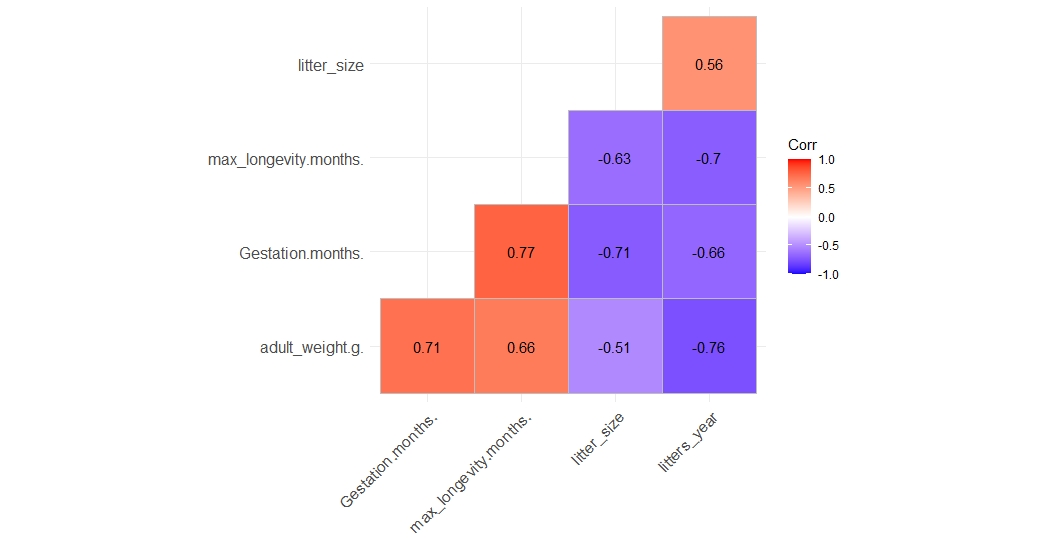

### S61nametree.pdf

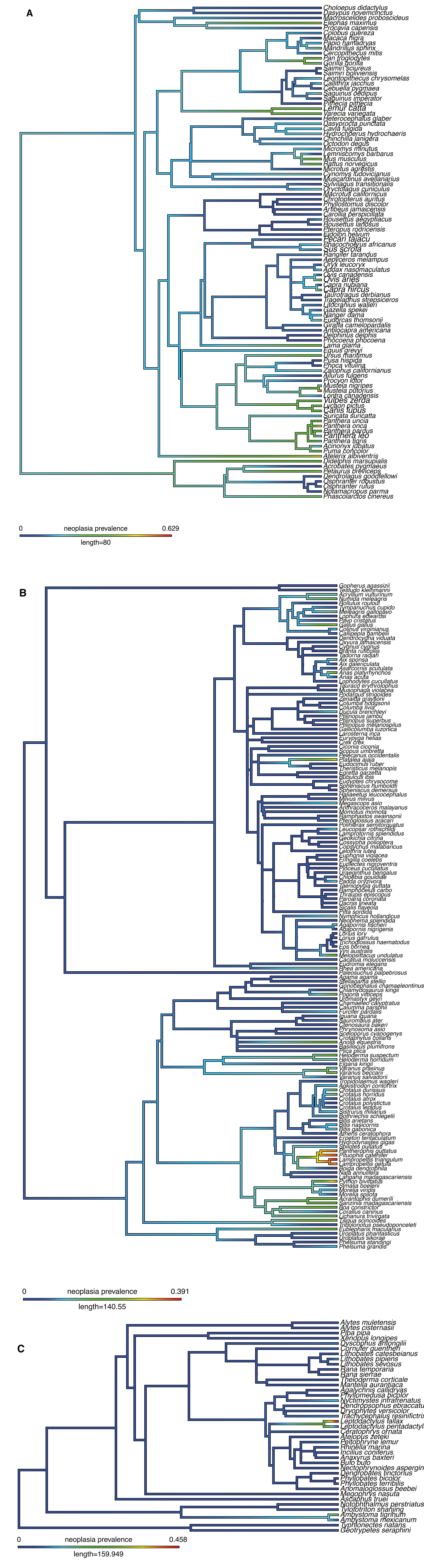

### S62wgtpropl.pdf

#### Prop Malignant vs. BMR

$p - value : 0.69$ $R^2 : 0.027$ $\Lambda : 0.1$

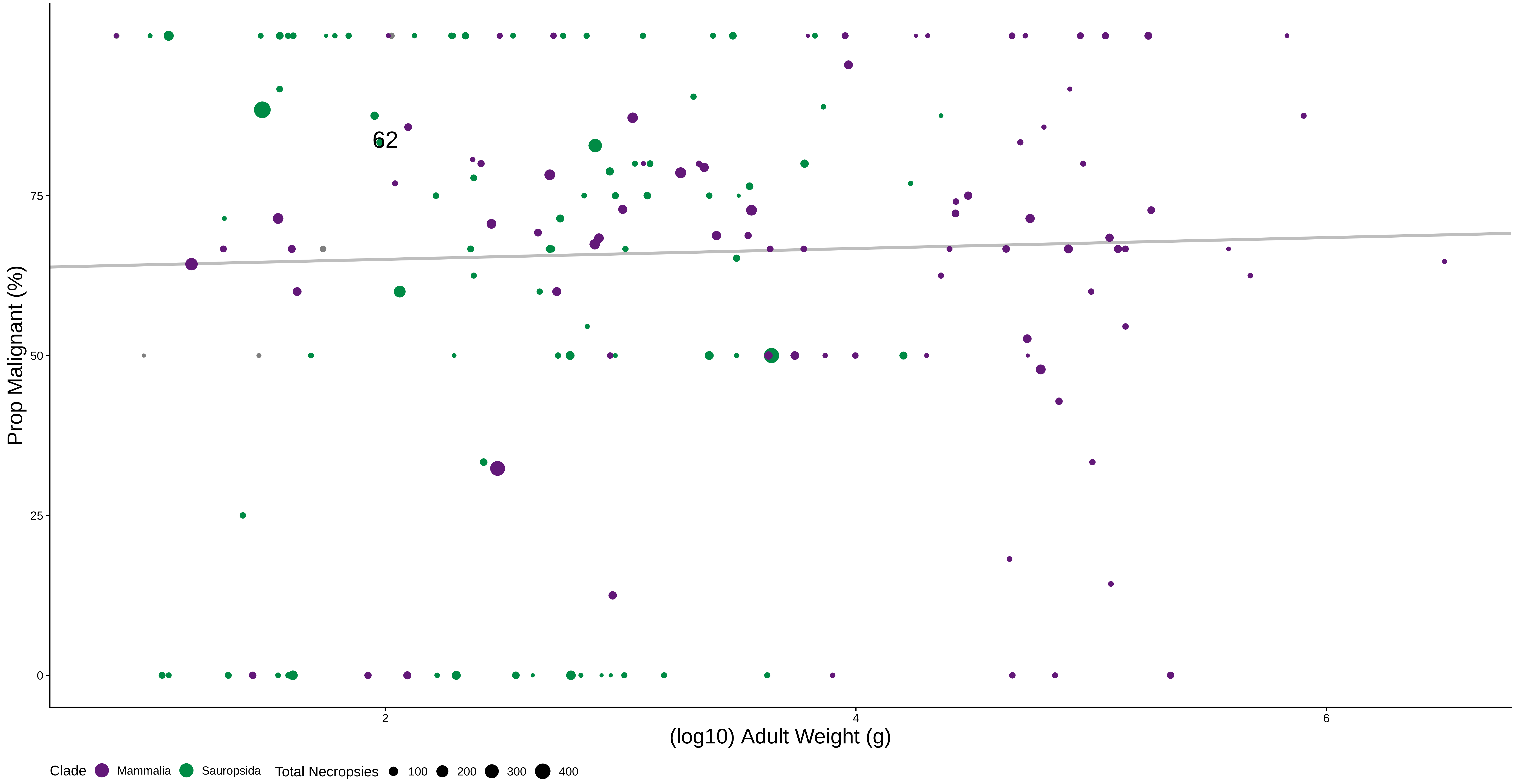

### S64gestprop.pdf

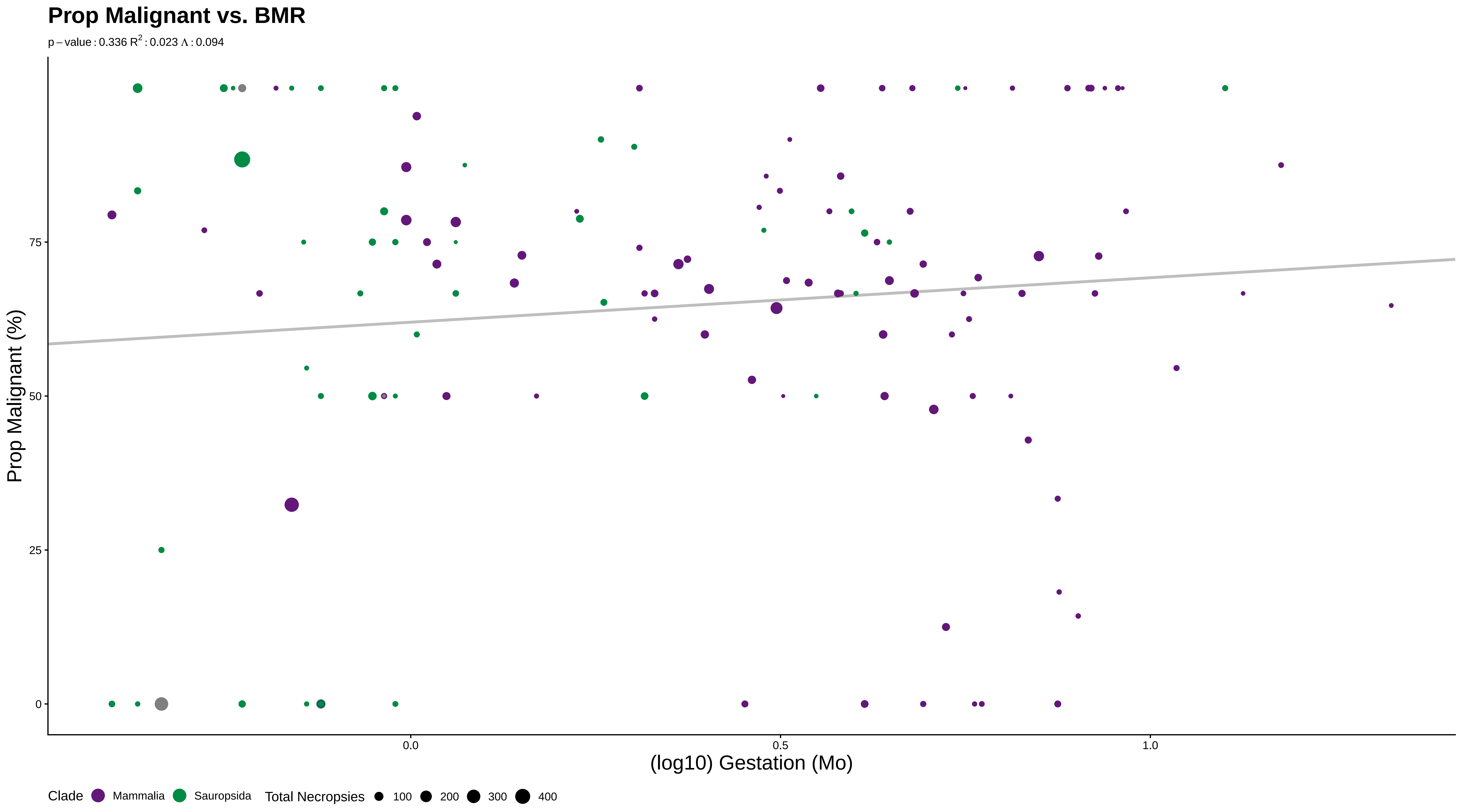

### S65litprop.pdf

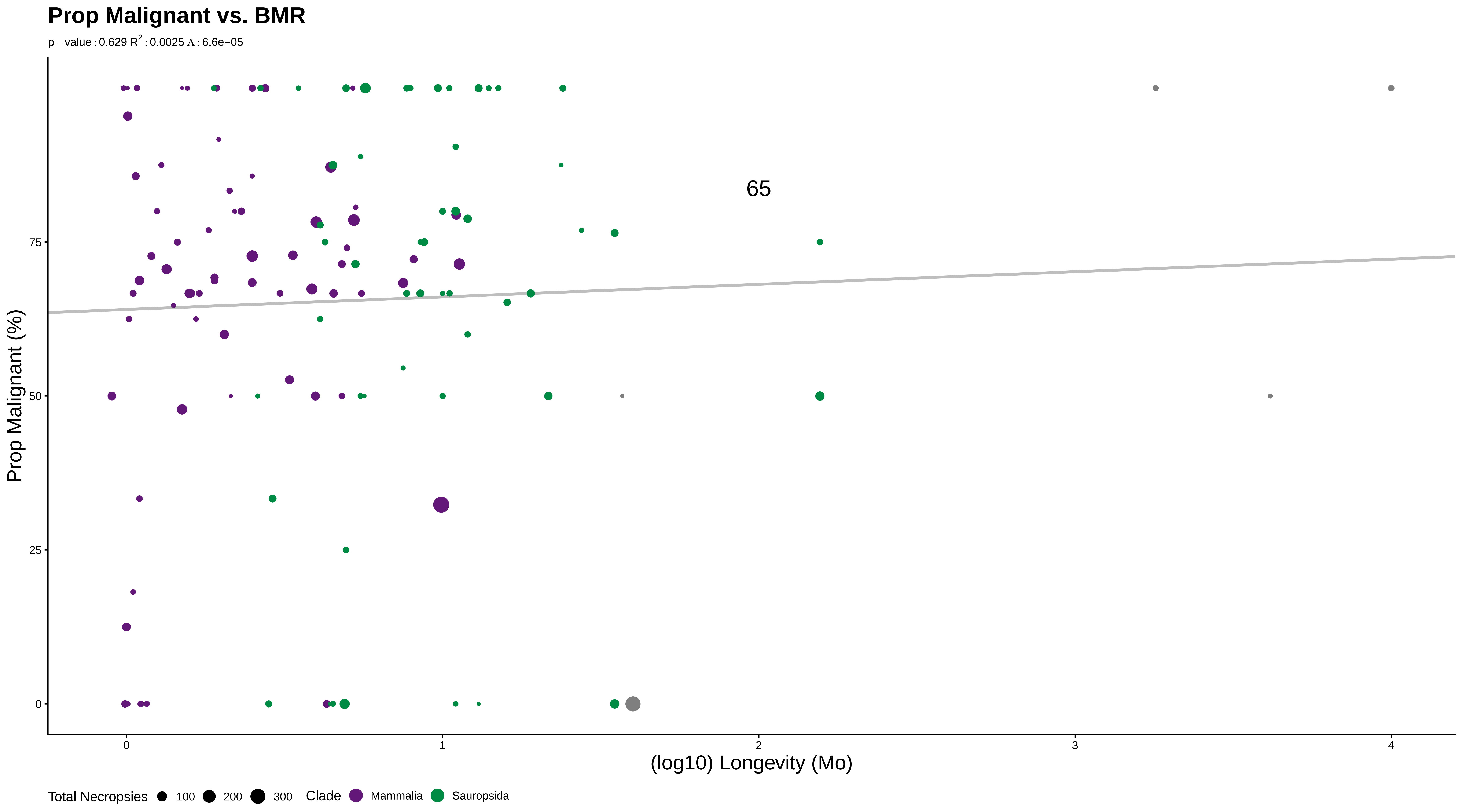

### S66longprop.pdf

##### Prop Malignant vs. BMR

p – value : 0.614 $R^2$ : 0.022 $\Lambda$ : 0.2

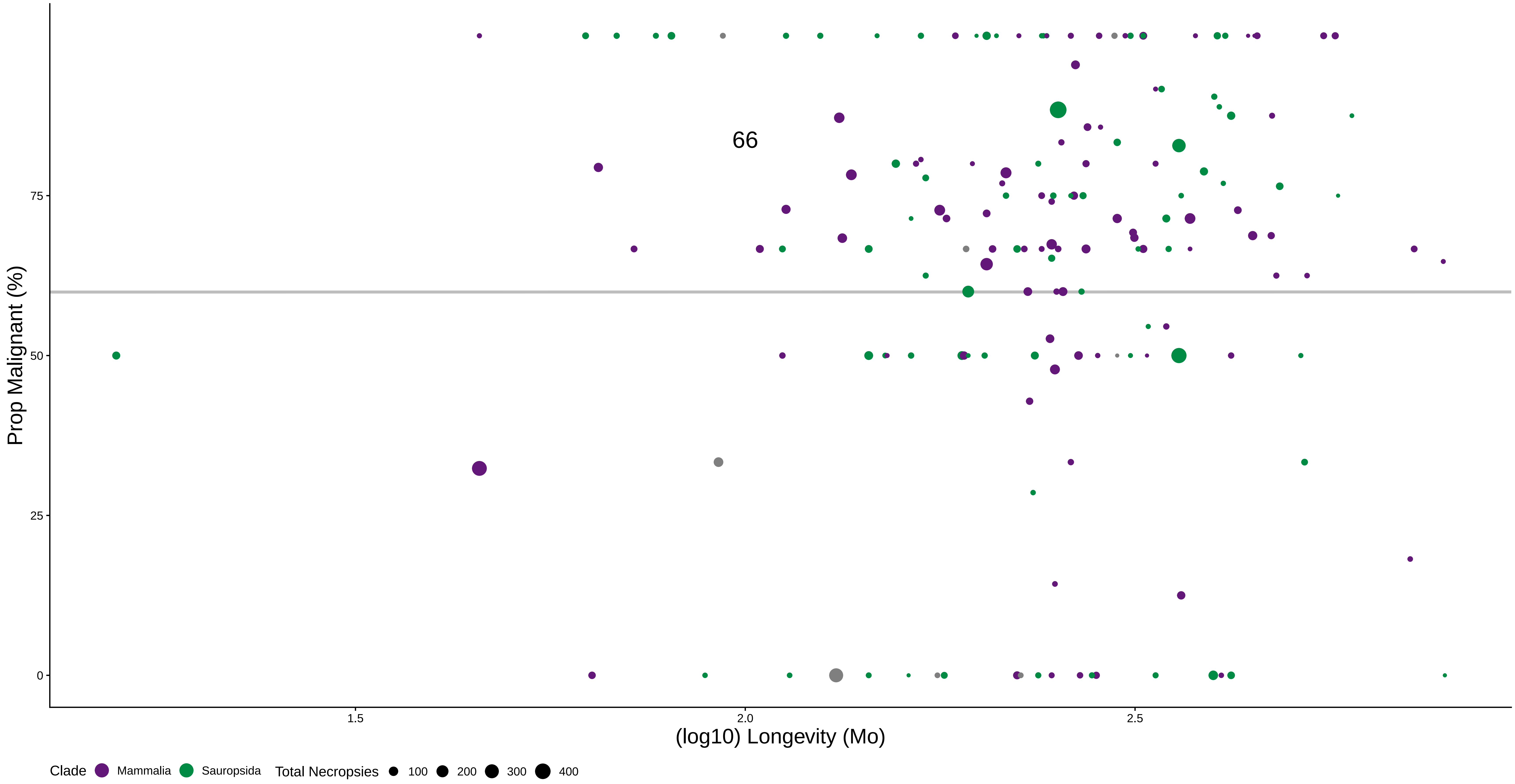

### S67bmrprop.pdf

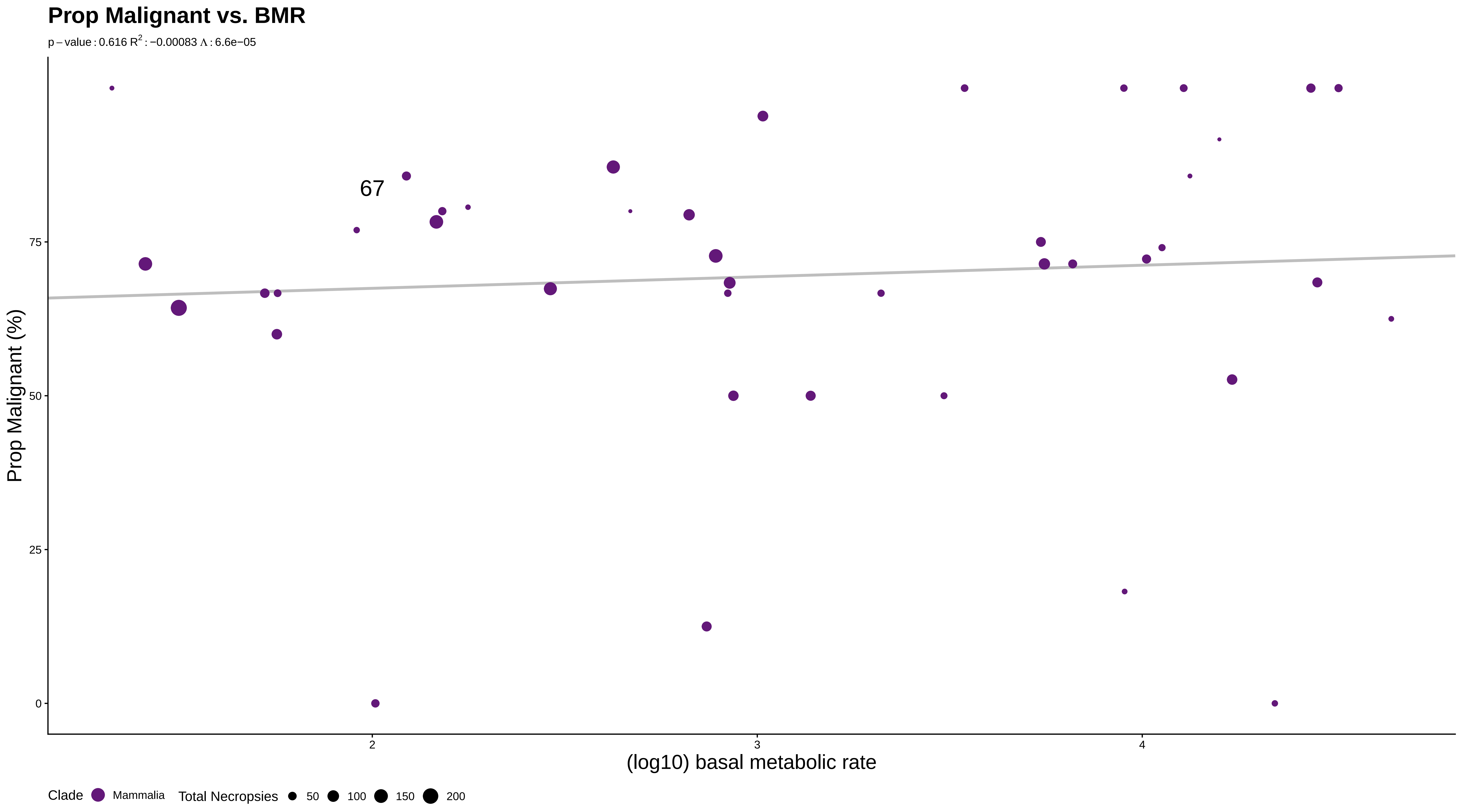

### S68wgtlongprop.pdf

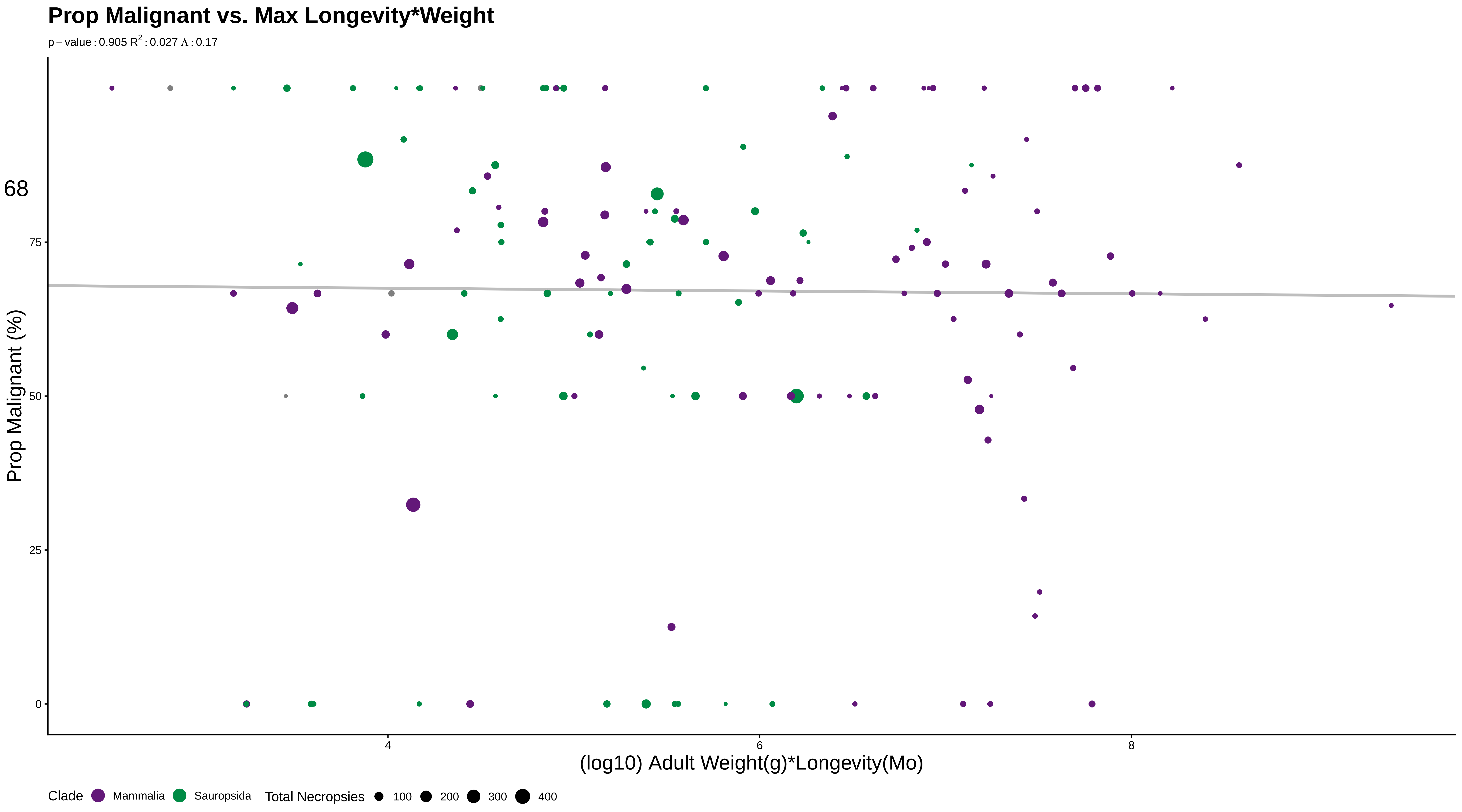

### S69lityearprop.pdf

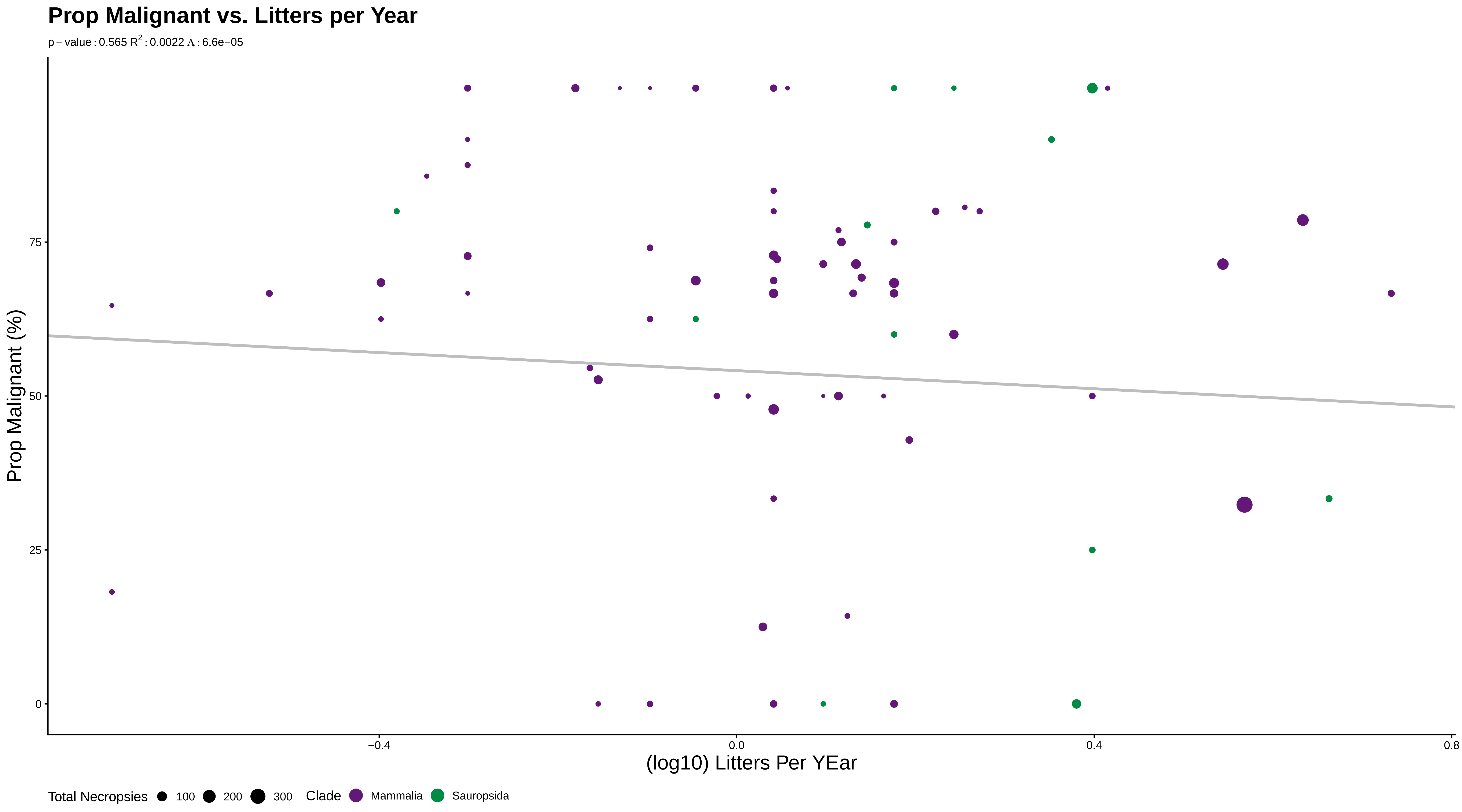

### S70femmatprop.pdf

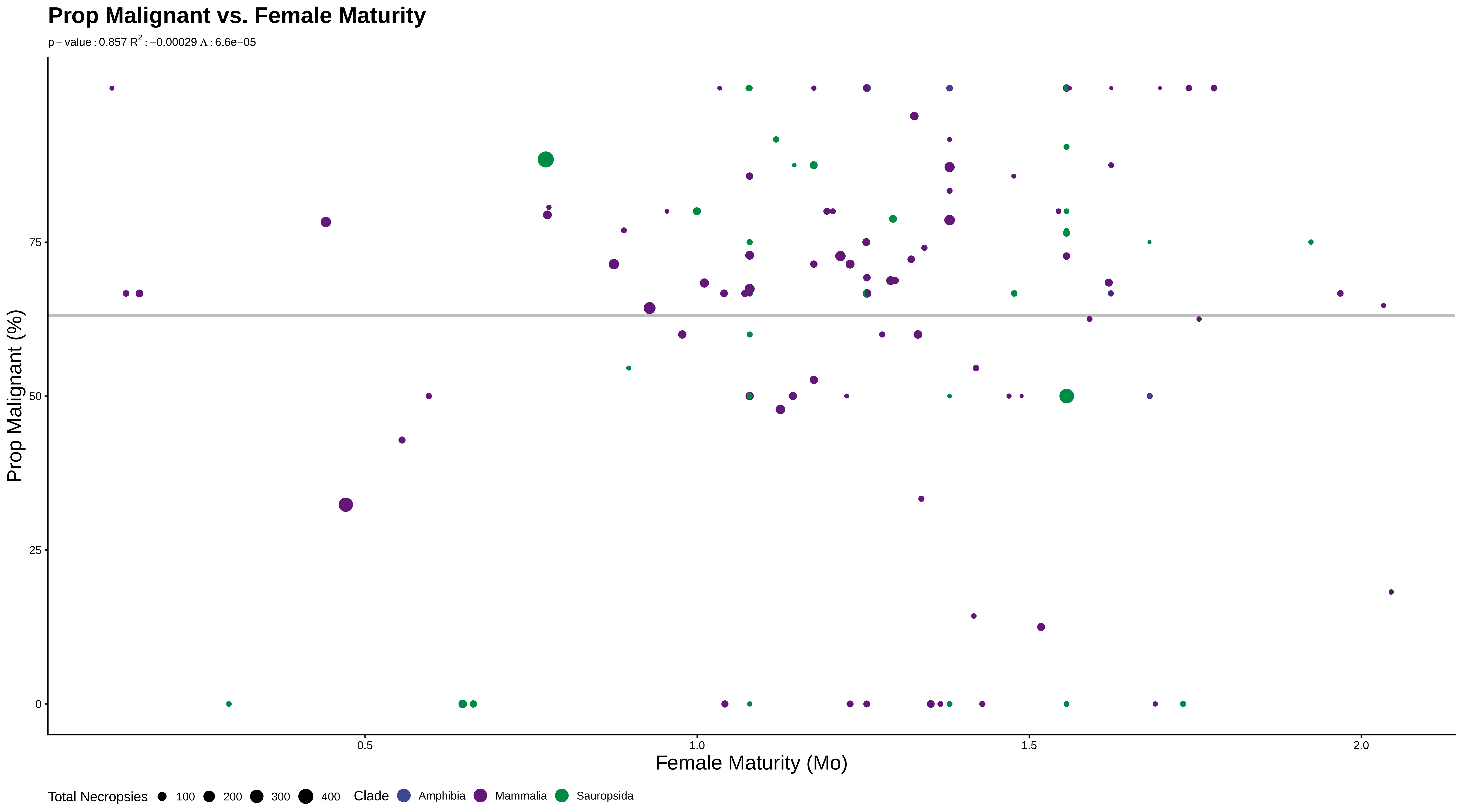

### S71malematprop.pdf

#### Prop Malignant vs. Male Maturity

p – value : 0.872 $R^2$ : 0.00051 $\Lambda$ : 6.4e−05

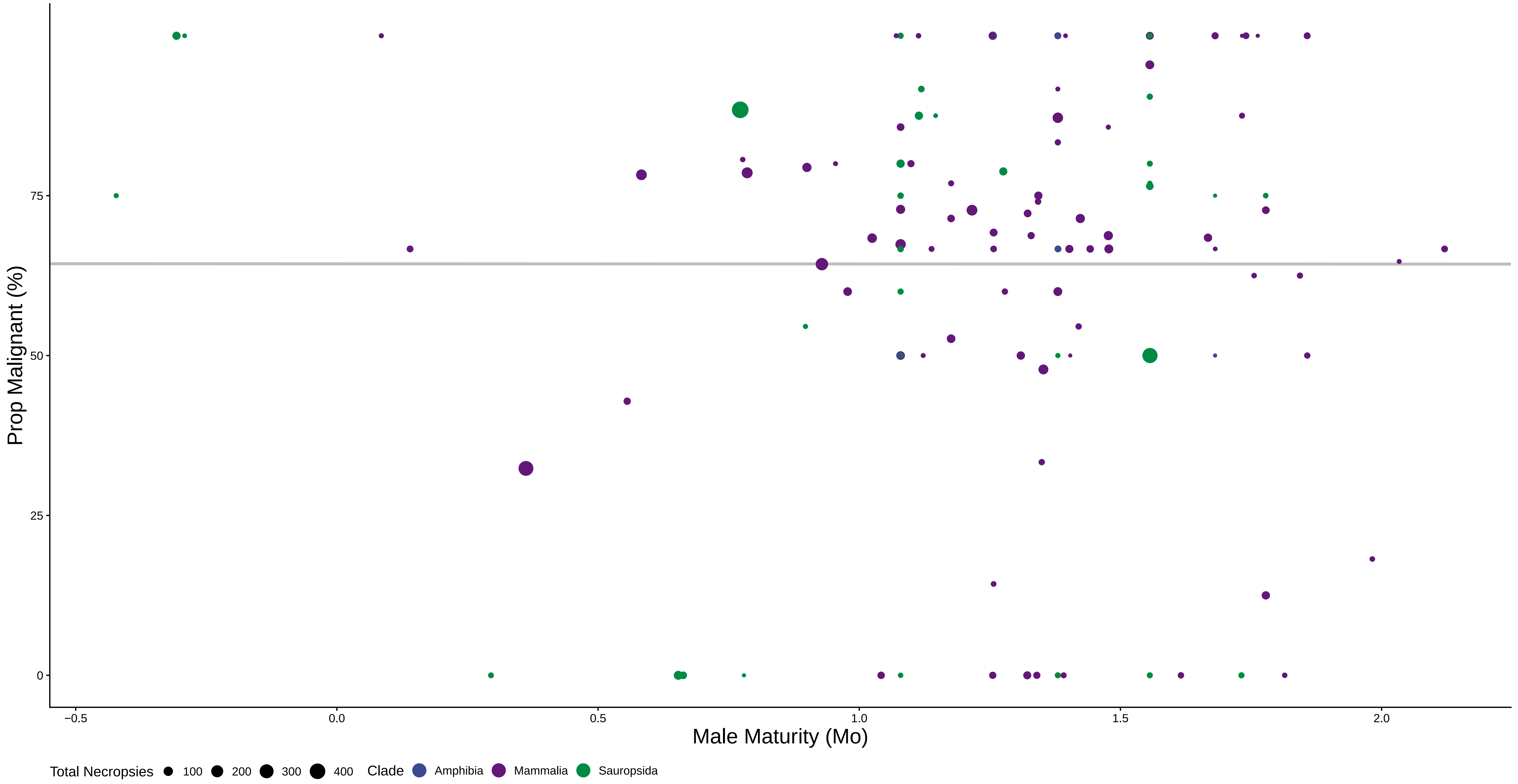

### S72weanprop.pdf

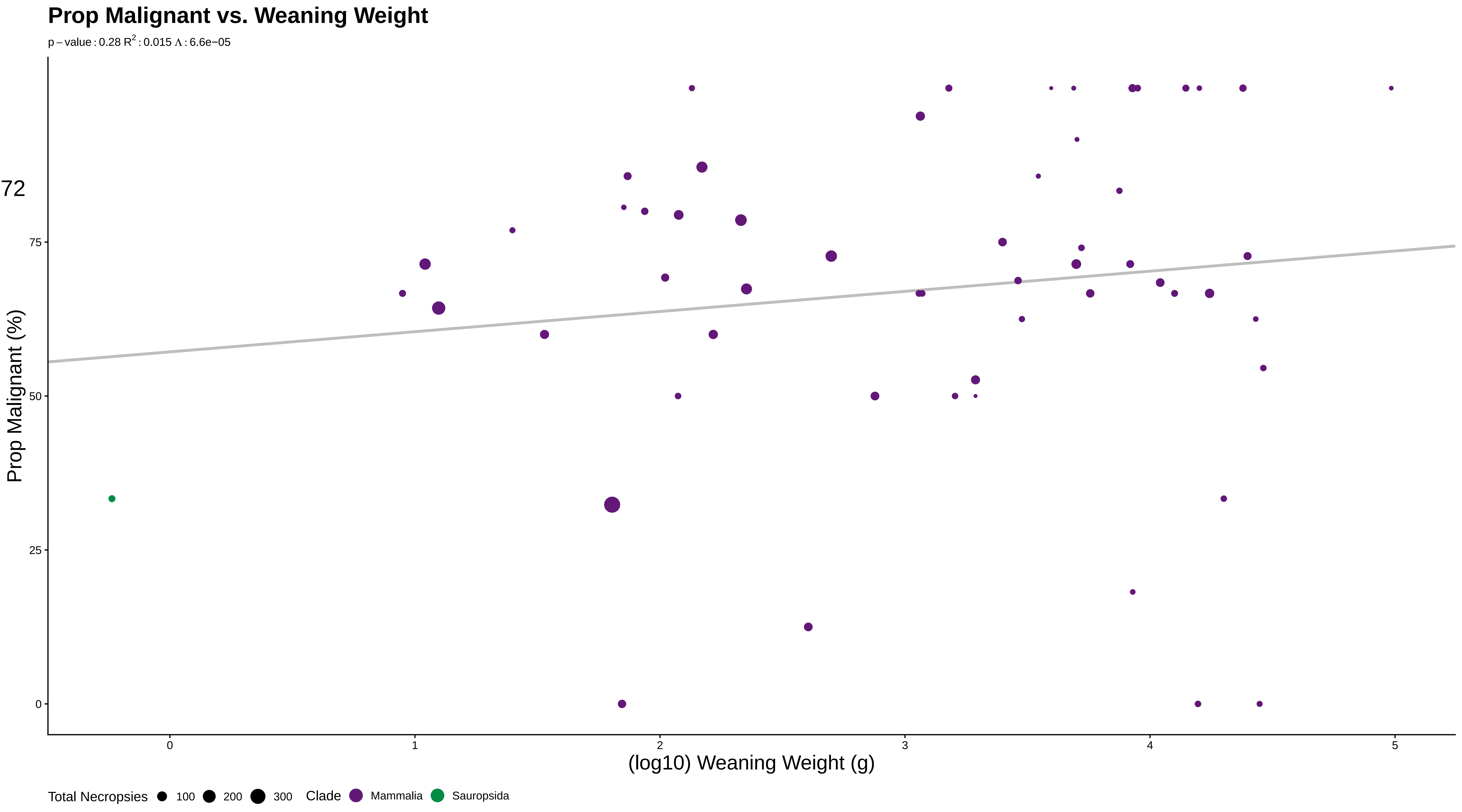

### S73growprop.pdf

#### Prop Malignant vs. Growth Rate

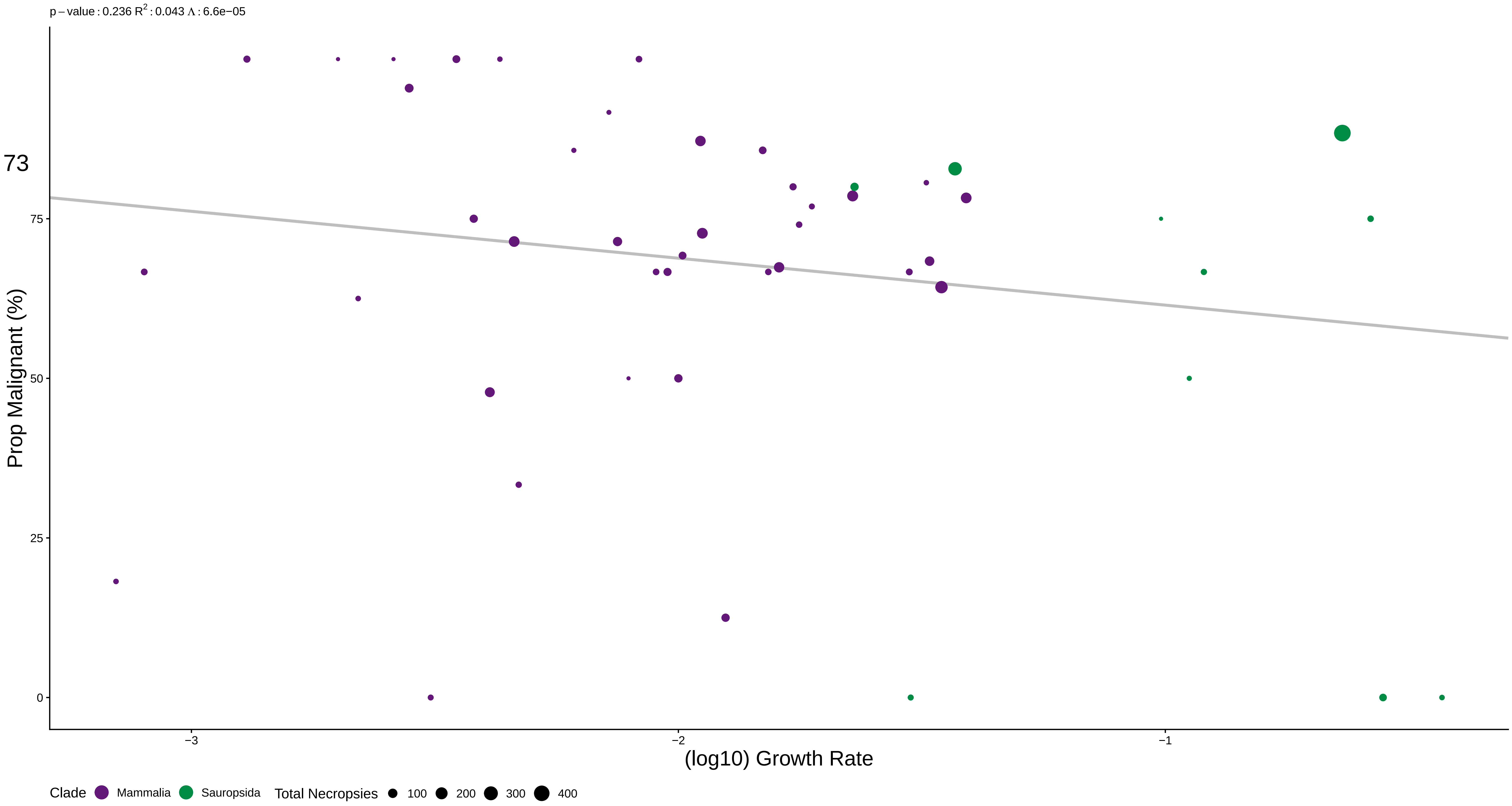

### S74wgtgestprop.pdf

#### Prop Malignant vs. Max Longevity+Gestation

p – value : 0.5846613 $R^2$ : 0.016 Λ : 6.6e−05

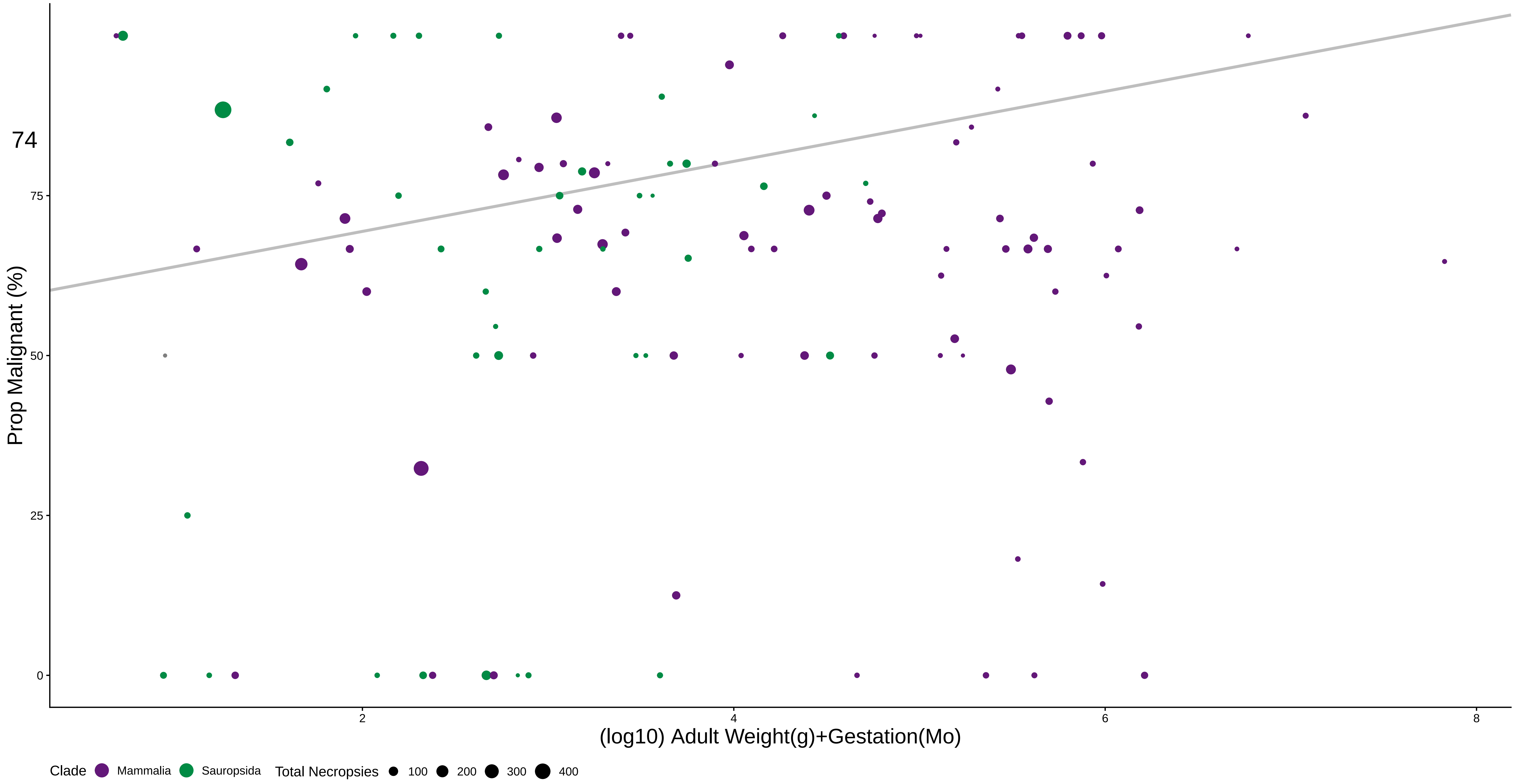

### sqViolin.pdf

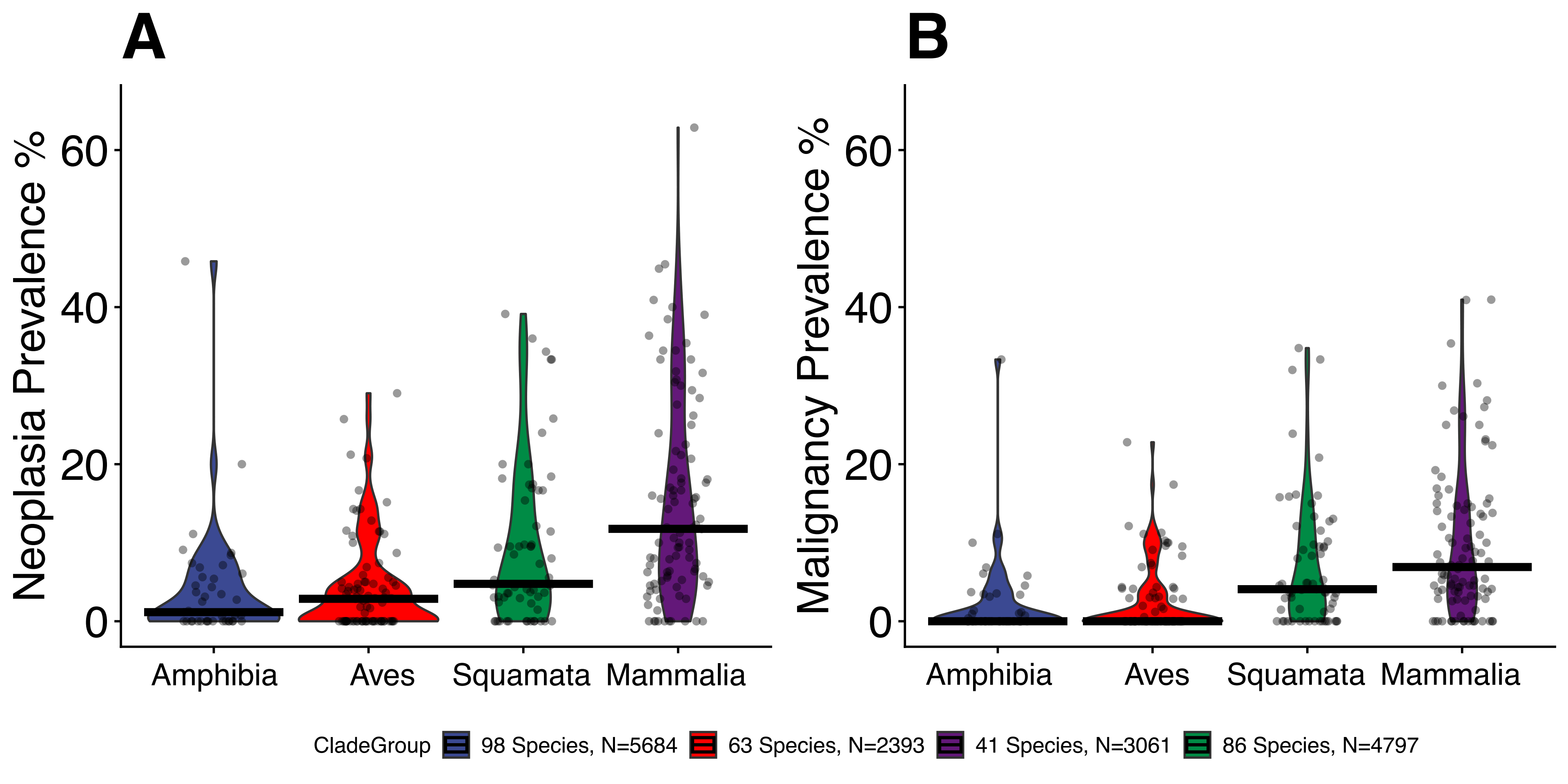
